# Long-read shotgun metagenomics identifies distinct urogenital microbial community states associated with functional potential and urinary tract infection during pregnancy

**DOI:** 10.64898/2026.11.04.762207

**Authors:** Melline Fontes Noronha, Ben Ferneyhough, Matthew J. Roddis, Nick J. Parkinson, Alan J Wolfe

## Abstract

Urinary tract infections (UTIs) are among the most common bacterial infections during pregnancy and are associated with adverse maternal and neonatal outcomes. Although previous studies used enhanced culture, 16S rRNA gene sequencing, and/or short-read shotgun metagenomic sequencing to characterize the urogenital microbiome of adult females, few have assessed the urogenital microbiome during pregnancy and thus this microbiome remains poorly understood. Here, we applied Oxford Nanopore long-read shotgun metagenomic sequencing to characterize urogenital microbial composition, functional potential, metagenome-assembled genomes (MAGs), and the urogenital virome of pregnant females.

Taxonomic profiling identified three distinct microbial community states predominated by *Lactobacillus crispatus*, *Gardnerella* species, and *L. iners*. The *Gardnerella*-predominated community was associated with a higher frequency of UTI during pregnancy. Functional profiles strongly reflected these community states, demonstrating that taxonomic differences were accompanied by distinct functional potential.

Genome-resolved analysis recovered 415 MAGs, including 29 high-quality genomes, revealing substantial genomic diversity among urogenital microbes. Viral sequences, predominantly bacteriophages, also were detected in most samples, highlighting an additional component of the pregnancy-associated urogenital microbiome.

These findings demonstrate that pregnant females possess distinct urogenital microbial community states and show that they differ not only in taxonomic composition but also in their functional potential.

## INTRODUCTION

Urinary tract infections (UTIs) are among the most common bacterial infections during pregnancy and are associated with important maternal and neonatal complications, including pyelonephritis, preterm delivery, and low birth weight. Asymptomatic bacteriuria (ASB), defined as the presence of significant bacteriuria in the absence of urinary symptoms, is estimated to occur in approximately 2–10% of pregnant individuals. Because untreated ASB can progress to infections of the bladder (cystitis) and sometimes the kidney (pyelonephritis), screening by urine culture during early prenatal care is recommended (US Preventive Services Task Force et al., 2019; American College of Obstetricians and Gynecologists’ Committee on Clinical Consensus–Obstetrics, 2023). However, conventional urine culture captures only a limited fraction of the microbes present within the female urogenital tract (Brubaker et al., 2023; Moreland et al., 2025).

Historically, the urinary tract was considered sterile in the absence of clinical infection. However, recent studies challenge this paradigm, using enhanced culture methods, 16S rRNA gene sequencing, and/or short-read shotgun metagenomic sequencing to demonstrate the presence of diverse bacterial communities in the female urinary tract (Wolfe et al., 2012; Hilt et al., 2014; Pearce et al., 2014; Giron-Villalobos et al., 2026). Some studies used enhanced culture methods and/or 16S rRNA gene sequencing to identify bacterial communities in asymptomatic pregnant female urine, most commonly finding the genera *Lactobacillus* and/or *Gardnerella* (Ollberding et al., 2016; Koedooder et al., 2021; Sung et al., 2024; Mora-Vargas et al., 2025). One study compared first-trimester midstream voided urine and vaginal swabs, showing that these microbial communities are related but distinct (Sung et al., 2024). Thus, bacterial detection alone does not imply infection, emphasizing the need to examine the broader microbial community in urinary health and disease.

An important consideration in studies of pregnant females is the method of specimen collection. It is unethical to catheterize a pregnant female without clinical cause and thus samples must be obtained by the midstream voided urine method. Microbial profiles obtained from voided urine are not equivalent to those obtained directly from the bladder by catheterization. During voiding, urine may acquire microbes from the urethra, vulvovagina, and surrounding skin. Accordingly, consensus recommendations for urobiome research propose the term “urogenital sample” for voided urine specimens, reserving terminology specific to the urinary bladder for samples collected directly from the bladder (Brubaker et al., 2021). Thus, microbial communities detected in voided urine should be interpreted as representing the broader urogenital microbial environment rather than the bladder microbiome exclusively.

Studies of the pregnancy urogenital microbiome have relied predominantly on enhanced culture and 16S rRNA gene sequencing (Sung et al., 2024). Although these approaches provide important information regarding bacterial community composition, they offer limited resolution of microbial genomic diversity and provide little direct information about microbial community functional potential. Consequently, it remains unclear whether distinct urogenital microbial community states during pregnancy also differ in their functional capabilities, which microbial genomes contribute to these functional differences, and whether these community-level features are associated with UTI and other clinical characteristics. Viral communities within the pregnancy-associated urogenital environment also remain understudied.

Shotgun metagenomic sequencing provides an opportunity to move beyond taxonomic profiling by simultaneously characterizing microbial composition, gene content, and functional potential, while enabling the reconstruction of metagenome-assembled genomes (MAGs). Long-read sequencing further provides improved genomic contiguity and can facilitate genome reconstruction and species- and strain-level characterization within complex microbial communities (Kim et al., 2024). Together, these approaches provide a framework for determining not only which microbes are present, but also how their genomic potentials differ across microbial community states.

Here, we used Oxford Nanopore long-read shotgun metagenomic sequencing to characterize urogenital microbial communities in midstream voided urine samples collected from pregnant females. We investigated microbial community structure, reconstructed microbial genomes, characterized functional potential, examined viral communities, and evaluated associations with clinical metadata, including UTI during pregnancy. By integrating taxonomic, genome-resolved, functional, and viral analyses, we sought to determine whether distinct urogenital microbial community states during pregnancy are accompanied by differences in genomic potential and whether these community characteristics are associated with clinically relevant outcomes.

## MATERIALS AND METHODS

### Study Population

This study represents a secondary analysis of samples and clinical data collected as part of a previously published study (Dhillon et al., 2026). The study protocol was approved by the Institutional Review Board [LU 216104] and registered on ClinicalTrials.gov (NCT 05554081). Patients were eligible at their initial prenatal visit if they were age >18 years, gestational age <20 weeks, and preferred English for medical communication. Patients with a history of recurrent UTI or ongoing antibiotic use were excluded. All participants provided informed consent in accordance with institutional guidelines. Out of the total number of participants in the original study, a total of 139 samples had sufficient remaining volume to permit metagenomic sequencing. One sample returned insufficient sequence reads and was excluded from further analysis; thus, 138 samples were included in the final analysis.

Participant demographic and clinical characteristics are summarized in **Supplemental Table 1**. Not all participants reported these characteristics; thus, the values for each characteristic are calculated only for those participants who provided information concerning that characteristic. UTI during pregnancy was defined as urogenital symptoms with a standard urine culture showing ³10^5^ colony forming units per milliliter of a bacterial species considered to be a uropathogen.

### Genome extraction, library preparation, and long-read shotgun metagenomic sequencing

For each sample, 2 mL of frozen urine was used as the input. Samples were thawed and warmed (37 °C, 30 min) to prevent co-precipitation of lipids and proteins, and cells were recovered by centrifugation (14,000 × g, 5 min). Host cells were selectively lysed and free host DNA removed by enzymatic host depletion (ZymoBIOMICS HostZero chemistry: host-depletion solution, followed by microbial selection buffer with nuclease digestion at 37 °C and subsequent proteinase K inactivation). The retained microbial fraction was spiked with a fixed titer of an internal calibration standard (ZymoBIOMICS Spike-in Control) to enable absolute quantification, then lysed by heating (95 °C, 5 min) and mechanical bead-beating (FastPrep, MP Biomedicals), and sheared by sonication (Bioruptor, Diagenode). Microbial DNA was captured on a silica spin column, washed, eluted in nuclease-free water, and quantified by fluorometry (Quant-iT PicoGreen/AccuBlue, read on a BMG PHERAStar against a DNA standard curve). Sample tracking, reagent lot control, and yield calculation were managed within the laboratory information management system (LIMS).

Sequencing libraries were prepared from extracted DNA (target ∼100 ng; maximum 400 ng input) using the Oxford Nanopore ligation-based native barcoding kit (SQK-NBD114.24). DNA was end-repaired and A-tailed (NEBNext FFPE DNA Repair and NEBNext Ultra II End Prep), purified with AMPure XP beads (Beckman Coulter), ligated to native barcodes and then to sequencing adapters, with bead-based clean-up and size selection between steps. Barcoded libraries were quantified, pooled, and eluted ready for loading. Library preparation was performed manually or by an equivalent semi-automated protocol on a Tecan Fluent liquid-handling workstation.

Pooled libraries were loaded onto R10.4.1 flow cells and sequenced on a GridION device (Oxford Nanopore Technologies) under MinKNOW control, typically as overnight runs (∼14 h). Basecalling was performed with the current Oxford Nanopore basecaller (Dorado).

### Sequence Quality Control and Preprocessing

Raw Oxford Nanopore sequencing reads were assessed using NanoStat v1.6.0 (De Coster et al., 2018) to characterize sequencing yield, read-length distributions, and read-quality metrics. Reads were subsequently quality-filtered using NanoFilt v2.8.0 (De Coster et al., 2018), applying a minimum Phred quality score threshold of Q≥12. No length-based filtering was applied. Sequencing and quality-control metrics were summarized across samples before and after quality filtering.

### Removal of Non-target Sequences

To identify and remove non-target sequences, quality-filtered reads were sequentially aligned against the reference genomes of the sequencing kit-associated bacterial contaminants *Allobacillus halotolerans* strain BCRC 17939 (RefSeq accession GCF_028743495.1) and *Imtechella halotolerans* strain JCM 17677 (RefSeq accession GCF_028743515.2), followed by alignment against the human reference genome GRCh38 (RefSeq accession GCF_000001405.39) to remove host-derived reads, as dictated by our approved IRB protocol.

Alignments were performed using minimap2 v2.24 (r1122) (Li, 2018) with the Oxford Nanopore preset (-x map-ont) and secondary alignments disabled (--secondary=no). Resulting alignments were sorted using SAMtools (Danecek et al., 2021), and unmapped reads were extracted using the SAM flag -f 4 and converted back to FASTQ format. Reads remaining after sequential removal of spike-in-associated and human-derived sequences were retained for downstream taxonomic and functional analyses.

### Taxonomic Profiling

Species-level taxonomic profiles generated from shotgun metagenomic sequencing were obtained using Kraken2 (Wood et al., 2019) for taxonomic classification followed by Bracken (Lu et al., 2025) for abundance re-estimation. Relative abundance tables were imported into R version 4.2.1 (R Core Team, 2022) for downstream community analyses.

### Microbial community clustering

To identify major microbial community structures, Bray–Curtis dissimilarity matrices were calculated from species-level relative abundance profiles using the vegan package (Oksanen et al., 2022). The optimal number of microbial community clusters was initially evaluated using partitioning around medoids (PAM) (Kaufman and Rousseeuw, 1990) and average silhouette width (Rousseeuw, 1987). Candidate solutions were further assessed based on cluster separation, sample distribution, and biological interpretability. Hierarchical clustering was subsequently performed using Ward’s minimum variance method (Ward.D2), and the dendrogram was partitioned into the selected number of clusters using the cutree() function. The resulting three-cluster solution was used for all downstream analyses.

### Clinical metadata association analysis

Boruta feature selection (Kursa and Rudnicki, 2010) was used to identify demographic and clinical variables associated with microbial community cluster membership. Variables identified as important by Boruta were subsequently evaluated for association with cluster membership using Fisher’s exact test for categorical variables.

### Alpha diversity analysis

Within-sample microbial diversity was assessed using Observed richness, ACE, Chao1, Shannon, Simpson, inverse Simpson, and Pielou’s evenness calculated from species-level profiles using the *phyloseq* package (McMurdie and Holmes, 2013). Pielou’s evenness was calculated as Shannon diversity divided by the natural logarithm of Observed richness. Differences among the three microbiome community state types were assessed using the Kruskal–Wallis test.

### Beta diversity analysis

Between-sample microbial community differences were evaluated using Bray–Curtis dissimilarity calculated from species-level relative abundance profiles. Principal coordinates analysis (PCoA) was used to visualize patterns of community composition. Statistical differences among microbial community clusters were assessed using PERMANOVA (adonis2), and homogeneity of multivariate dispersion was evaluated using PERMDISP (betadisper), as implemented in the vegan package (Oksanen et al., 2022).

### Differential abundance analysis

Differential abundance analysis was performed at the species level using ANCOM-BC2 (Analysis of Composition of Microbiomes with Bias Correction, version 2), implemented in the ANCOMBC R package (Lin and Peddada, 2020). Pairwise comparisons were conducted among the three microbial community clusters. For each comparison, the phyloseq object was subset to the two clusters of interest, and cluster membership was included as the fixed effect in the ANCOM-BC2 model. Species with a prevalence below 5% were excluded prior to analysis. Bias correction, structural zero detection, and negative lower-bound adjustment were enabled, and p values were adjusted for multiple testing using the Benjamini–Hochberg false discovery rate (FDR) procedure. Species with an FDR-adjusted q value <0.05 were considered differentially abundant.

### Functional annotation

Functional annotation of quality-filtered metagenomic reads was performed using eggNOG-mapper v2.1.13 (Cantalapiedra et al., 2021), with sequence similarity searches conducted using DIAMOND (Buchfink et al., 2021) in sensitive mode against the eggNOG protein database. Functional annotations included KEGG Orthologs (KOs) and pathways (Kanehisa et al., 2025), Enzyme Commission (EC) numbers, Gene Ontology (GO) terms, Clusters of Orthologous Groups (COGs), Pfam protein domains, CAZy families, and BiGG reactions. Sample-specific annotation files were merged to generate abundance matrices for each functional category and subsequently imported into R as individual *phyloseq* objects (McMurdie and Holmes, 2013).

### Functional comparative analysis

KO and KEGG pathway profiles were filtered to retain features detected in at least five samples with a minimum total count of 10 across the dataset. For beta-diversity analyses, counts were transformed to relative abundances, and Bray–Curtis dissimilarities were calculated. PCoA was used to visualize differences in functional composition among the previously defined *L. crispatus*-, *Gardnerella*-, and *L. iners*-predominated microbiome communities. Differences among community state types were assessed using PERMANOVA with 9,999 permutations, and homogeneity of multivariate dispersion was evaluated using PERMDISP with 9,999 permutations (Anderson, 2001, 2006).

### Clinical associations with functional composition

Associations between clinical and demographic variables and pathway-level functional composition were evaluated independently using PERMANOVA with 4,999 permutations. Homogeneity of multivariate dispersion was assessed using PERMDISP, and *P* values were adjusted for multiple testing using the Benjamini–Hochberg FDR procedure. For UTI during pregnancy, an additional Bray–Curtis PERMANOVA model including microbiome community state type followed by UTI was performed among samples with available UTI data (*n*=61) to evaluate the association between UTI and functional composition after accounting for microbiome community structure.

### Differential abundance analysis of KEGG Orthology (KOs) identifiers and KEGG pathways

DESeq2 (Love et al., 2014) was performed on filtered, untransformed count matrices, with size factors estimated using the poscounts method. One-versus-rest comparisons were performed for each microbiome community state. Features were considered significantly enriched at a Benjamini–Hochberg-adjusted *p*<0.05 and log_2_ fold change >1.

For KO visualization, the 20 most significantly enriched KOs from each community state were selected, and log_2_ fold changes across all three comparisons were displayed in a heatmap. Values were truncated at −4 and +4 for visualization only, while feature selection and statistical testing were based on the original values. KOs were annotated using eggNOG-derived KO identifiers and functional descriptions.

KEGG pathway profiles were generated by aggregating eggNOG-mapper annotations according to KEGG pathway identifiers (Kanehisa et al., 2025). Redundant KO and map representations sharing the same five-digit KEGG pathway identifier were collapsed into a single pathway representation. For visualization, all significantly enriched pathways from the *L. crispatus*- and *L. iners*-predominated communities and the 20 most significantly enriched pathways from the *Gardnerella*-predominated community were displayed in a heatmap. Log_2_ fold changes from all three comparisons were shown, with positive and negative values indicating relative enrichment and depletion, respectively.

### Taxon–functional pathway association analysis

Species-level taxonomic profiles were integrated with KO profiles across 138 samples. Taxa were retained if detected in ≥10% of samples with a mean relative abundance ≥0.1%, and KOs were retained if detected in ≥10% of samples. Taxonomic and KO count matrices were independently centered log-ratio (CLR) transformed (Aitchison, 1982) after addition of a pseudocount of 0.5. Pairwise taxon–KO associations were assessed using Spearman correlations, with significance defined as Benjamini–Hochberg-adjusted *p*<0.05 (Benjamini and Hochberg, 1995)and |ρ| ≥0.40.

Significantly associated KOs were mapped to KEGG pathways using eggNOG-mapper annotations. Taxon–pathway associations were retained when supported by ≥3 significant KOs and ≥70% directional concordance. Association strength was represented by the mean Spearman correlation among KOs supporting the predominant direction. For heatmap visualization, broad KEGG overview pathways were excluded, and the top 30 pathways were ranked by the number of associated taxa, supporting KOs, and mean absolute association strength.

For network analysis, a more stringent threshold of ≥15 supporting KOs and ≥70% directional concordance was applied. The resulting bipartite taxon–pathway network was visualized using the *igraph* (Csárdi G, Nepusz T, Traag V, Horvát Sz, Zanini F, Noom D, Müller K, 2026) and *ggraph* (Pedersen, 2025) R packages, with edge direction representing the predominant correlation direction and edge width representing the number of supporting KOs.

### Use of Artificial Intelligence

A locally deployed large language model (Llama 3.1 8B, via Ollama (Grattafiori et al., 2024)) was used in a two-stage AI-assisted interpretation workflow. An Evidence Agent first generated structured summaries of author-generated taxon–pathway and KO-level statistical results without biological interpretation. A Microbiome Biology Agent subsequently used these reports to assist with conservative biological interpretation while distinguishing study-supported observations, established biological knowledge, hypotheses, and limitations. The LLM was not used for primary data processing, statistical testing, or significance assessment. All AI-assisted outputs were reviewed by the authors, who retained responsibility for the final scientific interpretation and conclusions.

### Metagenome assembly and genome binning

Quality-filtered, decontaminated Oxford Nanopore reads were assembled independently for each sample using Flye v2.9-b1778 in metagenome mode (--meta) with --nano-hq (Kolmogorov et al., 2020). Metagenome assembled genomes (MAGs) were recovered using MetaBAT2 v2.15 (Kang et al., 2019) and evaluated with CheckM v1.2.4 using the lineage_wf workflow (Parks et al., 2015). MAGs with ≥90% completeness and ≤5% contamination were classified as high quality, whereas those with ≥50% but <90% completeness and ≤10% contamination were classified as medium quality.

### Genome dereplication and phylogenomics

MAGs with ≥50% completeness and ≤10% contamination were dereplicated using dRep v3.6.2 (Olm et al., 2017), with primary and secondary clustering at 90% and 99% average nucleotide identity (ANI), respectively, using FastANI (Jain et al., 2018). One representative genome per secondary cluster was retained for downstream analyses.

### MAG taxonomic assignment

Taxonomic assignments were determined for representative MAGs from each dRep secondary cluster by BLAST (Camacho et al., 2009) comparison against a curated bacterial reference genome database. Matches with ≥90% nucleotide identity and an alignment length ≥1,000 bp were retained, and the best-supported reference hit was selected based on bit score, sequence identity, alignment length, and E-value. Taxonomy of the best-matching reference genome was assigned to the corresponding dRep cluster and propagated to its member MAGs.

### Phylogenomic reconstruction

Using anvi’o v9 (Eren et al., 2020), bacterial single-copy marker genes from the Bacteria_71 HMM collection were identified, and the best hit for each marker was concatenated and aligned using MUSCLE (Edgar, 2004). A maximum-likelihood tree was inferred using FastTree (Price et al., 2010) and visualized in R using *ggtree (Yu et al., 2017)* and *ggtreeExtra (Xu et al., 2021)*.

### Comparative genomic analysis

Orthologous protein groups among 15 selected MAGs were identified using Proteinortho v6.3.6 (Lechner et al., 2011) with DIAMOND sequence similarity searches using an E-value threshold of 1 × 10⁻⁵, minimum sequence coverage of 50%, and minimum amino acid identity of 25%. Comparative genomic visualization was performed using anvi’o v9 (Eren et al., 2020), with detailed gene-content comparison focused on the six *Fannyhessea vaginae* MAGs.

### Putative antibiotic resistance-associated proteins

Protein-coding sequences predicted using Prokka v1.14.6 (Seemann, 2014) were screened for putative antibiotic resistance-associated proteins based on annotated gene product descriptions and predefined resistance-associated keywords. Proteins were grouped into resistance categories, and category counts across high-quality MAGs (*n* = 29) were visualized according to the phylogenomic tree using *ComplexHeatmap* (Gu et al., 2016).This analysis represents a screening of putative resistance-associated proteins based on Prokka functional annotations rather than dedicated antimicrobial resistance prediction; therefore, identified proteins were considered candidate resistance determinants rather than confirmed antibiotic resistance genes.

### Mobile genetic element-associated proteins

Prokka functional annotations from high-quality MAGs were screened for proteins associated with mobile genetic elements and related functions, including transposases, integrases/recombinases, phage-associated proteins, CRISPR/Cas systems, and toxin/antitoxin systems. Annotation counts were summarized across MAGs and used to compare the distribution of these functional categories among recovered genomes.

### Virome identification and analysis

Putative viral sequences were identified from assembled metagenomic contigs using geNomad v1.12.0 (Camargo et al., 2024). Contigs classified as viral were retained, whereas plasmid sequences were excluded, and viral taxonomy was assigned using the geNomad reference database. Viral richness was calculated as the number of viral contigs detected per sample and compared among the *L. crispatus, Gardnerella*-, and *L. iners*-predominated communities using the Kruskal–Wallis test followed by pairwise Wilcoxon rank-sum tests with Benjamini–Hochberg correction. Viral community composition was evaluated using Bray–Curtis dissimilarity and PCoA, with differences among community states assessed by PERMANOVA with 4,999 permutations. Associations between viral richness and clinical and demographic variables were evaluated using non-parametric tests with FDR correction.

### Data Availability

Raw sequencing data are available through the NCBI Sequence Read Archive under BioProject PRJNA1465356, “Detection methods of asymptomatic bacteriuria in an obstetric population.”

## RESULTS

### Participant Demographic and Clinical Characteristics

The participants were predominantly White (40.8%), followed by Other (30.8%) and Black (20.8%) with a mean age of 29.5 ± 5.3. Pre-pregnancy BMI and BMI at collection were 29.9 ± 7.1 kg/m² and 33.5 ± 6.2 kg/m², respectively. Gestational age at sample collection and at delivery were 69.5 ± 21.2 days and 260.7 ± 29.5 days, respectively. The median gravity was 2 (IQR 1-3). During pregnancy, a total of 16 urinary tract infections were reported (**Supplemental Table 1**).

### Sequencing and Quality Control

Oxford Nanopore sequencing of 138 urogenital samples generated a total of 53,395,690 raw reads, with a median sequencing depth of 341,706 reads per sample. Following quality filtering at Q≥12, 50,223,320 reads were retained, corresponding to 94.1% of the raw sequencing reads.

The quality-filtered dataset comprised 96.9 Gb of sequence data. Retained reads had a mean length of 1,809.5 bp and a median length of 1,525 bp, with an N50 read length of 2,382 bp. The mean Phred quality score was 17.5 (median, 17.4), and 81.8% of retained reads had quality scores greater than Q15.

Sequential removal of reads mapping to the *Allobacillus halotolerans* and *Imtechella halotolerans* reference genomes, followed by removal of reads mapping to the human reference genome (GRCh38), resulted in 9,117,654 microbial reads retained for downstream taxonomic and functional analyses. The final microbial dataset contained a median of 21,479 reads per sample, with individual sample counts ranging from 97 to 476,370 reads.

Taxonomic classification successfully assigned 8,638,189 reads, representing 94.7% of the microbial reads retained after filtering with a median of 20,516 classified reads per sample. These results indicate that the quality-control and decontamination pipeline was efficiently enriched for microbial sequences while maintaining sufficient sequencing depth for downstream taxonomic and functional analyses.

### Microbial Community Clustering and Composition

Across all participants, the microbial communities were predominated by the genera *Lactobacillus* (prevalence, 92.0%; mean relative abundance, 58.8%) and *Gardnerella* (65.9%; 24.0%), followed by *Streptococcus* (65.2%; 2.7%), *Corynebacterium* (65.2%; 2.5%), and *Fannyhessea* (formerly *Atopobium*) (37.7%; 2.3%) (**Supplemental Figure 1A**).

Hierarchical clustering of species-level relative abundance profiles identified three microbial community clusters comprising 42, 54, and 42 samples, respectively (**Figure 1A**). Species-level analysis revealed three distinct microbial community state types. Cluster 1 was characterized by a highly *Lactobacillus crispatus*-predominant microbiome, with *L. crispatus* present in all Cluster 1 samples with a mean relative abundance of 84.9% (**Figure 1B**). Cluster 2 was characterized by a polymicrobial *Gardnerella-*predominant community composed primarily of *G. swidsinskii*, *G. vaginalis*, *G. greenwoodii*, and *G. leopoldii*, together with *Fannyhessea vaginae* (**Figure 1B**). In contrast, Cluster 3 was predominated by *Lactobacillus iners* (relative abundance 61.8%), followed by *L. jensenii* (22.2%) (**Figure 1B**, **Supplemental Figure 1B**). These three microbial community clusters closely resemble the well-described vaginal community state types (CSTs) (Ravel et al., 2011; France et al., 2020). Cluster 1 broadly corresponds to CST I (*L. crispatus-*predominant), Cluster 3 to a combination of CST III and CST V (*L. iners*- and *L. jensenii*-predominant, respectively), and Cluster 2 resembling the low-*Lactobacillus*, anaerobe-rich communities characteristic of CST IV.

**Figure 1.**
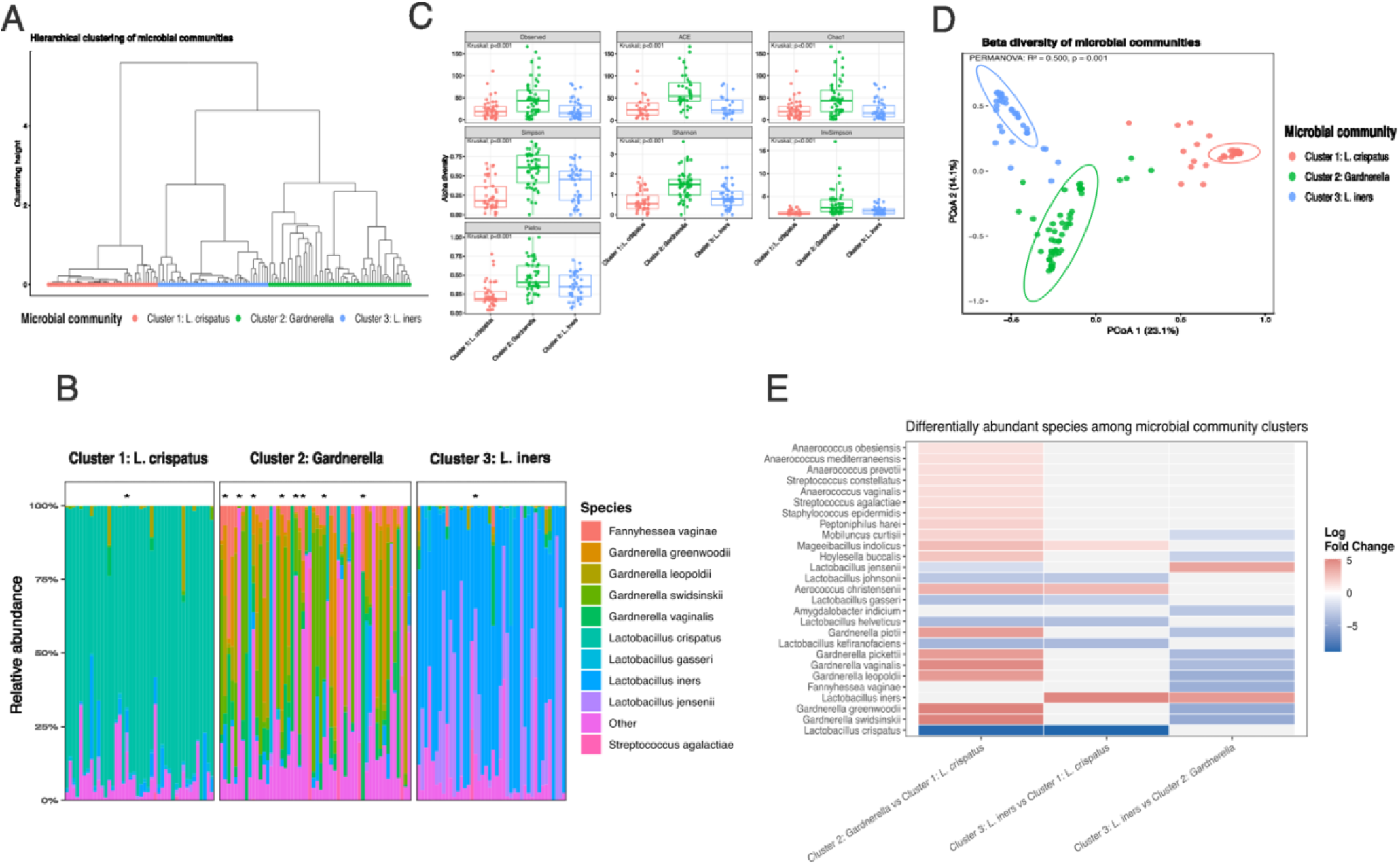
Characterization of urogenital microbial community clusters identified from Oxford Nanopore shotgun metagenomic sequencing. (A) Hierarchical clustering of samples based on Bray–Curtis dissimilarities of species-level relative abundances, identifying three distinct microbial community clusters. (B) Species-level relative abundance profiles for individual samples grouped by microbial community cluster. The three clusters were characterized by predominance of *L. crispatus* (Cluster 1), *Gardnerella* species (Cluster 2), and *L. iners* (Cluster 3); *, UTI during pregnancy. (C) Comparison of alpha diversity metrics across microbial community clusters, including observed richness, Chao1 richness, Shannon diversity, Simpson diversity, inverse Simpson diversity, Fisher’s alpha, Pielou’s evenness, and Faith’s phylogenetic diversity. Statistical significance was assessed using the Kruskal–Wallis test with pairwise Wilcoxon rank-sum tests and Benjamini–Hochberg correction for multiple comparisons. (D) PCoA of Bray–Curtis distances showing separation among the three microbial community clusters. Ellipses represent the 95% confidence intervals for each cluster. (E) Heatmap of differentially abundant bacterial species among microbial community clusters. Colors represent log_2_ fold changes, with red indicating enrichment and blue indicating depletion relative to the comparison group.

To identify clinical variables associated with microbial community composition, Boruta feature selection was applied to demographic and clinical metadata. UTI during pregnancy was the only variable confirmed as predictive of microbial community cluster membership. Consistent with this finding, UTI prevalence differed significantly among clusters (Fisher’s exact test, *p*=0.024), with the highest prevalence observed in the *Gardnerella*-predominant community (33.3%), compared with the *L. crispatus*-predominant (6.2%) and *L. iners*-predominant (4.8%) communities.

### Measures of Alpha Diversity

Alpha diversity differed significantly among the three microbial community clusters (Kruskal-Wallis p<0.001 for all measures) (**Figure 1C**). The *Gardnerella*-predominant community (Cluster 2) exhibited the highest species richness, evenness and overall diversity, whereas the *L. crispatus*-predominant community (Cluster 1) showed the lowest diversity, consistent with a community predominated by a single species. The *L. iners* cluster (Cluster 3) displayed intermediate diversity values (**Figure 1C**).

### Beta Diversity

PCoA using Bray-Curtis Dissimilarity distances demonstrated statistically significant differences in microbial community composition among the three clusters (PERMANOVA, R²=0.500, p=0.001) (**Figure 1D**). Homogeneity of multivariate dispersion also differed significantly among clusters (PERMDISP, p=0.001), indicating that some of the observed compositional differences were due to differences of within-cluster variability (*i.e.*, unequal dispersion).

### Statistical analysis

Differential abundance analysis identified characteristic species-level differences among the three clusters. ANCOM-BC2 revealed the largest number of differentially abundant species between the *Gardnerella*-predominant and *L. crispatus*-predominant clusters, with 24 significantly different species. The *Gardnerella*-predominant cluster was enriched in multiple *Gardnerella* species, including *G. greenwoodii, G. swidsinskii, G. vaginalis, G. piotii, G. pickettii,* and *G. leopoldii*, as well as several anaerobes *(e.g., Hoylesella buccalis* and several *Anaerococcus* species) and a few facultative anaerobes *(e.g., Streptococcus agalactiae* and *S. constellatus*). In contrast, the *L. crispatus*-predominant cluster was enriched in *L. crispatus* and several other *Lactobacillus* species, including *L. kefiranofaciens, L. helveticus*, *L. acidophilus*, *L. gasseri* and *L. jensenii* among others. Eight species differed significantly between the *L. iners*- and *L. crispatus*-predominant clusters. Besides *L. iners*, Cluster 3 was enriched in *Aerococcus christensenii* and *Mageeibacillus indolicus*. In addition to *L. crispatus*, Cluster 1 was enriched in several *Lactobacillus* species. Twelve species differed between the *L. iners*- and *Gardnerella*-predominant clusters, with multiple *Gardnerella* species, *F. vaginae*, and a few anaerobes enriched in Cluster 2, but only *L. iners* and *L. jensenii* in Cluster 3 (**Figure 1E; Supplemental Table 2).**

### Functional Profiles

Functional annotation of metagenomic reads identified 8,361 KEGG Orthology (KO) groups across the 138 samples. After filtering to retain KOs detected in at least five samples and with a minimum total abundance of 10 counts across the dataset, 4,313 KOs were retained for downstream analyses. Overall, predicted functional profiles closely reflected taxonomic community structure, with both KO- and pathway-level variation corresponding to the three clusters or microbiome community state types.

Among the clinical and demographic variables evaluated, UTI during pregnancy was associated with predicted pathway-level functional composition in univariate analysis (PERMANOVA, R²=0.041, *p*=0.030), although this association did not remain significant after multiple-testing correction (FDR=0.179). Among samples with available UTI data (*n*=61), the association between UTI and functional composition was no longer significant after accounting for microbiome community state (R²=0.024, *p*=0.086), whereas community state remained strongly associated with functional composition (R²=0.244, *p*<0.001). After FDR correction, no other clinical or demographic variables were significantly associated with predicted pathway-level functional composition.

### KO-level functional composition

KO-level functional composition also differed significantly among microbiome community states. Bray–Curtis PCoA revealed separation among the *L. crispatus-, Gardnerella-*, and *L. iners*-predominant communities, with PCoA1 and PCoA2 explaining 23.4% and 14.9% of the variation, respectively (38.3% combined). Microbiome community state explained 25.5% of the variation in KO composition (PERMANOVA, R²=0.255, F=23.16, p<0.001). Multivariate dispersion did not differ significantly among community states (PERMDISP, F=0.613, p=0.548), indicating that the observed differences in functional composition were not attributable to unequal within-group dispersion (**Figure 2A**).

**Figure 2.**
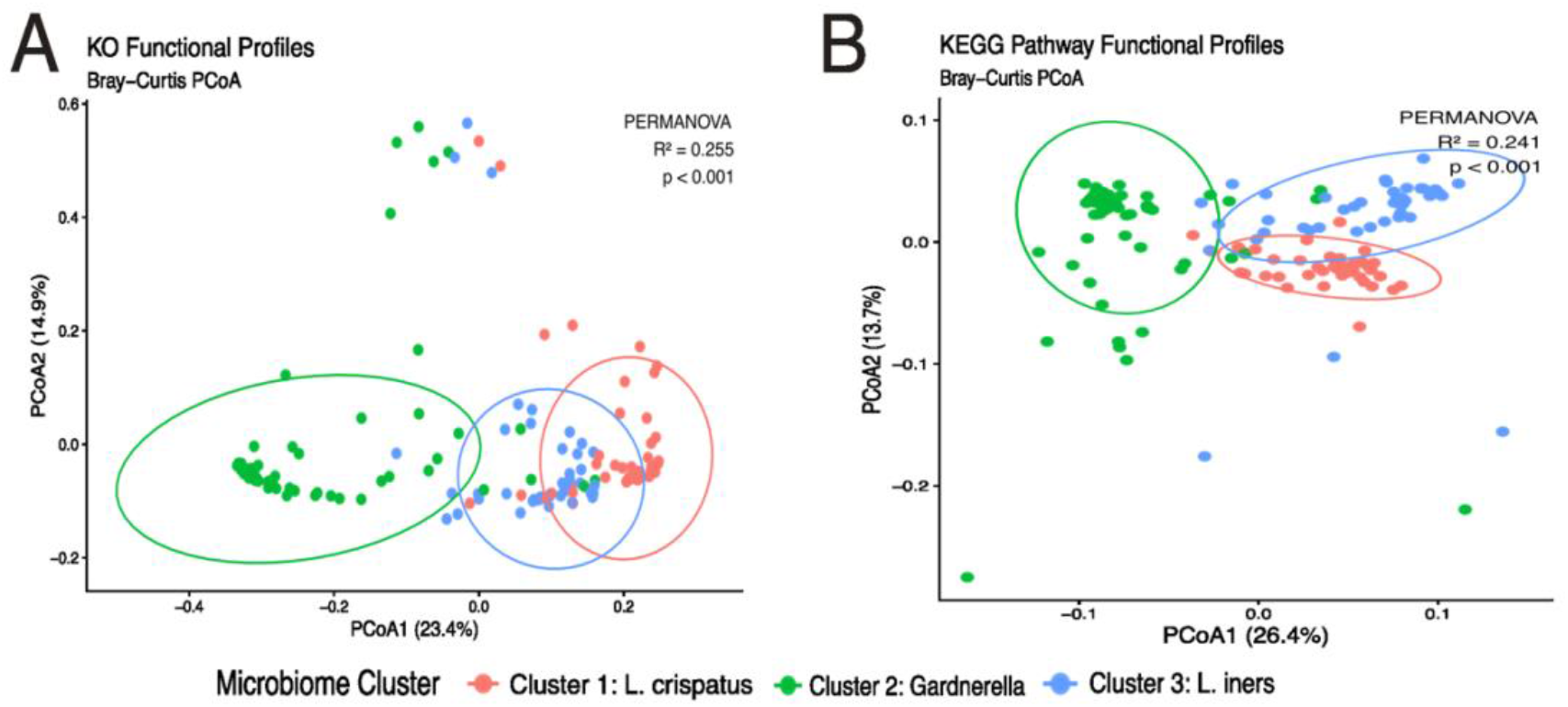
Functional differences among urogenital microbiome community states. (A) PCoA of KEGG Orthology (KO) functional profiles based on Bray–Curtis dissimilarity. Samples are colored according to the taxonomically defined urogenital microbiome clusters: *L. crispatus-, Gardnerella*-, and *L. iners*-predominated communities. PCoA1 and PCoA2 explained 23.4% and 14.9% of the variation in KO composition, respectively. Differences in functional composition among microbiome clusters were evaluated using PERMANOVA with 9,999 permutations, and homogeneity of multivariate dispersion was assessed using PERMDISP. Differences among urogenital microbiome clusters were assessed by PERMANOVA (R²=0.255, F=23.16, p<0.001); homogeneity of multivariate dispersion was confirmed by PERMDISP (F=0.613, p=0.548). (B) PCoA of KEGG pathway profiles based on Bray–Curtis dissimilarity. Each point represents one sample, and colors indicate the taxonomically defined *L. crispatus-, Gardnerella-,* and *L. iners*-predominated urogenital microbiome clusters. Ellipses represent the distribution of samples within each community state. PCoA1 and PCoA2 explained 26.4% and 13.7% of the total variation, respectively. Differences in functional composition among microbiome clusters were evaluated using PERMANOVA with 9,999 permutations, and homogeneity of multivariate dispersion was assessed using PERMDISP. Differences among urogenital microbiome clusters were assessed by PERMANOVA (R²=0.241, F=21.49, p<0.001); homogeneity of multivariate dispersion was confirmed by PERMDISP (F=0.245, p=0.785).

### Pathway-level functional composition

A similar pattern was observed at the KEGG pathway level. Bray–Curtis PCoA showed separation among the three microbiome community states, with PCoA1 and PCoA2 explaining 26.4% and 13.7% of the variation, respectively (40.1% combined). Microbiome community state explained 24.1% of the variation in pathway composition (PERMANOVA, R²=0.241, F=21.49, p<0.001). As observed for KO-level profiles, multivariate dispersion did not differ significantly among community states (PERMDISP, F=0.245, p=0.785), supporting differences in functional composition rather than differences in within-group variability (**Figure 2B**).

Across both functional resolutions, the *L. crispatus*- and *L. iners*-predominant communities occupied more similar regions of the ordination space, whereas the *Gardnerella*-predominant community was more clearly separated. This consistent pattern indicates greater functional similarity between the two *Lactobacillus*-predominant community states, with the *Gardnerella*-predominant community representing a more distinct functional state. *Cluster-enriched KO analysis*. Differential abundance analysis identified distinct sets of KOs associated with each microbiome community state. KOs were evaluated using DESeq2 with one-versus-rest comparisons, contrasting each community state against the other two communities combined. KOs with a Benjamini–Hochberg adjusted p<0.05 and log_2_ fold change >1 were considered significantly enriched. The *L. crispatus-Gardnerella-* and *L. iners-*predominated clusters were enriched with 249, 736, and 300 KOs, respectively (**Supplemental Table 3**). To visualize community-specific functional signatures, only the top 20 enriched KOs from each community state were selected (**Figure 3**). The *L. crispatus*-predominant community was characterized by enrichment of KOs associated with carbohydrate utilization and transport, including lactose/cellobiose-specific phosphotransferase system (PTS) components and mannitol metabolism. Additional enriched functions included ABC-type bacteriocin/lantibiotic export systems, major facilitator superfamily transport, iron transport-associated functions, and several mobile genetic element-associated functions, including transposases and a putative reverse transcriptase. The *Gardnerella*-predominant community exhibited enrichment of a distinct set of metabolic functions, including KOs associated with carbohydrate metabolism, glycosyl hydrolase, phosphoglycerate mutase, aminotransferase activity, thioredoxin-associated functions, and ABC transport. The *L. iners*-predominant community was characterized by enrichment of several membrane- and transport-associated functions, including FtsX-like permeases, sugar transporters, bacterial solute-binding proteins, and other transmembrane transport functions, together with KOs associated with cell adhesion, peptidase activity, and energy transduction. Overall, these findings reveal distinct KO-level functional signatures for each of the three microbiome community states.

**Figure 3.**
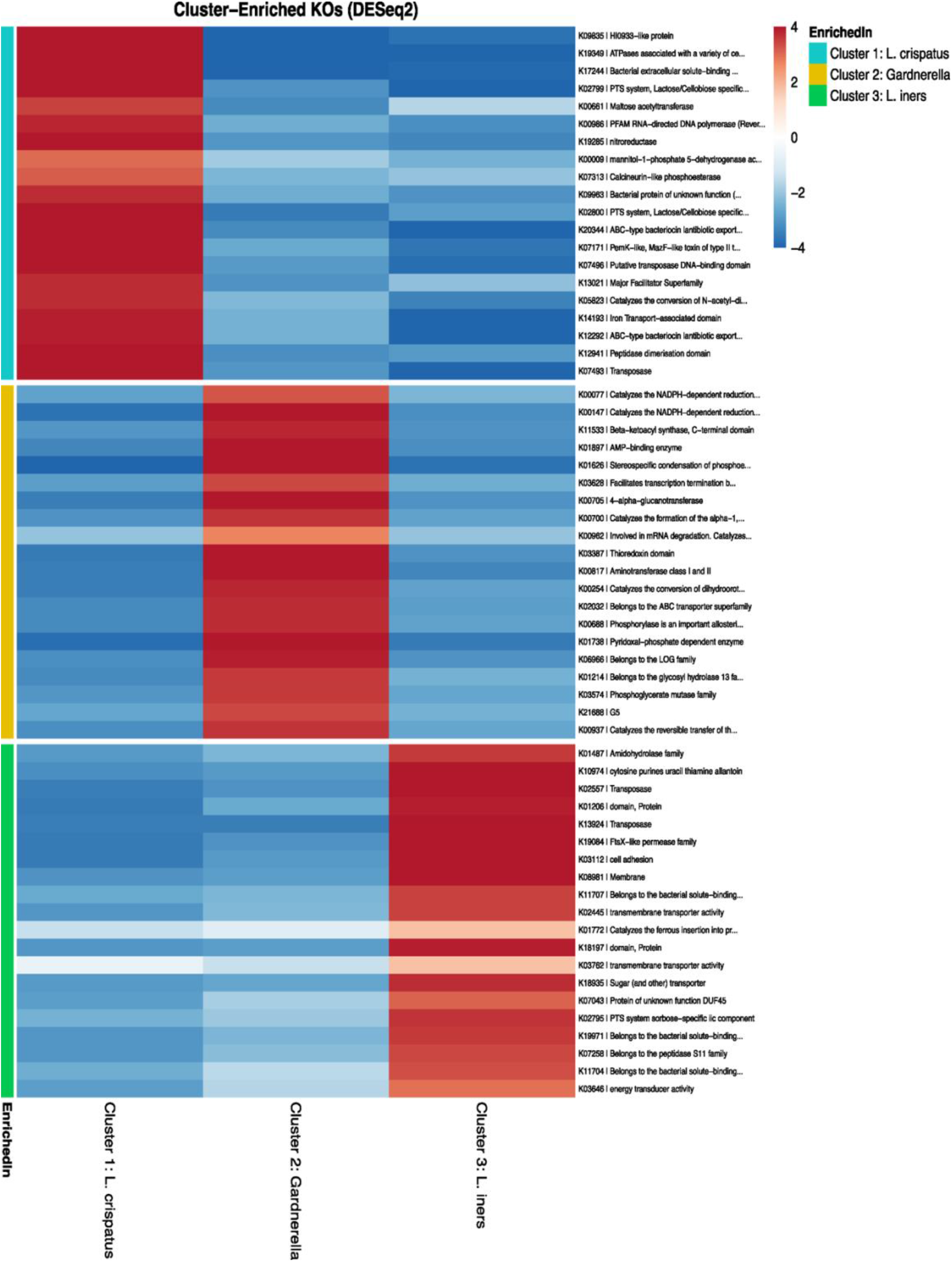
Heatmap of differentially abundant KOs across urogenital microbiome community states. Differential abundance was assessed using DESeq2 with one-versus-rest comparisons. KOs with Benjamini–Hochberg adjusted p<0.05 and log_2_ fold change >1 were considered significantly enriched. The top 20 enriched KOs for each microbiome community state are shown. Heatmap cells represent the DESeq2 log_2_ fold change for each KO across the three one-versus-rest comparisons, with positive values indicating relative enrichment and negative values indicating relative depletion. For visualization, log_2_ fold-change values were truncated at −4 and +4; statistical testing and KO selection were performed using the original, untruncated values. The annotation bar indicates the community state in which each KO was significantly enriched. KOs are labeled by KEGG Orthology identifier and eggNOG-derived functional annotation values. The colored annotation bar indicates the microbiome cluster in which each KO was significantly enriched. KOs are labeled by KEGG Orthology identifier and eggNOG-derived functional annotation.

### Cluster-enriched KEGG pathways

Differential abundance analysis also identified distinct KEGG pathway enrichment patterns for each of the three microbiome community states (**Figure 4; Supplemental Table 4**). Following removal of redundant KO and map representations of the same KEGG pathways, 11 pathways were significantly enriched in the *L. crispatus*-predominant community, 57 in the *Gardnerella*-predominant community, and 10 in the *L. iners*-predominant community (adjusted p<0.05 and log_2_ fold change >1).

**Figure 4.**
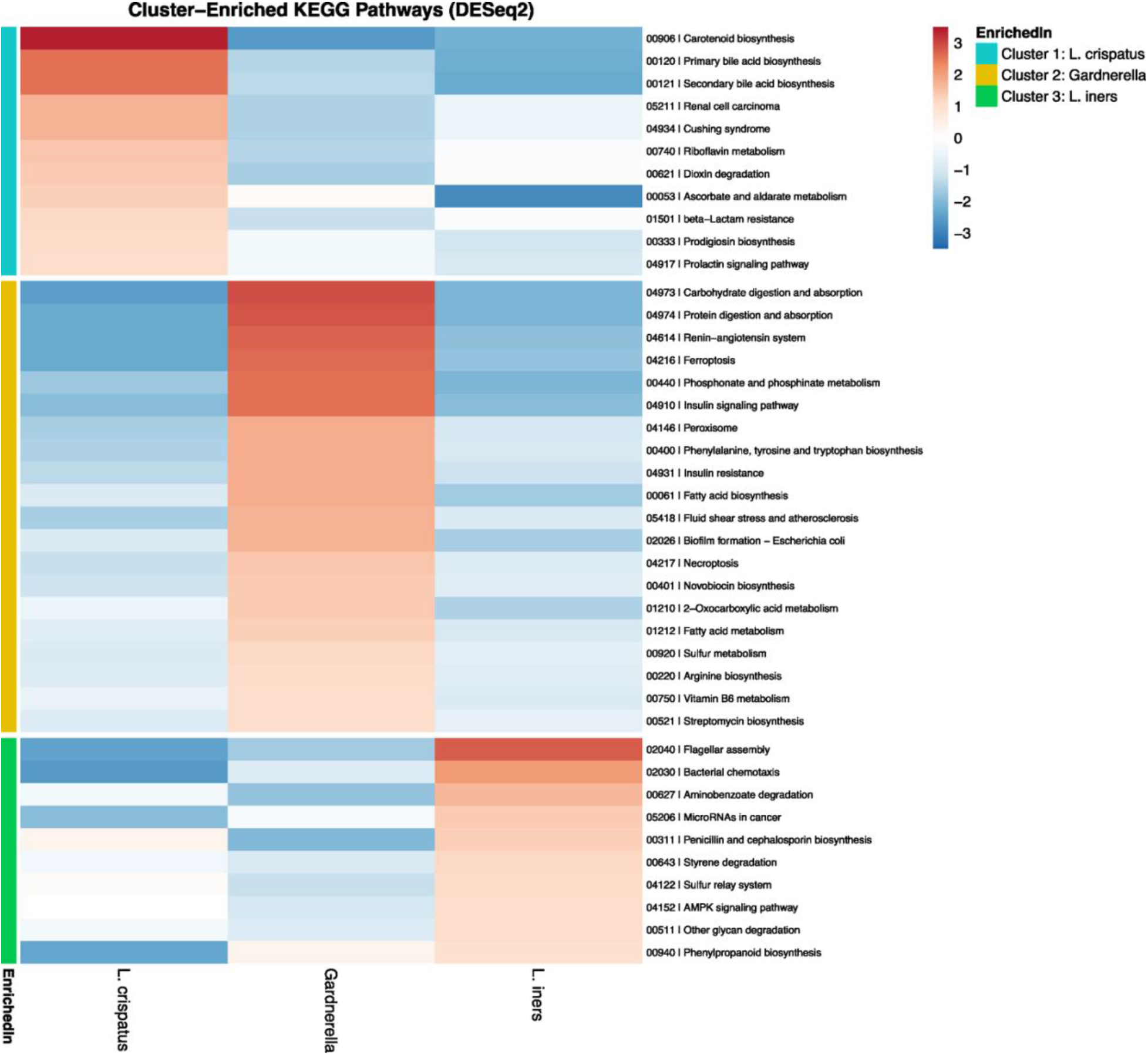
Cluster-enriched KEGG pathway profiles across urogenital microbiome community states. Heatmap showing differentially abundant KEGG pathways among the *L. crispatus*-, *Gardnerella*-, and *L. iners*-predominated community states. Differential abundance was assessed using DESeq2 with one-versus-rest comparisons, contrasting each community state against the other two combined. Pathways with a Benjamini–Hochberg adjusted *p*<0.05 and log_2_ fold change >1 were considered significantly enriched. The heatmap includes all significantly enriched pathways for the *L. crispatus*- and *L. iners*-predominated communities and the top 20 for the *Gardnerella*-predominated community, ranked by adjusted *P* value. Cells represent DESeq2 log_2_ fold changes across the three comparisons, with positive and negative values indicating relative enrichment and depletion, respectively. Rows are grouped according to the community state in which each pathway was significantly enriched.

The *L. crispatus*-predominant community was enriched in pathways annotated as riboflavin metabolism; carotenoid biosynthesis; primary and secondary bile acid biosynthesis; and ascorbate and aldarate metabolism. In contrast, the *Gardnerella*-predominant community was enriched in pathways associated with fatty acid biosynthesis and metabolism; phenylalanine, tyrosine, and tryptophan biosynthesis; arginine biosynthesis; sulfur metabolism; vitamin B6 metabolism; 2-oxocarboxylic acid metabolism; phosphonate and phosphinate metabolism; and biofilm formation. The *L. iners*-predominant community displayed a distinct functional signature that included pathways annotated as bacterial chemotaxis; flagellar assembly; other glycan degradation; aminobenzoate degradation; styrene degradation; sulfur relay systems; and penicillin and cephalosporin biosynthesis.

Although the *L. crispatus*- and *L. iners*-predominated communities occupied relatively similar regions of the pathway-level ordination space, differential abundance analysis identified distinct sets of enriched pathways within each community state. Together with the pathway-level beta-diversity results, these findings indicate that taxonomically defined microbiome communities differ not only in their overall predicted functional composition but also in specific functional signatures, with the *Gardnerella*-predominant community exhibiting the broadest set of enriched pathways.

### Taxon–KEGG pathway associations reveal distinct functional signatures across urogenital taxa

Correlation analysis revealed distinct relationships between bacterial taxa and KEGG functional pathways (**Figure 5**). Following identification of significant taxon–KO correlations (adjusted p<0.05, |ρ|≥0.40), pathway-level associations were retained when supported by at least three significantly correlated KOs with ≥70% directional concordance. The resulting profiles showed substantial taxon-specific structure, with related taxa frequently exhibiting similar association patterns across multiple metabolic and cellular pathways. Importantly, these correlations represent associations between taxon abundance and functional profiles and do not establish that the associated KOs or pathways originated from, or were encoded by, the correlated taxon.

**Figure 5.**
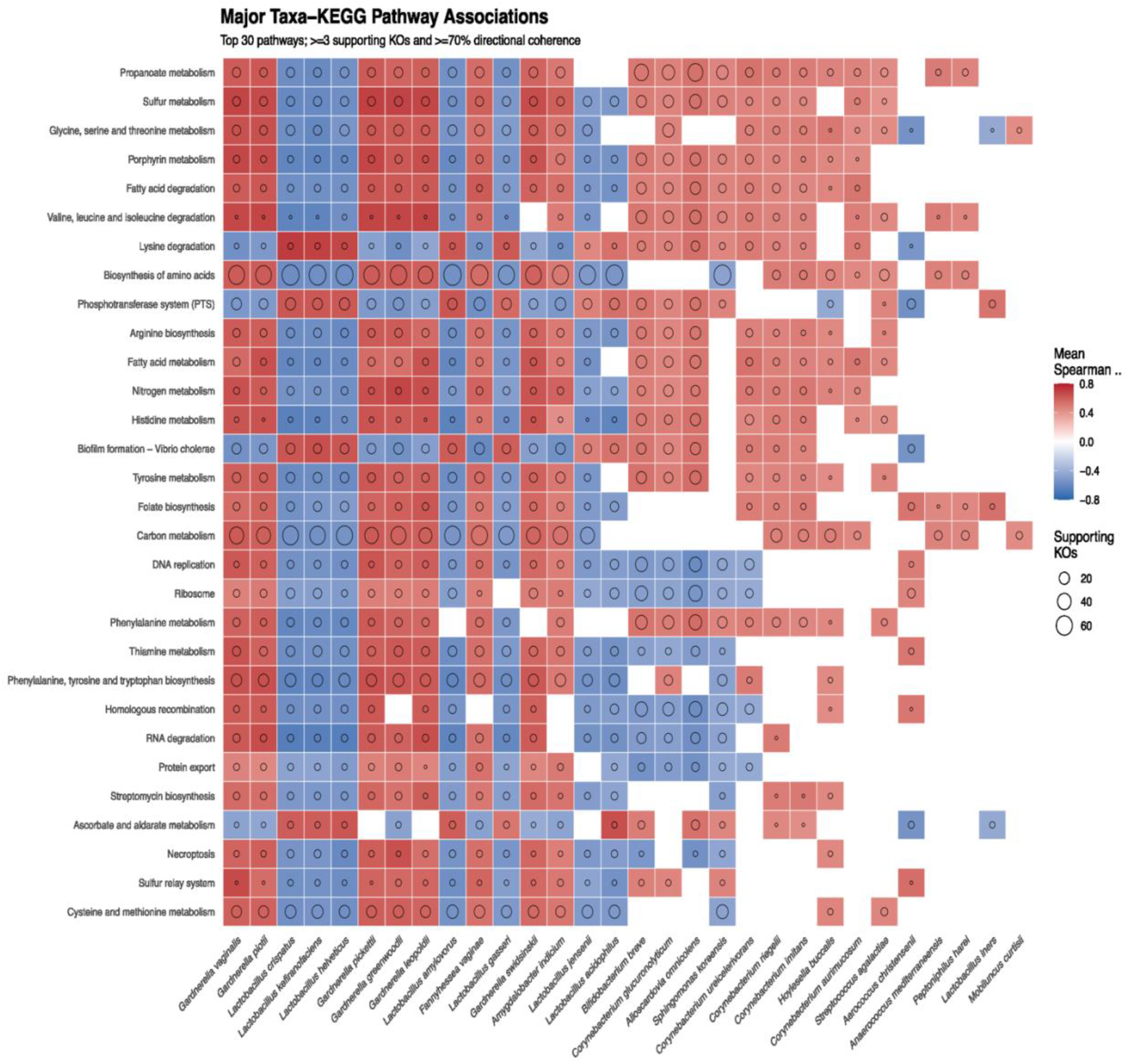
Associations between urogenital bacterial taxa and KEGG functional pathways. Heatmap showing associations between bacterial species and the top 30 KEGG pathways derived from significant taxon–KEGG Orthology (KO) correlations. Taxonomic and KO abundance profiles were centered log-ratio (CLR) transformed, and associations were evaluated using Spearman correlation. Taxon–KO associations with Benjamini–Hochberg adjusted *P*<0.05 and |ρ|≥0.40 were mapped to KEGG pathways. Taxon–pathway associations were retained when supported by at least three significantly associated KOs and ≥70% directional concordance. Tile color represents the mean Spearman correlation coefficient among KOs supporting the predominant direction, and circle size represents the number of significantly associated KOs. The 30 pathways shown were ranked by the number of associated taxa, total KO support, and mean absolute association strength. Associations represent correlations between taxonomic and functional profiles and do not establish that the associated functions originated from or were encoded by the correlated taxon.

A prominent contrasting pattern was observed between *Gardnerella* and *Lactobacillus* species. Multiple *Gardnerella* species were positively associated with pathways involved in propanoate metabolism, sulfur metabolism, fatty acid degradation and metabolism, amino acid biosynthesis, arginine biosynthesis, nitrogen metabolism, histidine metabolism, folate biosynthesis, and cysteine and methionine metabolism, among others. Most of these pathways were negatively associated with *Lactobacillus* species, including *L. crispatus, L. gasseri,* and *L. jensenii,* but not *L. iners*. Conversely, pathways including the phosphotransferase system (PTS), lysine degradation, biofilm formation-related functions, and ascorbate and aldarate metabolism showed positive associations with the same *Lactobacillus* species and negative associations with multiple *Gardnerella* species. Taken together, these results demonstrate contrasting functional association profiles between members of the *Gardnerella*-predominant cluster and members of the *L. crispatus*-predominant cluster.

Distinct functional association profiles were also observed for other prevalent taxa, including *Fannyhessea, Alloscardovia, Bifidobacterium,* and *Corynebacterium*. Together, these results indicate that differences in taxonomic composition across urogenital microbiome community states are accompanied by coordinated differences in functional potential, with particularly contrasting patterns associated with *Gardnerella* and *Lactobacillus* species.

### Taxon–pathway association network

To identify the strongest and most consistently supported taxon–function relationships, we further restricted the analysis to pathway associations supported by at least 15 significantly correlated KOs with ≥70% directional concordance. The resulting network comprised 25 bacterial taxa, 58 KEGG pathways, and 272 taxon–pathway associations (**Figure 6**). *Alloscardovia omnicolens* was the most highly connected taxonomic hub, with 37 pathway associations (24 positive and 13 negative), followed by *Bifidobacterium breve* with 34 (23 positive and 11 negative) and *Corynebacterium glucuronolyticum* with 26 (17 positive and 9 negative).

**Figure 6.**
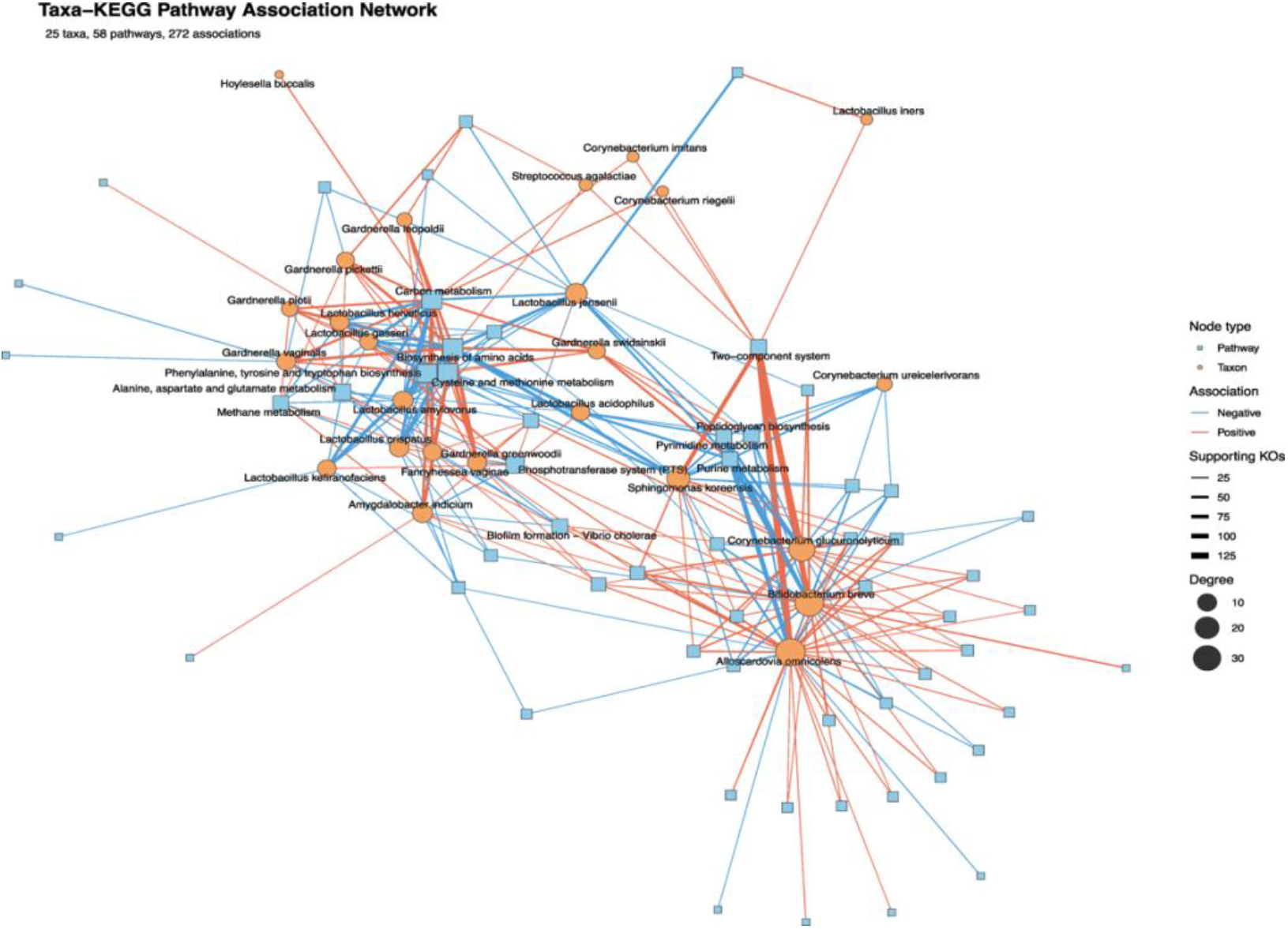
Taxon–KEGG pathway association network. Network representation of associations between bacterial taxa and KEGG pathways derived from KO-level Spearman correlations across 138 samples. Taxon–pathway associations were retained when supported by ≥15 significantly correlated KOs and ≥70% directional concordance. The final network comprised 25 bacterial taxa, 58 KEGG pathways, and 272 associations. Orange circles represent bacterial taxa, and blue squares represent KEGG pathways; node size is proportional to degree. Edges represent taxon–pathway associations, with red and blue indicating positive and negative associations, respectively, and edge width proportional to the number of supporting KOs. All bacterial taxa and the 12 KEGG pathways with the highest degree are labeled. The force-directed layout represents network connectivity rather than quantitative distance. Associations represent correlations between taxonomic and functional profiles and do not establish that the associated functions originated from or were encoded by the connected taxon.

A pronounced directional pattern was again observed among *Lactobacillus* and *Gardnerella* species. Associations involving several *Lactobacillus* species were predominantly negative, including *L. jensenii* (14 of 15 associations), *L. amylovorus* (11 of 12), *L. crispatus* (9 of 11), *L. helveticus* (8 of 10), *L. acidophilus* (8 of 9), *L. kefiranofaciens* (8 of 9), and *L. gasseri* (7 of 8). In contrast, associations involving several *Gardnerella* species were predominantly positive, including *G. greenwoodii* (8 of 10), *G. pickettii* (7 of 8), *G. swidsinskii* (7 of 7), *G. piotii* (6 of 6), and *G. leopoldii* (5 of 5), whereas *G. vaginalis* exhibited an equal distribution of positive and negative associations.

At the pathway level, biosynthesis of amino acids and carbon metabolism were the most highly connected functional hubs, each associated with 17 taxa, followed by phenylalanine, tyrosine, and tryptophan biosynthesis and cysteine and methionine metabolism, each associated with 16 taxa. Several pathways exhibited strongly directional relationships; the two-component system was positively associated with all eight connected taxa, whereas purine and pyrimidine metabolism were predominantly negatively associated, with seven of eight associations negative for each pathway.

Together, these findings reveal a highly structured taxon–function network within the urogenital microbiome, characterized by contrasting functional associations among *Lactobacillus* and *Gardnerella* species and by central roles for carbon and amino acid metabolism.

### LLM-assisted interpretation of taxon–pathway associations

To facilitate biological interpretation of the taxon–pathway association analysis, an LLM-assisted approach was used to prioritize statistically supported associations based on correlation strength, number of supporting KOs, consistency of KO-level associations, and biological relevance. The LLM was used only for interpretation and prioritization of associations identified through the statistical analysis and was not used to generate correlations or determine statistical significance. This prioritization highlighted two recurrent functional patterns, characterized by *Lactobacillus*-associated carbohydrate transport functions and *Gardnerella*-associated biosynthetic and central metabolic functions.

*Lactobacillus* species displayed a consistent functional signature characterized by positive associations with the phosphotransferase system (PTS), which is involved in carbohydrate transport and phosphorylation. Positive associations with PTS were observed for *L. amylovorus* (mean Spearman ρ=0.627; 17 supporting KOs), *L. crispatus* (ρ=0.625; 20 KOs), *L. kefiranofaciens* (ρ=0.621; 18 KOs), and *L. helveticus* (ρ=0.620; 20 KOs). These pathway-level associations were supported by recurrent KO-level correlations involving PTS-associated functions, including *scrA*, lactose/cellobiose-specific PTS components, and *mtlD*. In *L. crispatus*-associated profiles, for example, individual correlations reached ρ=0.803 for *mtlD*, ρ=0.796 for *scrA*, and ρ=0.785 for lactose/cellobiose-specific PTS components. These findings demonstrate a consistent positive association between the abundance of multiple *Lactobacillus* species and predicted carbohydrate transport and utilization functions.

In contrast, *Gardnerella* species exhibited positive associations with pathways involved in amino acid biosynthesis and central carbon metabolism. Biosynthesis of amino acids was positively associated with *G. greenwoodii* (mean ρ=0.343; 62 supporting KOs), *G. pickettii* (ρ=0.448; 54 KOs), and *G. leopoldii* (ρ=0.510; 47 KOs). These associations were supported by KOs involved in amino acid biosynthetic processes, including *pheA*, *tyrA*, *aroB*, *cysK*, and aminotransferase-associated functions. Notably, *G. pickettii* was also positively associated with phenylalanine, tyrosine, and tryptophan biosynthesis (ρ=0.592; 21 KOs), with strong KO-level associations involving *pheA* (ρ=0.811), *tyrA* (ρ=0.708), and *aroB* (ρ=0.689). Carbon metabolism was additionally associated with *G. pickettii* (ρ=0.395; 40 KOs) and *G. greenwoodii* (ρ=0.303; 50 KOs), further supporting a broader relationship between *Gardnerella* abundance and predicted central metabolic functions.

Overall, LLM-assisted prioritization highlighted contrasting functional signatures across the urogenital microbiome, with *Lactobacillus* species consistently associated with PTS-mediated carbohydrate transport and utilization functions, whereas *Gardnerella* species were preferentially associated with amino acid biosynthesis and central carbon metabolism. The recurrence of these patterns across multiple species within each genus, together with support from multiple correlated KOs, suggests that these associations reflect broader community-level differences in functional potential rather than isolated taxon–function relationships. Importantly, because these analyses are based on correlations between taxonomic and metagenomic functional abundances, they represent functional co-variation and predicted metabolic potential but do not establish direct taxonomic attribution, gene expression, or metabolic activity.

### Metagenome-assembled genomes

A total of 415 metagenome-assembled genomes (MAGs) were recovered from the dataset. Dereplication using dRep v3.6.2 grouped these genomes into 107 primary clusters at 90% average nucleotide identity (ANI) and 362 secondary clusters at 99% ANI. Based on CheckM completeness and contamination estimates, 29 MAGs were classified as high quality and 116 as medium quality. One representative genome from each secondary cluster was retained for downstream analyses.

Among the high-quality MAGs, the most frequently recovered taxa were *F. vaginae* (n=6), *L. iners* (n=5), and *G. swidsinskii* (n=3), followed by *G. leopoldii* and *G. greenwoodii* (n=2 each). Additional high-quality MAGs represented several other urogenital taxa, while some genomes could not be confidently classified to the species level.

### Comparative genomic analysis of high-quality MAGs

The 29 high-quality MAGs were examined to characterize genomic and functional variation among members of the urogenital microbiome. These genomes represented multiple prevalent taxa, including *Lactobacillus, Gardnerella, Fannyhessea, Streptococcus*, and *Escherichia* species.

Phylogenomic reconstruction based on concatenated bacterial single-copy marker genes separated the recovered genomes according to their taxonomic assignments (**Figure 7A**). Genome completeness was generally >95%, with contamination remaining <5%, supporting the quality of the genomes included in downstream comparative analyses.

**Figure 7.**
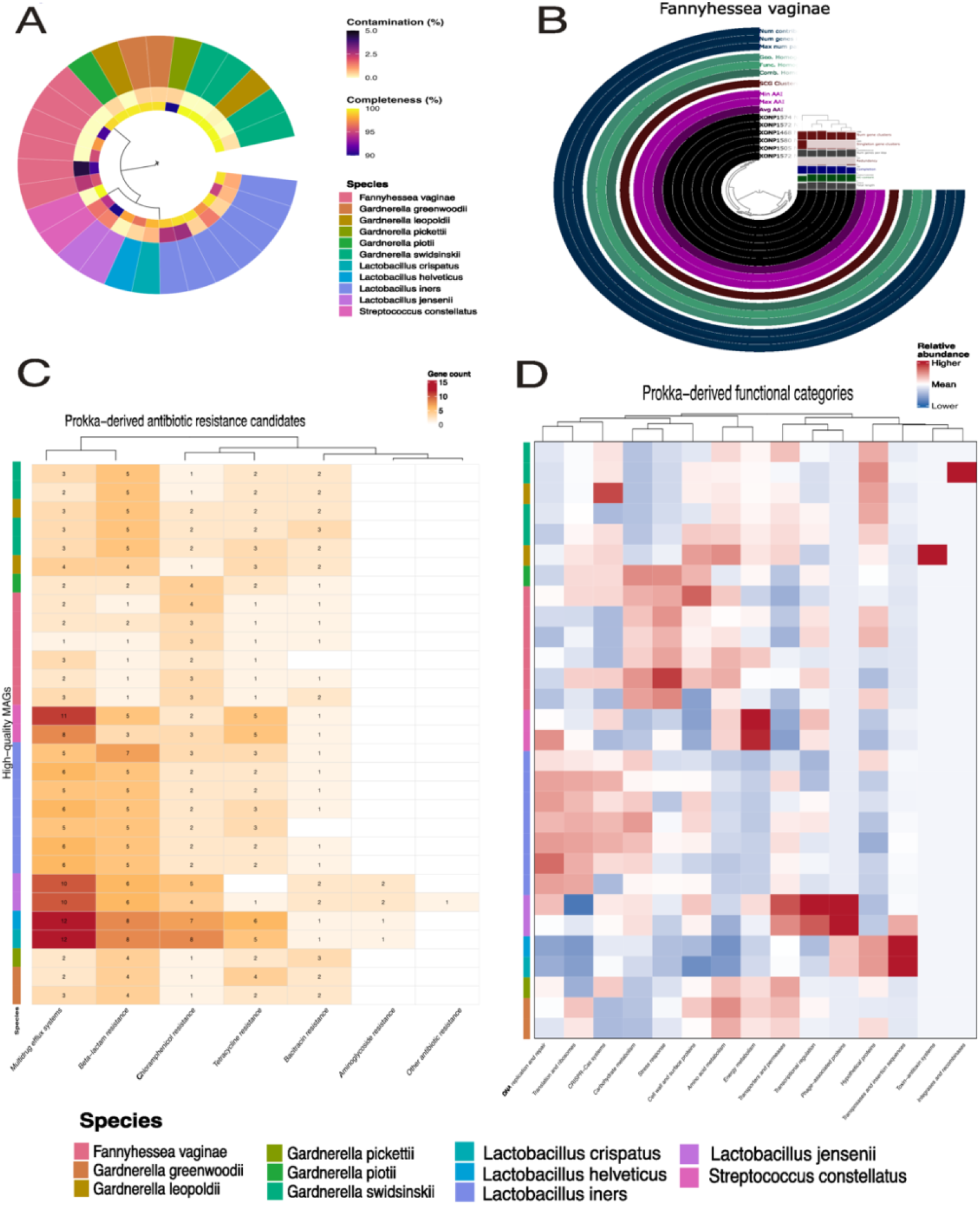
Characterization of 29 recovered high-quality MAGs. (A) Circular phylogenomic tree constructed from concatenated bacterial single-copy marker genes (Bacteria_71), showing taxonomic relationships among recovered MAGs. The innermost annotation ring represents contamination (%), followed by completeness (%), and the outer ring denotes species-level taxonomic assignment. (B) Comparative genomic analysis of six *F. vaginae* MAGs. Orthologous protein groups were identified using Proteinortho, and comparative genomic features were visualized using anvi’o. Concentric layers summarize gene-cluster characteristics, including amino acid identity (AAI), single-copy gene (SCG) clustering, combinatorial, functional, and geometric homogeneity, maximum paralog number, gene-cluster size, and the number of contributing genomes. (C) Heatmap showing putative antibiotic resistance-associated functional categories inferred from Prokka gene product annotations and keyword-based classification. Color intensity represents the number of annotated proteins assigned to each resistance-associated category within each genome. (D) Heatmap showing the distribution of predicted mobile genetic element-, defense-, and toxin-associated proteins. Predicted functional categories included transposase-, integrase/recombinase-, phage-, CRISPR/Cas-, and toxin/antitoxin-associated proteins. Color intensity represents the number of annotated proteins assigned to each category within each genome.

Six high-quality MAGs were identified as *F. vaginae*. Comparative genomic analysis revealed largely conserved gene-content profiles with variation in orthologous gene clusters among the six MAGs (**Figure 7B**). Concentric rings summarize gene-cluster characteristics, including amino acid identity (AAI), single-copy gene (SCG) clustering, combinatorial, functional, and geometric homogeneity, maximum paralog number, gene-cluster size, and the number of contributing genomes. Overall, these patterns indicate modest genomic variation among the *F. vaginae* MAGs, consistent with strain-level heterogeneity.

Functional annotation revealed putative antibiotic resistance-associated functions across the recovered MAGs (**Figure 7C**). Multidrug efflux-associated functions were broadly distributed among the genomes, whereas annotations associated with tetracycline, aminoglycoside, β-lactam, and macrolide resistance showed more restricted and taxon-specific distributions.

High-quality MAGs also differed in the distribution of mobile genetic element-, defense-, and toxin-associated functions (**Figure 7D**). Transposase-, integrase/recombinase-, phage-, CRISPR/Cas-, and toxin/antitoxin-associated functions showed variable distributions across genomes and taxa, indicating heterogeneity in the accessory functional repertoire of the recovered MAGs.

The previously identified association of bacterial chemotaxis and flagellar assembly pathways with Cluster 3 is curious as *L. iners* is well-documented as non-motile. Therefore, we sought associations between chemotaxis and flagellar genes and members of the *L. iners-*predominant cluster and found a strong association with the flagellated species *Sphingomonas koreensis* (mean ρ=0.481, p=4.56x10^-14^) (**Supplemental Table 5**). However, *S. koreensis* is a known sequencing kit-associated contaminant; thus, it is possible that these associations are a consequence of low-level contamination (equivalent to <1000 cfu/mL) (Ferneyhough et al., 2025).

### Virome characterization reveals increased viral richness in Gardnerella-predominant communities

Using geNomad, viral sequences were detected in 116 of 138 samples (84.1%). A total of 1,802 viral sequences were identified across the metagenomic assemblies. Most classified viral sequences were assigned to the class Caudoviricetes (85.9%, 1,548/1802), indicating that the detected urogenital virome was predominantly composed of tailed bacteriophages (**Figure 8A**). At the family level, however, the vast majority (97.2%, 1751/1802) of viral sequences remained unclassified. Only a small proportion (2.8%, 51/1802) were assigned to families, including Kyanoviridae, Herelleviridae, Aliceevansviridae, and Rountreeviridae. The high proportion of unclassified viral sequences highlights the limited taxonomic resolution currently available for many viruses detected within the urogenital microbiome.

**Figure 8.**
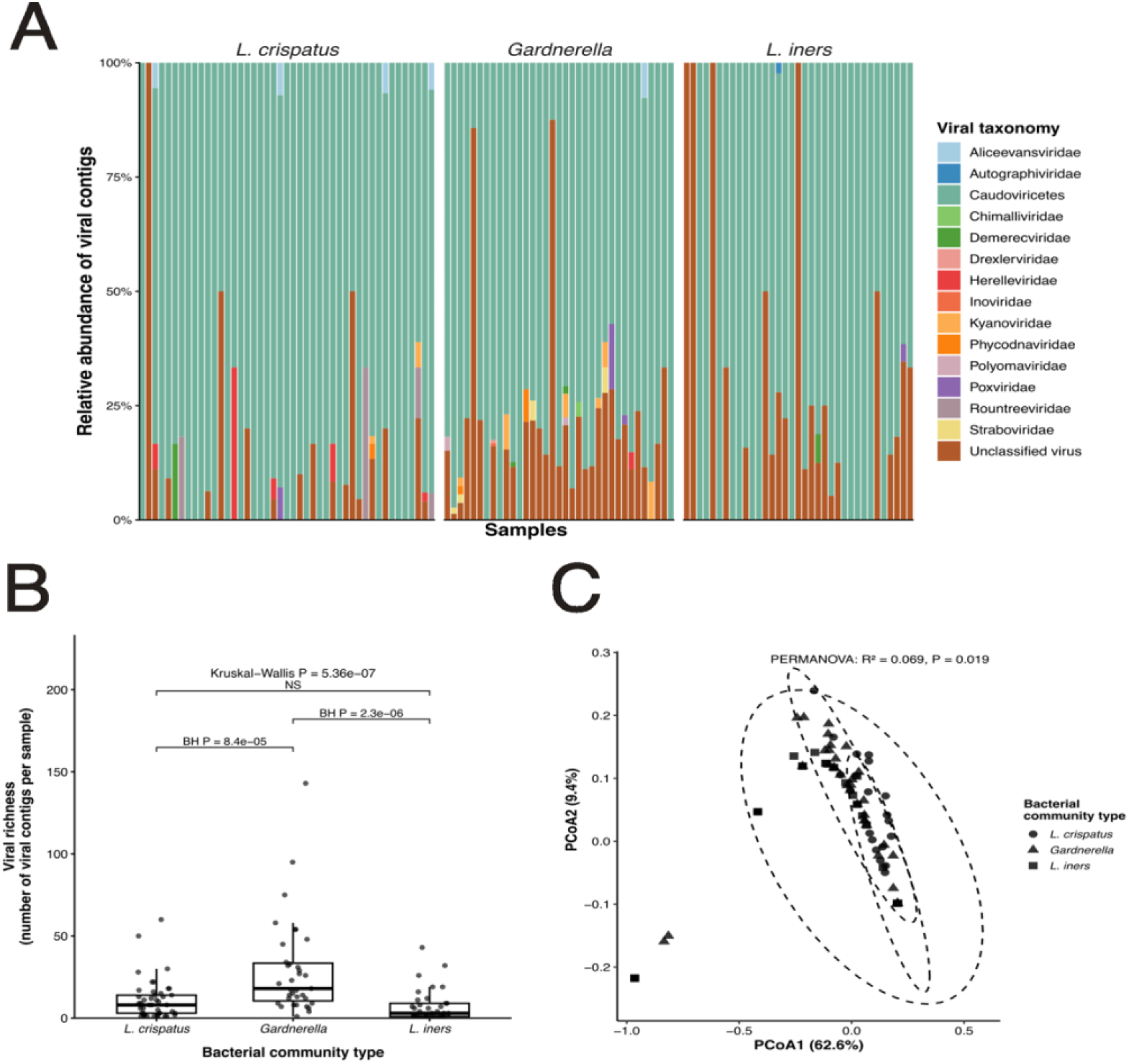
Viral richness and community composition across urogenital microbiome community states. (A) Relative abundance of the most prevalent viral taxa across samples, grouped according to bacterial community state. (B) Viral richness, measured as the number of viral contigs detected per sample, across *L. crispatus*-, *Gardnerella*-, and *L. iners*-predominated community states. Overall differences were assessed using the Kruskal–Wallis test, followed by Benjamini–Hochberg-adjusted pairwise Wilcoxon rank-sum tests. (C) PCoA of Bray–Curtis dissimilarities based on viral taxonomic profiles. Differences among bacterial community states were evaluated using PERMANOVA.

To determine whether viral richness differed according to bacterial community structure, the number of viral contigs detected per sample was compared among the three previously identified microbiome community states (**Figure 8B**). Mean viral richness was 27.5 viral contigs per sample in *Gardnerella*-predominant communities, compared with 9.7 and 6.8 viral contigs per sample in the *L. crispatus*- and *L. iners*-predominant communities, respectively. Thus, viral richness differed significantly among the three community states (Kruskal–Wallis, p=1.89×10⁻⁶). Pairwise comparisons showed that *Gardnerella*-predominant communities contained significantly more viral contigs than both *L. crispatus*-predominant (adjusted p=8.7×10⁻⁵) and *L. iners*-predominant communities (adjusted p=6.6×10⁻⁶), whereas no significant difference was observed between the two *Lactobacillus*-predominant community states.

Viral community composition also differed significantly among bacterial community states (**Figure 8C**). Bray–Curtis PCoA showed differences in viral taxonomic profiles among the *L. crispatus-, Gardnerella*-, and *L. iners*-predominant communities, with PCoA1 and PCoA2 explaining 62.6% and 9.4% of the variation, respectively (72.0% combined). Bacterial community state explained 6.9% of the variation in viral community composition (PERMANOVA, R²=0.069, p=0.019), indicating that differences in bacterial community structure were accompanied by differences in virome composition.

Viral richness differed strongly according to bacterial community state, suggesting a close relationship between bacterial community structure and viral richness within the urogenital microbiome. However, exploratory analyses evaluating associations between viral richness and available clinical and demographic variables identified no statistically significant relationships after correction for multiple testing.

## DISCUSSION

In this cohort of pregnant females, long-read shotgun metagenomic sequencing of midstream voided urine samples identified three recurring urogenital community states predominated by *L. crispatus, Gardnerella* species, and *L. iners*. These states differed not only in taxonomic diversity but also functional potential, indicating that the observed clustering reflects broader ecological variation rather than taxonomic identity alone.

### Urogenital microbial community states and UTI during pregnancy

The three microbial community states identified *de novo* from species-level shotgun metagenomic profiles of midstream voided urine resemble community state types previously described in studies of the vaginal microbiome. *L. crispatus*-predominated communities exhibited comparatively low bacterial diversity, consistent with the well-established association between *L. crispatus* predominance and stable, low-diversity vaginal microbial communities. *L. iners*-predominated communities displayed an intermediate ecological profile, consistent with previous studies describing *L. iners* as a species capable of persisting across a broad range of vaginal community configurations, including transitional states between *Lactobacillus*-predominated and more diverse communities. The *Gardnerella*-predominated community was most diverse, resembling the anaerobic communities frequently associated with vaginal dysbiosis and bacterial vaginosis (Ravel et al., 2011; Ma et al., 2012; Petrova et al., 2017; Van De Wijgert and Jespers, 2017).

The *Gardnerella*-predominated state showed the highest frequency of UTI during pregnancy, making it the clearest clinical association detected. *Gardnerella* species have been associated with bacterial vaginosis and urinary tract dysbiosis and have increasingly been investigated in the context of recurrent UTI. Experimental studies have demonstrated that transient bladder exposure to *Gardnerella* can trigger recurrence of uropathogenic *Escherichia coli* infection from intracellular reservoirs in murine models, suggesting a potential mechanism through which changes in the urogenital microbial environment may influence UTI susceptibility (Gilbert et al., 2017). These findings raise the possibility that *Gardnerella*-rich communities may contribute to an ecological environment permissive to UTI rather than functioning simply as communities defined by the abundance of a single microbe. However, the observational design of the present study does not establish whether the *Gardnerella*-predominant state precedes, results from it, or is otherwise causally related to UTI. Our findings therefore support an association between this community configuration and UTI during pregnancy but do not establish a causal relationship.

The pronounced taxonomic differentiation among the three community states supports their interpretation as distinct ecological configurations. Community state explained 43.2% of the variation in bacterial composition by Bray–Curtis PERMANOVA. Although significant differences in multivariate dispersion indicate that within-group variability also contributed to the observed pattern, the magnitude of the association demonstrates substantial compositional differentiation among the *L. crispatus*-, *Gardnerella*-, and *L. iners*-predominant communities.

### Functional differentiation among urogenital community states

A central finding of this study was the strong correspondence between taxonomic community structure and predicted functional potential. Both KO- and KEGG pathway-level profiles differed significantly among the three community states, with community state explaining approximately one-quarter of predicted functional variation. These findings indicate that the identified taxonomic configurations are accompanied by distinct functional gene repertoires rather than representing differences in taxonomic composition alone.

The *Gardnerella*-predominant community exhibited the broadest functional differentiation, with 57 significantly enriched KEGG pathways compared with 11 and 10 in the *L. crispatus-* and *L. iners*-predominant communities, respectively. Enriched pathways included functions associated with amino acid biosynthesis, fatty acid metabolism, sulfur metabolism, vitamin B6 metabolism, 2-oxocarboxylic acid metabolism, phosphonate and phosphinate metabolism, and biofilm formation. The greater functional breadth of these communities may partly reflect their greater bacterial diversity, because a more taxonomically diverse community can collectively encode a broader repertoire of metabolic functions. Accordingly, these pathway-level differences should be interpreted as properties of the overall community and not as functions encoded or expressed specifically by *Gardnerella*.

The *L. crispatus*-predominant community displayed a smaller but distinct set of enriched pathways, including riboflavin metabolism, carotenoid biosynthesis, primary and secondary bile acid biosynthesis, and ascorbate and aldarate metabolism. The riboflavin signal is particularly interesting in the context of previous comparative genomic work. France et al. reported genes that encode the riboflavin biosynthesis pathway in the *L. crispatus* core genome but not in the *L. iners* core genome, supporting functional differentiation between these two *Lactobacillus* species (France et al., 2016). Nevertheless, pathway labels such as bile acid biosynthesis and carotenoid biosynthesis should be interpreted cautiously because KEGG pathways may contain genes shared across broader metabolic processes, and pathway enrichment does not necessarily indicate completion or activity of the entire annotated pathway.

The *L. iners*-predominant community also exhibited a distinct functional profile, including enrichment of pathways annotated as bacterial chemotaxis, flagellar assembly, glycan degradation, aminobenzoate degradation, styrene degradation, sulfur relay systems, and penicillin and cephalosporin biosynthesis. The enrichment of flagellar assembly and bacterial chemotaxis pathways is notable because *L. iners* itself is considered non-motile (Brockmann, 2026). These results therefore should not be interpreted as evidence of motility by *L. iners*. Rather, they represent community-level functional signals that may originate from other organisms present within *L. iners*-predominated communities or from KOs shared with functions represented in these KEGG pathways. Taxonomic attribution of underlying genes will be required to determine which microbes contribute to these signals.

The predicted functional distinction between *L. crispatus*- and *L. iners*-predominant communities was apparent despite their relative proximity in functional ordination space. This finding is consistent with previous comparative genomic studies demonstrating substantial differences in genome size and metabolic repertoire between *L. crispatus* and *L. iners* (France et al., 2016). It also agrees with a shotgun metagenomic study of oral, gut, and vaginal microbiomes in pregnancy that demonstrated fine-scale taxonomic and functional variation among *Lactobacillus*- and *Gardnerella*-associated communities (Goltsman et al., 2018). Thus, predominance by members of the same genus does not necessarily imply equivalent community-level functional potential.

Integration of taxonomic and functional profiles provided additional evidence of coordinated community-level functional differences. Taxon–pathway associations were organized around highly connected pathways involving central carbon and amino acid metabolism. Multiple *Lactobacillus* species were positively associated with the phosphotransferase system (PTS), whereas several *Gardnerella* species were positively associated with amino acid biosynthetic and central carbon metabolic pathways. These relationships represent co-variation between taxonomic and functional abundances but do not establish that the associated genes or pathways originated from the correlated taxa, were transcriptionally active, or resulted in metabolic activity.

The predominance of carbohydrate transport and amino acid metabolism among these associations may reflect differences in nutrient acquisition strategies across urogenital community configurations. Recent paired metagenomic and metabolomic analyses have similarly demonstrated relationships between urinary microbial composition and biochemical profiles, including species-specific microbe–metabolite associations in recurrent UTI (Neugent et al., 2025). Direct metabolomic or transcriptomic measurements will be required to determine whether the functional potential identified here translates into differences in metabolic activity.

Functional composition was substantially more strongly associated with microbial community state than with UTI status. Although UTI during pregnancy showed a nominal association with pathway-level composition, this relationship did not remain significant after correction for multiple testing. Furthermore, after accounting for community state, UTI explained only an additional 2.4% of functional variation and was no longer significant, whereas community state remained strongly associated with functional composition. These findings suggest that functional differences associated with UTI may partly reflect the underlying microbial community configuration rather than UTI status itself.

### Genome-resolved diversity of the pregnancy-associated urogenital microbiome

Long-read metagenomic assembly provided an additional level of resolution by enabling recovery of 415 MAGs, including 29 genomes meeting the completeness and contamination thresholds applied to high-quality MAGs. Frequently recovered high-quality genomes included *F. vaginae, L. iners*, and multiple *Gardnerella* species, demonstrating that several microbes, characteristic of the community-level profiles, could also be reconstructed at the genome level.

Phylogenomic and orthologous gene analyses demonstrated both taxonomic conservation and substantial variation in accessory gene content. Genomes assigned to the same species generally clustered together and shared a large proportion of orthologous gene families, whereas differences in accessory gene repertoires were apparent both within and among taxa. The diversity observed among *Gardnerella* genomes is consistent with previous studies demonstrating substantial genomic heterogeneity within this genus and emphasizes the limitations of interpreting *Gardnerella* as a functionally uniform taxonomic group (Ahmed et al., 2012). Such studies found exceptionally high genomic diversity among 17 *Gardnerella* isolates, including substantial differences in gene content and distinct genomic clades (Yeoman et al., 2010). These studies found extensive genomic heterogeneity, rearrangements, mobile elements, and lateral gene transfer, with genomic differences corresponding to differences in predicted metabolic potential.

Exploratory annotation-based screening further identified differences in the distribution of putative antibiotic resistance-associated functions among recovered genomes. Multidrug efflux-associated annotations were broadly distributed, whereas annotations potentially related to tetracycline, aminoglycoside, β-lactam, and macrolide resistance were more restricted across taxa. Annotation-based screening also found differences in the distribution of mobile genetic element-, defense-, and toxin-associated functions. However, these observations should be interpreted cautiously. The resistance-associated analysis was based on Prokka functional annotations and keyword classification rather than comparison against a dedicated curated antimicrobial resistance database. Consequently, the identified proteins represent candidate resistance-associated functions rather than confirmed resistance genes, and no inference regarding phenotypic antimicrobial susceptibility can be made from these data.

### Urogenital virome differs across bacterial community states

The urogenital virome remains substantially less characterized than its bacterial counterpart, particularly during pregnancy. In the present study, viral sequences were detected in the majority of samples, and most classified sequences belonged to Caudoviricetes, consistent with a virome predominantly composed of bacteriophages. However, most viral sequences could not be classified at the family level, emphasizing the limited representation and taxonomic resolution of human-associated viral sequences in current reference databases.

Previous studies have similarly demonstrated extensive bacteriophage diversity in the female urinary tract. Santiago-Rodriguez et al. detected diverse viral communities in urine from individuals with and without UTI, with most identifiable viral sequences corresponding to bacteriophages (Santiago-Rodriguez et al., 2015). Garretto et al. demonstrated that partial and complete viral genomes, including bacteriophages, could be reconstructed directly from female bladder metagenomes (Garretto et al., 2018). Miller-Ensminger et al. identified 457 phage sequences across 181 bacterial genomes representative of the female urinary microbiota, with 86% of genomes containing at least one predicted phage sequence and most phages showing little or no similarity to sequences in public databases (Miller-Ensminger et al., 2018). Collectively, these studies indicate that bacteriophages are common, diverse, and incompletely characterized components of female urogenital microbial ecosystems.

Within this context, the substantially greater viral richness observed in *Gardnerella*-predominated communities is particularly notable. These communities contained, on average, 27.5 viral contigs per sample compared with 9.7 and 6.8 in *L. crispatus*- and *L. iners*-predominated communities, respectively. Viral community composition also differed according to bacterial community state, which explained 6.9% of the variation in viral taxonomic profiles. Together, these findings indicate that the ecological differences distinguishing the three bacterial community states extend to the viral component of the urogenital microbiome.

The increased viral richness of *Gardnerella*-predominant samples may partly reflect their greater bacterial diversity. More diverse bacterial communities potentially provide a broader range of hosts for bacteriophages and prophages, although direct phage–host relationships could not be determined from the present data. Conversely, both *Lactobacillus*-predominant community states exhibited relatively low viral richness despite their distinct bacterial and functional characteristics. This pattern suggests that viral richness in this cohort may be more closely related to the polymicrobial structure accompanying *Gardnerella* predominance than to differences between the two *Lactobacillus*-predominant states.

Clinical and demographic variables were not significantly associated with viral richness after correction for multiple testing. This observation, together with the strong differences according to bacterial community state, further supports a close relationship between bacterial and viral community structure. However, the highly individualized distribution of viral sequences and limited taxonomic classification constrain biological interpretation. Larger longitudinal studies will be required to determine whether changes in the urogenital virome precede, accompany, or result from alterations in bacterial community structure during pregnancy.

### Strengths and limitations

A major strength of this study is the application of long-read shotgun metagenomics to characterize multiple dimensions of the pregnancy-associated urogenital microbiome within the same cohort. This approach enabled species-level taxonomic profiling, functional characterization, genome reconstruction, comparative genomic analyses, and virome characterization, providing information that cannot be obtained from 16S rRNA gene sequencing alone. However, several limitations should temper interpretation. First, because samples were collected as midstream voided urine, the microbial profiles likely reflect the broader urogenital microbial environment rather than the bladder alone. Microorganisms originating from the urethra, vulva, vagina, and surrounding skin may contribute to the detected profiles. Although this limits anatomical localization of individual microbes, it also provides an integrated view of the urogenital microbial environment encountered in clinically collected urine during pregnancy. Second, the cross-sectional design supports association but not causation, particularly for the relationship between community state and UTI during pregnancy. Third, metagenomic functional profiles represent genomic potential rather than gene expression or metabolic activity. Similarly, taxon–function correlations indicate co-variation and cannot directly assign functions to specific organisms. Metatranscriptomic, metabolomic, and targeted genome-resolved analyses will be necessary to establish which community members encode and actively express the identified functions. Fourth, urine is a low-biomass sample; thus, a substantial proportion of sequencing reads originated from host DNA or reagent-derived background ("kitome") contamination. Although rigorous quality-control and decontamination procedures minimized these effects, they inevitably reduced the number of microbial reads available for downstream analyses, likely limiting the recovery of low-abundance microbes, viral genomes, and high-quality MAGs. Finally, viral characterization remains limited by incomplete reference databases and the highly individualized nature of the detected viral sequences. Likewise, the screening of putative antibiotic resistance-associated proteins was exploratory and based on general functional annotations rather than a dedicated AMR prediction framework. These analyses should therefore be viewed as hypothesis-generating components of the study. Despite these limitations, the present study demonstrates that long-read shotgun metagenomics can generate biologically meaningful information across multiple layers of the urogenital microbiome and provides a foundation for future multi-omics investigations of pregnancy-associated urinary health.

## CONCLUSIONS

Overall, our data show that pregnancy-associated midstream voided urine samples contain distinct urogenital community states that differ in both composition and inferred functional potential. The *Gardnerella*-predominant state was associated with higher UTI frequency during pregnancy, but the biological basis and temporal direction of this relationship remain unresolved. Future work integrating longitudinal sampling, direct bladder-adjacent sampling, metabolomics, and cultivation-based experiments will be needed to test the ecological mechanisms suggested by these metagenomic observations.

## AUTHOR STATEMENTS

### Author contributions

Conceptualization: MFN, AJW

Wet Lab Analysis: BF, MJR

Data Curation: MFN

Formal Analysis: MFN

Funding Acquisition: AJW

Investigation: All authors

Project administration: NJP, AJW

Resources: NJP, AJW

Supervision: NJP, AJW

Visualization: MFN

Writing – original draft: MFN

Writing – review & editing: All authors

### Ethical approval

The study protocol was approved by the Institutional Review Board [LU 216104].

### Consent for publication

Not applicable

## Acknowledgments

We would like to acknowledge the Loyola clinicians who recruited participants and collected samples, the participants themselves for consenting to the various studies, members of the Wolfe lab who processed the samples, and Mike Fischer for his financial support.

## REFERENCE

Ahmed, A., Earl, J., Retchless, A., Hillier, S. L., Rabe, L. K., Cherpes, T. L., et al. (2012). Comparative Genomic Analyses of 17 Clinical Isolates of Gardnerella vaginalis Provide Evidence of Multiple Genetically Isolated Clades Consistent with Subspeciation into Genovars. J. Bacteriol. 194, 3922–3937. doi: 10.1128/JB.00056-12

Aitchison, J. (1982). The Statistical Analysis of Compositional Data. J. R. Stat. Soc. Ser. B Stat. Methodol. 44, 139–160. doi: 10.1111/j.2517-6161.1982.tb01195.x

American College of Obstetricians and Gynecologists’ Committee on Clinical Consensus–Obstetrics (2023). Urinary Tract Infections in Pregnant Individuals. Obstet. Gynecol. 142, 435–445. doi: 10.1097/AOG.0000000000005269

Anderson, M. J. (2001). A new method for non-parametric multivariate analysis of variance. Austral Ecol. 1, 32–46. doi: 10.1111/j.1442-9993.2001.01070.pp.x

Anderson, M. J. (2006). Distance-Based Tests for Homogeneity of Multivariate Dispersions. Biometrics 62, 245–253. doi: 10.1111/j.1541-0420.2005.00440.x

Benjamini, Y., and Hochberg, Y. (1995). Controlling the False Discovery Rate: A Practical and Powerful Approach to Multiple Testing. J. R. Stat. Soc. Ser. B Stat. Methodol. 57, 289–300. doi: 10.1111/j.2517-6161.1995.tb02031.x

Brockmann, E. (2026). “Lactobacillus,” in Bergey’s Manual of Systematics of Archaea and Bacteria, eds. M. E. Trujillo, S. Dedysh, P. DeVos, B. Hedlund, P. Kämpfer, F. A. Rainey, et al. (Wiley), 1–56. doi: 10.1002/9781118960608.gbm00604.pub2

Brubaker, L., Chai, T. C., Horsley, H., Khasriya, R., Moreland, R. B., and Wolfe, A. J. (2023). Tarnished gold—the “standard” urine culture: reassessing the characteristics of a criterion standard for detecting urinary microbes. Front. Urol. 3, 1206046. doi: 10.3389/fruro.2023.1206046

Brubaker, L., Gourdine, J.-P. F., Siddiqui, N. Y., Holland, A., Halverson, T., Limeria, R., et al. (2021). Forming Consensus To Advance Urobiome Research. mSystems 6. doi: 10.1128/msystems.01371-20

Buchfink, B., Reuter, K., and Drost, H.-G. (2021). Sensitive protein alignments at tree-of-life scale using DIAMOND. Nat. Methods 18, 366–368. doi: 10.1038/s41592-021-01101-x

Camacho, C., Coulouris, G., Avagyan, V., Ma, N., Papadopoulos, J., Bealer, K., et al. (2009). BLAST+: architecture and applications. BMC Bioinformatics 10, 421. doi: 10.1186/1471-2105-10-421

Camargo, A. P., Roux, S., Schulz, F., Babinski, M., Xu, Y., Hu, B., et al. (2024). Identification of mobile genetic elements with geNomad. Nat. Biotechnol. 42, 1303–1312. doi: 10.1038/s41587-023-01953-y

Cantalapiedra, C. P., Hernández-Plaza, A., Letunic, I., Bork, P., and Huerta-Cepas, J. (2021). eggNOG-mapper v2: Functional Annotation, Orthology Assignments, and Domain Prediction at the Metagenomic Scale. Mol. Biol. Evol. 38, 5825–5829. doi: 10.1093/molbev/msab293

Csárdi G, Nepusz T, Traag V, Horvát Sz, Zanini F, Noom D, Müller K (2026). igraph: Network Analysis and Visualization in R. doi: 10.5281/zenodo.7682609

Danecek, P., Bonfield, J. K., Liddle, J., Marshall, J., Ohan, V., Pollard, M. O., et al. (2021). Twelve years of SAMtools and BCFtools. GigaScience 10, giab008. doi: 10.1093/gigascience/giab008

De Coster, W., D’Hert, S., Schultz, D. T., Cruts, M., and Van Broeckhoven, C. (2018). NanoPack: visualizing and processing long-read sequencing data. Bioinformatics 34, 2666–2669. doi: 10.1093/bioinformatics/bty149

Dhillon, N., Kothawala, A., Khemmani, M., Nwachokor, J., Adams, W., Wolfe, A. J., et al. (2026). The Peezy Device Compared With Midstream Clean Catch in an Obstetric Population: A Randomized Controlled Trial. Am. J. Perinatol., a-2884–8743. doi: 10.1055/a-2884-8743

Edgar, R. C. (2004). MUSCLE: multiple sequence alignment with high accuracy and high throughput. Nucleic Acids Res. 32, 1792–1797. doi: 10.1093/nar/gkh340

Eren, A. M., Kiefl, E., Shaiber, A., Veseli, I., Miller, S. E., Schechter, M. S., et al. (2020). Community-led, integrated, reproducible multi-omics with anvi’o. Nat. Microbiol. 6, 3–6. doi: 10.1038/s41564-020-00834-3

Ferneyhough, B., Roddis, M., Millington, S., Quirk, J., Clements, C., West, S., et al. (2025). A highly accurate nanopore-based sequencing workflow for culture and PCR-free microbial metagenomic profiling of urogenital samples. BMC Urol. 25, 41. doi: 10.1186/s12894-025-01723-9

France, M. T., Ma, B., Gajer, P., Brown, S., Humphrys, M. S., Holm, J. B., et al. (2020). VALENCIA: a nearest centroid classification method for vaginal microbial communities based on composition. Microbiome 8, 166. doi: 10.1186/s40168-020-00934-6

France, M. T., Mendes-Soares, H., and Forney, L. J. (2016). Genomic Comparisons of Lactobacillus crispatus and Lactobacillus iners Reveal Potential Ecological Drivers of Community Composition in the Vagina. Appl. Environ. Microbiol. 82, 7063–7073. doi: 10.1128/AEM.02385-16

Garretto, A., Thomas-White, K., Wolfe, A. J., and Putonti, C. (2018). Detecting viral genomes in the female urinary microbiome. J. Gen. Virol. 99, 1141–1146. doi: 10.1099/jgv.0.001097

Gilbert, N. M., O’Brien, V. P., and Lewis, A. L. (2017). Transient microbiota exposures activate dormant Escherichia coli infection in the bladder and drive severe outcomes of recurrent disease. PLOS Pathog. 13, e1006238. doi: 10.1371/journal.ppat.1006238

Giron-Villalobos, D., Schultsz, C., Wolfe, A. J., De Vos, M. G. J., Van De Wijgert, J. H. H. M., Dutilh, B. E., et al. (2026). Cross-study metagenomics analysis reveals distinct microbial signatures of urinary tract infections. doi: 10.64898/2026.06.17.732905

Goltsman, D. S. A., Sun, C. L., Proctor, D. M., DiGiulio, D. B., Robaczewska, A., Thomas, B. C., et al. (2018). Metagenomic analysis with strain-level resolution reveals fine-scale variation in the human pregnancy microbiome. Genome Res. 28, 1467–1480. doi: 10.1101/gr.236000.118

Grattafiori, A., Dubey, A., Jauhri, A., Pandey, A., Kadian, A., Al-Dahle, A., et al. (2024). The Llama 3 Herd of Models. doi: 10.48550/arXiv.2407.21783

Gu, Z., Eils, R., and Schlesner, M. (2016). Complex heatmaps reveal patterns and correlations in multidimensional genomic data. Bioinformatics 32, 2847–2849. doi: 10.1093/bioinformatics/btw313

Hilt, E. E., McKinley, K., Pearce, M. M., Rosenfeld, A. B., Zilliox, M. J., Mueller, E. R., et al. (2014). Urine Is Not Sterile: Use of Enhanced Urine Culture Techniques To Detect Resident Bacterial Flora in the Adult Female Bladder. J. Clin. Microbiol. 52, 871–876. doi: 10.1128/JCM.02876-13

Jain, C., Rodriguez-R, L. M., Phillippy, A. M., Konstantinidis, K. T., and Aluru, S. (2018). High throughput ANI analysis of 90K prokaryotic genomes reveals clear species boundaries. Nat. Commun. 9, 5114. doi: 10.1038/s41467-018-07641-9

Kanehisa, M., Furumichi, M., Sato, Y., Matsuura, Y., and Ishiguro-Watanabe, M. (2025). KEGG: biological systems database as a model of the real world. Nucleic Acids Res. 53, D672–D677. doi: 10.1093/nar/gkae909

Kang, D. D., Li, F., Kirton, E., Thomas, A., Egan, R., An, H., et al. (2019). MetaBAT 2: an adaptive binning algorithm for robust and efficient genome reconstruction from metagenome assemblies. PeerJ 7, e7359. doi: 10.7717/peerj.7359

Kaufman, L., and Rousseeuw, P. J. (1990). Finding Groups in Data: An Introduction to Cluster Analysis., 1st Edn. Wiley. doi: 10.1002/9780470316801

Kim, C., Pongpanich, M., and Porntaveetus, T. (2024). Unraveling metagenomics through long-read sequencing: a comprehensive review. J. Transl. Med. 22, 111. doi: 10.1186/s12967-024-04917-1

Koedooder, R., Maghdid, D. M., Beckers, N. G. M., Schoenmakers, S., Kok, D. J., and Laven, J. S. E. (2021). Dynamics of the urinary microbiome in pregnancy and the coincidental predictive value of the microbiota for IVF/IVF–ICSI outcome. Reprod. Biomed. Online 43, 871–879. doi: 10.1016/j.rbmo.2021.07.018

Kolmogorov, M., Bickhart, D. M., Behsaz, B., Gurevich, A., Rayko, M., Shin, S. B., et al. (2020). metaFlye: scalable long-read metagenome assembly using repeat graphs. Nat. Methods 17, 1103–1110. doi: 10.1038/s41592-020-00971-x

Kursa, M. B., and Rudnicki, W. R. (2010). Feature Selection with the **Boruta** Package. J. Stat. Softw. 36, 1–13. doi: 10.18637/jss.v036.i11

Lechner, M., Findeiß, S., Steiner, L., Marz, M., Stadler, P. F., and Prohaska, S. J. (2011). Proteinortho: Detection of (Co-)orthologs in large-scale analysis. BMC Bioinformatics 12, 124. doi: 10.1186/1471-2105-12-124

Li, H. (2018). Minimap2: pairwise alignment for nucleotide sequences. Bioinformatics 34, 3094–3100. doi: 10.1093/bioinformatics/bty191

Lin, H., and Peddada, S. D. (2020). Analysis of compositions of microbiomes with bias correction. Nat. Commun. 11, 3514. doi: 10.1038/s41467-020-17041-7

Love, M. I., Huber, W., and Anders, S. (2014). Moderated estimation of fold change and dispersion for RNA-seq data with DESeq2. Genome Biol. 15, 550. doi: 10.1186/s13059-014-0550-8

Lu, J., Breitwieser, F. P., Thielen, P., and Salzberg, S. L. (2025). Bracken: estimating species abundance in metagenomics data. PeerJ Comput. Sci. doi: 10.7717/peerj-cs.104

Ma, B., Forney, L. J., and Ravel, J. (2012). Vaginal Microbiome: Rethinking Health and Disease. Annu. Rev. Microbiol. 66, 371–389. doi: 10.1146/annurev-micro-092611-150157

McMurdie, P. J., and Holmes, S. (2013). phyloseq: An R Package for Reproducible Interactive Analysis and Graphics of Microbiome Census Data. PLoS ONE 8, e61217. doi: 10.1371/journal.pone.0061217

Miller-Ensminger, T., Garretto, A., Brenner, J., Thomas-White, K., Zambom, A., Wolfe, A. J., et al. (2018). Bacteriophages of the Urinary Microbiome. J. Bacteriol. 200. doi: 10.1128/JB.00738-17

Mora-Vargas, C. D., Grobeisen-Duque, O., Villavicencio-Carrisoza, O., Aguilar-Ayala, D. A., Castellanos-Diaz, O., Martinez-Salazar, M. G., et al. (2025). Urobiome Signatures of Recurrent Urinary Tract Infections in Adolescent Pregnancy: A Longitudinal Study. Microorganisms 13, 2406. doi: 10.3390/microorganisms13102406

Moreland, R. B., Brubaker, L., Tinawi, L., and Wolfe, A. J. (2025). Rapid and accurate testing for urinary tract infection: new clothes for the emperor. Clin. Microbiol. Rev. 38, e00129–24. doi: 10.1128/cmr.00129-24

Neugent, M. L., Neha V. Hulyalkar, Saenz, C. N., Zimmern, P. E., Shulaev, V., and De Nisco, N. J. (2025). Urinary biochemical ecology reveals microbiome-metabolite interactions and metabolic markers of recurrent urinary tract infection. Npj Biofilms Microbiomes 11, 216. doi: 10.1038/s41522-025-00844-1

Oksanen, J., Simpson, G. L., Blanchet, F. G., Roeland Kindt, Legendre, P., Minchin, P. R., et al. (2022). vegan: Community Ecology Package. doi: 10.32614/CRAN.package.vegan

Ollberding, N. J., Völgyi, E., Macaluso, M., Kumar, R., Morrow, C., Tylavsky, F. A., et al. (2016). Urinary Microbiota Associated with Preterm Birth: Results from the Conditions Affecting Neurocognitive Development and Learning in Early Childhood (CANDLE) Study. PLOS ONE 11, e0162302. doi: 10.1371/journal.pone.0162302

Olm, M. R., Brown, C. T., Brooks, B., and Banfield, J. F. (2017). dRep: a tool for fast and accurate genomic comparisons that enables improved genome recovery from metagenomes through de-replication. ISME J. 11, 2864–2868. doi: 10.1038/ismej.2017.126

Parks, D. H., Imelfort, M., Skennerton, C. T., Hugenholtz, P., and Tyson, G. W. (2015). CheckM: assessing the quality of microbial genomes recovered from isolates, single cells, and metagenomes. Genome Res. 25, 1043–1055. doi: 10.1101/gr.186072.114

Pearce, M. M., Hilt, E. E., Rosenfeld, A. B., Zilliox, M. J., Thomas-White, K., Fok, C., et al. (2014). The Female Urinary Microbiome: a Comparison of Women with and without Urgency Urinary Incontinence. mBio 5, e01283–14. doi: 10.1128/mBio.01283-14

Pedersen, T. L. (2025). ggraph: An Implementation of Grammar of Graphics for Graphs and Networks. Available at: https://CRAN.R-project.org/package=ggraph

Petrova, M. I., Reid, G., Vaneechoutte, M., and Lebeer, S. (2017). Lactobacillus iners : Friend or Foe? Trends Microbiol. 25, 182–191. doi: 10.1016/j.tim.2016.11.007

Price, M. N., Dehal, P. S., and Arkin, A. P. (2010). FastTree 2 – Approximately Maximum-Likelihood Trees for Large Alignments. PLoS ONE 5, e9490. doi: 10.1371/journal.pone.0009490

R Core Team (2022). R: A Language and Environment for Statistical Computing. Available at: https://www.r-project.org/

Ravel, J., Gajer, P., Abdo, Z., Schneider, G. M., Koenig, S. S. K., McCulle, S. L., et al. (2011). Vaginal microbiome of reproductive-age women. Proc. Natl. Acad. Sci. 108, 4680–4687. doi: 10.1073/pnas.1002611107

Rousseeuw, P. J. (1987). Silhouettes: A graphical aid to the interpretation and validation of cluster analysis. J. Comput. Appl. Math. 20, 53–65. doi: 10.1016/0377-0427(87)90125-7

Santiago-Rodriguez, T. M., Ly, M., Bonilla, N., and Pride, D. T. (2015). The human urine virome in association with urinary tract infections. Front. Microbiol. 6. doi: 10.3389/fmicb.2015.00014

Seemann, T. (2014). Prokka: rapid prokaryotic genome annotation. Bioinformatics 30, 2068–2069. doi: 10.1093/bioinformatics/btu153

Sung, J., Larsen, P., Halverson, T. M., Waters, T. P., Goodman, J. R., and Wolfe, A. J. (2024). First trimester “clean catch” urine and vaginal swab sample distinct microbiological niches. Microbiol. Spectr. 12, e02638–23. doi: 10.1128/spectrum.02638-23

US Preventive Services Task Force, Owens, D. K., Davidson, K. W., Krist, A. H., Barry, M. J., Cabana, M., et al. (2019). Screening for Asymptomatic Bacteriuria in Adults: US Preventive Services Task Force Recommendation Statement. JAMA 322, 1188. doi: 10.1001/jama.2019.13069

Van De Wijgert, J. H. H. M., and Jespers, V. (2017). The global health impact of vaginal dysbiosis. Res. Microbiol. 168, 859–864. doi: 10.1016/j.resmic.2017.02.003

Wolfe, A. J., Toh, E., Shibata, N., Rong, R., Kenton, K., FitzGerald, M., et al. (2012). Evidence of Uncultivated Bacteria in the Adult Female Bladder. J. Clin. Microbiol. 50, 1376–1383. doi: 10.1128/JCM.05852-11

Wood, D. E., Lu, J., and Langmead, B. (2019). Improved metagenomic analysis with Kraken 2. Genome Biol. 20, 257. doi: 10.1186/s13059-019-1891-0

Xu, S., Dai, Z., Guo, P., Fu, X., Liu, S., Zhou, L., et al. (2021). ggtreeExtra: Compact Visualization of Richly Annotated Phylogenetic Data. Mol. Biol. Evol. 38, 4039–4042. doi: 10.1093/molbev/msab166

Yeoman, C. J., Yildirim, S., Thomas, S. M., Durkin, A. S., Torralba, M., Sutton, G., et al. (2010). Comparative Genomics of Gardnerella vaginalis Strains Reveals Substantial Differences in Metabolic and Virulence Potential. PLoS ONE 5, e12411. doi: 10.1371/journal.pone.0012411

Yu, G., Smith, D. K., Zhu, H., Guan, Y., and Lam, T. T. (2017). GGTREE : an R package for visualization and annotation of phylogenetic trees with their covariates and other associated data. Methods Ecol. Evol. 8, 28–36. doi: 10.1111/2041-210X.12628

