## Supplemental Material for "Long-read shotgun metagenomics identifies distinct urogenital microbial community states associated with functional potential and urinary tract infection during pregnancy"

### Supplemental Information

**Supplemental Table 1. Participant Demographic and Clinical Characteristics**

| | |
|---|---|
| **Age, years** (n=129) [a] | 29.5 ± 5.3 |
| **Race** (n=130) | |
| White | 53 (40.8%) |
| Black | 27 (20.8%) |
| Asian | 6 (4.6%) |
| Native American | 2 (1.5%) |
| Other | 40 (30.8%) |
| Prefer not to answer | 2 (1.5%) |
| **Pre-pregnancy BMI, kg/m²** (n=128) | 29.9 ± 7.1 |
| **BMI at collection, kg/m²** (n=101) | 33.5 ± 6.2 |
| **Gestational age at sample collection, days** (n=126) | 69.5 ± 21.2 |
| **Gestational age at delivery, days** (n=80) | 260.7 ± 29.5 |
| **Gravidity** (n=129) | 2 (IQR 1–3) |
| **UTI during pregnancy** (n=104) | 16 |

[a] Denominators (n) vary by variable due to missing data. Percentages are calculated based on the number of non-missing observations for each variable, as noted.

Abbreviations: BMI, body mass index; IQR, interquartile range; UTI, urinary tract infection.

**Supplemental Table 2. Differentially abundant species identified by ANCOM-BC2 across pairwise comparisons of the three urinary microbial community clusters.**

| Comparison | Species | LogFoldChange | q_value | Direction |
|---|---|---|---|---|
| Cluster 2: *Gardnerella* vs Cluster 1: *L. crispatus* | *Lactobacillus crispatus* | -8.7808187 | 4.31E-55 | Enriched in Cluster 1: *L. crispatus* |
| Cluster 2: *Gardnerella* vs Cluster 1: *L. crispatus* | *Lactobacillus helveticus* | -3.7919784 | 2.69E-16 | Enriched in Cluster 1: *L. crispatus* |
| Cluster 2: *Gardnerella* vs Cluster 1: *L. crispatus* | *Lactobacillus kefiranofaciens* | -4.0549633 | 3.27E-16 | Enriched in Cluster 1: *L. crispatus* |
| Cluster 2: *Gardnerella* vs Cluster 1: *L. crispatus* | *Gardnerella greenwoodii* | 5.22856617 | 1.15E-13 | Enriched in Cluster 2: *Gardnerella* |
| Cluster 2: *Gardnerella* vs Cluster 1: *L. crispatus* | *Gardnerella swidsinskii* | 5.20685528 | 7.97E-11 | Enriched in Cluster 2: *Gardnerella* |
| Cluster 2: *Gardnerella* vs Cluster 1: *L. crispatus* | *Gardnerella vaginalis* | 4.85276642 | 2.49E-10 | Enriched in Cluster 2: *Gardnerella* |
| Cluster 2: *Gardnerella* vs Cluster 1: *L. crispatus* | *Lactobacillus johnsonii* | -3.0832883 | 7.93E-10 | Enriched in Cluster 1: *L. crispatus* |
| Cluster 2: *Gardnerella* vs Cluster 1: *L. crispatus* | *Gardnerella piotii* | 4.09831377 | 1.65E-08 | Enriched in Cluster 2: *Gardnerella* |
| Cluster 2: *Gardnerella* vs Cluster 1: *L. crispatus* | *Gardnerella pickettii* | 4.15108126 | 2.95E-08 | Enriched in Cluster 2: *Gardnerella* |
| Cluster 2: *Gardnerella* vs Cluster 1: *L. crispatus* | *Aerococcus christensenii* | 3.23873983 | 7.82E-08 | Enriched in Cluster 2: *Gardnerella* |
| Cluster 2: *Gardnerella* vs Cluster 1: *L. crispatus* | *Gardnerella leopoldii* | 4.17565527 | 2.30E-07 | Enriched in Cluster 2: *Gardnerella* |

| | | | | |
|---|---|---|---|---|
| Cluster 2: *Gardnerella* vs Cluster 1: *L. crispatus* | *Mageeibacillus indolicus* | 2.78977107 | 5.81E-07 | Enriched in Cluster 2: *Gardnerella* |
| Cluster 2: *Gardnerella* vs Cluster 1: *L. crispatus* | *Lactobacillus gasseri* | -3.5442632 | 8.69E-07 | Enriched in Cluster 1: *L. crispatus* |
| Cluster 2: *Gardnerella* vs Cluster 1: *L. crispatus* | *Hoylesella buccalis* | 2.42791746 | 2.98E-05 | Enriched in Cluster 2: *Gardnerella* |
| Cluster 2: *Gardnerella* vs Cluster 1: *L. crispatus* | *Mobiluncus curtisii* | 1.81133502 | 0.00265289 | Enriched in Cluster 2: *Gardnerella* |
| Cluster 2: *Gardnerella* vs Cluster 1: *L. crispatus* | *Staphylococcus epidermidis* | 1.71803022 | 0.00407839 | Enriched in Cluster 2: *Gardnerella* |
| Cluster 2: *Gardnerella* vs Cluster 1: *L. crispatus* | *Peptoniphilus harei* | 1.89363672 | 0.00654528 | Enriched in Cluster 2: *Gardnerella* |
| Cluster 2: *Gardnerella* vs Cluster 1: *L. crispatus* | *Streptococcus agalactiae* | 1.47150284 | 0.01139402 | Enriched in Cluster 2: *Gardnerella* |
| Cluster 2: *Gardnerella* vs Cluster 1: *L. crispatus* | *Streptococcus constellatus* | 1.42975431 | 0.01385764 | Enriched in Cluster 2: *Gardnerella* |
| Cluster 2: *Gardnerella* vs Cluster 1: *L. crispatus* | *Anaerococcus vaginalis* | 1.43408563 | 0.01791357 | Enriched in Cluster 2: *Gardnerella* |
| Cluster 2: *Gardnerella* vs Cluster 1: *L. crispatus* | *Anaerococcus prevotii* | 1.39080497 | 0.02185058 | Enriched in Cluster 2: *Gardnerella* |
| Cluster 2: *Gardnerella* vs Cluster 1: *L. crispatus* | *Lactobacillus jensenii* | -1.9026784 | 0.02333966 | Enriched in Cluster 1: *L. crispatus* |
| Cluster 2: *Gardnerella* vs Cluster 1: *L. crispatus* | *Anaerococcus obesiensis* | 1.2616402 | 0.03440085 | Enriched in Cluster 2: *Gardnerella* |
| Cluster 2: *Gardnerella* vs Cluster 1: *L. crispatus* | *Anaerococcus mediterraneensis* | 1.35817013 | 0.04019766 | Enriched in Cluster 2: *Gardnerella* |

| | | | | |
|---|---|---|---|---|
| Cluster 3: *L. iners* vs Cluster 1: *L. crispatus* | *Lactobacillus crispatus* | -8.9578014 | 2.77E-53 | Enriched in Cluster 1: *L. crispatus* |
| Cluster 3: *L. iners* vs Cluster 1: *L. crispatus* | *Lactobacillus helveticus* | -3.5423759 | 1.93E-13 | Enriched in Cluster 1: *L. crispatus* |
| Cluster 3: *L. iners* vs Cluster 1: *L. crispatus* | *Lactobacillus kefiranofaciens* | -3.8100988 | 1.93E-13 | Enriched in Cluster 1: *L. crispatus* |
| Cluster 3: *L. iners* vs Cluster 1: *L. crispatus* | *Lactobacillus iners* | 5.01613067 | 2.40E-10 | Enriched in Cluster 3: *L. iners* |
| Cluster 3: *L. iners* vs Cluster 1: *L. crispatus* | *Lactobacillus johnsonii* | -3.0553969 | 6.31E-09 | Enriched in Cluster 1: *L. crispatus* |
| Cluster 3: *L. iners* vs Cluster 1: L. crispatus | *Aerococcus christensenii* | 2.92358191 | 7.36E-06 | Enriched in Cluster 3: *L. iners* |
| Cluster 3: *L. iners* vs Cluster 1: *L. crispatus* | *Lactobacillus gasseri* | -2.7012979 | 0.00109283 | Enriched in Cluster 1: *L. crispatus* |
| Cluster 3: *L. iners* vs Cluster 1: *L. crispatus* | *Mageeibacillus indolicus* | 1.47869001 | 0.044862 | Enriched in Cluster 3: *L. iners* |
| Cluster 3: *L. iners* vs Cluster 2: *Gardnerella* | *Fannyhessea vaginae* | -4.4702863 | 4.91E-09 | Enriched in Cluster 2: *Gardnerella* |
| Cluster 3: *L. iners* vs Cluster 2: *Gardnerella* | *Gardnerella greenwoodii* | -4.7089225 | 4.91E-09 | Enriched in Cluster 2: *Gardnerella* |
| Cluster 3: *L. iners* vs Cluster 2: *Gardnerella* | *Lactobacillus iners* | 4.22119786 | 1.24E-08 | Enriched in Cluster 3: *L. iners* |
| Cluster 3: *L. iners* vs Cluster 2: *Gardnerella* | *Gardnerella swidsinskii* | -4.7457055 | 2.07E-08 | Enriched in Cluster 2: *Gardnerella* |
| Cluster 3: *L. iners* vs Cluster 2: *Gardnerella* | *Gardnerella leopoldii* | -4.4907094 | 2.40E-08 | Enriched in Cluster 2: *Gardnerella* |

| | | | | |
|---|---|---|---|---|
| Cluster 3: *L. iners* vs Cluster 2: *Gardnerella* | *Amygdalobacter indicium* | -3.1638725 | 3.37E-07 | Enriched in Cluster 2: *Gardnerella* |
| Cluster 3: *L. iners* vs Cluster 2: *Gardnerella* | *Gardnerella pickettii* | -3.8213016 | 1.02E-06 | Enriched in Cluster 2: *Gardnerella* |
| Cluster 3: *L. iners* vs Cluster 2: *Gardnerella* | *Lactobacillus jensenii* | 3.81043652 | 1.19E-06 | Enriched in Cluster 3: *L. iners* |
| Cluster 3: *L. iners* vs Cluster 2: *Gardnerella* | *Gardnerella vaginalis* | -3.7695413 | 3.14E-06 | Enriched in Cluster 2: *Gardnerella* |
| Cluster 3: *L. iners* vs Cluster 2: *Gardnerella* | *Gardnerella piotii* | -3.3971244 | 8.29E-06 | Enriched in Cluster 2: *Gardnerella* |
| Cluster 3: *L. iners* vs Cluster 2: *Gardnerella* | *Hoylesella buccalis* | -2.5385515 | 1.69E-05 | Enriched in Cluster 2: *Gardnerella* |
| Cluster 3: *L. iners* vs Cluster 2: *Gardnerella* | *Mobiluncus curtisii* | -2.0815696 | 0.00050572 | Enriched in Cluster 2: *Gardnerella* |

**Supplemental Table 3. Differentially abundant KEGG Orthology (KO) identifiers among microbial community clusters.**

| Feature | baseMean | log2FoldChange | pvalue | padj | EnrichedIn | Description |
|---|---|---|---|---|---|---|
| **K19349** | 4.949301965 | 6.018215457 | 1.47E-50 | 3.89E-47 | Cluster 1: L. crispatus | ATPases associated with a variety of cellular activities |
| **K09960** | 4.059889406 | 5.995016465 | 3.89E-31 | 1.47E-28 | Cluster 1: L. crispatus | Uncharacterized protein conserved in bacteria (DUF2213) |
| **K09961** | 3.589691739 | 5.723208238 | 5.82E-25 | 1.18E-22 | Cluster 1: L. crispatus | Protein of unknown function (DUF1073) |
| **K17244** | 2.909310907 | 5.56547305 | 8.44E-34 | 5.58E-31 | Cluster 1: L. crispatus | Bacterial extracellular solute-binding protein |
| **K01535** | 3.147179006 | 5.464290186 | 5.31E-18 | 3.26E-16 | Cluster 1: L. crispatus | Cation transporter/ATPase, N-terminus |
| **K09835** | 2.56883627 | 5.395260591 | 1.22E-28 | 3.23E-26 | Cluster 1: L. crispatus | HI0933-like protein |
| **K00986** | 14.35821431 | 4.713189107 | 3.05E-21 | 3.84E-19 | Cluster 1: L. crispatus | PFAM RNA-directed DNA polymerase (Reverse transcriptase) |
| **K02488** | 1.659990487 | 4.706557076 | 2.65E-15 | 8.32E-14 | Cluster 1: L. crispatus | GGDEF domain |
| **K16209** | 3.006154113 | 4.632602598 | 2.07E-23 | 3.41E-21 | Cluster 1: L. crispatus | Transporter |
| **K12292** | 9.421492859 | 4.58600494 | 1.91E-17 | 9.87E-16 | Cluster 1: L. crispatus | ABC-type bacteriocin lantibiotic exporters, contain an N-terminal double-glycine peptidase domain |
| **K14193** | 1.428448951 | 4.482967032 | 2.83E-21 | 3.73E-19 | Cluster 1: L. crispatus | Iron Transport-associated domain |

| | | | | | | |
|---|---|---|---|---|---|---|
| **K07313** | 2.966495704 | 4.452381405 | 5.71E-18 | 3.43E-16 | Cluster 1: L. crispatus | Calcineurin-like phosphoesterase |
| **K12941** | 2.651201829 | 4.367428519 | 8.71E-27 | 1.92E-24 | Cluster 1: L. crispatus | Peptidase dimerisation domain |
| **K07496** | 42.54868662 | 4.263390882 | 1.50E-28 | 3.59E-26 | Cluster 1: L. crispatus | Putative transposase DNA-binding domain |
| **K01577** | 4.023371302 | 4.259039229 | 4.00E-16 | 1.49E-14 | Cluster 1: L. crispatus | Belongs to the TPP enzyme family |
| **K07493** | 232.2822929 | 4.206829889 | 1.23E-22 | 1.80E-20 | Cluster 1: L. crispatus | Transposase |
| **K11621** | 1.871789003 | 4.105735616 | 1.64E-19 | 1.40E-17 | Cluster 1: L. crispatus | Putative adhesin |
| **K12293** | 1.483251829 | 4.070212103 | 1.64E-14 | 4.20E-13 | Cluster 1: L. crispatus | Transport protein ComB |
| **K06977** | 2.402824459 | 4.000955141 | 5.29E-15 | 1.49E-13 | Cluster 1: L. crispatus | acetyltransferase |
| **K00009** | 3.38428704 | 3.991529087 | 1.99E-21 | 2.76E-19 | Cluster 1: L. crispatus | mannitol-1-phosphate 5-dehydrogenase activity |
| **K07749** | 3.537585715 | 3.983363154 | 3.22E-20 | 2.84E-18 | Cluster 1: L. crispatus | Involved in the catabolism of oxalate and in the adapatation to low pH via the induction of the oxalate-dependent acid tolerance response (ATR). Catalyzes the transfer of the CoA moiety from formyl-CoA to oxalate |
| **K02054** | 1.585721995 | 3.915220977 | 7.49E-16 | 2.57E-14 | Cluster 1: L. crispatus | Binding-protein-dependent transport system inner membrane component |

| | | | | | | |
|---|---|---|---|---|---|---|
| **K13021** | 0.791480538 | 3.863345351 | 8.25E-16 | 2.79E-14 | Cluster 1: L. crispatus | Major Facilitator Superfamily |
| **K15524** | 3.569417287 | 3.847647304 | 9.35E-07 | 3.68E-06 | Cluster 1: L. crispatus | Glycosyl hydrolases family 38 N-terminal domain |
| **K02799** | 4.582621017 | 3.819578123 | 8.54E-17 | 3.76E-15 | Cluster 1: L. crispatus | PTS system, Lactose/Cellobiose specific IIB subunit |
| **K02800** | 4.582621017 | 3.819578123 | 8.54E-17 | 3.76E-15 | Cluster 1: L. crispatus | PTS system, Lactose/Cellobiose specific IIB subunit |
| **K05823** | 2.692939383 | 3.803469468 | 2.39E-18 | 1.62E-16 | Cluster 1: L. crispatus | Catalyzes the conversion of N-acetyl-diaminopimelate to diaminopimelate and acetate |
| **K19285** | 1.989032803 | 3.796945302 | 3.14E-20 | 2.84E-18 | Cluster 1: L. crispatus | nitroreductase |
| **K01607** | 1.713305232 | 3.719023865 | 1.30E-18 | 9.29E-17 | Cluster 1: L. crispatus | Carboxymuconolactone decarboxylase family |
| **K12308** | 4.850997726 | 3.704699149 | 3.44E-29 | 1.14E-26 | Cluster 1: L. crispatus | #NAME? |
| **K00547** | 2.359747532 | 3.700069664 | 1.07E-18 | 7.82E-17 | Cluster 1: L. crispatus | homocysteine S-methyltransferase |
| **K13731** | 2.479161489 | 3.661465115 | 3.68E-14 | 8.38E-13 | Cluster 1: L. crispatus | N-acetylmuramoyl-L-alanine amidase |
| **K02205** | 1.481319474 | 3.589816662 | 1.79E-08 | 9.21E-08 | Cluster 1: L. crispatus | amino acid |
| **K16236** | 1.481319474 | 3.589816662 | 1.79E-08 | 9.21E-08 | Cluster 1: L. crispatus | amino acid |

| | | | | | | |
|---|---|---|---|---|---|---|
| **K05910** | 4.500469924 | 3.506404517 | 3.70E-16 | 1.42E-14 | Cluster 1: L. crispatus | NADH oxidase |
| **K00661** | 3.985435083 | 3.497738693 | 3.42E-37 | 3.01E-34 | Cluster 1: L. crispatus | Maltose acetyltransferase |
| **K16235** | 1.870754791 | 3.485878159 | 1.62E-13 | 3.02E-12 | Cluster 1: L. crispatus | amino acid |
| **K10716** | 4.895914827 | 3.481906208 | 6.09E-24 | 1.07E-21 | Cluster 1: L. crispatus | Ion channel |
| **K05340** | 2.03928709 | 3.478995187 | 3.95E-15 | 1.20E-13 | Cluster 1: L. crispatus | sugar transport |
| **K20345** | 1.835856029 | 3.463270234 | 3.85E-14 | 8.68E-13 | Cluster 1: L. crispatus | Transport protein ComB |
| **K07474** | 1.235569631 | 3.387733322 | 3.92E-13 | 6.36E-12 | Cluster 1: L. crispatus | Terminase small subunit |
| **K00156** | 4.67077866 | 3.375169976 | 1.62E-20 | 1.64E-18 | Cluster 1: L. crispatus | Belongs to the TPP enzyme family |
| **K21929** | 1.051626185 | 3.367037316 | 8.19E-14 | 1.68E-12 | Cluster 1: L. crispatus | Uracil-DNA glycosylase |
| **K03817** | 1.464725714 | 3.355809187 | 1.46E-14 | 3.82E-13 | Cluster 1: L. crispatus | Acetyltransferase (GNAT) domain |
| **K20344** | 10.21021967 | 3.329500045 | 1.23E-16 | 5.32E-15 | Cluster 1: L. crispatus | ABC-type bacteriocin lantibiotic exporters, contain an N-terminal double-glycine peptidase domain |
| **K01610** | 5.066834012 | 3.325406398 | 9.23E-23 | 1.43E-20 | Cluster 1: L. crispatus | Phosphoenolpyruvate carboxykinase |

| | | | | | | |
|---|---|---|---|---|---|---|
| **K11104** | 3.045155269 | 3.324098741 | 1.04E-09 | 7.11E-09 | Cluster 1: L. crispatus | Transporter |
| **K13531** | 0.780081398 | 3.322102239 | 4.71E-15 | 1.38E-13 | Cluster 1: L. crispatus | 6-O-methylguanine DNA methyltransferase, DNA binding domain |
| **K03322** | 4.088225147 | 3.303875717 | 3.82E-31 | 1.47E-28 | Cluster 1: L. crispatus | H( )-stimulated, divalent metal cation uptake system |
| **K18445** | 0.62778138 | 3.271166591 | 4.72E-11 | 4.60E-10 | Cluster 1: L. crispatus | Nudix hydrolase |
| **K03724** | 7.753991799 | 3.256704951 | 4.62E-21 | 5.18E-19 | Cluster 1: L. crispatus | DEAD DEAH box helicase |
| **K07707** | 2.335579392 | 3.234509892 | 9.78E-13 | 1.43E-11 | Cluster 1: L. crispatus | LytTr DNA-binding domain |
| **K08093** | 1.217277535 | 3.233383363 | 3.37E-10 | 2.55E-09 | Cluster 1: L. crispatus | Orotidine 5'-phosphate decarboxylase / HUMPS family |
| **K02055** | 1.946520415 | 3.233159855 | 6.35E-12 | 7.62E-11 | Cluster 1: L. crispatus | ABC transporter |
| **K21562** | 2.141196733 | 3.184655191 | 4.55E-14 | 1.00E-12 | Cluster 1: L. crispatus | helix_turn_helix, cAMP Regulatory protein |
| **K03078** | 1.175149805 | 3.171165606 | 7.62E-08 | 3.49E-07 | Cluster 1: L. crispatus | Orotidine 5'-phosphate decarboxylase / HUMPS family |
| **K02069** | 2.177287897 | 3.105980353 | 6.44E-18 | 3.70E-16 | Cluster 1: L. crispatus | Uncharacterised protein family (UPF0014) |
| **K07171** | 4.038656551 | 3.09724832 | 2.68E-19 | 2.14E-17 | Cluster 1: L. crispatus | PemK-like, MazF-like toxin of type II toxin-antitoxin system |

| | | | | | | |
|---|---|---|---|---|---|---|
| **K02053** | 0.91367392 | 3.026866019 | 8.69E-08 | 3.94E-07 | Cluster 1: L. crispatus | Binding-protein-dependent transport system inner membrane component |
| **K08987** | 1.085742651 | 3.00934975 | 3.02E-13 | 5.14E-12 | Cluster 1: L. crispatus | Protein of unknown function (DUF1304) |
| **K06386** | 1.219879255 | 3.008620233 | 1.98E-08 | 1.01E-07 | Cluster 1: L. crispatus | Peptidase family M23 |
| **K10778** | 0.819755644 | 2.940802157 | 2.04E-09 | 1.34E-08 | Cluster 1: L. crispatus | 6-O-methylguanine DNA methyltransferase, DNA binding domain |
| **K18908** | 5.760671039 | 2.936405496 | 2.35E-08 | 1.19E-07 | Cluster 1: L. crispatus | MATE efflux family protein |
| **K01447** | 4.066734996 | 2.898893258 | 2.86E-14 | 6.81E-13 | Cluster 1: L. crispatus | N-acetylmuramoyl-L-alanine amidase |
| **K03827** | 0.759518396 | 2.845065332 | 2.49E-06 | 9.30E-06 | Cluster 1: L. crispatus | Acetyltransferase (GNAT) domain |
| **K13993** | 0.953262679 | 2.832193695 | 1.82E-13 | 3.36E-12 | Cluster 1: L. crispatus | Belongs to the small heat shock protein (HSP20) family |
| **K09963** | 9.897171097 | 2.826116496 | 4.22E-17 | 2.14E-15 | Cluster 1: L. crispatus | Bacterial protein of unknown function (DUF871) |
| **K10254** | 11.48779295 | 2.821200665 | 4.61E-19 | 3.58E-17 | Cluster 1: L. crispatus | Myosin-crossreactive antigen |
| **K18347** | 5.799376791 | 2.796330831 | 5.12E-09 | 2.97E-08 | Cluster 1: L. crispatus | Belongs to the D-isomer specific 2-hydroxyacid dehydrogenase family |
| **K05966** | 1.784021455 | 2.790415102 | 1.21E-12 | 1.73E-11 | Cluster 1: L. crispatus | 2-(5''-triphosphoribosyl)-3'-dephosphocoenzyme-A synthase |

| | | | | | | |
|---|---|---|---|---|---|---|
| **K00158** | 12.86259154 | 2.7668981 | 6.42E-18 | 3.70E-16 | Cluster 1: L. crispatus | Belongs to the TPP enzyme family |
| **K13714** | 2.599861726 | 2.755437283 | 5.12E-10 | 3.71E-09 | Cluster 1: L. crispatus | N-acetylmuramoyl-L-alanine amidase |
| **K19424** | 0.591769314 | 2.739589243 | 0.015910158 | 0.040209308 | Cluster 1: L. crispatus | Glycosyl transferases group 1 |
| **K00574** | 2.965115522 | 2.739566927 | 1.49E-16 | 6.23E-15 | Cluster 1: L. crispatus | cyclopropane-fatty-acyl-phospholipid synthase |
| **K09861** | 4.075385087 | 2.738807354 | 3.20E-21 | 3.85E-19 | Cluster 1: L. crispatus | Belongs to the UPF0246 family |
| **K07706** | 3.185488344 | 2.737664496 | 1.33E-08 | 7.06E-08 | Cluster 1: L. crispatus | GHKL domain |
| **K21053** | 3.220702992 | 2.735238182 | 1.14E-09 | 7.77E-09 | Cluster 1: L. crispatus | Catalyzes the hydrolytic deamination of adenine to hypoxanthine. Plays an important role in the purine salvage pathway and in nitrogen catabolism |
| **K13256** | 0.53827213 | 2.716573387 | 1.69E-07 | 7.30E-07 | Cluster 1: L. crispatus | Phosphate-starvation-inducible E |
| **K18702** | 0.580824245 | 2.711819175 | 7.18E-06 | 2.55E-05 | Cluster 1: L. crispatus | Involved in the catabolism of oxalate and in the adapatation to low pH via the induction of the oxalate-dependent acid tolerance response (ATR). Catalyzes the transfer of the CoA moiety from formyl-CoA to oxalate |
| **K03811** | 2.07705407 | 2.703965515 | 1.94E-11 | 2.08E-10 | Cluster 1: L. crispatus | nicotinamide mononucleotide transporter |
| **K01486** | 3.175097524 | 2.675078436 | 5.18E-09 | 2.99E-08 | Cluster 1: L. crispatus | Adenine deaminase C-terminal domain |

| | | | | | | |
|---|---|---|---|---|---|---|
| **K01060** | 0.845522174 | 2.657593894 | 0.000676891 | 0.002026836 | Cluster 1: L. crispatus | Acetyl xylan esterase (AXE1) |
| **K03486** | 0.898292447 | 2.643012627 | 1.28E-09 | 8.66E-09 | Cluster 1: L. crispatus | UTRA |
| **K00015** | 1.911977418 | 2.640089751 | 4.66E-11 | 4.56E-10 | Cluster 1: L. crispatus | Belongs to the D-isomer specific 2-hydroxyacid dehydrogenase family |
| **K08968** | 0.787968283 | 2.61051181 | 9.89E-13 | 1.43E-11 | Cluster 1: L. crispatus | GAF domain-containing protein |
| **K11065** | 1.05260444 | 2.574587237 | 1.66E-12 | 2.32E-11 | Cluster 1: L. crispatus | Thiol-specific peroxidase that catalyzes the reduction of hydrogen peroxide and organic hydroperoxides to water and alcohols, respectively. Plays a role in cell protection against oxidative stress by detoxifying peroxides |
| **K01081** | 10.02131971 | 2.550005237 | 1.45E-13 | 2.74E-12 | Cluster 1: L. crispatus | Belongs to the 5'-nucleotidase family |
| **K02052** | 1.947400319 | 2.539969337 | 1.18E-10 | 1.03E-09 | Cluster 1: L. crispatus | Part of the ABC transporter complex PotABCD involved in spermidine putrescine import. Responsible for energy coupling to the transport system |
| **K16924** | 1.171150508 | 2.533159259 | 2.27E-10 | 1.84E-09 | Cluster 1: L. crispatus | UPF0397 protein |
| **K12999** | 3.870328906 | 2.520699754 | 6.35E-12 | 7.62E-11 | Cluster 1: L. crispatus | Glycosyltransferase, group 2 family protein |
| **K05822** | 0.388286953 | 2.504983346 | 1.15E-06 | 4.47E-06 | Cluster 1: L. crispatus | Catalyzes the transfer of an acetyl group from acetyl-CoA to tetrahydrodipicolinate |
| **K19055** | 0.665175212 | 2.502176136 | 8.03E-08 | 3.66E-07 | Cluster 1: L. crispatus | Aminoacyl-tRNA editing domain |

| | | | | | | |
|---|---|---|---|---|---|---|
| **K03973** | 0.668626588 | 2.469276893 | 3.50E-07 | 1.45E-06 | Cluster 1: L. crispatus | PspC domain |
| **K01442** | 2.889082919 | 2.449887071 | 4.50E-11 | 4.42E-10 | Cluster 1: L. crispatus | Linear amide C-N hydrolase, choloylglycine hydrolase family protein |
| **K01246** | 1.612828318 | 2.446184531 | 2.16E-14 | 5.37E-13 | Cluster 1: L. crispatus | glycosylase |
| **K02236** | 2.414456809 | 2.439467598 | 5.99E-10 | 4.30E-09 | Cluster 1: L. crispatus | Bacterial Peptidase A24 N-terminal domain |
| **K18892** | 5.367541478 | 2.417411173 | 7.35E-08 | 3.37E-07 | Cluster 1: L. crispatus | ABC transporter transmembrane region |
| **K03077** | 4.127028466 | 2.415593347 | 1.60E-13 | 2.99E-12 | Cluster 1: L. crispatus | Links the arabinose metabolic pathway to the pentose phosphate pathway and allows the bacteria to use arabinose as an energy source |
| **K03206** | 1.319634197 | 2.402824194 | 9.19E-08 | 4.14E-07 | Cluster 1: L. crispatus | NADPH-dependent FMN reductase |
| **K00851** | 5.109181731 | 2.392117199 | 2.46E-16 | 9.85E-15 | Cluster 1: L. crispatus | Belongs to the FGGY kinase family |
| **K22441** | 1.44254217 | 2.387733997 | 1.22E-08 | 6.53E-08 | Cluster 1: L. crispatus | Acetyltransferase (GNAT) domain |
| **K01778** | 1.90749344 | 2.384605902 | 2.99E-11 | 3.11E-10 | Cluster 1: L. crispatus | Catalyzes the stereoinversion of LL-2,6-diaminoheptanedioate (L,L-DAP) to meso-diaminoheptanedioate (meso- DAP), a precursor of L-lysine and an essential component of the bacterial peptidoglycan |
| **K03484** | 7.183403125 | 2.380391326 | 4.95E-11 | 4.75E-10 | Cluster 1: L. crispatus | Periplasmic binding protein domain |

| | | | | | | |
|---|---|---|---|---|---|---|
| **K03885** | 5.026228477 | 2.377514601 | 1.26E-12 | 1.77E-11 | Cluster 1: L. crispatus | NADH dehydrogenase |
| **K00390** | 0.759918054 | 2.368387515 | 0.003251677 | 0.008982927 | Cluster 1: L. crispatus | Sulfate reduction |
| **K01643** | 3.976584429 | 2.352673666 | 4.97E-11 | 4.76E-10 | Cluster 1: L. crispatus | Citrate (pro-3S)-lyase alpha chain |
| **K06872** | 2.44812728 | 2.32777221 | 1.86E-13 | 3.40E-12 | Cluster 1: L. crispatus | TPM domain |
| **K04086** | 8.66198916 | 2.317821736 | 1.74E-10 | 1.45E-09 | Cluster 1: L. crispatus | AAA domain (Cdc48 subfamily) |
| **K02041** | 2.17170732 | 2.313155455 | 2.46E-09 | 1.57E-08 | Cluster 1: L. crispatus | Part of the ABC transporter complex PhnCDE involved in phosphonates import. Responsible for energy coupling to the transport system |
| **K00674** | 0.97509908 | 2.302408264 | 9.59E-09 | 5.27E-08 | Cluster 1: L. crispatus | Catalyzes the transfer of an acetyl group from acetyl-CoA to tetrahydrodipicolinate |
| **K03652** | 2.637600665 | 2.298411167 | 4.87E-14 | 1.06E-12 | Cluster 1: L. crispatus | Belongs to the DNA glycosylase MPG family |
| **K01625** | 0.859871412 | 2.297066555 | 7.01E-09 | 3.98E-08 | Cluster 1: L. crispatus | KDPG and KHG aldolase |
| **K00712** | 0.503351838 | 2.244679086 | 0.013374406 | 0.034175834 | Cluster 1: L. crispatus | Glycosyl transferases group 1 |
| **K02044** | 8.621768477 | 2.231877891 | 9.79E-10 | 6.77E-09 | Cluster 1: L. crispatus | Phosphonate ABC transporter |
| **K01662** | 7.439534621 | 2.21725034 | 5.02E-17 | 2.45E-15 | Cluster 1: L. crispatus | Catalyzes the acyloin condensation reaction between C atoms 2 and 3 of pyruvate and glyceraldehyde 3- |

| | | | | | | |
|---|---|---|---|---|---|---|
| | | | | | | phosphate to yield 1-deoxy-D-xylulose-5-phosphate (DXP) |
| **K02042** | 5.924349204 | 2.216031891 | 4.88E-11 | 4.72E-10 | Cluster 1: L. crispatus | Binding-protein-dependent transport system inner membrane component |
| **K03455** | 7.528991242 | 2.205625253 | 7.99E-08 | 3.64E-07 | Cluster 1: L. crispatus | TrkA C-terminal domain protein |
| **K09704** | 1.27070096 | 2.202188511 | 0.000582561 | 0.001750333 | Cluster 1: L. crispatus | Metal-independent alpha-mannosidase (GH125) |
| **K18891** | 4.691864843 | 2.193183192 | 2.09E-06 | 7.91E-06 | Cluster 1: L. crispatus | ABC transporter, ATP-binding protein |
| **K18829** | 0.163124357 | 2.189843101 | 0.005882053 | 0.015787095 | Cluster 1: L. crispatus | SpoVT / AbrB like domain |
| **K01581** | 7.287630611 | 2.187616151 | 1.99E-07 | 8.52E-07 | Cluster 1: L. crispatus | Orn/Lys/Arg decarboxylase, C-terminal domain |
| **K03307** | 2.411534468 | 2.157276822 | 2.91E-06 | 1.08E-05 | Cluster 1: L. crispatus | Belongs to the sodium solute symporter (SSF) (TC 2.A.21) family |
| **K08693** | 4.955123024 | 2.134841144 | 5.40E-06 | 1.96E-05 | Cluster 1: L. crispatus | Belongs to the 5'-nucleotidase family |
| **K07106** | 8.248002627 | 2.12908343 | 4.73E-13 | 7.49E-12 | Cluster 1: L. crispatus | Bacterial protein of unknown function (DUF871) |
| **K00135** | 7.478108001 | 2.128232561 | 2.84E-13 | 4.98E-12 | Cluster 1: L. crispatus | Belongs to the aldehyde dehydrogenase family |
| **K07004** | 5.31079467 | 2.127920031 | 3.38E-09 | 2.08E-08 | Cluster 1: L. crispatus | Belongs to the 5'-nucleotidase family |

| | | | | | | |
|---|---|---|---|---|---|---|
| **K03790** | 1.510294137 | 2.12248056 | 7.84E-11 | 7.14E-10 | Cluster 1: L. crispatus | Acetyltransferase (GNAT) domain |
| **K12296** | 1.249202541 | 2.112489407 | 2.55E-07 | 1.08E-06 | Cluster 1: L. crispatus | Belongs to the sigma-70 factor family |
| **K06909** | 1.490824809 | 2.10367556 | 7.33E-05 | 0.000236408 | Cluster 1: L. crispatus | Terminase RNAseH like domain |
| **K03488** | 2.957077792 | 2.102393467 | 1.72E-09 | 1.14E-08 | Cluster 1: L. crispatus | CAT RNA binding domain |
| **K20112** | 1.333218021 | 2.099944209 | 3.19E-07 | 1.33E-06 | Cluster 1: L. crispatus | Phosphoenolpyruvate-dependent sugar phosphotransferase system, EIIA 2 |
| **K11751** | 5.167683538 | 2.076943154 | 4.44E-08 | 2.13E-07 | Cluster 1: L. crispatus | Belongs to the 5'-nucleotidase family |
| **K06994** | 12.8591277 | 2.075718735 | 1.70E-10 | 1.42E-09 | Cluster 1: L. crispatus | MMPL family |
| **K01226** | 5.107764314 | 2.063307879 | 2.61E-08 | 1.30E-07 | Cluster 1: L. crispatus | Alpha amylase, catalytic domain protein |
| **K16012** | 5.770413809 | 2.063155845 | 1.01E-11 | 1.16E-10 | Cluster 1: L. crispatus | ABC transporter, CydDC cysteine exporter (CydDC-E) family, permease ATP-binding protein CydC |
| **K14155** | 4.861331914 | 2.055254422 | 5.40E-14 | 1.16E-12 | Cluster 1: L. crispatus | Aminotransferase, class I |
| **K00794** | 1.062885227 | 2.054131666 | 4.23E-09 | 2.51E-08 | Cluster 1: L. crispatus | Catalyzes the formation of 6,7-dimethyl-8-ribityllumazine by condensation of 5-amino-6-(D-ribitylamino)uracil with 3,4-dihydroxy-2-butanone 4-phosphate. This is the penultimate step in the biosynthesis of riboflavin |

| | | | | | | |
|---|---|---|---|---|---|---|
| **K01582** | 7.57377644 | 2.033158955 | 2.48E-08 | 1.25E-07 | Cluster 1: L. crispatus | Orn/Lys/Arg decarboxylase, C-terminal domain |
| **K03476** | 3.95517047 | 2.032720706 | 3.90E-09 | 2.36E-08 | Cluster 1: L. crispatus | Beta-lactamase superfamily domain |
| **K03284** | 6.873213078 | 2.015827672 | 2.53E-19 | 2.09E-17 | Cluster 1: L. crispatus | CorA-like Mg2+ transporter protein |
| **K02371** | 0.927946405 | 2.014584017 | 6.40E-05 | 0.000207751 | Cluster 1: L. crispatus | Nitronate monooxygenase |
| **K07305** | 0.942434942 | 2.011440467 | 7.35E-10 | 5.19E-09 | Cluster 1: L. crispatus | peptide methionine sulfoxide reductase |
| **K01119** | 10.24029709 | 1.984025302 | 2.09E-08 | 1.07E-07 | Cluster 1: L. crispatus | Belongs to the 5'-nucleotidase family |
| **K00865** | 5.27985503 | 1.974945024 | 3.79E-13 | 6.22E-12 | Cluster 1: L. crispatus | Belongs to the glycerate kinase type-1 family |
| **K02809** | 12.66332499 | 1.972564006 | 7.25E-07 | 2.89E-06 | Cluster 1: L. crispatus | Phosphotransferase system |
| **K02810** | 12.66332499 | 1.972564006 | 7.25E-07 | 2.89E-06 | Cluster 1: L. crispatus | Phosphotransferase system |
| **K01644** | 1.78127509 | 1.971116438 | 3.35E-08 | 1.64E-07 | Cluster 1: L. crispatus | Belongs to the HpcH HpaI aldolase family |
| **K00425** | 5.081561248 | 1.941733533 | 2.92E-09 | 1.83E-08 | Cluster 1: L. crispatus | Ubiquinol oxidase |
| **K03492** | 1.157311458 | 1.936392996 | 0.001987908 | 0.005627078 | Cluster 1: L. crispatus | UTRA domain |

| | | | | | | |
|---|---|---|---|---|---|---|
| **K02315** | 4.112057753 | 1.927928904 | 1.60E-07 | 6.97E-07 | Cluster 1: L. crispatus | IstB-like ATP binding protein |
| **K07776** | 0.355473885 | 1.918070278 | 0.005736482 | 0.015427748 | Cluster 1: L. crispatus | Transcriptional regulatory protein, C terminal |
| **K02756** | 12.73503713 | 1.912556458 | 5.11E-08 | 2.41E-07 | Cluster 1: L. crispatus | Phosphotransferase system |
| **K02757** | 12.73503713 | 1.912556458 | 5.11E-08 | 2.41E-07 | Cluster 1: L. crispatus | Phosphotransferase system |
| **K18893** | 0.472002113 | 1.905557636 | 0.00406112 | 0.011075751 | Cluster 1: L. crispatus | ABC-type multidrug transport system, ATPase and permease components |
| **K00805** | 3.971756245 | 1.891080868 | 9.08E-14 | 1.84E-12 | Cluster 1: L. crispatus | Belongs to the FPP GGPP synthase family |
| **K00426** | 2.69092798 | 1.890901986 | 3.00E-10 | 2.32E-09 | Cluster 1: L. crispatus | Cytochrome d ubiquinol oxidase subunit II |
| **K03183** | 1.576436995 | 1.888582576 | 2.30E-10 | 1.85E-09 | Cluster 1: L. crispatus | Methyltransferase required for the conversion of demethylmenaquinol (DMKH2) to menaquinol (MKH2) |
| **K01512** | 0.279841186 | 1.878965317 | 6.91E-05 | 0.000223145 | Cluster 1: L. crispatus | Belongs to the acylphosphatase family |
| **K02068** | 1.30959506 | 1.874232106 | 3.39E-07 | 1.40E-06 | Cluster 1: L. crispatus | ABC transporter, ATP-binding protein |
| **K03693** | 0.821446194 | 1.872274974 | 0.010627239 | 0.027516215 | Cluster 1: L. crispatus | Penicillin binding protein transpeptidase domain |
| **K12551** | 0.821446194 | 1.872274974 | 0.010627239 | 0.027516215 | Cluster 1: L. crispatus | Penicillin binding protein transpeptidase domain |

| | | | | | | |
|---|---|---|---|---|---|---|
| **K07729** | 2.816577703 | 1.859623476 | 1.38E-11 | 1.54E-10 | Cluster 1: L. crispatus | Helix-turn-helix XRE-family like proteins |
| **K01910** | 2.799645186 | 1.858866582 | 2.37E-07 | 1.01E-06 | Cluster 1: L. crispatus | Acetylation of prosthetic group (2-(5"-phosphoribosyl)- 3'-dephosphocoenzyme-A) of the gamma subunit of citrate lyase |
| **K19159** | 0.68913933 | 1.857093404 | 0.001570359 | 0.004498175 | Cluster 1: L. crispatus | Antitoxin component of a toxin-antitoxin (TA) module |
| **K06221** | 4.157983272 | 1.850728774 | 4.39E-08 | 2.11E-07 | Cluster 1: L. crispatus | Reductase |
| **K00467** | 4.331118791 | 1.844761731 | 4.62E-05 | 0.000151821 | Cluster 1: L. crispatus | L-lactate dehydrogenase (FMN-dependent) and related alpha-hydroxy acid dehydrogenases |
| **K10530** | 4.355773408 | 1.836812951 | 4.98E-05 | 0.000163731 | Cluster 1: L. crispatus | L-lactate dehydrogenase (FMN-dependent) and related alpha-hydroxy acid dehydrogenases |
| **K06921** | 10.82751237 | 1.828503623 | 7.26E-07 | 2.89E-06 | Cluster 1: L. crispatus | Archaea bacterial proteins of unknown function |
| **K02808** | 9.113149425 | 1.826774309 | 9.75E-06 | 3.44E-05 | Cluster 1: L. crispatus | Phosphotransferase system |
| **K00600** | 6.01294503 | 1.8250225 | 2.85E-13 | 4.98E-12 | Cluster 1: L. crispatus | Catalyzes the reversible interconversion of serine and glycine with tetrahydrofolate (THF) serving as the one-carbon carrier. This reaction serves as the major source of one-carbon groups required for the biosynthesis of purines, thymidylate, methionine, and other important biomolecules. Also exhibits THF-independent aldolase activity toward beta-hydroxyamino acids, producing glycine and aldehydes, via a retro-aldol mechanism |

| | | | | | | |
|---|---|---|---|---|---|---|
| **K00793** | 1.520933813 | 1.798504101 | 2.11E-09 | 1.38E-08 | Cluster 1: L. crispatus | Riboflavin synthase |
| **K01585** | 7.891741648 | 1.795091803 | 5.63E-10 | 4.06E-09 | Cluster 1: L. crispatus | Orn/Lys/Arg decarboxylase, C-terminal domain |
| **K15580** | 38.82763252 | 1.790576769 | 5.27E-09 | 3.04E-08 | Cluster 1: L. crispatus | ABC transporter, substratebinding protein |
| **K15583** | 5.910680252 | 1.784176283 | 3.46E-09 | 2.12E-08 | Cluster 1: L. crispatus | Belongs to the ABC transporter superfamily |
| **K07341** | 3.947239115 | 1.772620731 | 7.16E-08 | 3.31E-07 | Cluster 1: L. crispatus | Fic/DOC family |
| **K11752** | 4.891041429 | 1.771672968 | 5.79E-12 | 7.05E-11 | Cluster 1: L. crispatus | Converts 2,5-diamino-6-(ribosylamino)-4(3h)-pyrimidinone 5'-phosphate into 5-amino-6-(ribosylamino)-2,4(1h,3h)- pyrimidinedione 5'-phosphate |
| **K07304** | 2.369641988 | 1.766006479 | 1.69E-09 | 1.12E-08 | Cluster 1: L. crispatus | Has an important function as a repair enzyme for proteins that have been inactivated by oxidation. Catalyzes the reversible oxidation-reduction of methionine sulfoxide in proteins to methionine |
| **K01424** | 5.487125108 | 1.75626511 | 1.29E-11 | 1.45E-10 | Cluster 1: L. crispatus | L-asparaginase, type I |
| **K02246** | 0.913770201 | 1.749418099 | 2.10E-06 | 7.93E-06 | Cluster 1: L. crispatus | Competence protein |
| **K12267** | 2.357769278 | 1.729335753 | 2.85E-09 | 1.79E-08 | Cluster 1: L. crispatus | Has an important function as a repair enzyme for proteins that have been inactivated by oxidation. Catalyzes the reversible oxidation-reduction of methionine sulfoxide in proteins to methionine |

| | | | | | | |
|---|---|---|---|---|---|---|
| **K09155** | 6.447397277 | 1.726095009 | 3.22E-13 | 5.40E-12 | Cluster 1: L. crispatus | PAS domain |
| **K03499** | 9.274805291 | 1.717498634 | 6.02E-13 | 9.20E-12 | Cluster 1: L. crispatus | TrkA C-terminal domain protein |
| **K10542** | 0.62108708 | 1.711612597 | 0.018862154 | 0.046862605 | Cluster 1: L. crispatus | Part of the ABC transporter complex RbsABC involved in ribose import. Responsible for energy coupling to the transport system |
| **K03483** | 9.934199275 | 1.683292098 | 4.50E-09 | 2.66E-08 | Cluster 1: L. crispatus | Mga helix-turn-helix domain |
| **K00567** | 1.543288218 | 1.674057466 | 9.35E-08 | 4.21E-07 | Cluster 1: L. crispatus | 6-O-methylguanine DNA methyltransferase, DNA binding domain |
| **K07636** | 8.027931833 | 1.671443574 | 1.55E-09 | 1.03E-08 | Cluster 1: L. crispatus | Histidine kinase |
| **K05964** | 0.517224606 | 1.656030521 | 0.000544975 | 0.001645045 | Cluster 1: L. crispatus | Apo-citrate lyase phosphoribosyl-dephospho-CoA transferase |
| **K15773** | 0.183460009 | 1.639694276 | 0.016877357 | 0.042369867 | Cluster 1: L. crispatus | Peptidyl-tyrosine sulfation |
| **K03491** | 1.765348981 | 1.633691809 | 0.000586956 | 0.001761535 | Cluster 1: L. crispatus | Mga helix-turn-helix domain |
| **K03654** | 5.034711243 | 1.632961332 | 4.29E-08 | 2.07E-07 | Cluster 1: L. crispatus | ATP-dependent DNA helicase RecQ |
| **K00215** | 1.709534698 | 1.623447948 | 3.43E-08 | 1.68E-07 | Cluster 1: L. crispatus | Catalyzes the conversion of 4-hydroxy-tetrahydrodipicolinate (HTPA) to tetrahydrodipicolinate |

| | | | | | | |
|---|---|---|---|---|---|---|
| **K14652** | 7.497106877 | 1.620116069 | 2.61E-10 | 2.05E-09 | Cluster 1: L. crispatus | Catalyzes the conversion of GTP to 2,5-diamino-6-ribosylamino-4(3H)-pyrimidinone 5'-phosphate (DARP), formate and pyrophosphate |
| **K01179** | 0.283211022 | 1.616480708 | 0.016937752 | 0.042481104 | Cluster 1: L. crispatus | regulation of response to stimulus |
| **K01488** | 4.102989357 | 1.610196826 | 3.35E-10 | 2.55E-09 | Cluster 1: L. crispatus | Catalyzes the hydrolytic deamination of adenine to hypoxanthine. Plays an important role in the purine salvage pathway and in nitrogen catabolism |
| **K14337** | 0.687916162 | 1.59188134 | 0.007693612 | 0.020379969 | Cluster 1: L. crispatus | Carotene biosynthesis associated membrane protein |
| **K14339** | 0.687916162 | 1.59188134 | 0.007693612 | 0.020379969 | Cluster 1: L. crispatus | Carotene biosynthesis associated membrane protein |
| **K03303** | 10.27403388 | 1.590071766 | 1.27E-07 | 5.65E-07 | Cluster 1: L. crispatus | L-lactate permease |
| **K03079** | 2.851688733 | 1.589904676 | 1.16E-05 | 4.06E-05 | Cluster 1: L. crispatus | Xylose isomerase-like TIM barrel |
| **K01768** | 0.51540531 | 1.588392547 | 0.008000041 | 0.021128109 | Cluster 1: L. crispatus | Adenylate cyclase, family 3 (some proteins contain HAMP domain) |
| **K02755** | 14.59859629 | 1.581652934 | 3.97E-07 | 1.63E-06 | Cluster 1: L. crispatus | phosphotransferase system |
| **K13571** | 0.868449724 | 1.576560193 | 0.003988414 | 0.010892866 | Cluster 1: L. crispatus | Catalyzes the covalent attachment of the prokaryotic ubiquitin-like protein modifier Pup to the proteasomal substrate proteins, thereby targeting them for proteasomal degradation. This tagging system is termed pupylation. The ligation reaction involves the side-chain carboxylate of the C-terminal glutamate of |

| | | | | | | |
|---|---|---|---|---|---|---|
| | | | | | | Pup and the side-chain amino group of a substrate lysine |
| **K01613** | 0.696464744 | 1.576315341 | 0.003653608 | 0.010030331 | Cluster 1: L. crispatus | Belongs to the phosphatidylserine decarboxylase family |
| **K10823** | 3.927871416 | 1.566987342 | 2.48E-09 | 1.58E-08 | Cluster 1: L. crispatus | Belongs to the ABC transporter superfamily |
| **K22445** | 0.636838255 | 1.561493832 | 0.012042314 | 0.031028049 | Cluster 1: L. crispatus | Belongs to the aldehyde dehydrogenase family |
| **K04083** | 3.007653915 | 1.558112068 | 2.97E-10 | 2.31E-09 | Cluster 1: L. crispatus | Redox regulated molecular chaperone. Protects both thermally unfolding and oxidatively damaged proteins from irreversible aggregation. Plays an important role in the bacterial defense system toward oxidative stress |
| **K01218** | 0.272700465 | 1.537999374 | 0.003323126 | 0.009170716 | Cluster 1: L. crispatus | regulation of response to stimulus |
| **K01821** | 0.266397364 | 1.536221898 | 0.007081009 | 0.018832774 | Cluster 1: L. crispatus | Belongs to the 4-oxalocrotonate tautomerase family |
| **K01181** | 0.383012941 | 1.535936402 | 0.001304719 | 0.003807473 | Cluster 1: L. crispatus | Polysaccharide deacetylase |
| **K01679** | 5.44977077 | 1.523151698 | 1.59E-10 | 1.34E-09 | Cluster 1: L. crispatus | Involved in the TCA cycle. Catalyzes the stereospecific interconversion of fumarate to L-malate |
| **K01190** | 7.874644914 | 1.519660967 | 1.66E-06 | 6.39E-06 | Cluster 1: L. crispatus | Belongs to the glycosyl hydrolase 2 family |
| **K02761** | 23.16068459 | 1.504694753 | 4.96E-08 | 2.35E-07 | Cluster 1: L. crispatus | The phosphoenolpyruvate-dependent sugar phosphotransferase system (PTS), a major carbohydrate active - transport system, catalyzes the phosphorylation of incoming sugar substrates |

| | | | | | | |
|---|---|---|---|---|---|---|
| | | | | | | concomitant with their translocation across the cell membrane |
| **K20814** | 0.667189459 | 1.486108099 | 0.015357775 | 0.038999888 | Cluster 1: L. crispatus | Pup-ligase protein |
| **K07054** | 0.569896351 | 1.483737733 | 0.009898779 | 0.025705679 | Cluster 1: L. crispatus | Putative neutral zinc metallopeptidase |
| **K21401** | 0.683070591 | 1.480992234 | 0.008669391 | 0.022736706 | Cluster 1: L. crispatus | Dehydrogenases (flavoproteins) |
| **K01193** | 7.682423785 | 1.463717144 | 7.61E-06 | 2.70E-05 | Cluster 1: L. crispatus | invertase |
| **K03095** | 1.013712505 | 1.42961729 | 0.004097602 | 0.01115646 | Cluster 1: L. crispatus | Belongs to the SprT family |
| **K00104** | 7.431974825 | 1.429147045 | 2.41E-06 | 9.04E-06 | Cluster 1: L. crispatus | FAD linked oxidases, C-terminal domain |
| **K01966** | 0.791815321 | 1.423667418 | 0.01716429 | 0.04292698 | Cluster 1: L. crispatus | Acetyl-CoA carboxylase, carboxyltransferase component (subunits alpha and beta) |
| **K16148** | 0.673573716 | 1.419173501 | 0.016092656 | 0.040592841 | Cluster 1: L. crispatus | Glycogen synthase, Corynebacterium family |
| **K03781** | 1.019398841 | 1.402485539 | 0.00804592 | 0.021228047 | Cluster 1: L. crispatus | Catalase |
| **K01961** | 1.80449741 | 1.398877042 | 0.0089537 | 0.0234358 | Cluster 1: L. crispatus | Acetyl-CoA carboxylase biotin carboxylase subunit |
| **K18231** | 7.533410589 | 1.390212301 | 4.45E-05 | 0.000146386 | Cluster 1: L. crispatus | ATPases associated with a variety of cellular activities |

| | | | | | | |
|---|---|---|---|---|---|---|
| **K00728** | 0.952518613 | 1.387851531 | 0.009278865 | 0.024190998 | Cluster 1: L. crispatus | Dolichyl-phosphate-mannose--protein |
| **K01586** | 5.112917216 | 1.374397097 | 8.86E-07 | 3.50E-06 | Cluster 1: L. crispatus | Specifically catalyzes the decarboxylation of meso-diaminopimelate (meso-DAP) to L-lysine |
| **K03778** | 7.632793646 | 1.374065302 | 4.59E-06 | 1.67E-05 | Cluster 1: L. crispatus | Belongs to the D-isomer specific 2-hydroxyacid dehydrogenase family |
| **K09384** | 5.242627088 | 1.367901073 | 2.97E-05 | 9.91E-05 | Cluster 1: L. crispatus | Uncharacterized conserved protein (DUF2075) |
| **K01007** | 8.87344875 | 1.311895123 | 3.68E-06 | 1.36E-05 | Cluster 1: L. crispatus | Catalyzes the phosphorylation of pyruvate to phosphoenolpyruvate |
| **K01227** | 6.025037381 | 1.302975068 | 1.22E-05 | 4.26E-05 | Cluster 1: L. crispatus | mannosyl-glycoprotein |
| **K07010** | 3.223304014 | 1.295175099 | 5.90E-07 | 2.38E-06 | Cluster 1: L. crispatus | peptidase C26 |
| **K00662** | 2.01568714 | 1.280677215 | 0.018470999 | 0.046020667 | Cluster 1: L. crispatus | Aminoglycoside 3-N-acetyltransferase |
| **K05833** | 4.789347759 | 1.249014724 | 6.42E-06 | 2.30E-05 | Cluster 1: L. crispatus | ABC transporter, ATP-binding protein |
| **K02395** | 0.74676309 | 1.238509881 | 0.003061044 | 0.008482914 | Cluster 1: L. crispatus | amidase activity |
| **K02013** | 4.561807666 | 1.237878308 | 4.11E-06 | 1.51E-05 | Cluster 1: L. crispatus | abc transporter atp-binding protein |
| **K20861** | 8.996092432 | 1.231971753 | 0.000481019 | 0.001461877 | Cluster 1: L. crispatus | Sucrose-6F-phosphate phosphohydrolase |

| | | | | | | |
|---|---|---|---|---|---|---|
| **K01739** | 1.038786811 | 1.228521753 | 0.016496533 | 0.041492707 | Cluster 1: L. crispatus | Cys/Met metabolism PLP-dependent enzyme |
| **K07150** | 3.231065255 | 1.220650809 | 3.43E-05 | 0.000113683 | Cluster 1: L. crispatus | Protein of unknown function (DUF554) |
| **K11737** | 14.02152435 | 1.195547652 | 0.000132349 | 0.000419105 | Cluster 1: L. crispatus | Amino acid permease |
| **K20480** | 2.255086238 | 1.182337973 | 0.003967808 | 0.010859048 | Cluster 1: L. crispatus | Helix-turn-helix |
| **K01919** | 1.219268727 | 1.164188762 | 0.013069998 | 0.03351249 | Cluster 1: L. crispatus | glutamate--cysteine ligase |
| **K00841** | 11.80665037 | 1.163387401 | 0.000535983 | 0.001623316 | Cluster 1: L. crispatus | Aminotransferase |
| **K00001** | 3.84546346 | 1.153022488 | 0.002291987 | 0.006412223 | Cluster 1: L. crispatus | alcohol dehydrogenase |
| **K01223** | 13.71380223 | 1.114350592 | 0.003603298 | 0.009902508 | Cluster 1: L. crispatus | Belongs to the glycosyl hydrolase 1 family |
| **K18930** | 1.002598855 | 1.102191246 | 0.018791977 | 0.046732214 | Cluster 1: L. crispatus | FAD linked oxidase |
| **K10353** | 5.29886851 | 1.100828351 | 0.004118888 | 0.011202867 | Cluster 1: L. crispatus | deoxynucleoside kinase |
| **K07467** | 4.046440646 | 1.083402698 | 0.008758496 | 0.022947607 | Cluster 1: L. crispatus | Replication initiation factor |
| **K09458** | 2.356142141 | 1.073296208 | 0.019870716 | 0.049183282 | Cluster 1: L. crispatus | Catalyzes the condensation reaction of fatty acid synthesis by the addition to an acyl acceptor of two carbons from malonyl-ACP |

| | | | | | | |
|---|---|---|---|---|---|---|
| **K07124** | 6.427821483 | 1.031211195 | 0.000187144 | 0.000584216 | Cluster 1: L. crispatus | Belongs to the short-chain dehydrogenases reductases (SDR) family |
| **K01478** | 4.876268184 | 1.026443393 | 9.01E-05 | 0.000288698 | Cluster 1: L. crispatus | Arginine |
| **K00849** | 4.540263043 | 1.008191855 | 0.000146415 | 0.000461436 | Cluster 1: L. crispatus | Catalyzes the transfer of the gamma-phosphate of ATP to D-galactose to form alpha-D-galactose-1-phosphate (Gal-1-P) |
| **K07675** | 13.70676583 | 5.279648013 | 7.71E-33 | 9.70E-31 | Cluster 2: Gardnerella | Histidine kinase |
| **K09818** | 16.45920972 | 5.01970051 | 1.34E-36 | 3.21E-34 | Cluster 2: Gardnerella | Zinc-uptake complex component A periplasmic |
| **K03291** | 3.620727871 | 4.934623872 | 5.40E-11 | 2.28E-10 | Cluster 2: Gardnerella | Major facilitator Superfamily |
| **K03457** | 7.576813954 | 4.89257598 | 3.04E-27 | 1.04E-25 | Cluster 2: Gardnerella | Permease for cytosine/purines, uracil, thiamine, allantoin |
| **K20263** | 4.32091753 | 4.846882772 | 8.67E-36 | 1.88E-33 | Cluster 2: Gardnerella | Histidine kinase |
| **K02083** | 3.902248525 | 4.787017949 | 4.96E-21 | 5.79E-20 | Cluster 2: Gardnerella | COG0624 Acetylornithine deacetylase Succinyl-diaminopimelate desuccinylase and related deacylases |
| **K08168** | 9.812624142 | 4.739692856 | 2.53E-26 | 7.48E-25 | Cluster 2: Gardnerella | Major Facilitator Superfamily |
| **K15515** | 16.21269508 | 4.720212511 | 3.13E-29 | 1.70E-27 | Cluster 2: Gardnerella | belongs to the iron- containing alcohol dehydrogenase family |
| **K07271** | 6.080162619 | 4.716556035 | 3.20E-25 | 7.56E-24 | Cluster 2: Gardnerella | LicD family |

| | | | | | | |
|---|---|---|---|---|---|---|
| **K01858** | 7.982304387 | 4.713402973 | 2.32E-31 | 2.41E-29 | Cluster 2: Gardnerella | Myo-inositol-1-phosphate synthase |
| **K16328** | 6.397656858 | 4.688928592 | 5.66E-39 | 3.38E-36 | Cluster 2: Gardnerella | pfkB family carbohydrate kinase |
| **K01466** | 3.974542387 | 4.669553086 | 5.83E-25 | 1.32E-23 | Cluster 2: Gardnerella | Catalyzes the conversion of allantoin (5-ureidohydantoin) to allantoic acid by hydrolytic cleavage of the five-member hydantoin ring |
| **K18120** | 4.913071064 | 4.669050407 | 2.67E-27 | 9.26E-26 | Cluster 2: Gardnerella | Iron-containing alcohol dehydrogenase |
| **K11733** | 10.17918553 | 4.652214118 | 7.68E-40 | 6.12E-37 | Cluster 2: Gardnerella | amino acid |
| **K05341** | 11.62562431 | 4.649744173 | 1.60E-27 | 5.98E-26 | Cluster 2: Gardnerella | Alpha-amylase domain |
| **K11533** | 47.42341746 | 4.592950824 | 4.25E-29 | 2.16E-27 | Cluster 2: Gardnerella | Beta-ketoacyl synthase, C-terminal domain |
| **K21688** | 15.01187843 | 4.576881225 | 2.55E-29 | 1.56E-27 | Cluster 2: Gardnerella | G5 |
| **K07589** | 3.400194075 | 4.55421751 | 1.04E-20 | 1.15E-19 | Cluster 2: Gardnerella | Catalyzes the conversion of 7,8-dihydroneopterin to 6-hydroxymethyl-7,8-dihydropterin |
| **K03153** | 7.298376976 | 4.530211624 | 8.63E-31 | 7.93E-29 | Cluster 2: Gardnerella | Glycine oxidase |
| **K00632** | 7.790402898 | 4.518854434 | 3.32E-35 | 5.29E-33 | Cluster 2: Gardnerella | Thiolase, C-terminal domain |
| **K02050** | 9.586081269 | 4.507938409 | 1.66E-30 | 1.41E-28 | Cluster 2: Gardnerella | Binding-protein-dependent transport system inner membrane component |

| | | | | | | |
|---|---|---|---|---|---|---|
| **K03313** | 8.888013888 | 4.493582925 | 3.09E-29 | 1.70E-27 | Cluster 2: Gardnerella | Na( ) H( ) antiporter that extrudes sodium in exchange for external protons |
| **K07862** | 8.172907778 | 4.468013816 | 7.39E-29 | 3.61E-27 | Cluster 2: Gardnerella | Involved in the import of serine and threonine into the cell, with the concomitant import of sodium (symport system) |
| **K07084** | 8.043530842 | 4.408923061 | 3.12E-32 | 3.72E-30 | Cluster 2: Gardnerella | Na H antiporter family protein |
| **K00878** | 10.38444963 | 4.408322238 | 7.18E-34 | 1.01E-31 | Cluster 2: Gardnerella | Catalyzes the phosphorylation of the hydroxyl group of 4-methyl-5-beta-hydroxyethylthiazole (THZ) |
| **K15553** | 3.926244167 | 4.401520656 | 3.25E-31 | 3.24E-29 | Cluster 2: Gardnerella | NMT1-like family |
| **K01697** | 5.679173928 | 4.396270992 | 2.60E-27 | 9.13E-26 | Cluster 2: Gardnerella | Pyridoxal-phosphate dependent enzyme |
| **K06287** | 9.943387715 | 4.370061268 | 3.70E-29 | 1.96E-27 | Cluster 2: Gardnerella | Maf-like protein |
| **K07161** | 2.633388255 | 4.35233966 | 1.54E-18 | 1.27E-17 | Cluster 2: Gardnerella | Transposase |
| **K03149** | 7.106125412 | 4.333919525 | 5.91E-29 | 2.94E-27 | Cluster 2: Gardnerella | Catalyzes the rearrangement of 1-deoxy-D-xylulose 5-phosphate (DXP) to produce the thiazole phosphate moiety of thiamine. Sulfur is provided by the thiocarboxylate moiety of the carrier protein ThiS. In vitro, sulfur can be provided by H(2)S |
| **K14475** | 2.711615214 | 4.331823726 | 2.65E-13 | 1.35E-12 | Cluster 2: Gardnerella | energy transducer activity |
| **K10975** | 3.003945077 | 4.322920208 | 3.64E-17 | 2.57E-16 | Cluster 2: Gardnerella | Permease for cytosine/purines, uracil, thiamine, allantoin |

| | | | | | | |
|---|---|---|---|---|---|---|
| **K03387** | 11.48134516 | 4.318439084 | 1.14E-36 | 3.03E-34 | Cluster 2: Gardnerella | Thioredoxin domain |
| **K06999** | 5.053254637 | 4.308138547 | 5.68E-26 | 1.54E-24 | Cluster 2: Gardnerella | Phospholipase/Carboxylesterase |
| **K03525** | 5.076477673 | 4.305063808 | 2.18E-27 | 7.78E-26 | Cluster 2: Gardnerella | Catalyzes the phosphorylation of pantothenate (Pan), the first step in CoA biosynthesis |
| **K00705** | 31.00257713 | 4.293664924 | 2.82E-29 | 1.61E-27 | Cluster 2: Gardnerella | 4-alpha-glucanotransferase |
| **K01991** | 2.740644204 | 4.289176311 | 8.37E-24 | 1.48E-22 | Cluster 2: Gardnerella | Transposase |
| **K14153** | 8.818583933 | 4.266579645 | 9.15E-31 | 8.10E-29 | Cluster 2: Gardnerella | Phosphomethylpyrimidine kinase |
| **K14170** | 5.170851758 | 4.238697502 | 7.45E-26 | 1.98E-24 | Cluster 2: Gardnerella | Prephenate dehydratase |
| **K20859** | 5.510023374 | 4.238441915 | 5.95E-27 | 1.82E-25 | Cluster 2: Gardnerella | DNA polymerase alpha chain like domain |
| **K08681** | 2.776528352 | 4.236350148 | 1.78E-30 | 1.43E-28 | Cluster 2: Gardnerella | Catalyzes the hydrolysis of glutamine to glutamate and ammonia as part of the biosynthesis of pyridoxal 5'-phosphate. The resulting ammonia molecule is channeled to the active site of PdxS |
| **K00651** | 6.930445295 | 4.232551976 | 3.36E-23 | 5.47E-22 | Cluster 2: Gardnerella | Transfers an acetyl group from acetyl-CoA to L-homoserine, forming acetyl-L-homoserine |
| **K10907** | 6.507360444 | 4.220578354 | 7.06E-30 | 4.96E-28 | Cluster 2: Gardnerella | Aminotransferase class I and II |
| **K00077** | 8.042121484 | 4.211575818 | 3.32E-30 | 2.48E-28 | Cluster 2: Gardnerella | Catalyzes the NADPH-dependent reduction of ketopantoate into pantoic acid |

| | | | | | | |
|---|---|---|---|---|---|---|
| **K05343** | 12.02888496 | 4.210406236 | 1.94E-30 | 1.49E-28 | Cluster 2: Gardnerella | Alpha-amylase domain |
| **K00936** | 7.480441399 | 4.187371099 | 1.73E-22 | 2.41E-21 | Cluster 2: Gardnerella | ATPase histidine kinase DNA gyrase B HSP90 domain protein |
| **K02049** | 9.290590663 | 4.185201921 | 1.48E-28 | 6.79E-27 | Cluster 2: Gardnerella | ATPases associated with a variety of cellular activities |
| **K01176** | 31.99107582 | 4.169941169 | 1.56E-27 | 5.91E-26 | Cluster 2: Gardnerella | Aamy_C |
| **K07053** | 5.703893387 | 4.156623334 | 1.94E-28 | 8.44E-27 | Cluster 2: Gardnerella | DNA polymerase alpha chain like domain |
| **K00982** | 14.58160799 | 4.141836791 | 3.18E-25 | 7.56E-24 | Cluster 2: Gardnerella | Involved in the regulation of glutamine synthetase GlnA, a key enzyme in the process to assimilate ammonia. When cellular nitrogen levels are high, the C-terminal adenylyl transferase (AT) inactivates GlnA by covalent transfer of an adenylyl group from ATP to specific tyrosine residue of GlnA, thus reducing its activity. Conversely, when nitrogen levels are low, the N-terminal adenylyl removase (AR) activates GlnA by removing the adenylyl group by phosphorolysis, increasing its activity. The regulatory region of GlnE binds the signal transduction protein PII (GlnB) which indicates the nitrogen status of the cell |
| **K13829** | 8.324601871 | 4.12291237 | 1.37E-29 | 8.83E-28 | Cluster 2: Gardnerella | Catalyzes the conversion of 3-deoxy-D-arabino-heptulosonate 7-phosphate (DAHP) to dehydroquinate (DHQ) |
| **K07089** | 7.277370362 | 4.120512771 | 1.31E-27 | 5.13E-26 | Cluster 2: Gardnerella | Predicted permease |

| | | | | | | |
|---|---|---|---|---|---|---|
| **K01950** | 9.022653612 | 4.113342928 | 8.21E-31 | 7.85E-29 | Cluster 2: Gardnerella | Catalyzes the ATP-dependent amidation of deamido-NAD to form NAD. Uses L-glutamine as a nitrogen source |
| **K05838** | 6.035553584 | 4.110430555 | 1.26E-22 | 1.83E-21 | Cluster 2: Gardnerella | Tetratricopeptide repeat |
| **K10007** | 5.079397203 | 4.096419744 | 5.54E-23 | 8.60E-22 | Cluster 2: Gardnerella | Binding-protein-dependent transport system inner membrane component |
| **K15634** | 4.359224536 | 4.093846108 | 5.85E-25 | 1.32E-23 | Cluster 2: Gardnerella | Phosphoglycerate mutase family |
| **K14215** | 4.305542781 | 4.089221904 | 1.73E-28 | 7.67E-27 | Cluster 2: Gardnerella | Catalyzes the condensation of isopentenyl diphosphate (IPP) with allylic pyrophosphates generating different type of terpenoids |
| **K21273** | 4.305542781 | 4.089221904 | 1.73E-28 | 7.67E-27 | Cluster 2: Gardnerella | Catalyzes the condensation of isopentenyl diphosphate (IPP) with allylic pyrophosphates generating different type of terpenoids |
| **K01214** | 22.52893994 | 4.079070956 | 2.83E-29 | 1.61E-27 | Cluster 2: Gardnerella | Belongs to the glycosyl hydrolase 13 family |
| **K05810** | 6.818906218 | 4.073254556 | 2.38E-28 | 1.01E-26 | Cluster 2: Gardnerella | Multi-copper polyphenol oxidoreductase laccase |
| **K03719** | 2.167557831 | 4.045634053 | 5.10E-27 | 1.63E-25 | Cluster 2: Gardnerella | helix_turn_helix ASNC type |
| **K07243** | 11.89313048 | 4.040429439 | 4.95E-26 | 1.36E-24 | Cluster 2: Gardnerella | Iron permease FTR1 family |
| **K00240** | 4.834656568 | 4.012273799 | 5.91E-27 | 1.82E-25 | Cluster 2: Gardnerella | 4Fe-4S dicluster domain |

| | | | | | | |
|---|---|---|---|---|---|---|
| **K01389** | 12.81423967 | 3.997407135 | 5.17E-24 | 9.73E-23 | Cluster 2: Gardnerella | Peptidase family M13 |
| **K01415** | 12.90531224 | 3.995395668 | 6.20E-24 | 1.12E-22 | Cluster 2: Gardnerella | Peptidase family M13 |
| **K01735** | 9.901947949 | 3.994746315 | 9.06E-32 | 9.84E-30 | Cluster 2: Gardnerella | Catalyzes the conversion of 3-deoxy-D-arabino-heptulosonate 7-phosphate (DAHP) to dehydroquinate (DHQ) |
| **K03980** | 26.36095367 | 3.983166416 | 4.74E-27 | 1.53E-25 | Cluster 2: Gardnerella | serine threonine protein kinase |
| **K00812** | 27.70954685 | 3.976095749 | 1.14E-28 | 5.32E-27 | Cluster 2: Gardnerella | Aminotransferase, class I II |
| **K22010** | 2.522347529 | 3.968899877 | 7.63E-25 | 1.70E-23 | Cluster 2: Gardnerella | Response regulator receiver domain protein |
| **K08372** | 9.751905983 | 3.953671007 | 9.03E-24 | 1.57E-22 | Cluster 2: Gardnerella | Domain present in PSD-95, Dlg, and ZO-1/2. |
| **K08776** | 16.17805572 | 3.952742337 | 1.04E-23 | 1.77E-22 | Cluster 2: Gardnerella | Peptidase family M1 domain |
| **K01354** | 12.29070776 | 3.94786011 | 3.51E-22 | 4.67E-21 | Cluster 2: Gardnerella | Peptidase, S9A B C family, catalytic domain protein |
| **K00528** | 11.27518125 | 3.944118332 | 8.94E-24 | 1.57E-22 | Cluster 2: Gardnerella | Pyridine nucleotide-disulphide oxidoreductase |
| **K01802** | 7.430773032 | 3.93950272 | 1.50E-27 | 5.79E-26 | Cluster 2: Gardnerella | FKBP-type peptidyl-prolyl cis-trans isomerase |
| **K06215** | 6.8965769 | 3.937102305 | 3.79E-27 | 1.24E-25 | Cluster 2: Gardnerella | Catalyzes the formation of pyridoxal 5'-phosphate from ribose 5-phosphate (RBP), glyceraldehyde 3-phosphate (G3P) and ammonia. The ammonia is |

| | | | | | | |
|---|---|---|---|---|---|---|
| | | | | | | provided by the PdxT subunit. Can also use ribulose 5-phosphate and dihydroxyacetone phosphate as substrates, resulting from enzyme-catalyzed isomerization of RBP and G3P, respectively |
| **K07114** | 11.57689339 | 3.929181464 | 3.62E-26 | 1.03E-24 | Cluster 2: Gardnerella | von Willebrand factor (vWF) type A domain |
| **K03684** | 7.180550905 | 3.921941984 | 5.77E-25 | 1.32E-23 | Cluster 2: Gardnerella | 3'-5' exonuclease |
| **K03628** | 12.88422744 | 3.918366277 | 4.10E-29 | 2.13E-27 | Cluster 2: Gardnerella | Facilitates transcription termination by a mechanism that involves Rho binding to the nascent RNA, activation of Rho's RNA-dependent ATPase activity, and release of the mRNA from the DNA template |
| **K08300** | 15.660514 | 3.905870478 | 2.77E-24 | 5.61E-23 | Cluster 2: Gardnerella | Ribonuclease E/G family |
| **K02823** | 3.055820078 | 3.903294152 | 1.56E-25 | 3.96E-24 | Cluster 2: Gardnerella | Iron-sulfur cluster binding domain of dihydroorotate dehydrogenase B |
| **K10005** | 3.785653657 | 3.903015953 | 2.65E-19 | 2.42E-18 | Cluster 2: Gardnerella | Belongs to the bacterial solute-binding protein 3 family |
| **K00937** | 16.98443709 | 3.902834081 | 2.83E-29 | 1.61E-27 | Cluster 2: Gardnerella | Catalyzes the reversible transfer of the terminal phosphate of ATP to form a long-chain polyphosphate (polyP) |
| **K09760** | 9.874025518 | 3.901682969 | 9.67E-25 | 2.12E-23 | Cluster 2: Gardnerella | RmuC family |
| **K07442** | 5.270295078 | 3.894207874 | 1.07E-22 | 1.60E-21 | Cluster 2: Gardnerella | Catalyzes the S-adenosyl-L-methionine-dependent formation of N(1)-methyladenine at position 58 (m1A58) in tRNA |

| | | | | | | |
|---|---|---|---|---|---|---|
| **K03147** | 11.88978248 | 3.884436914 | 5.03E-23 | 7.86E-22 | Cluster 2: Gardnerella | Catalyzes the synthesis of the hydroxymethylpyrimidine phosphate (HMP-P) moiety of thiamine from aminoimidazole ribotide (AIR) in a radical S-adenosyl-L-methionine (SAM)-dependent reaction |
| **K02006** | 2.824124236 | 3.877704466 | 9.22E-22 | 1.17E-20 | Cluster 2: Gardnerella | Psort location CytoplasmicMembrane, score |
| **K22306** | 5.176662917 | 3.874692319 | 1.66E-22 | 2.34E-21 | Cluster 2: Gardnerella | Phosphoglycerate mutase family |
| **K15554** | 2.173805096 | 3.874160033 | 6.82E-28 | 2.81E-26 | Cluster 2: Gardnerella | Binding-protein-dependent transport system inner membrane component |
| **K03578** | 18.42337041 | 3.85949856 | 2.05E-26 | 6.20E-25 | Cluster 2: Gardnerella | Helicase associated domain (HA2) Add an annotation |
| **K09761** | 4.946807883 | 3.838307664 | 3.77E-27 | 1.24E-25 | Cluster 2: Gardnerella | Specifically methylates the N3 position of the uracil ring of uridine 1498 (m3U1498) in 16S rRNA. Acts on the fully assembled 30S ribosomal subunit |
| **K09118** | 12.80535083 | 3.837649499 | 1.46E-22 | 2.09E-21 | Cluster 2: Gardnerella | Uncharacterised protein family (UPF0182) |
| **K08301** | 16.28763335 | 3.835266191 | 2.20E-24 | 4.50E-23 | Cluster 2: Gardnerella | Ribonuclease E/G family |
| **K01880** | 9.469850417 | 3.834350578 | 9.11E-25 | 2.02E-23 | Cluster 2: Gardnerella | Catalyzes the attachment of glycine to tRNA(Gly) |
| **K01885** | 22.51726668 | 3.82567164 | 4.49E-23 | 7.11E-22 | Cluster 2: Gardnerella | Catalyzes the attachment of glutamate to tRNA(Glu) in a two-step reaction glutamate is first activated by ATP to form Glu-AMP and then transferred to the acceptor end of tRNA(Glu) |

| | | | | | | |
|---|---|---|---|---|---|---|
| **K00773** | 13.02509112 | 3.821146365 | 2.36E-26 | 7.04E-25 | Cluster 2: Gardnerella | Catalyzes the base-exchange of a guanine (G) residue with the queuine precursor 7-aminomethyl-7-deazaguanine (PreQ1) at position 34 (anticodon wobble position) in tRNAs with GU(N) anticodons (tRNA-Asp, -Asn, -His and -Tyr). Catalysis occurs through a double-displacement mechanism. The nucleophile active site attacks the C1' of nucleotide 34 to detach the guanine base from the RNA, forming a covalent enzyme-RNA intermediate. The proton acceptor active site deprotonates the incoming PreQ1, allowing a nucleophilic attack on the C1' of the ribose to form the product. After dissociation, two additional enzymatic reactions on the tRNA convert PreQ1 to queuine (Q), resulting in the hypermodified nucleoside queuosine (7-(((4,5-cis-dihydroxy-2- cyclopenten-1-yl)amino)methyl)-7-deazaguanosine) |
| **K07452** | 2.865449267 | 3.820554953 | 9.39E-15 | 5.49E-14 | Cluster 2: Gardnerella | AAA domain (dynein-related subfamily) |
| **K03727** | 10.95955755 | 3.817568498 | 4.22E-23 | 6.78E-22 | Cluster 2: Gardnerella | DEAD DEAH box helicase |
| **K00688** | 22.18450279 | 3.816415566 | 3.43E-24 | 6.84E-23 | Cluster 2: Gardnerella | Phosphorylase is an important allosteric enzyme in carbohydrate metabolism. Enzymes from different sources differ in their regulatory mechanisms and in their natural substrates. However, all known phosphorylases share catalytic and structural properties |
| **K05364** | 8.359102515 | 3.812308072 | 1.30E-22 | 1.88E-21 | Cluster 2: Gardnerella | penicillin-binding protein |

| | | | | | | |
|---|---|---|---|---|---|---|
| **K00556** | 4.769713849 | 3.810346836 | 1.02E-25 | 2.65E-24 | Cluster 2: Gardnerella | RNA methyltransferase TrmH family |
| **K06188** | 2.501745577 | 3.799786317 | 3.45E-18 | 2.77E-17 | Cluster 2: Gardnerella | Belongs to the MIP aquaporin (TC 1.A.8) family |
| **K08744** | 4.503251053 | 3.798738828 | 4.18E-23 | 6.75E-22 | Cluster 2: Gardnerella | Belongs to the CDP-alcohol phosphatidyltransferase class-I family |
| **K03587** | 12.98539833 | 3.785470666 | 1.54E-24 | 3.22E-23 | Cluster 2: Gardnerella | Penicillin-binding protein, transpeptidase domain protein |
| **K13787** | 6.036433838 | 3.78403001 | 5.64E-27 | 1.78E-25 | Cluster 2: Gardnerella | Belongs to the FPP GGPP synthase family |
| **K02334** | 8.976355904 | 3.775505526 | 4.19E-15 | 2.49E-14 | Cluster 2: Gardnerella | DNA-directed DNA polymerase |
| **K00210** | 4.984779776 | 3.772369332 | 1.22E-22 | 1.79E-21 | Cluster 2: Gardnerella | Prephenate dehydrogenase |
| **K01633** | 3.702555946 | 3.768410992 | 1.41E-19 | 1.32E-18 | Cluster 2: Gardnerella | Catalyzes the conversion of 7,8-dihydroneopterin to 6-hydroxymethyl-7,8-dihydropterin |
| **K11263** | 11.71034308 | 3.76530463 | 1.40E-19 | 1.31E-18 | Cluster 2: Gardnerella | Carbamoyl-phosphate synthase L chain, ATP binding domain protein |
| **K01139** | 11.40452062 | 3.759752773 | 4.47E-23 | 7.11E-22 | Cluster 2: Gardnerella | In eubacteria ppGpp (guanosine 3'-diphosphate 5'-diphosphate) is a mediator of the stringent response that coordinates a variety of cellular activities in response to changes in nutritional abundance |
| **K00970** | 11.60784564 | 3.752807429 | 2.30E-23 | 3.79E-22 | Cluster 2: Gardnerella | Probable RNA and SrmB- binding site of polymerase A |
| **K10213** | 5.373985324 | 3.735904502 | 1.10E-19 | 1.06E-18 | Cluster 2: Gardnerella | Inosine-uridine preferring nucleoside hydrolase |

| | | | | | | |
|---|---|---|---|---|---|---|
| **K09812** | 5.391541578 | 3.732886124 | 3.04E-26 | 8.77E-25 | Cluster 2: Gardnerella | Cell division ATP-binding protein FtsE |
| **K00962** | 18.98943674 | 3.732502899 | 4.40E-26 | 1.24E-24 | Cluster 2: Gardnerella | Involved in mRNA degradation. Catalyzes the phosphorolysis of single-stranded polyribonucleotides processively in the 3'- to 5'-direction |
| **K00817** | 10.46180921 | 3.730008197 | 8.83E-29 | 4.22E-27 | Cluster 2: Gardnerella | Aminotransferase class I and II |
| **K03924** | 5.334142674 | 3.729370202 | 4.73E-24 | 8.97E-23 | Cluster 2: Gardnerella | ATPase family associated with various cellular activities (AAA) |
| **K07259** | 7.636910722 | 3.726354138 | 2.74E-23 | 4.49E-22 | Cluster 2: Gardnerella | D-Ala-D-Ala carboxypeptidase 3 (S13) family |
| **K12524** | 8.530269842 | 3.726087655 | 4.53E-24 | 8.67E-23 | Cluster 2: Gardnerella | Homoserine dehydrogenase |
| **K03386** | 4.032022171 | 3.725751174 | 1.29E-27 | 5.12E-26 | Cluster 2: Gardnerella | C-terminal domain of 1-Cys peroxiredoxin |
| **K03695** | 22.25177285 | 3.718482934 | 9.94E-24 | 1.71E-22 | Cluster 2: Gardnerella | Part of a stress-induced multi-chaperone system, it is involved in the recovery of the cell from heat-induced damage, in cooperation with DnaK, DnaJ and GrpE |
| **K01897** | 31.85491233 | 3.715864038 | 3.98E-24 | 7.73E-23 | Cluster 2: Gardnerella | AMP-binding enzyme |
| **K01494** | 3.813618017 | 3.712580587 | 1.84E-22 | 2.54E-21 | Cluster 2: Gardnerella | Belongs to the dCTP deaminase family |
| **K13640** | 2.749963058 | 3.708859098 | 1.55E-24 | 3.22E-23 | Cluster 2: Gardnerella | transcriptional regulator, MerR family |
| **K08289** | 7.428967296 | 3.707995059 | 6.86E-22 | 8.86E-21 | Cluster 2: Gardnerella | Involved in the de novo purine biosynthesis. Catalyzes the transfer of formate to 5-phospho-ribosyl- |

| | | | | | | |
|---|---|---|---|---|---|---|
| | | | | | | glycinamide (GAR), producing 5-phospho-ribosyl-N-formylglycinamide (FGAR). Formate is provided by PurU via hydrolysis of 10-formyl-tetrahydrofolate |
| **K02283** | 9.827581779 | 3.705178223 | 8.43E-23 | 1.27E-21 | Cluster 2: Gardnerella | Type II IV secretion system protein |
| **K01673** | 2.931552531 | 3.704858816 | 4.91E-22 | 6.44E-21 | Cluster 2: Gardnerella | Reversible hydration of carbon dioxide |
| **K01524** | 10.3636774 | 3.701706562 | 9.05E-26 | 2.38E-24 | Cluster 2: Gardnerella | Ppx/GppA phosphatase family |
| **K01155** | 1.581452502 | 3.681981521 | 6.17E-11 | 2.58E-10 | Cluster 2: Gardnerella | Psort location Cytoplasmic, score 8.96 |
| **K14260** | 13.14756594 | 3.6818013 | 6.37E-19 | 5.57E-18 | Cluster 2: Gardnerella | Aminotransferase, class I II |
| **K00989** | 5.109106952 | 3.680759216 | 7.69E-23 | 1.17E-21 | Cluster 2: Gardnerella | Phosphorolytic exoribonuclease that removes nucleotide residues following the -CCA terminus of tRNA and adds nucleotides to the ends of RNA molecules by using nucleoside diphosphates as substrates |
| **K02484** | 8.006306718 | 3.68037544 | 1.05E-20 | 1.16E-19 | Cluster 2: Gardnerella | ATPase histidine kinase DNA gyrase B HSP90 domain protein |
| **K11939** | 5.694681747 | 3.677958315 | 2.57E-21 | 3.13E-20 | Cluster 2: Gardnerella | EamA-like transporter family |
| **K07768** | 7.493632064 | 3.675417863 | 1.46E-19 | 1.35E-18 | Cluster 2: Gardnerella | ATPase histidine kinase DNA gyrase B HSP90 domain protein |
| **K04518** | 5.551478163 | 3.667740064 | 1.16E-24 | 2.52E-23 | Cluster 2: Gardnerella | Prephenate dehydratase |

| | | | | | | |
|---|---|---|---|---|---|---|
| **K13788** | 6.876096739 | 3.663666211 | 1.65E-24 | 3.40E-23 | Cluster 2: Gardnerella | phosphate acetyltransferase |
| **K07739** | 1.175014578 | 3.655691212 | 1.24E-14 | 7.17E-14 | Cluster 2: Gardnerella | histone acetyltransferase, ELP3 family |
| **K15555** | 1.745583408 | 3.645094556 | 7.15E-22 | 9.18E-21 | Cluster 2: Gardnerella | ATPases associated with a variety of cellular activities |
| **K01129** | 7.425518422 | 3.636780532 | 4.76E-26 | 1.32E-24 | Cluster 2: Gardnerella | Phosphohydrolase-associated domain |
| **K11788** | 6.095745664 | 3.628667204 | 2.03E-20 | 2.16E-19 | Cluster 2: Gardnerella | Belongs to the GARS family |
| **K02226** | 4.438316173 | 3.62261079 | 3.85E-21 | 4.58E-20 | Cluster 2: Gardnerella | Phosphoglycerate mutase family |
| **K04095** | 5.734106195 | 3.596848283 | 1.39E-20 | 1.50E-19 | Cluster 2: Gardnerella | Fic/DOC family |
| **K10008** | 2.998108396 | 3.584163085 | 2.81E-19 | 2.55E-18 | Cluster 2: Gardnerella | ATP-binding protein of ABC transporter for glutamate K02028 |
| **K04517** | 5.765516222 | 3.572740211 | 1.96E-25 | 4.87E-24 | Cluster 2: Gardnerella | Prephenate dehydrogenase |
| **K03523** | 3.372327386 | 3.56814677 | 3.45E-22 | 4.61E-21 | Cluster 2: Gardnerella | BioY family |
| **K19427** | 2.680575509 | 3.541694765 | 0.000120634 | 0.000273026 | Cluster 2: Gardnerella | Glycosyl transferase family 2 |
| **K01057** | 3.582772743 | 3.53803907 | 1.10E-18 | 9.26E-18 | Cluster 2: Gardnerella | Glucosamine-6-phosphate isomerases/6-phosphogluconolactonase |

| | | | | | | |
|---|---|---|---|---|---|---|
| **K00931** | 6.448504719 | 3.537060427 | 2.48E-22 | 3.39E-21 | Cluster 2: Gardnerella | Catalyzes the transfer of a phosphate group to glutamate to form L-glutamate 5-phosphate |
| **K15738** | 13.26029477 | 3.534372819 | 1.41E-22 | 2.02E-21 | Cluster 2: Gardnerella | ABC transporter |
| **K00254** | 6.749455442 | 3.528644094 | 3.81E-24 | 7.45E-23 | Cluster 2: Gardnerella | Catalyzes the conversion of dihydroorotate to orotate with quinone as electron acceptor |
| **K09819** | 7.390499433 | 3.524080428 | 6.26E-20 | 6.18E-19 | Cluster 2: Gardnerella | ABC 3 transport family |
| **K09811** | 6.579814816 | 3.500637153 | 6.39E-20 | 6.29E-19 | Cluster 2: Gardnerella | Part of the ABC transporter FtsEX involved in cellular division |
| **K04761** | 3.13737809 | 3.498197942 | 1.21E-19 | 1.15E-18 | Cluster 2: Gardnerella | LysR substrate binding domain protein |
| **K01589** | 8.602521487 | 3.481716184 | 3.80E-22 | 5.02E-21 | Cluster 2: Gardnerella | Catalyzes the ATP-dependent conversion of 5-aminoimidazole ribonucleotide (AIR) and HCO(3)(-) to N5- carboxyaminoimidazole ribonucleotide (N5-CAIR) |
| **K01507** | 3.645264048 | 3.478280776 | 2.32E-21 | 2.84E-20 | Cluster 2: Gardnerella | Catalyzes the hydrolysis of inorganic pyrophosphate (PPi) forming two phosphate ions |
| **K01299** | 1.016805782 | 3.477182382 | 3.43E-09 | 1.21E-08 | Cluster 2: Gardnerella | Broad specificity carboxypetidase that releases amino acids sequentially from the C-terminus, including neutral, aromatic, polar and basic residues |
| **K10006** | 1.524439623 | 3.464105036 | 1.75E-16 | 1.17E-15 | Cluster 2: Gardnerella | Binding-protein-dependent transport system inner membrane component |
| **K01740** | 5.988232018 | 3.452700218 | 2.53E-16 | 1.66E-15 | Cluster 2: Gardnerella | Aminotransferase class-V |

| | | | | | | |
|---|---|---|---|---|---|---|
| **K01736** | 8.336031099 | 3.44644077 | 8.07E-21 | 9.06E-20 | Cluster 2: Gardnerella | Catalyzes the anti-1,4-elimination of the C-3 phosphate and the C-6 proR hydrogen from 5-enolpyruvylshikimate-3-phosphate (EPSP) to yield chorismate, which is the branch point compound that serves as the starting substrate for the three terminal pathways of aromatic amino acid biosynthesis. This reaction introduces a second double bond into the aromatic ring system |
| **K01626** | 9.195528817 | 3.443947237 | 7.42E-21 | 8.44E-20 | Cluster 2: Gardnerella | Stereospecific condensation of phosphoenolpyruvate (PEP) and D-erythrose-4-phosphate (E4P) giving rise to 3-deoxy-D- arabino-heptulosonate-7-phosphate (DAHP) |
| **K03786** | 1.535868305 | 3.443316085 | 1.16E-19 | 1.11E-18 | Cluster 2: Gardnerella | Catalyzes a trans-dehydration via an enolate intermediate |
| **K01940** | 6.081074309 | 3.43470512 | 1.76E-20 | 1.88E-19 | Cluster 2: Gardnerella | Belongs to the argininosuccinate synthase family. Type 1 subfamily |
| **K08296** | 1.677688454 | 3.428158211 | 8.63E-18 | 6.68E-17 | Cluster 2: Gardnerella | Phosphoglycerate mutase family |
| **K07230** | 2.324797663 | 3.425391498 | 1.24E-20 | 1.35E-19 | Cluster 2: Gardnerella | Fe2+ transport protein |
| **K05516** | 8.483483368 | 3.418707965 | 6.08E-24 | 1.11E-22 | Cluster 2: Gardnerella | DnaJ molecular chaperone homology domain |
| **K00656** | 15.90688303 | 3.411532773 | 1.85E-21 | 2.29E-20 | Cluster 2: Gardnerella | Pyruvate formate lyase-like |
| **K07391** | 10.11023337 | 3.402039838 | 7.24E-20 | 7.09E-19 | Cluster 2: Gardnerella | Magnesium chelatase, subunit ChlI C-terminal |

| | | | | | | |
|---|---|---|---|---|---|---|
| **K00239** | 9.742311139 | 3.386370044 | 1.48E-22 | 2.10E-21 | Cluster 2: Gardnerella | Succinate dehydrogenase flavoprotein subunit |
| **K00147** | 7.965439923 | 3.386019827 | 1.19E-22 | 1.75E-21 | Cluster 2: Gardnerella | Catalyzes the NADPH-dependent reduction of L-glutamate 5-phosphate into L-glutamate 5-semialdehyde and phosphate. The product spontaneously undergoes cyclization to form 1-pyrroline-5- carboxylate |
| **K01091** | 9.143132627 | 3.380967391 | 4.48E-32 | 5.09E-30 | Cluster 2: Gardnerella | HAD-hyrolase-like |
| **K00700** | 13.83403717 | 3.378274826 | 5.32E-22 | 6.94E-21 | Cluster 2: Gardnerella | Catalyzes the formation of the alpha-1,6-glucosidic linkages in glycogen by scission of a 1,4-alpha-linked oligosaccharide from growing alpha-1,4-glucan chains and the subsequent attachment of the oligosaccharide to the alpha-1,6 position |
| **K07248** | 1.147694952 | 3.377436245 | 9.38E-11 | 3.87E-10 | Cluster 2: Gardnerella | Belongs to the aldehyde dehydrogenase family |
| **K12297** | 1.548600645 | 3.370902147 | 3.49E-09 | 1.23E-08 | Cluster 2: Gardnerella | Specifically methylates the guanine in position 2445 (m2G2445) and the guanine in position 2069 (m7G2069) of 23S rRNA |
| **K10117** | 20.61853503 | 3.359714341 | 4.26E-21 | 5.01E-20 | Cluster 2: Gardnerella | Bacterial extracellular solute-binding protein |
| **K00640** | 3.784305561 | 3.354884053 | 1.68E-22 | 2.35E-21 | Cluster 2: Gardnerella | Bacterial transferase hexapeptide (six repeats) |
| **K01696** | 6.701643258 | 3.333112208 | 1.12E-15 | 6.99E-15 | Cluster 2: Gardnerella | The beta subunit is responsible for the synthesis of L-tryptophan from indole and L-serine |
| **K07503** | 3.097207097 | 3.330950385 | 3.51E-18 | 2.81E-17 | Cluster 2: Gardnerella | Cleaves both 3' and 5' ssDNA extremities of branched DNA structures |

| | | | | | | |
|---|---|---|---|---|---|---|
| **K01186** | 7.689887996 | 3.330499348 | 7.74E-13 | 3.83E-12 | Cluster 2: Gardnerella | BNR repeat-like domain |
| **K00602** | 11.83186161 | 3.329758355 | 2.44E-22 | 3.35E-21 | Cluster 2: Gardnerella | Bifunctional purine biosynthesis protein PurH |
| **K10227** | 1.925371392 | 3.328129449 | 1.29E-07 | 3.92E-07 | Cluster 2: Gardnerella | Bacterial extracellular solute-binding protein |
| **K21147** | 3.072690603 | 3.316400732 | 1.19E-16 | 8.12E-16 | Cluster 2: Gardnerella | ThiF family |
| **K03593** | 4.414332263 | 3.315602886 | 1.42E-19 | 1.32E-18 | Cluster 2: Gardnerella | Binds and transfers iron-sulfur (Fe-S) clusters to target apoproteins. Can hydrolyze ATP |
| **K03524** | 6.731478383 | 3.310159618 | 4.35E-24 | 8.39E-23 | Cluster 2: Gardnerella | Biotin/lipoate A/B protein ligase family |
| **K00826** | 7.693628181 | 3.306626544 | 1.60E-20 | 1.72E-19 | Cluster 2: Gardnerella | Amino-transferase class IV |
| **K01738** | 7.379876478 | 3.304276466 | 3.71E-24 | 7.34E-23 | Cluster 2: Gardnerella | Pyridoxal-phosphate dependent enzyme |
| **K00999** | 1.430600428 | 3.280172896 | 9.92E-19 | 8.53E-18 | Cluster 2: Gardnerella | CDP-alcohol phosphatidyltransferase |
| **K02897** | 2.876693926 | 3.278100091 | 1.44E-18 | 1.19E-17 | Cluster 2: Gardnerella | This is one of the proteins that binds to the 5S RNA in the ribosome where it forms part of the central protuberance |
| **K00278** | 10.58779661 | 3.25002961 | 2.09E-25 | 5.16E-24 | Cluster 2: Gardnerella | Succinate dehydrogenase flavoprotein subunit |
| **K10119** | 7.123665904 | 3.244473509 | 6.33E-17 | 4.41E-16 | Cluster 2: Gardnerella | ABC transporter permease |

| | | | | | | |
|---|---|---|---|---|---|---|
| **K00616** | 7.092261213 | 3.227822303 | 5.37E-24 | 1.00E-22 | Cluster 2: Gardnerella | Transaldolase is important for the balance of metabolites in the pentose-phosphate pathway |
| **K09820** | 2.886530489 | 3.216021992 | 1.02E-18 | 8.72E-18 | Cluster 2: Gardnerella | ATPases associated with a variety of cellular activities |
| **K00800** | 7.165878308 | 3.215166967 | 3.69E-20 | 3.83E-19 | Cluster 2: Gardnerella | Catalyzes the transfer of the enolpyruvyl moiety of phosphoenolpyruvate (PEP) to the 5-hydroxyl of shikimate-3- phosphate (S3P) to produce enolpyruvyl shikimate-3-phosphate and inorganic phosphate |
| **K13288** | 2.907686981 | 3.214337158 | 2.46E-17 | 1.79E-16 | Cluster 2: Gardnerella | 3'-to-5' exoribonuclease specific for small oligoribonucleotides |
| **K03574** | 10.85340129 | 3.196385621 | 1.08E-37 | 4.31E-35 | Cluster 2: Gardnerella | Phosphoglycerate mutase family |
| **K08325** | 1.205730237 | 3.19280204 | 0.000118102 | 0.000268324 | Cluster 2: Gardnerella | Iron-containing alcohol dehydrogenase |
| **K00297** | 4.593825783 | 3.192166052 | 1.18E-15 | 7.33E-15 | Cluster 2: Gardnerella | Methylenetetrahydrofolate reductase |
| **K07272** | 3.491473306 | 3.18258566 | 2.50E-09 | 8.97E-09 | Cluster 2: Gardnerella | Rhamnan synthesis protein F |
| **K01854** | 11.37383133 | 3.180432845 | 7.92E-22 | 1.01E-20 | Cluster 2: Gardnerella | UDP-galactopyranose mutase |
| **K05524** | 0.83746093 | 3.177848554 | 2.43E-16 | 1.60E-15 | Cluster 2: Gardnerella | 4Fe-4S binding domain |
| **K04069** | 6.215685315 | 3.17082759 | 4.11E-19 | 3.67E-18 | Cluster 2: Gardnerella | Activation of pyruvate formate-lyase under anaerobic conditions by generation of an organic free radical, using S- adenosylmethionine and reduced flavodoxin as cosubstrates to produce 5'-deoxy-adenosine |

| | | | | | | |
|---|---|---|---|---|---|---|
| **K06910** | 2.403582992 | 3.154754003 | 1.26E-15 | 7.81E-15 | Cluster 2: Gardnerella | Phosphatidylethanolamine-binding protein |
| **K16089** | 1.947655495 | 3.126854341 | 5.42E-07 | 1.55E-06 | Cluster 2: Gardnerella | TonB-dependent receptor |
| **K12257** | 1.452759112 | 3.126311906 | 2.21E-09 | 7.99E-09 | Cluster 2: Gardnerella | Part of the Sec protein translocase complex. Interacts with the SecYEG preprotein conducting channel. SecDF uses the proton motive force (PMF) to complete protein translocation after the ATP-dependent function of SecA |
| **K07460** | 1.239521878 | 3.122324101 | 1.38E-13 | 7.22E-13 | Cluster 2: Gardnerella | Belongs to the UPF0102 family |
| **K01952** | 22.28493534 | 3.098012713 | 7.40E-20 | 7.22E-19 | Cluster 2: Gardnerella | CobB/CobQ-like glutamine amidotransferase domain |
| **K01755** | 6.716326498 | 3.092658998 | 3.92E-19 | 3.51E-18 | Cluster 2: Gardnerella | argininosuccinate lyase |
| **K01092** | 5.603399284 | 3.092206711 | 4.19E-18 | 3.32E-17 | Cluster 2: Gardnerella | Inositol monophosphatase family |
| **K02032** | 41.55196709 | 3.071444541 | 7.27E-44 | 1.74E-40 | Cluster 2: Gardnerella | Belongs to the ABC transporter superfamily |
| **K03402** | 2.052931163 | 3.071141251 | 4.93E-17 | 3.48E-16 | Cluster 2: Gardnerella | Regulates arginine biosynthesis genes |
| **K10441** | 9.550942005 | 3.066000272 | 3.07E-16 | 2.00E-15 | Cluster 2: Gardnerella | ATPases associated with a variety of cellular activities |
| **K06346** | 3.565039591 | 3.060839374 | 1.85E-16 | 1.23E-15 | Cluster 2: Gardnerella | Putative single-stranded nucleic acids-binding domain |

| | | | | | | |
|---|---|---|---|---|---|---|
| **K07088** | 11.37209973 | 3.05935398 | 3.52E-17 | 2.49E-16 | Cluster 2: Gardnerella | Transporter, auxin efflux carrier (AEC) family protein |
| **K19180** | 1.571239157 | 3.049176597 | 4.88E-09 | 1.70E-08 | Cluster 2: Gardnerella | NAD dependent epimerase/dehydratase family |
| **K12341** | 0.789292801 | 3.047100275 | 0.000214838 | 0.000473238 | Cluster 2: Gardnerella | Hep Hag repeat protein |
| **K00764** | 9.580177419 | 3.033123385 | 1.02E-17 | 7.72E-17 | Cluster 2: Gardnerella | Catalyzes the formation of phosphoribosylamine from phosphoribosylpyrophosphate (PRPP) and glutamine |
| **K07284** | 43.23138119 | 3.021798186 | 4.61E-38 | 2.20E-35 | Cluster 2: Gardnerella | Sortase family |
| **K10440** | 5.810950781 | 3.010705698 | 4.23E-13 | 2.13E-12 | Cluster 2: Gardnerella | Branched-chain amino acid transport system / permease component |
| **K00024** | 4.820744395 | 2.999713424 | 1.52E-16 | 1.03E-15 | Cluster 2: Gardnerella | Belongs to the LDH MDH superfamily. LDH family |
| **K10548** | 1.398917026 | 2.959337099 | 5.57E-06 | 1.43E-05 | Cluster 2: Gardnerella | ATPases associated with a variety of cellular activities |
| **K01923** | 4.030813961 | 2.946751272 | 2.12E-18 | 1.71E-17 | Cluster 2: Gardnerella | Belongs to the SAICAR synthetase family |
| **K10439** | 10.53224444 | 2.939298929 | 1.06E-14 | 6.19E-14 | Cluster 2: Gardnerella | Periplasmic binding protein domain |
| **K00615** | 14.74955246 | 2.935437329 | 4.07E-20 | 4.19E-19 | Cluster 2: Gardnerella | Belongs to the transketolase family |
| **K09809** | 3.497265326 | 2.931496842 | 3.19E-07 | 9.33E-07 | Cluster 2: Gardnerella | Glycosyl transferase family 2 |

| | | | | | | |
|---|---|---|---|---|---|---|
| **K00886** | 2.575214875 | 2.927947616 | 2.56E-14 | 1.43E-13 | Cluster 2: Gardnerella | ROK family |
| **K10118** | 5.224925961 | 2.920138953 | 1.32E-12 | 6.45E-12 | Cluster 2: Gardnerella | Binding-protein-dependent transport system inner membrane component |
| **K01588** | 1.688211415 | 2.919596346 | 7.11E-18 | 5.53E-17 | Cluster 2: Gardnerella | Catalyzes the conversion of N5-carboxyaminoimidazole ribonucleotide (N5-CAIR) to 4-carboxy-5-aminoimidazole ribonucleotide (CAIR) |
| **K10547** | 1.140883944 | 2.911272777 | 1.59E-07 | 4.77E-07 | Cluster 2: Gardnerella | Branched-chain amino acid transport system / permease component |
| **K04070** | 0.877943357 | 2.910143491 | 5.65E-09 | 1.95E-08 | Cluster 2: Gardnerella | Radical SAM domain protein |
| **K21600** | 1.022509571 | 2.908510335 | 3.70E-14 | 2.03E-13 | Cluster 2: Gardnerella | Metal-sensitive transcriptional repressor |
| **K06997** | 4.2378983 | 2.891716449 | 2.58E-17 | 1.87E-16 | Cluster 2: Gardnerella | Pyridoxal 5'-phosphate (PLP)-binding protein, which is involved in PLP homeostasis |
| **K12510** | 2.577771111 | 2.891181103 | 8.60E-13 | 4.24E-12 | Cluster 2: Gardnerella | Type ii secretion system |
| **K01159** | 1.929059893 | 2.881558696 | 2.39E-13 | 1.22E-12 | Cluster 2: Gardnerella | Nuclease that resolves Holliday junction intermediates in genetic recombination. Cleaves the cruciform structure in supercoiled DNA by nicking to strands with the same polarity at sites symmetrically opposed at the junction in the homologous arms and leaves a 5'-terminal phosphate and a 3'-terminal hydroxyl group |
| **K21449** | 7.090436592 | 2.878441075 | 3.93E-11 | 1.67E-10 | Cluster 2: Gardnerella | Hep Hag repeat protein |

| | | | | | | |
|---|---|---|---|---|---|---|
| **K20444** | 1.09153426 | 2.857522323 | 7.74E-09 | 2.64E-08 | Cluster 2: Gardnerella | Transposase |
| **K02012** | 3.033896023 | 2.83942607 | 1.46E-08 | 4.85E-08 | Cluster 2: Gardnerella | Tat pathway signal sequence domain protein |
| **K03767** | 3.3656648 | 2.832509986 | 1.41E-14 | 8.13E-14 | Cluster 2: Gardnerella | PPIases accelerate the folding of proteins. It catalyzes the cis-trans isomerization of proline imidic peptide bonds in oligopeptides |
| **K13888** | 0.933496624 | 2.829205839 | 7.02E-09 | 2.41E-08 | Cluster 2: Gardnerella | Belongs to the membrane fusion protein (MFP) (TC 8.A.1) family |
| **K17329** | 0.963125556 | 2.815103081 | 2.79E-06 | 7.39E-06 | Cluster 2: Gardnerella | Bacterial extracellular solute-binding protein |
| **K07574** | 1.000427941 | 2.807036803 | 1.05E-10 | 4.30E-10 | Cluster 2: Gardnerella | CRS1_YhbY |
| **K02282** | 0.910815425 | 2.792658075 | 1.27E-07 | 3.87E-07 | Cluster 2: Gardnerella | CobQ/CobB/MinD/ParA nucleotide binding domain |
| **K03737** | 5.336782698 | 2.792577874 | 3.54E-11 | 1.52E-10 | Cluster 2: Gardnerella | Oxidoreductase required for the transfer of electrons from pyruvate to flavodoxin |
| **K01361** | 37.23222856 | 2.789771383 | 1.13E-12 | 5.57E-12 | Cluster 2: Gardnerella | Belongs to the peptidase S8 family |
| **K02021** | 3.050842343 | 2.778921957 | 1.25E-08 | 4.17E-08 | Cluster 2: Gardnerella | ABC transporter, ATP-binding protein |
| **K07736** | 2.033650216 | 2.765906515 | 3.42E-12 | 1.62E-11 | Cluster 2: Gardnerella | CarD-like/TRCF domain |
| **K16785** | 40.68747928 | 2.752055418 | 7.61E-41 | 9.09E-38 | Cluster 2: Gardnerella | ATPases associated with a variety of cellular activities |

| | | | | | | |
|---|---|---|---|---|---|---|
| **K01945** | 9.13977608 | 2.742206137 | 3.05E-14 | 1.69E-13 | Cluster 2: Gardnerella | Belongs to the GARS family |
| **K16650** | 3.09666103 | 2.73826759 | 3.48E-06 | 9.15E-06 | Cluster 2: Gardnerella | Glycosyltransferase like family 2 |
| **K13572** | 1.405439805 | 2.736795149 | 5.65E-10 | 2.14E-09 | Cluster 2: Gardnerella | WYL domain |
| **K09161** | 0.611479796 | 2.724106402 | 0.027213971 | 0.045333119 | Cluster 2: Gardnerella | YfbU domain |
| **K00014** | 4.764115896 | 2.69266332 | 7.19E-17 | 4.98E-16 | Cluster 2: Gardnerella | Shikimate dehydrogenase substrate binding domain |
| **K20490** | 1.893502829 | 2.685609556 | 2.74E-08 | 8.81E-08 | Cluster 2: Gardnerella | ABC transporter |
| **K01710** | 6.804221698 | 2.672888575 | 3.31E-14 | 1.82E-13 | Cluster 2: Gardnerella | Belongs to the NAD(P)-dependent epimerase dehydratase family. dTDP-glucose dehydratase subfamily |
| **K02008** | 0.692376814 | 2.663277481 | 9.81E-06 | 2.46E-05 | Cluster 2: Gardnerella | transmembrane transporter activity |
| **K10546** | 0.931730858 | 2.659373814 | 7.83E-06 | 1.98E-05 | Cluster 2: Gardnerella | Periplasmic binding protein domain |
| **K03575** | 3.744698178 | 2.657696762 | 2.46E-11 | 1.08E-10 | Cluster 2: Gardnerella | FES |
| **K09803** | 0.611556125 | 2.65379385 | 0.001369233 | 0.002756921 | Cluster 2: Gardnerella | Ribonuclease toxin, BrnT, of type II toxin-antitoxin system |
| **K07133** | 40.61540507 | 2.644180263 | 9.00E-19 | 7.76E-18 | Cluster 2: Gardnerella | Domain of unknown function (DUF4143) |

| | | | | | | |
|---|---|---|---|---|---|---|
| **K00286** | 3.312630778 | 2.644077022 | 7.31E-14 | 3.89E-13 | Cluster 2: Gardnerella | Catalyzes the reduction of 1-pyrroline-5-carboxylate (PCA) to L-proline |
| **K03148** | 5.139630007 | 2.642848572 | 1.25E-08 | 4.17E-08 | Cluster 2: Gardnerella | ThiF family |
| **K02429** | 2.369924993 | 2.634501993 | 2.49E-09 | 8.94E-09 | Cluster 2: Gardnerella | Major Facilitator Superfamily |
| **K01079** | 2.204682992 | 2.627137713 | 2.26E-15 | 1.39E-14 | Cluster 2: Gardnerella | haloacid dehalogenase-like hydrolase |
| **K06966** | 2.888695372 | 2.622855569 | 8.14E-11 | 3.38E-10 | Cluster 2: Gardnerella | Belongs to the LOG family |
| **K11358** | 1.10046641 | 2.619175415 | 3.75E-06 | 9.84E-06 | Cluster 2: Gardnerella | PFAM Aminotransferase class I and II |
| **K04079** | 2.218572515 | 2.617315537 | 2.40E-09 | 8.63E-09 | Cluster 2: Gardnerella | Molecular chaperone. Has ATPase activity |
| **K02033** | 27.80905912 | 2.610857078 | 1.41E-35 | 2.59E-33 | Cluster 2: Gardnerella | Binding-protein-dependent transport system inner membrane component |
| **K13730** | 2.131017145 | 2.607064662 | 1.62E-07 | 4.85E-07 | Cluster 2: Gardnerella | domain protein |
| **K01805** | 1.101727072 | 2.59686875 | 1.80E-06 | 4.88E-06 | Cluster 2: Gardnerella | Belongs to the xylose isomerase family |
| **K01823** | 16.99080983 | 2.584133023 | 3.64E-34 | 5.44E-32 | Cluster 2: Gardnerella | Involved in the biosynthesis of isoprenoids. Catalyzes the 1,3-allylic rearrangement of the homoallylic substrate isopentenyl (IPP) to its allylic isomer, dimethylallyl diphosphate (DMAPP) |
| **K02031** | 50.29578166 | 2.583268149 | 7.73E-37 | 2.60E-34 | Cluster 2: Gardnerella | Belongs to the ABC transporter superfamily |

| | | | | | | |
|---|---|---|---|---|---|---|
| **K20074** | 14.31334591 | 2.580664034 | 8.69E-37 | 2.60E-34 | Cluster 2: Gardnerella | Sigma factor PP2C-like phosphatases |
| **K03475** | 21.56736049 | 2.576706782 | 1.90E-15 | 1.17E-14 | Cluster 2: Gardnerella | PTS system sugar-specific permease component |
| **K14441** | 1.620350144 | 2.570240497 | 3.04E-09 | 1.08E-08 | Cluster 2: Gardnerella | Catalyzes the methylthiolation of an aspartic acid residue of ribosomal protein S12 |
| **K04488** | 3.091884693 | 2.564060952 | 5.79E-23 | 8.92E-22 | Cluster 2: Gardnerella | SUF system FeS assembly protein, NifU family |
| **K12990** | 0.962348329 | 2.559697977 | 8.58E-06 | 2.17E-05 | Cluster 2: Gardnerella | Glycosyl transferase family 2 |
| **K00655** | 10.52381586 | 2.554051852 | 3.94E-33 | 5.23E-31 | Cluster 2: Gardnerella | Phosphate acyltransferases |
| **K01933** | 5.35067005 | 2.541753981 | 9.96E-12 | 4.51E-11 | Cluster 2: Gardnerella | Phosphoribosylformylglycinamidine cyclo-ligase |
| **K06972** | 4.008626212 | 2.538213452 | 2.24E-09 | 8.08E-09 | Cluster 2: Gardnerella | Peptidase M16 inactive domain |
| **K17067** | 1.246741962 | 2.53613375 | 1.20E-05 | 2.98E-05 | Cluster 2: Gardnerella | Iron-containing alcohol dehydrogenase |
| **K06726** | 1.091786303 | 2.530370823 | 7.74E-08 | 2.40E-07 | Cluster 2: Gardnerella | RbsD / FucU transport protein family |
| **K01191** | 9.343091336 | 2.522616255 | 9.13E-07 | 2.53E-06 | Cluster 2: Gardnerella | Alpha mannosidase, middle domain |
| **K18334** | 1.691734122 | 2.519882212 | 6.60E-05 | 0.00015408 | Cluster 2: Gardnerella | Enolase C-terminal domain-like |

| | | | | | | |
|---|---|---|---|---|---|---|
| **K16214** | 0.827665719 | 2.517915881 | 0.0014734 | 0.00295174 | Cluster 2: Gardnerella | Zeta toxin |
| **K09693** | 1.818876535 | 2.508293769 | 1.18E-07 | 3.60E-07 | Cluster 2: Gardnerella | ABC transporter |
| **K06969** | 1.289671186 | 2.505335099 | 5.35E-08 | 1.68E-07 | Cluster 2: Gardnerella | Met-10+ like-protein |
| **K07137** | 2.243792533 | 2.487479234 | 5.68E-08 | 1.78E-07 | Cluster 2: Gardnerella | FAD dependent oxidoreductase |
| **K17330** | 0.942449386 | 2.467062482 | 1.40E-05 | 3.45E-05 | Cluster 2: Gardnerella | Binding-protein-dependent transport system inner membrane component |
| **K00133** | 10.19229904 | 2.463395614 | 1.39E-21 | 1.73E-20 | Cluster 2: Gardnerella | Catalyzes the NADPH-dependent formation of L-aspartate- semialdehyde (L-ASA) by the reductive dephosphorylation of L- aspartyl-4-phosphate |
| **K00048** | 1.393416847 | 2.453464267 | 3.24E-07 | 9.48E-07 | Cluster 2: Gardnerella | Iron-containing alcohol dehydrogenase |
| **K07451** | 1.609052417 | 2.445766563 | 2.30E-07 | 6.83E-07 | Cluster 2: Gardnerella | HNH endonuclease |
| **K07067** | 0.907680723 | 2.432008742 | 1.08E-05 | 2.69E-05 | Cluster 2: Gardnerella | DNA integrity scanning protein DisA |
| **K03743** | 2.626653742 | 2.429641867 | 3.65E-10 | 1.41E-09 | Cluster 2: Gardnerella | Belongs to the CinA family |
| **K16925** | 8.955130543 | 2.411371163 | 4.03E-25 | 9.45E-24 | Cluster 2: Gardnerella | ABC-type cobalt transport system, permease component |
| **K08600** | 1.079537853 | 2.410325981 | 4.16E-06 | 1.09E-05 | Cluster 2: Gardnerella | sortase, SrtB family |

| | | | | | | |
|---|---|---|---|---|---|---|
| **K07665** | 0.429333094 | 2.389420788 | 0.001146925 | 0.002332895 | Cluster 2: Gardnerella | Response regulator receiver domain |
| **K03406** | 0.97505661 | 2.388152005 | 0.000154495 | 0.000343802 | Cluster 2: Gardnerella | methyl-accepting chemotaxis protein |
| **K00997** | 4.446878025 | 2.383037192 | 1.04E-27 | 4.19E-26 | Cluster 2: Gardnerella | Transfers the 4'-phosphopantetheine moiety from coenzyme A to a Ser of acyl-carrier-protein |
| **K00694** | 0.984963966 | 2.377584564 | 0.012882072 | 0.02294199 | Cluster 2: Gardnerella | transferase activity, transferring glycosyl groups |
| **K07040** | 6.038002221 | 2.377342852 | 2.72E-26 | 7.92E-25 | Cluster 2: Gardnerella | Uncharacterized ACR, COG1399 |
| **K01804** | 1.103085903 | 2.376010526 | 9.95E-05 | 0.000227692 | Cluster 2: Gardnerella | Catalyzes the conversion of L-arabinose to L-ribulose |
| **K00003** | 11.1840673 | 2.370015121 | 1.22E-16 | 8.28E-16 | Cluster 2: Gardnerella | Homoserine dehydrogenase |
| **K02477** | 0.418286465 | 2.368332567 | 0.000209842 | 0.00046266 | Cluster 2: Gardnerella | Response regulator receiver domain |
| **K00786** | 0.688796853 | 2.367595702 | 0.005913081 | 0.011015015 | Cluster 2: Gardnerella | transferase activity, transferring glycosyl groups |
| **K00872** | 7.500661595 | 2.351732244 | 2.87E-17 | 2.07E-16 | Cluster 2: Gardnerella | Catalyzes the ATP-dependent phosphorylation of L-homoserine to L-homoserine phosphate |
| **K03320** | 13.89431065 | 2.349258307 | 3.80E-14 | 2.08E-13 | Cluster 2: Gardnerella | Ammonium Transporter Family |
| **K00266** | 2.803639158 | 2.345527069 | 2.85E-10 | 1.11E-09 | Cluster 2: Gardnerella | glutamate synthase (NADPH), homotetrameric |

| | | | | | | |
|---|---|---|---|---|---|---|
| **K03602** | 1.332911472 | 2.3413458 | 1.32E-11 | 5.83E-11 | Cluster 2: Gardnerella | Bidirectionally degrades single-stranded DNA into large acid-insoluble oligonucleotides, which are then degraded further into small acid-soluble oligonucleotides |
| **K06218** | 0.730899245 | 2.340869598 | 0.000126724 | 0.000285458 | Cluster 2: Gardnerella | nuclease activity |
| **K13953** | 2.076681312 | 2.339435019 | 1.92E-08 | 6.27E-08 | Cluster 2: Gardnerella | Alcohol dehydrogenase GroES-like domain |
| **K07481** | 0.674544615 | 2.338124049 | 0.005434988 | 0.010211967 | Cluster 2: Gardnerella | COG3039 Transposase and inactivated derivatives IS5 family |
| **K02939** | 5.81765513 | 2.337334855 | 3.21E-27 | 1.08E-25 | Cluster 2: Gardnerella | Binds to the 23S rRNA |
| **K02109** | 6.1754376 | 2.336369527 | 5.65E-24 | 1.05E-22 | Cluster 2: Gardnerella | Component of the F(0) channel, it forms part of the peripheral stalk, linking F(1) to F(0) |
| **K01803** | 14.96583982 | 2.331796343 | 2.28E-29 | 1.43E-27 | Cluster 2: Gardnerella | Phosphoglycerate kinase |
| **K03704** | 1.534879403 | 2.325836111 | 6.00E-20 | 6.00E-19 | Cluster 2: Gardnerella | 'Cold-shock' DNA-binding domain |
| **K19123** | 10.01400295 | 2.325231469 | 7.16E-07 | 2.01E-06 | Cluster 2: Gardnerella | DEAD-like helicases superfamily |
| **K00941** | 12.01880446 | 2.316487609 | 2.59E-25 | 6.26E-24 | Cluster 2: Gardnerella | Phosphomethylpyrimidine kinase |
| **K08302** | 0.649894845 | 2.29969737 | 6.98E-05 | 0.000162515 | Cluster 2: Gardnerella | ketose-bisphosphate aldolase |
| **K07315** | 1.404860936 | 2.293425555 | 5.74E-07 | 1.63E-06 | Cluster 2: Gardnerella | Sigma factor PP2C-like phosphatases |

| | | | | | | |
|---|---|---|---|---|---|---|
| **K20534** | 3.810084929 | 2.29022703 | 2.82E-10 | 1.10E-09 | Cluster 2: Gardnerella | Glycosyl transferase family 2 |
| **K09157** | 16.1755396 | 2.244211621 | 1.82E-10 | 7.24E-10 | Cluster 2: Gardnerella | UPF0210 protein |
| **K00761** | 7.743317281 | 2.242779356 | 7.38E-24 | 1.33E-22 | Cluster 2: Gardnerella | Catalyzes the conversion of uracil and 5-phospho-alpha- D-ribose 1-diphosphate (PRPP) to UMP and diphosphate |
| **K04477** | 0.529844248 | 2.241741321 | 1.09E-05 | 2.71E-05 | Cluster 2: Gardnerella | PHP domain protein |
| **K18303** | 0.507664189 | 2.23745123 | 0.005785805 | 0.010786329 | Cluster 2: Gardnerella | Belongs to the resistance-nodulation-cell division (RND) (TC 2.A.6) family |
| **K08884** | 48.49196453 | 2.237385093 | 1.19E-21 | 1.50E-20 | Cluster 2: Gardnerella | Protein tyrosine kinase |
| **K12111** | 1.128475055 | 2.235195396 | 0.002070242 | 0.004082407 | Cluster 2: Gardnerella | Psort location Cytoplasmic, score 8.87 |
| **K06409** | 0.636836513 | 2.227364034 | 0.002085054 | 0.004108226 | Cluster 2: Gardnerella | Stage V sporulation protein B |
| **K07082** | 10.72488957 | 2.225350924 | 1.09E-11 | 4.93E-11 | Cluster 2: Gardnerella | Functions as a peptidoglycan terminase that cleaves nascent peptidoglycan strands endolytically to terminate their elongation |
| **K02034** | 36.35517185 | 2.222917057 | 7.12E-26 | 1.91E-24 | Cluster 2: Gardnerella | Binding-protein-dependent transport system inner membrane component |
| **K00869** | 9.453403116 | 2.218738395 | 1.17E-20 | 1.29E-19 | Cluster 2: Gardnerella | GHMP kinases C terminal |
| **K00845** | 7.829461298 | 2.21826766 | 2.58E-11 | 1.12E-10 | Cluster 2: Gardnerella | ROK family |

| | | | | | | |
|---|---|---|---|---|---|---|
| **K04759** | 4.264653755 | 2.211814623 | 2.93E-08 | 9.37E-08 | Cluster 2: Gardnerella | Ferrous iron transport protein B |
| **K03711** | 2.17295775 | 2.20811226 | 5.38E-09 | 1.86E-08 | Cluster 2: Gardnerella | Belongs to the Fur family |
| **K05515** | 2.626691814 | 2.20191545 | 1.04E-08 | 3.52E-08 | Cluster 2: Gardnerella | Penicillin-binding protein 2 |
| **K18707** | 1.693078188 | 2.196823668 | 9.15E-07 | 2.54E-06 | Cluster 2: Gardnerella | tRNA methylthiotransferase YqeV |
| **K00863** | 2.009416747 | 2.195184934 | 2.66E-06 | 7.10E-06 | Cluster 2: Gardnerella | Dihydroxyacetone kinase |
| **K03326** | 1.19387468 | 2.194250364 | 6.63E-05 | 0.000154824 | Cluster 2: Gardnerella | Transporter, anaerobic C4-dicarboxylate uptake C (DcuC) family |
| **K21029** | 4.636761149 | 2.191171472 | 6.23E-08 | 1.95E-07 | Cluster 2: Gardnerella | ThiF family |
| **K01595** | 28.47315988 | 2.186297215 | 5.92E-12 | 2.71E-11 | Cluster 2: Gardnerella | Forms oxaloacetate, a four-carbon dicarboxylic acid source for the tricarboxylic acid cycle |
| **K10190** | 1.094282964 | 2.185303716 | 0.001347181 | 0.002717098 | Cluster 2: Gardnerella | Binding-protein-dependent transport system inner membrane component |
| **K03699** | 17.27012031 | 2.180063071 | 6.89E-23 | 1.06E-21 | Cluster 2: Gardnerella | Transporter associated domain |
| **K06962** | 0.922438953 | 2.176664673 | 1.32E-06 | 3.63E-06 | Cluster 2: Gardnerella | YacP-like NYN domain |
| **K19302** | 6.841919354 | 2.164577047 | 2.04E-19 | 1.87E-18 | Cluster 2: Gardnerella | Psort location CytoplasmicMembrane, score |

| | | | | | | |
|---|---|---|---|---|---|---|
| **K02005** | 2.154383596 | 2.148668177 | 2.08E-06 | 5.60E-06 | Cluster 2: Gardnerella | Belongs to the membrane fusion protein (MFP) (TC 8.A.1) family |
| **K06191** | 0.950846124 | 2.14797558 | 6.93E-07 | 1.95E-06 | Cluster 2: Gardnerella | Glutaredoxin |
| **K16783** | 0.474330127 | 2.134464194 | 0.000185579 | 0.000410299 | Cluster 2: Gardnerella | transmembrane transporter activity |
| **K16013** | 17.89708159 | 2.132991587 | 9.19E-16 | 5.81E-15 | Cluster 2: Gardnerella | ABC transporter transmembrane region |
| **K11031** | 15.54466043 | 2.129299343 | 4.37E-06 | 1.14E-05 | Cluster 2: Gardnerella | Thiol-activated cytolysin beta sandwich domain |
| **K02406** | 0.58490806 | 2.119592956 | 0.005747808 | 0.010732234 | Cluster 2: Gardnerella | Flagellin is the subunit protein which polymerizes to form the filaments of bacterial flagella |
| **K01733** | 12.52474744 | 2.116679962 | 8.42E-14 | 4.45E-13 | Cluster 2: Gardnerella | Threonine synthase N terminus |
| **K21572** | 1.219606145 | 2.113072096 | 0.001744621 | 0.003472103 | Cluster 2: Gardnerella | SusD family |
| **K03101** | 3.108900889 | 2.110127709 | 5.06E-19 | 4.48E-18 | Cluster 2: Gardnerella | This protein specifically catalyzes the removal of signal peptides from prolipoproteins |
| **K19165** | 0.481169979 | 2.109682232 | 5.29E-05 | 0.000124068 | Cluster 2: Gardnerella | Antitoxin component of a toxin-antitoxin (TA) module |
| **K00284** | 1.756199378 | 2.10594262 | 0.000287321 | 0.000623703 | Cluster 2: Gardnerella | Glutamate synthase |
| **K04078** | 2.508029019 | 2.09175984 | 8.97E-18 | 6.90E-17 | Cluster 2: Gardnerella | Binds to Cpn60 in the presence of Mg-ATP and suppresses the ATPase activity of the latter |

| | | | | | | |
|---|---|---|---|---|---|---|
| **K02431** | 0.491558756 | 2.091550671 | 0.003194738 | 0.006142737 | Cluster 2: Gardnerella | RbsD / FucU transport protein family |
| **K02968** | 1.533079521 | 2.086819358 | 5.24E-12 | 2.42E-11 | Cluster 2: Gardnerella | Binds directly to 16S ribosomal RNA |
| **K03979** | 17.68520341 | 2.086643941 | 8.23E-24 | 1.47E-22 | Cluster 2: Gardnerella | An essential GTPase which binds GTP, GDP and possibly (p)ppGpp with moderate affinity, with high nucleotide exchange rates and a fairly low GTP hydrolysis rate. Plays a role in control of the cell cycle, stress response, ribosome biogenesis and in those bacteria that undergo differentiation, in morphogenesis control |
| **K12065** | 0.402459792 | 2.08544052 | 0.025738483 | 0.043259475 | Cluster 2: Gardnerella | transfer protein TraB |
| **K01358** | 12.52239759 | 2.078252594 | 2.83E-22 | 3.82E-21 | Cluster 2: Gardnerella | Cleaves peptides in various proteins in a process that requires ATP hydrolysis. Has a chymotrypsin-like activity. Plays a major role in the degradation of misfolded proteins |
| **K02959** | 2.346362808 | 2.072121875 | 6.83E-17 | 4.74E-16 | Cluster 2: Gardnerella | Belongs to the bacterial ribosomal protein bS16 family |
| **K11693** | 1.38669941 | 2.069311145 | 1.48E-05 | 3.64E-05 | Cluster 2: Gardnerella | Methicillin resistance protein |
| **K02654** | 1.007939282 | 2.068682285 | 1.94E-08 | 6.31E-08 | Cluster 2: Gardnerella | Type IV leader peptidase family |
| **K01915** | 30.31330247 | 2.063362417 | 1.47E-23 | 2.49E-22 | Cluster 2: Gardnerella | glutamine synthetase |
| **K03429** | 0.627067118 | 2.05917275 | 0.000636671 | 0.001326628 | Cluster 2: Gardnerella | Monogalactosyldiacylglycerol synthase, C-terminal domain protein |

| | | | | | | |
|---|---|---|---|---|---|---|
| **K05346** | 0.980393647 | 2.051657957 | 0.000125408 | 0.000282759 | Cluster 2: Gardnerella | Putative sugar-binding domain |
| **K09800** | 1.332921892 | 2.049078925 | 0.000365797 | 0.000783383 | Cluster 2: Gardnerella | TamB, inner membrane protein subunit of TAM complex |
| **K01591** | 13.12225206 | 2.044821858 | 1.24E-21 | 1.55E-20 | Cluster 2: Gardnerella | Belongs to the OMP decarboxylase family. Type 2 subfamily |
| **K07568** | 1.718807722 | 2.042038546 | 6.91E-07 | 1.94E-06 | Cluster 2: Gardnerella | Transfers and isomerizes the ribose moiety from AdoMet to the 7-aminomethyl group of 7-deazaguanine (preQ1-tRNA) to give epoxyqueuosine (oQ-tRNA) |
| **K12684** | 0.904111702 | 2.040912892 | 0.008224998 | 0.015063406 | Cluster 2: Gardnerella | Type V secretory pathway, adhesin AidA |
| **K02996** | 3.834915614 | 2.040073064 | 5.46E-18 | 4.28E-17 | Cluster 2: Gardnerella | Belongs to the universal ribosomal protein uS9 family |
| **K06919** | 12.96469031 | 2.038316871 | 4.67E-06 | 1.21E-05 | Cluster 2: Gardnerella | Phage plasmid primase, P4 family |
| **K17331** | 0.445129936 | 2.038177872 | 0.006758432 | 0.012511738 | Cluster 2: Gardnerella | Binding-protein-dependent transport system inner membrane component |
| **K01790** | 3.091947725 | 2.029821575 | 1.43E-06 | 3.91E-06 | Cluster 2: Gardnerella | Catalyzes the reduction of dTDP-6-deoxy-L-lyxo-4-hexulose to yield dTDP-L-rhamnose |
| **K01210** | 0.959605606 | 2.025138349 | 0.004888632 | 0.009228934 | Cluster 2: Gardnerella | Belongs to the glycosyl hydrolase 5 (cellulase A) family |
| **K03496** | 22.69307466 | 2.021320803 | 1.53E-22 | 2.16E-21 | Cluster 2: Gardnerella | CobQ CobB MinD ParA nucleotide binding domain protein |
| **K02821** | 1.986075599 | 2.01444134 | 5.69E-09 | 1.96E-08 | Cluster 2: Gardnerella | Phosphoenolpyruvate-dependent sugar phosphotransferase system, EIIA 2 |

| | | | | | | |
|---|---|---|---|---|---|---|
| **K03685** | 8.467496892 | 2.011437352 | 1.84E-23 | 3.09E-22 | Cluster 2: Gardnerella | Digests double-stranded RNA. Involved in the processing of primary rRNA transcript to yield the immediate precursors to the large and small rRNAs (23S and 16S). Processes some mRNAs, and tRNAs when they are encoded in the rRNA operon. Processes pre- crRNA and tracrRNA of type II CRISPR loci if present in the organism |
| **K03299** | 2.865674807 | 2.000864791 | 7.27E-07 | 2.03E-06 | Cluster 2: Gardnerella | GntP family permease |
| **K02026** | 13.46012779 | 1.999671962 | 5.52E-20 | 5.57E-19 | Cluster 2: Gardnerella | ABC transporter permease |
| **K00549** | 18.09644449 | 1.996340044 | 4.41E-11 | 1.87E-10 | Cluster 2: Gardnerella | Catalyzes the transfer of a methyl group from 5-methyltetrahydrofolate to homocysteine resulting in methionine formation |
| **K00020** | 1.349134738 | 1.995617435 | 2.06E-05 | 5.04E-05 | Cluster 2: Gardnerella | NAD binding domain |
| **K07318** | 0.87892227 | 1.992932326 | 0.007454167 | 0.013704199 | Cluster 2: Gardnerella | D12 class N6 adenine-specific DNA methyltransferase |
| **K07704** | 1.652280653 | 1.989706991 | 0.001232802 | 0.002496946 | Cluster 2: Gardnerella | histidine kinase DNA gyrase B |
| **K06714** | 0.352217439 | 1.989316401 | 0.025818721 | 0.043302978 | Cluster 2: Gardnerella | PFAM sigma-54 factor interaction domain-containing protein, Propionate catabolism activator domain-containing protein |
| **K07277** | 0.915557342 | 1.98481372 | 0.000576156 | 0.001206846 | Cluster 2: Gardnerella | Outer membrane protein, OMP85 family |

| | | | | | | |
|---|---|---|---|---|---|---|
| **K00973** | 2.591798726 | 1.971987936 | 5.57E-07 | 1.59E-06 | Cluster 2: Gardnerella | Catalyzes the formation of dTDP-glucose, from dTTP and glucose 1-phosphate, as well as its pyrophosphorolysis |
| **K04656** | 1.290758054 | 1.970857261 | 0.000569828 | 0.001195687 | Cluster 2: Gardnerella | Belongs to the carbamoyltransferase HypF family |
| **K02279** | 0.44849276 | 1.968491369 | 0.002587113 | 0.005035179 | Cluster 2: Gardnerella | Flp pilus assembly protein CpaB |
| **K17624** | 0.552697915 | 1.968266275 | 0.025587521 | 0.043066321 | Cluster 2: Gardnerella | Glycosyl hydrolase 101 beta sandwich domain |
| **K16923** | 0.810568177 | 1.965501865 | 0.000125061 | 0.000282242 | Cluster 2: Gardnerella | ECF transporter, substrate-specific component |
| **K02011** | 2.106945028 | 1.963829737 | 2.90E-05 | 6.99E-05 | Cluster 2: Gardnerella | Binding-protein-dependent transport system inner membrane component |
| **K03588** | 23.82820679 | 1.956083469 | 9.07E-24 | 1.57E-22 | Cluster 2: Gardnerella | Belongs to the SEDS family |
| **K03216** | 7.408277078 | 1.953587462 | 6.76E-19 | 5.89E-18 | Cluster 2: Gardnerella | Belongs to the class IV-like SAM-binding methyltransferase superfamily. RNA methyltransferase TrmH family. TrmL subfamily |
| **K15581** | 6.175219888 | 1.946133236 | 3.11E-10 | 1.21E-09 | Cluster 2: Gardnerella | Binding-protein-dependent transport system inner membrane component |
| **K02517** | 0.735311761 | 1.944464229 | 0.003517817 | 0.006731451 | Cluster 2: Gardnerella | Lipid A biosynthesis |
| **K07006** | 2.561391003 | 1.943442572 | 4.47E-14 | 2.43E-13 | Cluster 2: Gardnerella | Pfam:Pyridox_oxidase |
| **K02563** | 17.49768545 | 1.942466182 | 1.89E-21 | 2.33E-20 | Cluster 2: Gardnerella | Cell wall formation. Catalyzes the transfer of a GlcNAc subunit on undecaprenyl-pyrophosphoryl- |

| | | | | | | |
|---|---|---|---|---|---|---|
| | | | | | | MurNAc-pentapeptide (lipid intermediate I) to form undecaprenyl-pyrophosphoryl-MurNAc-(pentapeptide)GlcNAc (lipid intermediate II) |
| **K12952** | 19.42200178 | 1.941355207 | 3.16E-14 | 1.74E-13 | Cluster 2: Gardnerella | E1-E2 ATPase |
| **K03470** | 6.929566798 | 1.939463758 | 3.03E-19 | 2.73E-18 | Cluster 2: Gardnerella | Endonuclease that specifically degrades the RNA of RNA- DNA hybrids |
| **K14534** | 0.416764964 | 1.93887613 | 0.025095532 | 0.042327678 | Cluster 2: Gardnerella | 4-hydroxyphenylacetate 3-hydroxylase C terminal |
| **K01652** | 15.24268266 | 1.938009073 | 4.10E-11 | 1.75E-10 | Cluster 2: Gardnerella | Thiamine pyrophosphate enzyme, central domain |
| **K08303** | 5.072422534 | 1.931611076 | 1.99E-06 | 5.36E-06 | Cluster 2: Gardnerella | Peptidase, U32 family |
| **K02935** | 3.151979983 | 1.928414314 | 2.12E-13 | 1.09E-12 | Cluster 2: Gardnerella | Forms part of the ribosomal stalk which helps the ribosome interact with GTP-bound translation factors. Is thus essential for accurate translation |
| **K00760** | 11.51771872 | 1.927695002 | 6.34E-22 | 8.23E-21 | Cluster 2: Gardnerella | Ligates lysine onto the cytidine present at position 34 of the AUA codon-specific tRNA(Ile) that contains the anticodon CAU, in an ATP-dependent manner. Cytidine is converted to lysidine, thus changing the amino acid specificity of the tRNA from methionine to isoleucine |
| **K10545** | 1.009297321 | 1.92342716 | 0.021489839 | 0.036607781 | Cluster 2: Gardnerella | Part of the ABC transporter complex RbsABC involved in ribose import. Responsible for energy coupling to the transport system |

| | | | | | | |
|---|---|---|---|---|---|---|
| **K00954** | 4.48119289 | 1.911861142 | 2.50E-16 | 1.65E-15 | Cluster 2: Gardnerella | Reversibly transfers an adenylyl group from ATP to 4'- phosphopantetheine, yielding dephospho-CoA (dPCoA) and pyrophosphate |
| **K18220** | 5.497031967 | 1.908801169 | 5.90E-05 | 0.000138188 | Cluster 2: Gardnerella | elongation factor G |
| **K03686** | 18.72852392 | 1.892855254 | 2.23E-23 | 3.70E-22 | Cluster 2: Gardnerella | ATP binding to DnaK triggers the release of the substrate protein, thus completing the reaction cycle. Several rounds of ATP-dependent interactions between DnaJ, DnaK and GrpE are required for fully efficient folding. Also involved, together with DnaK and GrpE, in the DNA replication of plasmids through activation of initiation proteins |
| **K01533** | 27.67032032 | 1.881576006 | 7.19E-08 | 2.24E-07 | Cluster 2: Gardnerella | E1-E2 ATPase |
| **K02499** | 0.872355643 | 1.880816711 | 0.000574951 | 0.00120538 | Cluster 2: Gardnerella | MazG family |
| **K22397** | 0.892053058 | 1.877767386 | 0.006691582 | 0.012397581 | Cluster 2: Gardnerella | Dihydrodipicolinate synthetase family |
| **K03625** | 2.616775497 | 1.87576162 | 8.68E-18 | 6.69E-17 | Cluster 2: Gardnerella | Involved in transcription antitermination. Required for transcription of ribosomal RNA (rRNA) genes. Binds specifically to the boxA antiterminator sequence of the ribosomal RNA (rrn) operons |
| **K11753** | 13.03280596 | 1.873476637 | 4.83E-21 | 5.66E-20 | Cluster 2: Gardnerella | riboflavin kinase |
| **K00919** | 2.399915991 | 1.872630681 | 2.51E-06 | 6.71E-06 | Cluster 2: Gardnerella | Catalyzes the phosphorylation of the position 2 hydroxy group of 4-diphosphocytidyl-2C-methyl-D-erythritol |

| | | | | | | |
|---|---|---|---|---|---|---|
| **K03687** | 3.895761816 | 1.868896888 | 2.83E-15 | 1.73E-14 | Cluster 2: Gardnerella | Participates actively in the response to hyperosmotic and heat shock by preventing the aggregation of stress-denatured proteins, in association with DnaK and GrpE. It is the nucleotide exchange factor for DnaK and may function as a thermosensor. Unfolded proteins bind initially to DnaJ |
| **K17734** | 0.790416128 | 1.866513661 | 0.005285338 | 0.009954261 | Cluster 2: Gardnerella | Belongs to the peptidase S8 family |
| **K01006** | 3.7894157 | 1.853924577 | 6.50E-06 | 1.67E-05 | Cluster 2: Gardnerella | Belongs to the PEP-utilizing enzyme family |
| **K07095** | 0.697633664 | 1.849954069 | 1.81E-05 | 4.45E-05 | Cluster 2: Gardnerella | Phosphoesterase |
| **K03296** | 1.112706125 | 1.84840708 | 0.000144647 | 0.000323089 | Cluster 2: Gardnerella | Belongs to the resistance-nodulation-cell division (RND) (TC 2.A.6) family |
| **K07046** | 0.425139227 | 1.842738112 | 0.002367507 | 0.004653242 | Cluster 2: Gardnerella | Amidohydrolase |
| **K03073** | 1.047409647 | 1.841773387 | 8.94E-11 | 3.70E-10 | Cluster 2: Gardnerella | Essential subunit of the Sec protein translocation channel SecYEG. Clamps together the 2 halves of SecY. May contact the channel plug during translocation |
| **K01807** | 9.390226782 | 1.833547706 | 2.06E-17 | 1.51E-16 | Cluster 2: Gardnerella | Catalyzes the reversible conversion of ribose-5-phosphate to ribulose 5-phosphate |
| **K00833** | 0.811903774 | 1.831069117 | 0.000364823 | 0.000781996 | Cluster 2: Gardnerella | Catalyzes the transfer of the alpha-amino group from S- adenosyl-L-methionine (SAM) to 7-keto-8-aminopelargonic acid (KAPA) to form 7,8-diaminopelargonic acid (DAPA). It is the only |

| | | | | | | |
|---|---|---|---|---|---|---|
| | | | | | | animotransferase known to utilize SAM as an amino donor |
| **K06206** | 0.460705846 | 1.825991628 | 0.006424815 | 0.011921824 | Cluster 2: Gardnerella | Belongs to the SfsA family |
| **K07738** | 3.520123964 | 1.824768464 | 7.31E-15 | 4.31E-14 | Cluster 2: Gardnerella | Negatively regulates transcription of bacterial ribonucleotide reductase nrd genes and operons by binding to NrdR- boxes |
| **K03168** | 30.40759353 | 1.818824133 | 1.22E-18 | 1.02E-17 | Cluster 2: Gardnerella | Releases the supercoiling and torsional tension of DNA, which is introduced during the DNA replication and transcription, by transiently cleaving and rejoining one strand of the DNA duplex. Introduces a single-strand break via transesterification at a target site in duplex DNA. The scissile phosphodiester is attacked by the catalytic tyrosine of the enzyme, resulting in the formation of a DNA-(5'-phosphotyrosyl)-enzyme intermediate and the expulsion of a 3'-OH DNA strand. The free DNA strand then undergoes passage around the unbroken strand, thus removing DNA supercoils. Finally, in the religation step, the DNA 3'-OH attacks the covalent intermediate to expel the active-site tyrosine and restore the DNA phosphodiester backbone |
| **K02470** | 42.53191828 | 1.813062492 | 1.19E-18 | 1.00E-17 | Cluster 2: Gardnerella | A type II topoisomerase that negatively supercoils closed circular double-stranded (ds) DNA in an ATP-dependent manner to modulate DNA topology and maintain chromosomes in an underwound state. Negative supercoiling favors strand separation, and DNA replication, transcription, recombination and repair, all of which involve strand separation. Also able to catalyze the interconversion of other |

| | | | | | | |
|---|---|---|---|---|---|---|
| | | | | | | topological isomers of dsDNA rings, including catenanes and knotted rings. Type II topoisomerases break and join 2 DNA strands simultaneously in an ATP-dependent manner |
| **K00995** | 8.45360135 | 1.812781099 | 1.80E-17 | 1.33E-16 | Cluster 2: Gardnerella | Belongs to the CDP-alcohol phosphatidyltransferase class-I family |
| **K09691** | 0.903392218 | 1.802059565 | 0.000504553 | 0.001067152 | Cluster 2: Gardnerella | ABC transporter, ATP-binding protein |
| **K16786** | 77.70287332 | 1.799358572 | 1.50E-15 | 9.27E-15 | Cluster 2: Gardnerella | ATPases associated with a variety of cellular activities |
| **K02822** | 0.791003747 | 1.797902108 | 1.94E-06 | 5.25E-06 | Cluster 2: Gardnerella | PTS system, Lactose/Cellobiose specific IIB subunit |
| **K01886** | 0.841657701 | 1.797470411 | 0.002222664 | 0.004372153 | Cluster 2: Gardnerella | Glutaminyl-tRNA synthetase |
| **K02747** | 0.863941608 | 1.791649031 | 0.002862631 | 0.005527286 | Cluster 2: Gardnerella | PTS system mannose fructose sorbose family IID component |
| **K02587** | 0.356852233 | 1.787181207 | 0.019134709 | 0.032900686 | Cluster 2: Gardnerella | nitrogenase, component 1 |
| **K11717** | 15.68849416 | 1.763909203 | 3.72E-18 | 2.96E-17 | Cluster 2: Gardnerella | Catalyzes the removal of elemental sulfur and selenium atoms from L-cysteine, L-cystine, L-selenocysteine, and L- selenocystine to produce L-alanine |
| **K03696** | 50.78985147 | 1.758139314 | 3.70E-16 | 2.39E-15 | Cluster 2: Gardnerella | Part of a stress-induced multi-chaperone system, it is involved in the recovery of the cell from heat-induced damage, in cooperation with DnaK, DnaJ and GrpE |

| | | | | | | |
|---|---|---|---|---|---|---|
| **K12511** | 0.457994959 | 1.754269091 | 0.000898534 | 0.001841948 | Cluster 2: Gardnerella | Type II secretion system |
| **K00703** | 1.707543601 | 1.753331749 | 3.57E-05 | 8.55E-05 | Cluster 2: Gardnerella | Synthesizes alpha-1,4-glucan chains using ADP-glucose |
| **K08217** | 1.5302174 | 1.753256622 | 0.005455931 | 0.010243264 | Cluster 2: Gardnerella | Transmembrane secretion effector |
| **K03072** | 2.832454658 | 1.752699367 | 7.73E-05 | 0.000179108 | Cluster 2: Gardnerella | Part of the Sec protein translocase complex. Interacts with the SecYEG preprotein conducting channel. SecDF uses the proton motive force (PMF) to complete protein translocation after the ATP-dependent function of SecA |
| **K01996** | 0.552620236 | 1.751792396 | 0.000753 | 0.001555461 | Cluster 2: Gardnerella | ABC transporter, ATP-binding protein |
| **K00099** | 1.522673509 | 1.749196063 | 0.000270787 | 0.000588345 | Cluster 2: Gardnerella | Catalyzes the NADP-dependent rearrangement and reduction of 1-deoxy-D-xylulose-5-phosphate (DXP) to 2-C-methyl-D-erythritol 4-phosphate (MEP) |
| **K03975** | 10.47547091 | 1.744908147 | 5.94E-13 | 2.97E-12 | Cluster 2: Gardnerella | SNARE associated Golgi protein |
| **K00639** | 0.884532613 | 1.739726113 | 0.000964177 | 0.00197293 | Cluster 2: Gardnerella | Catalyzes the decarboxylative condensation of pimeloyl- acyl-carrier protein and L-alanine to produce 8-amino-7- oxononanoate (AON), acyl-carrier protein , and carbon dioxide |
| **K22132** | 0.506393811 | 1.730152193 | 0.000295516 | 0.00063975 | Cluster 2: Gardnerella | ThiF family |
| **K03060** | 0.670394501 | 1.714443633 | 3.21E-08 | 1.02E-07 | Cluster 2: Gardnerella | Promotes RNA polymerase assembly. Latches the N- and C- terminal regions of the beta' subunit thereby |

| | | | | | | |
|---|---|---|---|---|---|---|
| | | | | | | facilitating its interaction with the beta and alpha subunits |
| **K02834** | 2.152348308 | 1.712547477 | 1.70E-12 | 8.24E-12 | Cluster 2: Gardnerella | One of several proteins that assist in the late maturation steps of the functional core of the 30S ribosomal subunit. Associates with free 30S ribosomal subunits (but not with 30S subunits that are part of 70S ribosomes or polysomes). Required for efficient processing of 16S rRNA. May interact with the 5'-terminal helix region of 16S rRNA |
| **K03316** | 10.53449855 | 1.709739878 | 4.95E-07 | 1.42E-06 | Cluster 2: Gardnerella | Sodium/hydrogen exchanger family |
| **K21936** | 0.536384986 | 1.708236464 | 0.017071759 | 0.029566307 | Cluster 2: Gardnerella | Radical SAM domain protein |
| **K08384** | 1.011979211 | 1.698856138 | 0.000785444 | 0.001618285 | Cluster 2: Gardnerella | Penicillin-binding protein, transpeptidase domain protein |
| **K07816** | 7.462026346 | 1.694950265 | 4.79E-12 | 2.22E-11 | Cluster 2: Gardnerella | Region found in RelA / SpoT proteins |
| **K03734** | 13.50082905 | 1.692306885 | 1.28E-11 | 5.70E-11 | Cluster 2: Gardnerella | Flavin transferase that catalyzes the transfer of the FMN moiety of FAD and its covalent binding to the hydroxyl group of a threonine residue in a target flavoprotein |
| **K00088** | 25.16219221 | 1.690836986 | 4.18E-20 | 4.28E-19 | Cluster 2: Gardnerella | Catalyzes the conversion of inosine 5'-phosphate (IMP) to xanthosine 5'-phosphate (XMP), the first committed and rate- limiting step in the de novo synthesis of guanine nucleotides, and therefore plays an important role in the regulation of cell growth |

| | | | | | | |
|---|---|---|---|---|---|---|
| **K02902** | 1.363190111 | 1.687827861 | 1.73E-11 | 7.63E-11 | Cluster 2: Gardnerella | Ribosomal L28 family |
| **K05363** | 2.003994782 | 1.680272954 | 0.000334048 | 0.000717963 | Cluster 2: Gardnerella | Methicillin resistance protein |
| **K02860** | 4.927097117 | 1.679284887 | 5.64E-13 | 2.84E-12 | Cluster 2: Gardnerella | An accessory protein needed during the final step in the assembly of 30S ribosomal subunit, possibly for assembly of the head region. Probably interacts with S19. Essential for efficient processing of 16S rRNA. May be needed both before and after RbfA during the maturation of 16S rRNA. It has affinity for free ribosomal 30S subunits but not for 70S ribosomes |
| **K00313** | 0.818512831 | 1.675968577 | 0.000952117 | 0.001949922 | Cluster 2: Gardnerella | FAD dependent oxidoreductase |
| **K02867** | 3.787049444 | 1.671574304 | 3.88E-13 | 1.97E-12 | Cluster 2: Gardnerella | Forms part of the ribosomal stalk which helps the ribosome interact with GTP-bound translation factors |
| **K02004** | 134.9334966 | 1.66982451 | 8.81E-15 | 5.16E-14 | Cluster 2: Gardnerella | FtsX-like permease family |
| **K00928** | 14.31114081 | 1.660853981 | 1.53E-08 | 5.06E-08 | Cluster 2: Gardnerella | Amino acid kinase family |
| **K16787** | 80.85426865 | 1.657835684 | 2.12E-13 | 1.09E-12 | Cluster 2: Gardnerella | ATPases associated with a variety of cellular activities |
| **K11720** | 0.501955329 | 1.654921077 | 0.012819595 | 0.022847749 | Cluster 2: Gardnerella | Permease, YjgP YjgQ family |
| **K01000** | 10.57951621 | 1.654367778 | 6.10E-16 | 3.89E-15 | Cluster 2: Gardnerella | First step of the lipid cycle reactions in the biosynthesis of the cell wall peptidoglycan |

| | | | | | | |
|---|---|---|---|---|---|---|
| **K02884** | 2.213007706 | 1.652980047 | 3.59E-12 | 1.69E-11 | Cluster 2: Gardnerella | This protein is located at the 30S-50S ribosomal subunit interface and may play a role in the structure and function of the aminoacyl-tRNA binding site |
| **K07173** | 2.708933374 | 1.652017332 | 2.78E-07 | 8.18E-07 | Cluster 2: Gardnerella | Involved in the synthesis of autoinducer 2 (AI-2) which is secreted by bacteria and is used to communicate both the cell density and the metabolic potential of the environment. The regulation of gene expression in response to changes in cell density is called quorum sensing. Catalyzes the transformation of S-ribosylhomocysteine (RHC) to homocysteine (HC) and 4,5- dihydroxy-2,3-pentadione (DPD) |
| **K02967** | 8.71029572 | 1.648319608 | 2.71E-17 | 1.96E-16 | Cluster 2: Gardnerella | Belongs to the universal ribosomal protein uS2 family |
| **K02601** | 12.86503537 | 1.644756268 | 2.39E-16 | 1.58E-15 | Cluster 2: Gardnerella | Participates in transcription elongation, termination and antitermination |
| **K06182** | 0.486990382 | 1.64121798 | 0.009713871 | 0.017681761 | Cluster 2: Gardnerella | Lytic transglycosylase with a strong preference for naked glycan strands that lack stem peptides |
| **K03589** | 8.04702764 | 1.640707295 | 7.25E-17 | 5.01E-16 | Cluster 2: Gardnerella | Cell division protein FtsQ |
| **K02990** | 2.926259413 | 1.639870355 | 3.67E-12 | 1.71E-11 | Cluster 2: Gardnerella | Binds together with S18 to 16S ribosomal RNA |
| **K02035** | 83.22538079 | 1.639104406 | 2.37E-14 | 1.34E-13 | Cluster 2: Gardnerella | Bacterial extracellular solute-binding proteins, family 5 Middle |
| **K01356** | 8.695647617 | 1.634719408 | 2.26E-16 | 1.50E-15 | Cluster 2: Gardnerella | Represses a number of genes involved in the response to DNA damage (SOS response), including recA and lexA. In the presence of single-stranded DNA, RecA interacts with LexA causing an autocatalytic cleavage which disrupts the DNA-binding part of LexA, leading |

| | | | | | | |
|---|---|---|---|---|---|---|
| | | | | | | to derepression of the SOS regulon and eventually DNA repair |
| **K02435** | 1.428575124 | 1.631622769 | 9.83E-09 | 3.33E-08 | Cluster 2: Gardnerella | Allows the formation of correctly charged Asn-tRNA(Asn) or Gln-tRNA(Gln) through the transamidation of misacylated Asp- tRNA(Asn) or Glu-tRNA(Gln) in organisms which lack either or both of asparaginyl-tRNA or glutaminyl-tRNA synthetases. The reaction takes place in the presence of glutamine and ATP through an activated phospho-Asp-tRNA(Asn) or phospho-Glu-tRNA(Gln) |
| **K01810** | 22.1165603 | 1.627157809 | 1.75E-16 | 1.17E-15 | Cluster 2: Gardnerella | Belongs to the GPI family |
| **K03925** | 4.837551052 | 1.626313031 | 1.31E-13 | 6.84E-13 | Cluster 2: Gardnerella | Belongs to the MraZ family |
| **K01624** | 13.98509961 | 1.620237854 | 1.78E-14 | 1.02E-13 | Cluster 2: Gardnerella | Fructose-bisphosphate aldolase class-II |
| **K08999** | 0.442755711 | 1.612126374 | 0.004870562 | 0.009202089 | Cluster 2: Gardnerella | Bifunctional nuclease |
| **K02986** | 8.028406446 | 1.61196213 | 1.68E-14 | 9.62E-14 | Cluster 2: Gardnerella | One of the primary rRNA binding proteins, it binds directly to 16S rRNA where it nucleates assembly of the body of the 30S subunit |
| **K02469** | 53.5424196 | 1.607289203 | 2.08E-14 | 1.18E-13 | Cluster 2: Gardnerella | A type II topoisomerase that negatively supercoils closed circular double-stranded (ds) DNA in an ATP-dependent manner to modulate DNA topology and maintain chromosomes in an underwound state. Negative supercoiling favors strand separation, and DNA replication, transcription, recombination and repair, all of which involve strand separation. Also able to catalyze the interconversion of other |

| | | | | | | |
|---|---|---|---|---|---|---|
| | | | | | | topological isomers of dsDNA rings, including catenanes and knotted rings. Type II topoisomerases break and join 2 DNA strands simultaneously in an ATP-dependent manner |
| **K06183** | 6.111140174 | 1.604122953 | 1.00E-12 | 4.93E-12 | Cluster 2: Gardnerella | Belongs to the pseudouridine synthase RsuA family |
| **K06925** | 3.002664582 | 1.601475038 | 8.00E-12 | 3.63E-11 | Cluster 2: Gardnerella | Threonylcarbamoyl adenosine biosynthesis protein TsaE |
| **K19002** | 13.92068118 | 1.596836875 | 3.23E-11 | 1.39E-10 | Cluster 2: Gardnerella | PFAM Glycosyl transferase, group 1 |
| **K15633** | 1.087562404 | 1.59354083 | 0.000751501 | 0.001553708 | Cluster 2: Gardnerella | Catalyzes the interconversion of 2-phosphoglycerate and 3-phosphoglycerate |
| **K01811** | 1.611506638 | 1.592255446 | 0.013917328 | 0.024584193 | Cluster 2: Gardnerella | Belongs to the glycosyl hydrolase 31 family |
| **K00554** | 12.02894658 | 1.590455593 | 6.94E-14 | 3.70E-13 | Cluster 2: Gardnerella | Belongs to the RNA methyltransferase TrmD family |
| **K00075** | 13.80826144 | 1.590266469 | 1.17E-15 | 7.28E-15 | Cluster 2: Gardnerella | Cell wall formation |
| **K00652** | 0.696452869 | 1.5901006 | 0.00135429 | 0.002729134 | Cluster 2: Gardnerella | Catalyzes the decarboxylative condensation of pimeloyl- acyl-carrier protein and L-alanine to produce 8-amino-7- oxononanoate (AON), acyl-carrier protein , and carbon dioxide |
| **K04092** | 0.448138214 | 1.587569674 | 0.002154272 | 0.004241111 | Cluster 2: Gardnerella | Chorismate mutase type II |
| **K00917** | 1.041942322 | 1.585830705 | 0.001801171 | 0.003569485 | Cluster 2: Gardnerella | Belongs to the carbohydrate kinase PfkB family |

| | | | | | | |
|---|---|---|---|---|---|---|
| **K12410** | 8.272493793 | 1.585351849 | 5.22E-12 | 2.41E-11 | Cluster 2: Gardnerella | Sir2 family |
| **K03657** | 83.94116222 | 1.57912494 | 2.38E-14 | 1.34E-13 | Cluster 2: Gardnerella | DNA helicase |
| **K03526** | 1.803181665 | 1.579121808 | 0.000170098 | 0.000377118 | Cluster 2: Gardnerella | Converts 2C-methyl-D-erythritol 2,4-cyclodiphosphate (ME-2,4cPP) into 1-hydroxy-2-methyl-2-(E)-butenyl 4-diphosphate |
| **K06381** | 0.51478927 | 1.578511634 | 0.012269282 | 0.021932671 | Cluster 2: Gardnerella | SpoIID LytB domain protein |
| **K01639** | 3.949052987 | 1.57849582 | 0.00043907 | 0.000932779 | Cluster 2: Gardnerella | Dihydrodipicolinate synthetase family |
| **K20903** | 0.47540906 | 1.578214315 | 0.007394823 | 0.013605564 | Cluster 2: Gardnerella | dehydratase |
| **K02988** | 9.837097168 | 1.577916617 | 1.06E-14 | 6.19E-14 | Cluster 2: Gardnerella | Located at the back of the 30S subunit body where it stabilizes the conformation of the head with respect to the body |
| **K00974** | 20.34666255 | 1.577710359 | 3.10E-15 | 1.87E-14 | Cluster 2: Gardnerella | Probable RNA and SrmB- binding site of polymerase A |
| **K05342** | 0.635349395 | 1.577232034 | 0.011189359 | 0.020122323 | Cluster 2: Gardnerella | Glycosyl hydrolase family 65 central catalytic domain |
| **K01924** | 25.22908667 | 1.57675922 | 7.97E-15 | 4.68E-14 | Cluster 2: Gardnerella | Belongs to the MurCDEF family |
| **K07007** | 2.305681244 | 1.571376219 | 0.000292424 | 0.00063363 | Cluster 2: Gardnerella | Flavoprotein family |
| **K03756** | 0.552497813 | 1.570381664 | 0.022211416 | 0.037702617 | Cluster 2: Gardnerella | Putrescine transporter |

| | | | | | | |
|---|---|---|---|---|---|---|
| **K02520** | 7.120645 | 1.569017698 | 4.86E-14 | 2.63E-13 | Cluster 2: Gardnerella | IF-3 binds to the 30S ribosomal subunit and shifts the equilibrum between 70S ribosomes and their 50S and 30S subunits in favor of the free subunits, thus enhancing the availability of 30S subunits on which protein synthesis initiation begins |
| **K01183** | 0.674561414 | 1.568579132 | 0.020782145 | 0.035528846 | Cluster 2: Gardnerella | Hep Hag repeat protein |
| **K01200** | 69.27928886 | 1.568452276 | 9.25E-08 | 2.85E-07 | Cluster 2: Gardnerella | Aamy_C |
| **K06960** | 0.32897159 | 1.568178072 | 0.000700611 | 0.001454788 | Cluster 2: Gardnerella | Belongs to the UPF0109 family |
| **K03110** | 13.28776507 | 1.566827471 | 3.43E-16 | 2.22E-15 | Cluster 2: Gardnerella | Involved in targeting and insertion of nascent membrane proteins into the cytoplasmic membrane. Acts as a receptor for the complex formed by the signal recognition particle (SRP) and the ribosome-nascent chain (RNC) |
| **K13276** | 0.68217373 | 1.566325452 | 0.025440219 | 0.042848571 | Cluster 2: Gardnerella | Belongs to the peptidase S8 family |
| **K03561** | 0.417389276 | 1.565996023 | 0.011797113 | 0.021119924 | Cluster 2: Gardnerella | Transporter MotA TolQ ExbB proton channel family protein |
| **K18979** | 0.67486048 | 1.563442817 | 0.010472813 | 0.018962139 | Cluster 2: Gardnerella | iron-sulfur cluster-binding protein |
| **K02493** | 9.07001921 | 1.55929284 | 1.39E-16 | 9.42E-16 | Cluster 2: Gardnerella | Methylates the class 1 translation termination release factors RF1 PrfA and RF2 PrfB on the glutamine residue of the universally conserved GGQ motif |
| **K00604** | 8.82183263 | 1.558739428 | 1.96E-14 | 1.12E-13 | Cluster 2: Gardnerella | Attaches a formyl group to the free amino group of methionyl-tRNA(fMet). The formyl group appears to |

| | | | | | | |
|---|---|---|---|---|---|---|
| | | | | | | play a dual role in the initiator identity of N-formylmethionyl-tRNA by promoting its recognition by IF2 and preventing the misappropriation of this tRNA by the elongation apparatus |
| **K01462** | 9.31396693 | 1.558680792 | 3.06E-15 | 1.85E-14 | Cluster 2: Gardnerella | Removes the formyl group from the N-terminal Met of newly synthesized proteins. Requires at least a dipeptide for an efficient rate of reaction. N-terminal L-methionine is a prerequisite for activity but the enzyme has broad specificity at other positions |
| **K06941** | 2.656392647 | 1.553882363 | 0.000122015 | 0.000275889 | Cluster 2: Gardnerella | Specifically methylates position 2 of adenine 2503 in 23S rRNA and position 2 of adenine 37 in tRNAs |
| **K04043** | 26.78413125 | 1.548853626 | 1.14E-13 | 6.00E-13 | Cluster 2: Gardnerella | Heat shock 70 kDa protein |
| **K00789** | 13.14222283 | 1.543991939 | 7.05E-16 | 4.47E-15 | Cluster 2: Gardnerella | Catalyzes the formation of S-adenosylmethionine (AdoMet) from methionine and ATP. The overall synthetic reaction is composed of two sequential steps, AdoMet formation and the subsequent tripolyphosphate hydrolysis which occurs prior to release of AdoMet from the enzyme |
| **K00174** | 0.795164579 | 1.543174892 | 0.002537369 | 0.00494239 | Cluster 2: Gardnerella | ferredoxin oxidoreductase |
| **K03074** | 1.119274185 | 1.5424602 | 0.000263559 | 0.000574208 | Cluster 2: Gardnerella | Part of the Sec protein translocase complex. Interacts with the SecYEG preprotein conducting channel. SecDF uses the proton motive force (PMF) to complete protein translocation after the ATP-dependent function of SecA |
| **K02337** | 38.82670224 | 1.536778136 | 5.90E-14 | 3.18E-13 | Cluster 2: Gardnerella | DNA polymerase III alpha subunit |

| | | | | | | |
|---|---|---|---|---|---|---|
| **K03117** | 0.361892815 | 1.536382783 | 0.003172153 | 0.006104224 | Cluster 2: Gardnerella | Twin arginine-targeting protein translocase, TatA E family |
| **K17686** | 28.37758314 | 1.53125626 | 1.32E-11 | 5.83E-11 | Cluster 2: Gardnerella | E1-E2 ATPase |
| **K00759** | 3.85164391 | 1.529686686 | 1.17E-12 | 5.75E-12 | Cluster 2: Gardnerella | Catalyzes a salvage reaction resulting in the formation of AMP, that is energically less costly than de novo synthesis |
| **K03310** | 5.16045543 | 1.526579918 | 0.000146596 | 0.000327138 | Cluster 2: Gardnerella | amino acid carrier protein |
| **K03118** | 1.152299638 | 1.525194115 | 0.00017538 | 0.000388469 | Cluster 2: Gardnerella | Part of the twin-arginine translocation (Tat) system that transports large folded proteins containing a characteristic twin-arginine motif in their signal peptide across membranes. Together with TatB, TatC is part of a receptor directly interacting with Tat signal peptides |
| **K06213** | 2.667817708 | 1.524812655 | 0.000613522 | 0.001281746 | Cluster 2: Gardnerella | Magnesium transporter |
| **K00975** | 2.430306193 | 1.521607008 | 8.49E-05 | 0.000196161 | Cluster 2: Gardnerella | Catalyzes the synthesis of ADP-glucose, a sugar donor used in elongation reactions on alpha-glucans |
| **K02600** | 18.53395943 | 1.519548904 | 4.76E-14 | 2.58E-13 | Cluster 2: Gardnerella | Participates in both transcription termination and antitermination |
| **K01906** | 0.55954401 | 1.516693268 | 0.005690504 | 0.010633546 | Cluster 2: Gardnerella | Catalyzes the decarboxylative condensation of pimeloyl- acyl-carrier protein and L-alanine to produce 8-amino-7- oxononanoate (AON), acyl-carrier protein , and carbon dioxide |
| **K01714** | 4.884524318 | 1.516028472 | 2.71E-06 | 7.21E-06 | Cluster 2: Gardnerella | Dihydrodipicolinate synthetase family |

| | | | | | | |
|---|---|---|---|---|---|---|
| **K06168** | 2.837464627 | 1.515053187 | 0.000122757 | 0.000277305 | Cluster 2: Gardnerella | Catalyzes the methylthiolation of N6-(dimethylallyl)adenosine (i(6)A), leading to the formation of 2- methylthio-N6-(dimethylallyl)adenosine (ms(2)i(6)A) at position 37 in tRNAs that read codons beginning with uridine |
| **K01338** | 2.340663726 | 1.510942626 | 0.000510667 | 0.001078175 | Cluster 2: Gardnerella | ATP-dependent serine protease that mediates the selective degradation of mutant and abnormal proteins as well as certain short-lived regulatory proteins. Required for cellular homeostasis and for survival from DNA damage and developmental changes induced by stress. Degrades polypeptides processively to yield small peptide fragments that are 5 to 10 amino acids long. Binds to DNA in a double-stranded, site-specific manner |
| **K02424** | 2.468317371 | 1.50996607 | 0.000338346 | 0.000726547 | Cluster 2: Gardnerella | Bacterial periplasmic substrate-binding proteins |
| **K00852** | 14.97181713 | 1.50664488 | 2.16E-13 | 1.11E-12 | Cluster 2: Gardnerella | Catalyzes the phosphorylation of ribose at O-5 in a reaction requiring ATP and magnesium. The resulting D-ribose-5- phosphate can then be used either for sythesis of nucleotides, histidine, and tryptophan, or as a component of the pentose phosphate pathway |
| **K03439** | 10.23316868 | 1.506417612 | 2.47E-14 | 1.38E-13 | Cluster 2: Gardnerella | Catalyzes the formation of N(7)-methylguanine at position 46 (m7G46) in tRNA |
| **K05808** | 5.854118557 | 1.505015207 | 5.72E-11 | 2.41E-10 | Cluster 2: Gardnerella | Required for dimerization of active 70S ribosomes into 100S ribosomes in stationary phase |
| **K03090** | 0.841998894 | 1.504186784 | 0.00649836 | 0.012048937 | Cluster 2: Gardnerella | RNA polymerase sigma-70 factor, sigma-B F G subfamily |

| | | | | | | |
|---|---|---|---|---|---|---|
| **K01870** | 37.80703671 | 1.498389912 | 1.60E-14 | 9.21E-14 | Cluster 2: Gardnerella | amino acids such as valine, to avoid such errors it has two additional distinct tRNA(Ile)-dependent editing activities. One activity is designated as 'pretransfer' editing and involves the hydrolysis of activated Val-AMP. The other activity is designated 'posttransfer' editing and involves deacylation of mischarged Val-tRNA(Ile) |
| **K01243** | 8.560408869 | 1.493240997 | 7.49E-13 | 3.73E-12 | Cluster 2: Gardnerella | Catalyzes the irreversible cleavage of the glycosidic bond in both 5'-methylthioadenosine (MTA) and S-adenosylhomocysteine (SAH AdoHcy) to adenine and the corresponding thioribose, 5'-methylthioribose and S-ribosylhomocysteine, respectively |
| **K06147** | 146.9518171 | 1.492950241 | 2.89E-12 | 1.39E-11 | Cluster 2: Gardnerella | ABC transporter transmembrane region |
| **K03075** | 0.7388194 | 1.490916883 | 4.21E-07 | 1.22E-06 | Cluster 2: Gardnerella | Preprotein translocase SecG subunit |
| **K02887** | 2.568250975 | 1.489556155 | 1.15E-11 | 5.16E-11 | Cluster 2: Gardnerella | Binds directly to 23S ribosomal RNA and is necessary for the in vitro assembly process of the 50S ribosomal subunit. It is not involved in the protein synthesizing functions of that subunit |
| **K02895** | 0.900149496 | 1.487368068 | 5.26E-07 | 1.51E-06 | Cluster 2: Gardnerella | One of the proteins that surrounds the polypeptide exit tunnel on the outside of the subunit |
| **K01770** | 0.559290332 | 1.486214138 | 0.003390182 | 0.006502838 | Cluster 2: Gardnerella | Involved in the biosynthesis of isopentenyl diphosphate (IPP) and dimethylallyl diphosphate (DMAPP), two major building blocks of isoprenoid compounds. Catalyzes the conversion of 4-diphosphocytidyl-2-C-methyl-D-erythritol 2-phosphate (CDP-ME2P) to 2-C-methyl-D-erythritol |

| | | | | | | |
|---|---|---|---|---|---|---|
| | | | | | | 2,4-cyclodiphosphate (ME-CPP) with a corresponding release of cytidine 5-monophosphate (CMP) |
| **K18918** | 0.594601068 | 1.484208279 | 0.020359521 | 0.034902486 | Cluster 2: Gardnerella | CopG domain protein DNA-binding domain protein |
| **K08313** | 0.342909791 | 1.481689719 | 0.015668075 | 0.027313419 | Cluster 2: Gardnerella | Transaldolase |
| **K02114** | 3.113859076 | 1.481450539 | 3.40E-11 | 1.47E-10 | Cluster 2: Gardnerella | Produces ATP from ADP in the presence of a proton gradient across the membrane |
| **K00854** | 2.133536252 | 1.478298927 | 0.001264207 | 0.002556222 | Cluster 2: Gardnerella | FGGY family of carbohydrate kinases, C-terminal domain |
| **K02077** | 19.23927591 | 1.477038577 | 1.66E-11 | 7.33E-11 | Cluster 2: Gardnerella | Zinc-uptake complex component A periplasmic |
| **K01881** | 22.58687372 | 1.471925011 | 1.58E-13 | 8.19E-13 | Cluster 2: Gardnerella | Catalyzes the attachment of proline to tRNA(Pro) in a two-step reaction proline is first activated by ATP to form Pro- AMP and then transferred to the acceptor end of tRNA(Pro). As ProRS can inadvertently accommodate and process non-cognate amino acids such as alanine and cysteine, to avoid such errors it has two additional distinct editing activities against alanine. One activity is designated as 'pretransfer' editing and involves the tRNA(Pro)-independent hydrolysis of activated Ala-AMP. The other activity is designated 'posttransfer' editing and involves deacylation of mischarged Ala-tRNA(Pro). The misacylated Cys- tRNA(Pro) is not edited by ProRS |
| **K04654** | 0.552310705 | 1.471825826 | 0.005058133 | 0.009533863 | Cluster 2: Gardnerella | hydrogenase expression formation protein HypD |

| | | | | | | |
|---|---|---|---|---|---|---|
| **K03154** | 0.258526533 | 1.470031506 | 0.010060488 | 0.018284841 | Cluster 2: Gardnerella | thiamine diphosphate biosynthetic process |
| **K02358** | 15.87789851 | 1.468425061 | 7.04E-13 | 3.51E-12 | Cluster 2: Gardnerella | This protein promotes the GTP-dependent binding of aminoacyl-tRNA to the A-site of ribosomes during protein biosynthesis |
| **K01998** | 1.221220439 | 1.452488499 | 0.002662833 | 0.00516992 | Cluster 2: Gardnerella | Belongs to the binding-protein-dependent transport system permease family |
| **K01597** | 9.110567773 | 1.449203061 | 1.47E-12 | 7.17E-12 | Cluster 2: Gardnerella | GHMP kinases N terminal domain |
| **K01999** | 1.884052674 | 1.446223174 | 0.000971695 | 0.001986614 | Cluster 2: Gardnerella | Periplasmic binding protein domain |
| **K02835** | 15.66000882 | 1.446153798 | 1.16E-11 | 5.18E-11 | Cluster 2: Gardnerella | Peptide chain release factor 1 directs the termination of translation in response to the peptide chain termination codons UAG and UAA |
| **K07042** | 3.250436538 | 1.44606962 | 9.95E-11 | 4.10E-10 | Cluster 2: Gardnerella | Single strand-specific metallo-endoribonuclease involved in late-stage 70S ribosome quality control and in maturation of the 3' terminus of the 16S rRNA |
| **K02666** | 0.420868186 | 1.445193998 | 0.019423927 | 0.033349989 | Cluster 2: Gardnerella | type II and III secretion system protein |
| **K06199** | 4.746302677 | 1.444514219 | 1.50E-10 | 6.06E-10 | Cluster 2: Gardnerella | Important for reducing fluoride concentration in the cell, thus reducing its toxicity |
| **K17828** | 11.86249495 | 1.444328395 | 3.79E-10 | 1.46E-09 | Cluster 2: Gardnerella | Belongs to the dihydroorotate dehydrogenase family. Type 1 subfamily |
| **K18955** | 0.783882905 | 1.438844309 | 0.000431272 | 0.000917845 | Cluster 2: Gardnerella | Acts as a transcriptional regulator. Probably redox-responsive. The apo- but not holo-form probably binds DNA |

| | | | | | | |
|---|---|---|---|---|---|---|
| **K01619** | 1.274274545 | 1.436857612 | 0.000525045 | 0.001104629 | Cluster 2: Gardnerella | Catalyzes a reversible aldol reaction between acetaldehyde and D-glyceraldehyde 3-phosphate to generate 2-deoxy- D-ribose 5-phosphate |
| **K09936** | 0.621151558 | 1.435881215 | 0.016828269 | 0.0292137 | Cluster 2: Gardnerella | protein conserved in bacteria |
| **K02948** | 2.931176511 | 1.433032945 | 3.77E-08 | 1.19E-07 | Cluster 2: Gardnerella | Located on the platform of the 30S subunit, it bridges several disparate RNA helices of the 16S rRNA. Forms part of the Shine-Dalgarno cleft in the 70S ribosome |
| **K03282** | 2.131115653 | 1.432612054 | 4.24E-10 | 1.63E-09 | Cluster 2: Gardnerella | Channel that opens in response to stretch forces in the membrane lipid bilayer. May participate in the regulation of osmotic pressure changes within the cell |
| **K00560** | 12.44545957 | 1.426200238 | 5.71E-12 | 2.62E-11 | Cluster 2: Gardnerella | Catalyzes the reductive methylation of 2'-deoxyuridine- 5'-monophosphate (dUMP) to 2'-deoxythymidine-5'-monophosphate (dTMP) while utilizing 5,10-methylenetetrahydrofolate (mTHF) as the methyl donor and reductant in the reaction, yielding dihydrofolate (DHF) as a by-product. This enzymatic reaction provides an intracellular de novo source of dTMP, an essential precursor for DNA biosynthesis |
| **K01938** | 21.59749554 | 1.425474668 | 1.19E-10 | 4.87E-10 | Cluster 2: Gardnerella | Formate-tetrahydrofolate ligase |
| **K02913** | 1.028259803 | 1.419374286 | 1.73E-06 | 4.71E-06 | Cluster 2: Gardnerella | Ribosomal protein L33 |
| **K02237** | 4.069903143 | 1.418337334 | 4.71E-10 | 1.80E-09 | Cluster 2: Gardnerella | Helix-hairpin-helix motif |

| | | | | | | |
|---|---|---|---|---|---|---|
| **K02014** | 2.343215789 | 1.417506622 | 0.000807316 | 0.001660487 | Cluster 2: Gardnerella | TonB dependent receptor |
| **K02911** | 1.043958244 | 1.416584614 | 2.02E-05 | 4.96E-05 | Cluster 2: Gardnerella | Belongs to the bacterial ribosomal protein bL32 family |
| **K00067** | 4.008415931 | 1.414637015 | 0.000524978 | 0.001104629 | Cluster 2: Gardnerella | Catalyzes the reduction of dTDP-6-deoxy-L-lyxo-4-hexulose to yield dTDP-L-rhamnose |
| **K02945** | 16.15850248 | 1.411873474 | 5.87E-13 | 2.95E-12 | Cluster 2: Gardnerella | Ribosomal protein S1 |
| **K06200** | 1.148891341 | 1.408212596 | 0.0071077 | 0.013127823 | Cluster 2: Gardnerella | Carbon starvation protein |
| **K10914** | 4.178163267 | 1.407321677 | 6.17E-06 | 1.59E-05 | Cluster 2: Gardnerella | helix_turn_helix, cAMP Regulatory protein |
| **K03977** | 18.65045277 | 1.403216886 | 5.40E-12 | 2.48E-11 | Cluster 2: Gardnerella | GTPase that plays an essential role in the late steps of ribosome biogenesis |
| **K02316** | 20.79663191 | 1.398315179 | 6.94E-12 | 3.16E-11 | Cluster 2: Gardnerella | RNA polymerase that catalyzes the synthesis of short RNA molecules used as primers for DNA polymerase during DNA replication |
| **K01995** | 0.905885824 | 1.397615645 | 0.003746267 | 0.00715142 | Cluster 2: Gardnerella | ABC transporter, ATP-binding protein |
| **K00859** | 5.340732197 | 1.391465005 | 1.72E-10 | 6.89E-10 | Cluster 2: Gardnerella | Catalyzes the phosphorylation of the 3'-hydroxyl group of dephosphocoenzyme A to form coenzyme A |
| **K02863** | 16.61307487 | 1.390843312 | 1.58E-09 | 5.74E-09 | Cluster 2: Gardnerella | Binds directly to 23S rRNA. The L1 stalk is quite mobile in the ribosome, and is involved in E site tRNA release |
| **K04655** | 0.635286616 | 1.388824719 | 0.010933861 | 0.019692485 | Cluster 2: Gardnerella | hydrogenase expression formation protein HypE |

| | | | | | | |
|---|---|---|---|---|---|---|
| **K07166** | 0.825629499 | 1.388775194 | 0.002369648 | 0.004653623 | Cluster 2: Gardnerella | Belongs to the UPF0237 family |
| **K03584** | 6.778486258 | 1.387032868 | 8.44E-11 | 3.50E-10 | Cluster 2: Gardnerella | Involved in DNA repair and RecF pathway recombination |
| **K02483** | 6.437557263 | 1.386750649 | 9.86E-05 | 0.00022584 | Cluster 2: Gardnerella | Response regulator receiver domain protein |
| **K03585** | 0.600303136 | 1.384742229 | 0.025690943 | 0.04320996 | Cluster 2: Gardnerella | Belongs to the membrane fusion protein (MFP) (TC 8.A.1) family |
| **K03536** | 4.167555289 | 1.383221595 | 2.88E-09 | 1.03E-08 | Cluster 2: Gardnerella | RNaseP catalyzes the removal of the 5'-leader sequence from pre-tRNA to produce the mature 5'-terminus. It can also cleave other RNA substrates such as 4.5S RNA. The protein component plays an auxiliary but essential role in vivo by binding to the 5'-leader sequence and broadening the substrate specificity of the ribozyme |
| **K02111** | 22.81802286 | 1.38124237 | 1.46E-09 | 5.34E-09 | Cluster 2: Gardnerella | Produces ATP from ADP in the presence of a proton gradient across the membrane. The alpha chain is a regulatory subunit |
| **K00791** | 13.10403674 | 1.378794242 | 3.63E-12 | 1.70E-11 | Cluster 2: Gardnerella | Catalyzes the transfer of a dimethylallyl group onto the adenine at position 37 in tRNAs that read codons beginning with uridine, leading to the formation of N6-(dimethylallyl)adenosine (i(6)A) |
| **K03546** | 2.373461558 | 1.378078299 | 0.000503395 | 0.001065646 | Cluster 2: Gardnerella | Exonuclease SbcC |
| **K01918** | 1.664162797 | 1.375160401 | 0.000678891 | 0.001410914 | Cluster 2: Gardnerella | Catalyzes the condensation of pantoate with beta-alanine in an ATP-dependent reaction via a pantoyl-adenylate intermediate |

| | | | | | | |
|---|---|---|---|---|---|---|
| **K02653** | 0.743075729 | 1.372632113 | 0.00283578 | 0.005483426 | Cluster 2: Gardnerella | Type II secretion system |
| **K02232** | 0.490744223 | 1.371814895 | 0.017842745 | 0.030812255 | Cluster 2: Gardnerella | Catalyzes amidations at positions B, D, E, and G on adenosylcobyrinic A,C-diamide. NH(2) groups are provided by glutamine, and one molecule of ATP is hydrogenolyzed for each amidation |
| **K01939** | 22.25007285 | 1.370823322 | 1.28E-11 | 5.72E-11 | Cluster 2: Gardnerella | Plays an important role in the de novo pathway of purine nucleotide biosynthesis. Catalyzes the first committed step in the biosynthesis of AMP from IMP |
| **K00134** | 16.13067346 | 1.370420062 | 3.02E-13 | 1.53E-12 | Cluster 2: Gardnerella | Belongs to the glyceraldehyde-3-phosphate dehydrogenase family |
| **K02108** | 13.42302452 | 1.368632143 | 2.72E-11 | 1.18E-10 | Cluster 2: Gardnerella | it plays a direct role in the translocation of protons across the membrane |
| **K02113** | 5.345821409 | 1.368309673 | 3.62E-11 | 1.55E-10 | Cluster 2: Gardnerella | F(1)F(0) ATP synthase produces ATP from ADP in the presence of a proton or sodium gradient. F-type ATPases consist of two structural domains, F(1) containing the extramembraneous catalytic core and F(0) containing the membrane proton channel, linked together by a central stalk and a peripheral stalk. During catalysis, ATP synthesis in the catalytic domain of F(1) is coupled via a rotary mechanism of the central stalk subunits to proton translocation |
| **K06207** | 24.45616165 | 1.367446855 | 3.55E-12 | 1.67E-11 | Cluster 2: Gardnerella | GTP-binding protein TypA |
| **K00384** | 16.17302014 | 1.366047353 | 6.98E-09 | 2.40E-08 | Cluster 2: Gardnerella | Belongs to the class-II pyridine nucleotide-disulfide oxidoreductase family |

| | | | | | | |
|---|---|---|---|---|---|---|
| **K00806** | 9.182915917 | 1.364068415 | 4.77E-14 | 2.58E-13 | Cluster 2: Gardnerella | Catalyzes the condensation of isopentenyl diphosphate (IPP) with allylic pyrophosphates generating different type of terpenoids |
| **K03601** | 19.336752 | 1.358802098 | 2.82E-10 | 1.10E-09 | Cluster 2: Gardnerella | Bidirectionally degrades single-stranded DNA into large acid-insoluble oligonucleotides, which are then degraded further into small acid-soluble oligonucleotides |
| **K02072** | 4.682206801 | 1.354150285 | 2.53E-11 | 1.10E-10 | Cluster 2: Gardnerella | Binding-protein-dependent transport system inner membrane component |
| **K04047** | 5.760778015 | 1.353371574 | 4.59E-10 | 1.76E-09 | Cluster 2: Gardnerella | Belongs to the Dps family |
| **K06142** | 0.387536531 | 1.351297904 | 0.012269448 | 0.021932671 | Cluster 2: Gardnerella | PFAM Outer membrane chaperone Skp (OmpH) |
| **K03177** | 9.946160102 | 1.349344173 | 6.15E-11 | 2.58E-10 | Cluster 2: Gardnerella | Responsible for synthesis of pseudouridine from uracil- 55 in the psi GC loop of transfer RNAs |
| **K02075** | 11.47486659 | 1.346940545 | 8.70E-10 | 3.25E-09 | Cluster 2: Gardnerella | ABC 3 transport family protein |
| **K02112** | 21.72496473 | 1.3456371 | 7.07E-11 | 2.95E-10 | Cluster 2: Gardnerella | Produces ATP from ADP in the presence of a proton gradient across the membrane. The catalytic sites are hosted primarily by the beta subunits |
| **K03648** | 6.242439446 | 1.343241287 | 3.67E-11 | 1.57E-10 | Cluster 2: Gardnerella | Excises uracil residues from the DNA which can arise as a result of misincorporation of dUMP residues by DNA polymerase or due to deamination of cytosine |
| **K00609** | 11.91173628 | 1.337531545 | 1.31E-09 | 4.81E-09 | Cluster 2: Gardnerella | Belongs to the ATCase OTCase family |

| | | | | | | |
|---|---|---|---|---|---|---|
| **K00762** | 6.396994747 | 1.33747229 | 9.03E-10 | 3.37E-09 | Cluster 2: Gardnerella | Catalyzes the transfer of a ribosyl phosphate group from 5-phosphoribose 1-diphosphate to orotate, leading to the formation of orotidine monophosphate (OMP) |
| **K00963** | 12.64588482 | 1.334828751 | 3.61E-12 | 1.69E-11 | Cluster 2: Gardnerella | UTP-glucose-1-phosphate uridylyltransferase |
| **K01174** | 0.927480239 | 1.331251095 | 0.004562432 | 0.008640423 | Cluster 2: Gardnerella | nuclease |
| **K03218** | 13.43225546 | 1.323965101 | 1.27E-11 | 5.68E-11 | Cluster 2: Gardnerella | Belongs to the class IV-like SAM-binding methyltransferase superfamily. RNA methyltransferase TrmH family |
| **K08316** | 5.09070401 | 1.323413048 | 2.76E-10 | 1.08E-09 | Cluster 2: Gardnerella | Conserved hypothetical protein 95 |
| **K19046** | 0.9448377 | 1.321986083 | 0.007646795 | 0.014025971 | Cluster 2: Gardnerella | CRISPR-associated protein Cse2 (CRISPR_cse2) |
| **K04096** | 9.613972042 | 1.320776073 | 5.19E-11 | 2.19E-10 | Cluster 2: Gardnerella | DNA recombination-mediator protein A |
| **K06173** | 14.17589751 | 1.316247237 | 6.32E-10 | 2.39E-09 | Cluster 2: Gardnerella | Formation of pseudouridine at positions 38, 39 and 40 in the anticodon stem and loop of transfer RNAs |
| **K02051** | 0.775482783 | 1.313769417 | 0.014897619 | 0.026122751 | Cluster 2: Gardnerella | NMT1-like family |
| **K03501** | 11.38584036 | 1.310681742 | 2.51E-10 | 9.86E-10 | Cluster 2: Gardnerella | Specifically methylates the N7 position of a guanine in 16S rRNA |
| **K00287** | 3.780793992 | 1.308778745 | 5.47E-10 | 2.08E-09 | Cluster 2: Gardnerella | dihydrofolate reductase |

| | | | | | | |
|---|---|---|---|---|---|---|
| **K07448** | 1.834919457 | 1.307525386 | 0.010073373 | 0.018294348 | Cluster 2: Gardnerella | Restriction endonuclease |
| **K02224** | 0.451068118 | 1.307219021 | 0.018381946 | 0.031674731 | Cluster 2: Gardnerella | Catalyzes the ATP-dependent amidation of the two carboxylate groups at positions a and c of cobyrinate, using either L-glutamine or ammonia as the nitrogen source |
| **K07403** | 0.42864603 | 1.306601337 | 0.027140126 | 0.045265109 | Cluster 2: Gardnerella | Nodulation efficiency protein D |
| **K03702** | 24.17146765 | 1.304451179 | 3.11E-12 | 1.48E-11 | Cluster 2: Gardnerella | Damaged site, the DNA wraps around one UvrB monomer. DNA wrap is dependent on ATP binding by UvrB and probably causes local melting of the DNA helix, facilitating insertion of UvrB beta-hairpin between the DNA strands. Then UvrB probes one DNA strand for the presence of a lesion. If a lesion is found the UvrA subunits dissociate and the UvrB-DNA preincision complex is formed. This complex is subsequently bound by UvrC and the second UvrB is released. If no lesion is found, the DNA wraps around the other UvrB subunit that will check the other stand for damage |
| **K02548** | 8.393971315 | 1.302436716 | 1.08E-05 | 2.69E-05 | Cluster 2: Gardnerella | Belongs to the MenA family. Type 1 subfamily |
| **K01873** | 34.77911531 | 1.302273324 | 4.90E-11 | 2.07E-10 | Cluster 2: Gardnerella | amino acids such as threonine, to avoid such errors, it has a posttransfer editing activity that hydrolyzes mischarged Thr-tRNA(Val) in a tRNA-dependent manner |
| **K06958** | 9.811711394 | 1.300050886 | 1.07E-10 | 4.40E-10 | Cluster 2: Gardnerella | Displays ATPase and GTPase activities |

| | | | | | | |
|---|---|---|---|---|---|---|
| **K12960** | 0.722321771 | 1.299902304 | 0.019835237 | 0.034031742 | Cluster 2: Gardnerella | Catalyzes the deamination of 5-methylthioadenosine and S-adenosyl-L-homocysteine into 5-methylthioinosine and S-inosyl-L- homocysteine, respectively. Is also able to deaminate adenosine |
| **K02518** | 1.05378824 | 1.29870626 | 7.30E-06 | 1.86E-05 | Cluster 2: Gardnerella | One of the essential components for the initiation of protein synthesis. Stabilizes the binding of IF-2 and IF-3 on the 30S subunit to which N-formylmethionyl-tRNA(fMet) subsequently binds. Helps modulate mRNA selection, yielding the 30S pre- initiation complex (PIC). Upon addition of the 50S ribosomal subunit IF-1, IF-2 and IF-3 are released leaving the mature 70S translation initation complex |
| **K04072** | 34.72286622 | 1.296052008 | 1.90E-10 | 7.53E-10 | Cluster 2: Gardnerella | belongs to the iron- containing alcohol dehydrogenase family |
| **K02864** | 8.632532112 | 1.289756517 | 5.87E-10 | 2.22E-09 | Cluster 2: Gardnerella | Forms part of the ribosomal stalk, playing a central role in the interaction of the ribosome with GTP-bound translation factors |
| **K01784** | 16.5062845 | 1.285324627 | 2.64E-12 | 1.27E-11 | Cluster 2: Gardnerella | Belongs to the NAD(P)-dependent epimerase dehydratase family |
| **K00981** | 9.520664086 | 1.28164236 | 2.30E-09 | 8.28E-09 | Cluster 2: Gardnerella | Cytidylyltransferase family |
| **K02879** | 2.8720868 | 1.281418812 | 9.54E-08 | 2.94E-07 | Cluster 2: Gardnerella | Ribosomal protein L17 |
| **K03527** | 2.134317831 | 1.278593465 | 0.002809342 | 0.005436703 | Cluster 2: Gardnerella | Catalyzes the conversion of 1-hydroxy-2-methyl-2-(E)- butenyl 4-diphosphate (HMBPP) into a mixture of isopentenyl diphosphate (IPP) and dimethylallyl diphosphate (DMAPP). Acts in the terminal step of the |

| | | | | | | |
|---|---|---|---|---|---|---|
| | | | | | | DOXP MEP pathway for isoprenoid precursor biosynthesis |
| **K02003** | 42.21865436 | 1.278063332 | 3.04E-10 | 1.18E-09 | Cluster 2: Gardnerella | ABC transporter, ATP-binding protein |
| **K03106** | 20.51711713 | 1.277103307 | 5.75E-11 | 2.41E-10 | Cluster 2: Gardnerella | Involved in targeting and insertion of nascent membrane proteins into the cytoplasmic membrane. Binds to the hydrophobic signal sequence of the ribosome-nascent chain (RNC) as it emerges from the ribosomes. The SRP-RNC complex is then targeted to the cytoplasmic membrane where it interacts with the SRP receptor FtsY |
| **K03768** | 5.98873134 | 1.275483767 | 1.15E-09 | 4.25E-09 | Cluster 2: Gardnerella | PPIases accelerate the folding of proteins. It catalyzes the cis-trans isomerization of proline imidic peptide bonds in oligopeptides |
| **K01925** | 17.60453893 | 1.275156667 | 7.74E-10 | 2.91E-09 | Cluster 2: Gardnerella | Cell wall formation. Catalyzes the addition of glutamate to the nucleotide precursor UDP-N-acetylmuramoyl-L-alanine (UMA) |
| **K08998** | 1.640563197 | 1.27369189 | 2.70E-06 | 7.19E-06 | Cluster 2: Gardnerella | Could be involved in insertion of integral membrane proteins into the membrane |
| **K01835** | 23.17805063 | 1.27276637 | 1.34E-09 | 4.91E-09 | Cluster 2: Gardnerella | Phosphoglucomutase/phosphomannomutase, alpha/beta/alpha domain II |
| **K02025** | 12.06395423 | 1.272506618 | 8.61E-10 | 3.22E-09 | Cluster 2: Gardnerella | Binding-protein-dependent transport system inner membrane component |
| **K03856** | 0.512925516 | 1.272281397 | 0.006879721 | 0.012726419 | Cluster 2: Gardnerella | 3-deoxy-7-phosphoheptulonate synthase |

| | | | | | | |
|---|---|---|---|---|---|---|
| **K00969** | 6.351472214 | 1.27187763 | 3.40E-10 | 1.32E-09 | Cluster 2: Gardnerella | Catalyzes the reversible adenylation of nicotinate mononucleotide (NaMN) to nicotinic acid adenine dinucleotide (NaAD) |
| **K00891** | 0.471891559 | 1.2674419 | 0.020371953 | 0.034902486 | Cluster 2: Gardnerella | Catalyzes the specific phosphorylation of the 3-hydroxyl group of shikimic acid using ATP as a cosubstrate |
| **K02519** | 33.57989776 | 1.265694199 | 1.44E-10 | 5.85E-10 | Cluster 2: Gardnerella | One of the essential components for the initiation of protein synthesis. Protects formylmethionyl-tRNA from spontaneous hydrolysis and promotes its binding to the 30S ribosomal subunits. Also involved in the hydrolysis of GTP during the formation of the 70S ribosomal complex |
| **K00790** | 21.7303145 | 1.265399066 | 1.84E-11 | 8.06E-11 | Cluster 2: Gardnerella | Cell wall formation. Adds enolpyruvyl to UDP-N-acetylglucosamine |
| **K00558** | 19.67627143 | 1.264657904 | 0.001515855 | 0.003034248 | Cluster 2: Gardnerella | C-5 cytosine-specific DNA methylase |
| **K00226** | 12.22104951 | 1.245317562 | 1.88E-08 | 6.15E-08 | Cluster 2: Gardnerella | Catalyzes the conversion of dihydroorotate to orotate with quinone as electron acceptor |
| **K00262** | 2.569844396 | 1.240516678 | 0.001738604 | 0.003465607 | Cluster 2: Gardnerella | Belongs to the Glu Leu Phe Val dehydrogenases family |
| **K11991** | 4.149397505 | 1.238239026 | 5.07E-08 | 1.59E-07 | Cluster 2: Gardnerella | Catalyzes the deamination of adenosine to inosine at the wobble position 34 of tRNA(Arg2) |
| **K04763** | 19.85680335 | 1.237700012 | 2.21E-11 | 9.69E-11 | Cluster 2: Gardnerella | recombinase XerD |
| **K08314** | 0.485542294 | 1.234760222 | 0.026307627 | 0.044061128 | Cluster 2: Gardnerella | Transaldolase |

| | | | | | | |
|---|---|---|---|---|---|---|
| **K10773** | 5.374983328 | 1.232733844 | 2.65E-10 | 1.04E-09 | Cluster 2: Gardnerella | DNA repair enzyme that has both DNA N-glycosylase activity and AP-lyase activity. The DNA N-glycosylase activity releases various damaged pyrimidines from DNA by cleaving the N- glycosidic bond, leaving an AP (apurinic apyrimidinic) site. The AP-lyase activity cleaves the phosphodiester bond 3' to the AP site by a beta-elimination, leaving a 3'-terminal unsaturated sugar and a product with a terminal 5'-phosphate |
| **K09772** | 3.060271611 | 1.230638829 | 1.06E-07 | 3.25E-07 | Cluster 2: Gardnerella | Cell division protein that is part of the divisome complex and is recruited early to the Z-ring. Probably stimulates Z-ring formation, perhaps through the cross-linking of FtsZ protofilaments. Its function overlaps with FtsA |
| **K01752** | 18.84609796 | 1.227914355 | 9.96E-10 | 3.71E-09 | Cluster 2: Gardnerella | Serine dehydratase alpha chain |
| **K02115** | 8.074728874 | 1.226210387 | 2.21E-09 | 7.99E-09 | Cluster 2: Gardnerella | Produces ATP from ADP in the presence of a proton gradient across the membrane. The gamma chain is believed to be important in regulating ATPase activity and the flow of protons through the CF(0) complex |
| **K03595** | 11.11053088 | 1.225058242 | 4.79E-09 | 1.67E-08 | Cluster 2: Gardnerella | An essential GTPase that binds both GDP and GTP, with rapid nucleotide exchange. Plays a role in 16S rRNA processing and 30S ribosomal subunit biogenesis and possibly also in cell cycle regulation and energy metabolism |
| **K01958** | 2.632579406 | 1.224946217 | 0.004223115 | 0.008036023 | Cluster 2: Gardnerella | Catalyzes a 2-step reaction, involving the ATP-dependent carboxylation of the covalently attached biotin in the first step and the transfer of the carboxyl group to pyruvate in the second |

| | | | | | | |
|---|---|---|---|---|---|---|
| **K02343** | 32.50775227 | 1.223554845 | 7.54E-09 | 2.58E-08 | Cluster 2: Gardnerella | DNA polymerase III is a complex, multichain enzyme responsible for most of the replicative synthesis in bacteria. This DNA polymerase also exhibits 3' to 5' exonuclease activity |
| **K00341** | 0.854649192 | 1.220296971 | 0.011539004 | 0.020704369 | Cluster 2: Gardnerella | NADH-quinone oxidoreductase subunit L |
| **K03466** | 44.12998706 | 1.219726054 | 2.48E-08 | 7.97E-08 | Cluster 2: Gardnerella | FtsK SpoIIIE family protein |
| **K19125** | 2.356851184 | 1.218027265 | 0.021432519 | 0.036562257 | Cluster 2: Gardnerella | CRISPR-associated protein (Cas_Cas5) |
| **K03424** | 10.71143287 | 1.21659974 | 6.35E-11 | 2.65E-10 | Cluster 2: Gardnerella | TatD related DNase |
| **K01262** | 20.54548799 | 1.215110005 | 2.53E-09 | 9.06E-09 | Cluster 2: Gardnerella | Aminopeptidase P, N-terminal domain |
| **K09014** | 18.96650934 | 1.209515586 | 2.25E-08 | 7.29E-08 | Cluster 2: Gardnerella | FeS assembly protein SufB |
| **K04567** | 24.13755517 | 1.209440571 | 2.86E-11 | 1.24E-10 | Cluster 2: Gardnerella | Belongs to the class-II aminoacyl-tRNA synthetase family |
| **K03545** | 16.55043159 | 1.205426942 | 4.34E-09 | 1.52E-08 | Cluster 2: Gardnerella | Involved in protein export. Acts as a chaperone by maintaining the newly synthesized protein in an open conformation. Functions as a peptidyl-prolyl cis-trans isomerase |
| **K04068** | 7.552134017 | 1.20366434 | 1.54E-08 | 5.08E-08 | Cluster 2: Gardnerella | Activation of anaerobic ribonucleoside-triphosphate reductase under anaerobic conditions by generation of an organic free radical, using S-adenosylmethionine and reduced flavodoxin as cosubstrates to produce 5'-deoxy-adenosine |

| | | | | | | |
|---|---|---|---|---|---|---|
| **K15582** | 6.072935182 | 1.201071252 | 0.000322572 | 0.000695173 | Cluster 2: Gardnerella | N-terminal TM domain of oligopeptide transport permease C |
| **K09015** | 16.16981388 | 1.200726185 | 1.13E-08 | 3.80E-08 | Cluster 2: Gardnerella | FeS assembly protein SufD |
| **K03046** | 51.14762387 | 1.199227375 | 1.55E-09 | 5.66E-09 | Cluster 2: Gardnerella | DNA-dependent RNA polymerase catalyzes the transcription of DNA into RNA using the four ribonucleoside triphosphates as substrates |
| **K03976** | 4.180801923 | 1.196667501 | 1.40E-08 | 4.67E-08 | Cluster 2: Gardnerella | Belongs to the prolyl-tRNA editing family. YbaK EbsC subfamily |
| **K03043** | 47.14281431 | 1.1924479 | 6.46E-10 | 2.44E-09 | Cluster 2: Gardnerella | DNA-dependent RNA polymerase catalyzes the transcription of DNA into RNA using the four ribonucleoside triphosphates as substrates |
| **K01869** | 35.34940578 | 1.19101736 | 2.17E-08 | 7.04E-08 | Cluster 2: Gardnerella | Belongs to the class-I aminoacyl-tRNA synthetase family |
| **K03111** | 18.35782772 | 1.188121153 | 4.36E-09 | 1.53E-08 | Cluster 2: Gardnerella | Single-strand binding protein family |
| **K03431** | 19.42206701 | 1.187175774 | 1.30E-09 | 4.79E-09 | Cluster 2: Gardnerella | Catalyzes the conversion of glucosamine-6-phosphate to glucosamine-1-phosphate |
| **K03664** | 3.566379353 | 1.18575118 | 1.38E-09 | 5.07E-09 | Cluster 2: Gardnerella | the 2 termini fold to resemble tRNA(Ala) and it encodes a tag peptide , a short internal open reading frame. During trans-translation Ala- aminoacylated tmRNA acts like a tRNA, entering the A-site of stalled ribosomes, displacing the stalled mRNA. The ribosome then switches to translate the ORF on the tmRNA |
| **K03647** | 4.689190249 | 1.185675209 | 1.33E-06 | 3.65E-06 | Cluster 2: Gardnerella | Probably involved in ribonucleotide reductase function |

| | | | | | | |
|---|---|---|---|---|---|---|
| **K00847** | 10.46971162 | 1.185625621 | 2.40E-06 | 6.43E-06 | Cluster 2: Gardnerella | pfkB family carbohydrate kinase |
| **K03040** | 19.64559199 | 1.18161412 | 3.17E-07 | 9.29E-07 | Cluster 2: Gardnerella | DNA-dependent RNA polymerase catalyzes the transcription of DNA into RNA using the four ribonucleoside triphosphates as substrates |
| **K03596** | 21.93949668 | 1.180232756 | 2.69E-09 | 9.63E-09 | Cluster 2: Gardnerella | Required for accurate and efficient protein synthesis under certain stress conditions. May act as a fidelity factor of the translation reaction, by catalyzing a one-codon backward translocation of tRNAs on improperly translocated ribosomes. Back-translocation proceeds from a post-translocation (POST) complex to a pre-translocation (PRE) complex, thus giving elongation factor G a second chance to translocate the tRNAs correctly. Binds to ribosomes in a GTP-dependent manner |
| **K01783** | 5.054242237 | 1.178304983 | 2.77E-09 | 9.89E-09 | Cluster 2: Gardnerella | Ribulose-phosphate 3-epimerase |
| **K02335** | 32.30859665 | 1.177937804 | 5.20E-09 | 1.80E-08 | Cluster 2: Gardnerella | In addition to polymerase activity, this DNA polymerase exhibits 5'-3' exonuclease activity |
| **K03086** | 14.84355257 | 1.17573881 | 4.23E-09 | 1.49E-08 | Cluster 2: Gardnerella | Sigma factors are initiation factors that promote the attachment of RNA polymerase to specific initiation sites and are then released. This sigma factor is the primary sigma factor during exponential growth |
| **K01304** | 6.600243998 | 1.175487013 | 4.75E-06 | 1.23E-05 | Cluster 2: Gardnerella | Removes 5-oxoproline from various penultimate amino acid residues except L-proline |
| **K05601** | 1.143678371 | 1.168319877 | 0.010840182 | 0.019553233 | Cluster 2: Gardnerella | Catalyzes the reduction of hydroxylamine to form NH(3) and H(2)O |

| | | | | | | |
|---|---|---|---|---|---|---|
| **K03798** | 32.71720585 | 1.166223198 | 1.57E-09 | 5.71E-09 | Cluster 2: Gardnerella | Acts as a processive, ATP-dependent zinc metallopeptidase for both cytoplasmic and membrane proteins. Plays a role in the quality control of integral membrane proteins |
| **K02836** | 14.4906751 | 1.152443149 | 7.88E-08 | 2.44E-07 | Cluster 2: Gardnerella | Peptide chain release factor 2 directs the termination of translation in response to the peptide chain termination codons UGA and UAA |
| **K03551** | 15.19242704 | 1.150765869 | 1.74E-08 | 5.72E-08 | Cluster 2: Gardnerella | The RuvA-RuvB complex in the presence of ATP renatures cruciform structure in supercoiled DNA with palindromic sequence, indicating that it may promote strand exchange reactions in homologous recombination. RuvAB is a helicase that mediates the Holliday junction migration by localized denaturation and reannealing |
| **K01239** | 23.54368393 | 1.150398024 | 1.99E-07 | 5.94E-07 | Cluster 2: Gardnerella | Inosine-uridine preferring nucleoside hydrolase |
| **K03281** | 33.38162026 | 1.145834526 | 1.00E-07 | 3.08E-07 | Cluster 2: Gardnerella | Voltage gated chloride channel |
| **K00951** | 27.0343444 | 1.144726795 | 3.11E-09 | 1.10E-08 | Cluster 2: Gardnerella | In eubacteria ppGpp (guanosine 3'-diphosphate 5'-diphosphate) is a mediator of the stringent response that coordinates a variety of cellular activities in response to changes in nutritional abundance |
| **K06187** | 7.631043371 | 1.144268984 | 1.89E-08 | 6.17E-08 | Cluster 2: Gardnerella | May play a role in DNA repair. It seems to be involved in an RecBC-independent recombinational process of DNA repair. It may act with RecF and RecO |
| **K01808** | 0.607361538 | 1.143872621 | 0.020948527 | 0.035762128 | Cluster 2: Gardnerella | isomerase B |

| | | | | | | |
|---|---|---|---|---|---|---|
| **K02313** | 30.46799194 | 1.137488045 | 8.63E-08 | 2.67E-07 | Cluster 2: Gardnerella | it binds specifically double-stranded DNA at a 9 bp consensus (dnaA box) 5'-TTATC CA A CA A-3'. DnaA binds to ATP and to acidic phospholipids |
| **K02433** | 19.71897485 | 1.137473051 | 1.09E-07 | 3.32E-07 | Cluster 2: Gardnerella | Allows the formation of correctly charged Gln-tRNA(Gln) through the transamidation of misacylated Glu-tRNA(Gln) in organisms which lack glutaminyl-tRNA synthetase. The reaction takes place in the presence of glutamine and ATP through an activated gamma-phospho-Glu-tRNA(Gln) |
| **K01465** | 15.57606283 | 1.137099169 | 9.32E-07 | 2.58E-06 | Cluster 2: Gardnerella | Belongs to the metallo-dependent hydrolases superfamily. DHOase family. Class I DHOase subfamily |
| **K07056** | 14.94824508 | 1.134529856 | 3.51E-08 | 1.12E-07 | Cluster 2: Gardnerella | Catalyzes the 2'-O-methylation of the ribose of cytidine 1402 (C1402) in 16S rRNA |
| **K03703** | 20.41741792 | 1.133808322 | 7.88E-09 | 2.68E-08 | Cluster 2: Gardnerella | The UvrABC repair system catalyzes the recognition and processing of DNA lesions. UvrC both incises the 5' and 3' sides of the lesion. The N-terminal half is responsible for the 3' incision and the C-terminal half is responsible for the 5' incision |
| **K01874** | 32.85652701 | 1.132763118 | 1.68E-07 | 5.04E-07 | Cluster 2: Gardnerella | Is required not only for elongation of protein synthesis but also for the initiation of all mRNA translation through initiator tRNA(fMet) aminoacylation |
| **K00820** | 23.34544778 | 1.130938452 | 3.60E-08 | 1.14E-07 | Cluster 2: Gardnerella | Catalyzes the first step in hexosamine metabolism, converting fructose-6P into glucosamine-6P using glutamine as a nitrogen source |
| **K01921** | 19.07894463 | 1.128469414 | 2.47E-08 | 7.97E-08 | Cluster 2: Gardnerella | Belongs to the D-alanine--D-alanine ligase family |

| | | | | | | |
|---|---|---|---|---|---|---|
| **K02909** | 1.869305352 | 1.124373386 | 0.000120619 | 0.000273026 | Cluster 2: Gardnerella | Binds the 23S rRNA |
| **K02342** | 14.84281142 | 1.118426951 | 1.20E-08 | 4.02E-08 | Cluster 2: Gardnerella | Exonuclease |
| **K06871** | 2.02406209 | 1.113282891 | 0.016828892 | 0.0292137 | Cluster 2: Gardnerella | six-Cys-in-45 modification radical SAM protein |
| **K03531** | 15.07796152 | 1.106804485 | 3.88E-08 | 1.23E-07 | Cluster 2: Gardnerella | Essential cell division protein that forms a contractile ring structure (Z ring) at the future cell division site. The regulation of the ring assembly controls the timing and the location of cell division. One of the functions of the FtsZ ring is to recruit other cell division proteins to the septum to produce a new cell wall between the dividing cells. Binds GTP and shows GTPase activity |
| **K01868** | 25.80107799 | 1.104504497 | 1.28E-07 | 3.87E-07 | Cluster 2: Gardnerella | Catalyzes the attachment of threonine to tRNA(Thr) in a two-step reaction L-threonine is first activated by ATP to form Thr-AMP and then transferred to the acceptor end of tRNA(Thr) |
| **K00763** | 21.76868815 | 1.098675755 | 1.60E-07 | 4.79E-07 | Cluster 2: Gardnerella | Catalyzes the synthesis of beta-nicotinate D-ribonucleotide from nicotinate and 5-phospho-D-ribose 1-phosphate at the expense of ATP |
| **K03816** | 7.212272568 | 1.098671342 | 2.16E-07 | 6.43E-07 | Cluster 2: Gardnerella | Converts the preformed base xanthine, a product of nucleic acid breakdown, to xanthosine 5'-monophosphate (XMP), so it can be reused for RNA or DNA synthesis |
| **K01972** | 26.03289443 | 1.095917074 | 2.76E-08 | 8.84E-08 | Cluster 2: Gardnerella | DNA ligase that catalyzes the formation of phosphodiester linkages between 5'-phosphoryl and 3'-hydroxyl groups in double-stranded DNA using NAD as a coenzyme and as the energy source for the |

| | | | | | | |
|---|---|---|---|---|---|---|
| | | | | | | reaction. It is essential for DNA replication and repair of damaged DNA |
| **K04077** | 21.72932693 | 1.09529636 | 4.95E-10 | 1.89E-09 | Cluster 2: Gardnerella | Prevents misfolding and promotes the refolding and proper assembly of unfolded polypeptides generated under stress conditions |
| **K06926** | 6.917828844 | 1.093770826 | 0.026229619 | 0.043961283 | Cluster 2: Gardnerella | AAA domain, putative AbiEii toxin, Type IV TA system |
| **K02963** | 1.337874412 | 1.093662614 | 4.80E-05 | 0.000112848 | Cluster 2: Gardnerella | Binds as a heterodimer with protein S6 to the central domain of the 16S rRNA, where it helps stabilize the platform of the 30S subunit |
| **K01887** | 26.52694984 | 1.091481274 | 7.46E-08 | 2.32E-07 | Cluster 2: Gardnerella | Arginyl-tRNA synthetase |
| **K02355** | 33.42024958 | 1.090661762 | 2.44E-07 | 7.21E-07 | Cluster 2: Gardnerella | Catalyzes the GTP-dependent ribosomal translocation step during translation elongation. During this step, the ribosome changes from the pre-translocational (PRE) to the post- translocational (POST) state as the newly formed A-site-bound peptidyl-tRNA and P-site-bound deacylated tRNA move to the P and E sites, respectively. Catalyzes the coordinated movement of the two tRNA molecules, the mRNA and conformational changes in the ribosome |
| **K02899** | 1.111480875 | 1.088836626 | 0.000131128 | 0.000294545 | Cluster 2: Gardnerella | Belongs to the bacterial ribosomal protein bL27 family |
| **K02340** | 11.21991584 | 1.086337065 | 1.87E-08 | 6.12E-08 | Cluster 2: Gardnerella | DNA polymerase III delta subunit |
| **K02956** | 2.213572786 | 1.080550211 | 6.36E-06 | 1.63E-05 | Cluster 2: Gardnerella | Forms an intersubunit bridge (bridge B4) with the 23S rRNA of the 50S subunit in the ribosome |

| | | | | | | |
|---|---|---|---|---|---|---|
| **K06950** | 8.148035804 | 1.074742223 | 6.36E-07 | 1.80E-06 | Cluster 2: Gardnerella | HD domain |
| **K03438** | 14.46447352 | 1.067891182 | 4.02E-07 | 1.16E-06 | Cluster 2: Gardnerella | Specifically methylates the N4 position of cytidine in position 1402 (C1402) of 16S rRNA |
| **K09762** | 10.23469872 | 1.063998631 | 5.47E-07 | 1.56E-06 | Cluster 2: Gardnerella | May be required for sporulation |
| **K01890** | 27.70602041 | 1.055866398 | 1.07E-07 | 3.29E-07 | Cluster 2: Gardnerella | Belongs to the phenylalanyl-tRNA synthetase beta subunit family. Type 1 subfamily |
| **K02039** | 6.03491124 | 1.052572488 | 9.65E-05 | 0.000221591 | Cluster 2: Gardnerella | Plays a role in the regulation of phosphate uptake |
| **K01641** | 13.59375168 | 1.050366482 | 7.86E-06 | 1.99E-05 | Cluster 2: Gardnerella | Hydroxymethylglutaryl-coenzyme A synthase N terminal |
| **K03498** | 3.371459333 | 1.049858991 | 0.011627938 | 0.020848292 | Cluster 2: Gardnerella | Cation transport protein |
| **K03076** | 27.63440965 | 1.048878163 | 9.50E-07 | 2.62E-06 | Cluster 2: Gardnerella | The central subunit of the protein translocation channel SecYEG. Consists of two halves formed by TMs 1-5 and 6-10. These two domains form a lateral gate at the front which open onto the bilayer between TMs 2 and 7, and are clamped together by SecE at the back. The channel is closed by both a pore ring composed of hydrophobic SecY resides and a short helix (helix 2A) on the extracellular side of the membrane which forms a plug. The plug probably moves laterally to allow the channel to open. The ring and the pore may move independently |
| **K00036** | 21.45425602 | 1.045088859 | 2.96E-06 | 7.83E-06 | Cluster 2: Gardnerella | Catalyzes the oxidation of glucose 6-phosphate to 6-phosphogluconolactone |

| | | | | | | |
|---|---|---|---|---|---|---|
| **K02434** | 20.10608612 | 1.040180641 | 5.63E-07 | 1.60E-06 | Cluster 2: Gardnerella | Allows the formation of correctly charged Asn-tRNA(Asn) or Gln-tRNA(Gln) through the transamidation of misacylated Asp- tRNA(Asn) or Glu-tRNA(Gln) in organisms which lack either or both of asparaginyl-tRNA or glutaminyl-tRNA synthetases. The reaction takes place in the presence of glutamine and ATP through an activated phospho-Asp-tRNA(Asn) or phospho-Glu-tRNA(Gln) |
| **K01560** | 11.04897887 | 1.035843863 | 4.39E-07 | 1.26E-06 | Cluster 2: Gardnerella | Haloacid dehalogenase-like hydrolase |
| **K03553** | 17.05335784 | 1.034372353 | 7.53E-07 | 2.10E-06 | Cluster 2: Gardnerella | Can catalyze the hydrolysis of ATP in the presence of single-stranded DNA, the ATP-dependent uptake of single-stranded DNA by duplex DNA, and the ATP-dependent hybridization of homologous single-stranded DNAs. It interacts with LexA causing its activation and leading to its autocatalytic cleavage |
| **K00033** | 24.15322871 | 1.032756781 | 3.46E-06 | 9.11E-06 | Cluster 2: Gardnerella | Catalyzes the oxidative decarboxylation of 6-phosphogluconate to ribulose 5-phosphate and CO(2), with concomitant reduction of NADP to NADPH |
| **K00927** | 17.93588403 | 1.027155397 | 7.14E-07 | 2.00E-06 | Cluster 2: Gardnerella | Belongs to the phosphoglycerate kinase family |
| **K03308** | 2.028760546 | 1.026041391 | 0.023563567 | 0.039799947 | Cluster 2: Gardnerella | Belongs to the sodium neurotransmitter symporter (SNF) (TC 2.A.22) family |
| **K01256** | 33.38169021 | 1.025392069 | 1.16E-07 | 3.54E-07 | Cluster 2: Gardnerella | Peptidase family M1 domain |
| **K01876** | 20.94774372 | 1.023649871 | 7.05E-07 | 1.98E-06 | Cluster 2: Gardnerella | Aspartyl-tRNA synthetase with relaxed tRNA specificity since it is able to aspartylate not only its cognate tRNA(Asp) but also tRNA(Asn). Reaction proceeds in two steps L-aspartate is first activated by |

| | | | | | | |
|---|---|---|---|---|---|---|
| | | | | | | ATP to form Asp-AMP and then transferred to the acceptor end of tRNA(Asp Asn) |
| **K02878** | 5.342088756 | 1.022785043 | 6.13E-07 | 1.74E-06 | Cluster 2: Gardnerella | Binds 23S rRNA and is also seen to make contacts with the A and possibly P site tRNAs |
| **K02952** | 3.877005712 | 1.01877857 | 3.75E-06 | 9.84E-06 | Cluster 2: Gardnerella | Located at the top of the head of the 30S subunit, it contacts several helices of the 16S rRNA. In the 70S ribosome it contacts the 23S rRNA (bridge B1a) and protein L5 of the 50S subunit (bridge B1b), connecting the 2 subunits |
| **K04485** | 19.50743931 | 1.008016407 | 2.63E-07 | 7.75E-07 | Cluster 2: Gardnerella | DNA-dependent ATPase involved in processing of recombination intermediates, plays a role in repairing DNA breaks. Stimulates the branch migration of RecA-mediated strand transfer reactions, allowing the 3' invading strand to extend heteroduplex DNA faster. Binds ssDNA in the presence of ADP but not other nucleotides, has ATPase activity that is stimulated by ssDNA and various branched DNA structures, but inhibited by SSB. Does not have RecA's homology-searching function |
| **K09903** | 7.225612509 | 1.003604572 | 2.41E-07 | 7.13E-07 | Cluster 2: Gardnerella | Catalyzes the reversible phosphorylation of UMP to UDP |
| **K19157** | 3.184743658 | 1.001947926 | 0.001818445 | 0.003600732 | Cluster 2: Gardnerella | Bacterial toxin of type II toxin-antitoxin system, YafQ |
| **K18197** | 21.98515734 | 4.64240639 | 6.05E-22 | 1.09E-19 | Cluster 3: L. iners | domain, Protein |
| **K13924** | 16.13207118 | 4.390347625 | 9.47E-22 | 1.56E-19 | Cluster 3: L. iners | Transposase |

| | | | | | | |
|---|---|---|---|---|---|---|
| **K02557** | 16.1032028 | 4.350729276 | 1.78E-23 | 3.90E-21 | Cluster 3: L. iners | Transposase |
| **K03112** | 32.08025887 | 4.296662379 | 1.99E-20 | 2.30E-18 | Cluster 3: L. iners | cell adhesion |
| **K01487** | 6.764059649 | 4.092862924 | 2.49E-23 | 4.91E-21 | Cluster 3: L. iners | Amidohydrolase family |
| **K10974** | 4.451564991 | 4.043437734 | 2.19E-19 | 2.05E-17 | Cluster 3: L. iners | cytosine purines uracil thiamine allantoin |
| **K01206** | 23.1115665 | 4.025377501 | 1.03E-23 | 2.91E-21 | Cluster 3: L. iners | domain, Protein |
| **K19084** | 7.327776999 | 3.973305759 | 3.56E-18 | 2.42E-16 | Cluster 3: L. iners | FtsX-like permease family |
| **K18935** | 4.586760922 | 3.836667485 | 2.22E-17 | 1.29E-15 | Cluster 3: L. iners | Sugar (and other) transporter |
| **K03762** | 5.772142207 | 3.710022815 | 5.68E-16 | 2.55E-14 | Cluster 3: L. iners | transmembrane transporter activity |
| **K11704** | 4.47543986 | 3.645460766 | 6.23E-16 | 2.73E-14 | Cluster 3: L. iners | Belongs to the bacterial solute-binding protein 9 family |
| **K03646** | 9.078179085 | 3.621953136 | 2.48E-19 | 2.12E-17 | Cluster 3: L. iners | energy transducer activity |
| **K08981** | 7.213960517 | 3.605781348 | 2.23E-27 | 1.47E-24 | Cluster 3: L. iners | Membrane |
| **K02538** | 8.028309687 | 3.582106798 | 4.99E-15 | 1.76E-13 | Cluster 3: L. iners | transcriptional antiterminator |

| | | | | | | |
|---|---|---|---|---|---|---|
| **K11707** | 4.605096411 | 3.542489797 | 2.76E-19 | 2.27E-17 | Cluster 3: L. iners | Belongs to the bacterial solute-binding protein 9 family |
| **K19971** | 4.558675951 | 3.522274001 | 7.36E-16 | 3.16E-14 | Cluster 3: L. iners | Belongs to the bacterial solute-binding protein 9 family |
| **K19972** | 1.992663768 | 3.506706597 | 6.85E-13 | 1.88E-11 | Cluster 3: L. iners | ABC 3 transport family |
| **K02445** | 6.149817313 | 3.413120695 | 1.86E-15 | 7.51E-14 | Cluster 3: L. iners | transmembrane transporter activity |
| **K08196** | 5.586198611 | 3.384646642 | 3.31E-12 | 8.05E-11 | Cluster 3: L. iners | Major Facilitator |
| **K11050** | 3.920133662 | 3.320387095 | 6.22E-10 | 1.03E-08 | Cluster 3: L. iners | ABC transporter |
| **K01113** | 4.855006366 | 3.286210655 | 2.88E-15 | 1.07E-13 | Cluster 3: L. iners | Associated with various cellular activities |
| **K11705** | 2.075392803 | 3.188477857 | 1.08E-09 | 1.66E-08 | Cluster 3: L. iners | ABC 3 transport family |
| **K01772** | 4.196044073 | 3.159111896 | 8.43E-17 | 4.26E-15 | Cluster 3: L. iners | Catalyzes the ferrous insertion into protoporphyrin IX |
| **K22212** | 0.95701625 | 3.153826003 | 7.44E-07 | 3.82E-06 | Cluster 3: L. iners | Malic enzyme |
| **K19509** | 2.719133292 | 3.089639359 | 6.76E-12 | 1.59E-10 | Cluster 3: L. iners | PTS system mannose/fructose/sorbose family IID component |
| **K01628** | 3.907291091 | 3.084555116 | 1.44E-12 | 3.73E-11 | Cluster 3: L. iners | Class II Aldolase and Adducin N-terminal domain |

| | | | | | | |
|---|---|---|---|---|---|---|
| **K03297** | 1.464408419 | 3.077437693 | 4.88E-11 | 1.05E-09 | Cluster 3: L. iners | Multidrug Resistance protein |
| **K03309** | 0.837983182 | 3.076104798 | 0.000183178 | 0.000551491 | Cluster 3: L. iners | Belongs to the dicarboxylate amino acid cation symporter (DAACS) (TC 2.A.23) family |
| **K19976** | 7.292272407 | 3.055157418 | 2.41E-15 | 9.18E-14 | Cluster 3: L. iners | Belongs to the bacterial solute-binding protein 9 family |
| **K19975** | 5.021221526 | 3.021158025 | 1.20E-13 | 3.63E-12 | Cluster 3: L. iners | Belongs to the bacterial solute-binding protein 9 family |
| **K07043** | 3.656489031 | 3.004093205 | 3.01E-15 | 1.10E-13 | Cluster 3: L. iners | Protein of unknown function DUF45 |
| **K03092** | 4.180430314 | 2.93912258 | 3.73E-15 | 1.34E-13 | Cluster 3: L. iners | Sigma-54 factor, core binding domain |
| **K00105** | 3.95383568 | 2.881015306 | 2.19E-06 | 1.02E-05 | Cluster 3: L. iners | C-terminal domain of alpha-glycerophosphate oxidase |
| **K20424** | 1.362903137 | 2.879595174 | 0.002208154 | 0.005597019 | Cluster 3: L. iners | alcohol dehydrogenase |
| **K19082** | 1.191859583 | 2.864783398 | 2.47E-10 | 4.55E-09 | Cluster 3: L. iners | response regulator |
| **K11709** | 2.301250978 | 2.839126609 | 1.59E-11 | 3.56E-10 | Cluster 3: L. iners | ABC 3 transport family |
| **K19083** | 1.695009626 | 2.78819949 | 5.97E-09 | 6.69E-08 | Cluster 3: L. iners | ABC transporter, ATP-binding protein |
| **K06610** | 0.562957129 | 2.656122334 | 0.000598681 | 0.001679374 | Cluster 3: L. iners | Major Facilitator Superfamily |

| | | | | | | |
|---|---|---|---|---|---|---|
| **K00433** | 1.715763182 | 2.592374693 | 1.69E-09 | 2.41E-08 | Cluster 3: L. iners | Alpha/beta hydrolase family |
| **K01426** | 5.027298807 | 2.590558659 | 2.34E-09 | 3.11E-08 | Cluster 3: L. iners | Belongs to the amidase family |
| **K16926** | 2.807227522 | 2.563258004 | 2.25E-13 | 6.53E-12 | Cluster 3: L. iners | Hypothetical bacterial integral membrane protein (Trep_Strep) |
| **K18104** | 3.684105116 | 2.553992348 | 1.14E-08 | 1.12E-07 | Cluster 3: L. iners | (ABC) transporter |
| **K07777** | 0.469942277 | 2.499892827 | 0.008593731 | 0.019385401 | Cluster 3: L. iners | Sensor histidine kinase |
| **K02171** | 2.595285627 | 2.484343246 | 7.90E-10 | 1.29E-08 | Cluster 3: L. iners | Copper transport repressor CopY TcrY |
| **K03300** | 0.91726895 | 2.466016208 | 4.00E-05 | 0.000136969 | Cluster 3: L. iners | Citrate transporter |
| **K07483** | 5.635692046 | 2.4440848 | 2.22E-09 | 3.00E-08 | Cluster 3: L. iners | the current gene model (or a revised gene model) may contain a frame shift |
| **K02771** | 0.759468063 | 2.428012838 | 0.002593372 | 0.006457235 | Cluster 3: L. iners | PTS system mannose/fructose/sorbose family IID component |
| **K08166** | 0.59997747 | 2.420332573 | 0.005995774 | 0.013959465 | Cluster 3: L. iners | Major Facilitator Superfamily |
| **K07757** | 4.197265569 | 2.399240829 | 1.79E-12 | 4.47E-11 | Cluster 3: L. iners | Sucrose-6F-phosphate phosphohydrolase |
| **K05881** | 1.038761914 | 2.320522482 | 1.71E-07 | 1.08E-06 | Cluster 3: L. iners | PTS system fructose IIA component |

| | | | | | | |
|---|---|---|---|---|---|---|
| **K02245** | 0.762232584 | 2.317908738 | 5.20E-09 | 5.96E-08 | Cluster 3: L. iners | competence protein ComGC |
| **K11085** | 3.904158552 | 2.310637393 | 3.83E-09 | 4.66E-08 | Cluster 3: L. iners | (ABC) transporter |
| **K01220** | 3.990698307 | 2.304723207 | 1.42E-06 | 6.89E-06 | Cluster 3: L. iners | Belongs to the glycosyl hydrolase 1 family |
| **K03797** | 5.624857501 | 2.274605509 | 3.06E-10 | 5.44E-09 | Cluster 3: L. iners | Belongs to the peptidase S41A family |
| **K05846** | 5.228621601 | 2.233287773 | 4.16E-08 | 3.22E-07 | Cluster 3: L. iners | Periplasmic glycine betaine choline-binding (lipo)protein of an ABC-type transport system (osmoprotectant binding protein) |
| **K07023** | 3.769223752 | 2.227075648 | 1.60E-10 | 3.18E-09 | Cluster 3: L. iners | HD containing hydrolase-like enzyme |
| **K03832** | 11.91040138 | 2.213649192 | 7.36E-07 | 3.79E-06 | Cluster 3: L. iners | energy transducer activity |
| **K05845** | 5.431628965 | 2.190855347 | 3.43E-07 | 1.96E-06 | Cluster 3: L. iners | Periplasmic glycine betaine choline-binding (lipo)protein of an ABC-type transport system (osmoprotectant binding protein) |
| **K20459** | 7.871399378 | 2.151204338 | 7.50E-06 | 2.99E-05 | Cluster 3: L. iners | abc transporter atp-binding protein |
| **K01251** | 6.504677214 | 2.147597306 | 5.92E-08 | 4.33E-07 | Cluster 3: L. iners | adenosylhomocysteinase activity |
| **K01263** | 9.876108868 | 2.145285615 | 1.36E-09 | 2.00E-08 | Cluster 3: L. iners | Cleaves the N-terminal amino acid of tripeptides |
| **K22457** | 0.954764629 | 2.113887226 | 0.00097828 | 0.002635475 | Cluster 3: L. iners | Aminotransferase |

| | | | | | | |
|---|---|---|---|---|---|---|
| **K03306** | 6.494424086 | 2.066778982 | 8.37E-13 | 2.20E-11 | Cluster 3: L. iners | Phosphate transporter family |
| **K05847** | 2.589177993 | 2.045212263 | 2.51E-06 | 1.14E-05 | Cluster 3: L. iners | ABC transporter, ATP-binding protein |
| **K07683** | 0.612989663 | 2.029673976 | 0.010012333 | 0.022309966 | Cluster 3: L. iners | Sensor histidine kinase |
| **K15539** | 4.942175529 | 2.008767593 | 3.16E-10 | 5.57E-09 | Cluster 3: L. iners | Helix-turn-helix domain |
| **K06198** | 8.810498007 | 2.004636727 | 1.29E-08 | 1.26E-07 | Cluster 3: L. iners | Competence protein |
| **K05879** | 2.827471074 | 1.995335346 | 6.61E-09 | 7.22E-08 | Cluster 3: L. iners | Dak2 |
| **K18889** | 14.28234707 | 1.977673399 | 3.38E-12 | 8.13E-11 | Cluster 3: L. iners | ABC transporter |
| **K01635** | 2.983024937 | 1.960340715 | 1.12E-05 | 4.33E-05 | Cluster 3: L. iners | Belongs to the aldolase LacD family |
| **K20114** | 2.096103197 | 1.917782389 | 0.004031151 | 0.009694427 | Cluster 3: L. iners | PTS system sugar-specific permease component |
| **K07484** | 23.95759275 | 1.915494193 | 2.23E-09 | 3.00E-08 | Cluster 3: L. iners | Transposase |
| **K07317** | 0.626434122 | 1.912965139 | 0.00216616 | 0.005511828 | Cluster 3: L. iners | type I restriction modification DNA specificity domain |
| **K17217** | 1.038174146 | 1.900355072 | 0.010377823 | 0.022994456 | Cluster 3: L. iners | cystathionine |

| | | | | | | |
|---|---|---|---|---|---|---|
| **K07778** | 1.103334594 | 1.88089132 | 0.000891927 | 0.002432753 | Cluster 3: L. iners | Histidine kinase |
| **K02530** | 2.226626726 | 1.880781498 | 0.000238791 | 0.000712399 | Cluster 3: L. iners | DeoR C terminal sensor domain |
| **K01485** | 6.218954994 | 1.878652069 | 5.58E-07 | 3.00E-06 | Cluster 3: L. iners | cytosine deaminase |
| **K05350** | 4.494034125 | 1.87792429 | 3.83E-05 | 0.000131861 | Cluster 3: L. iners | Belongs to the glycosyl hydrolase 1 family |
| **K00111** | 5.56434127 | 1.874453225 | 3.36E-07 | 1.92E-06 | Cluster 3: L. iners | C-terminal domain of alpha-glycerophosphate oxidase |
| **K02749** | 13.65606213 | 1.834218648 | 2.63E-07 | 1.55E-06 | Cluster 3: L. iners | phosphotransferase system, EIIB |
| **K02750** | 13.65606213 | 1.834218648 | 2.63E-07 | 1.55E-06 | Cluster 3: L. iners | phosphotransferase system, EIIB |
| **K07258** | 21.17620155 | 1.816688883 | 2.33E-19 | 2.09E-17 | Cluster 3: L. iners | Belongs to the peptidase S11 family |
| **K02795** | 9.899841501 | 1.814612606 | 9.21E-20 | 9.08E-18 | Cluster 3: L. iners | PTS system sorbose-specific iic component |
| **K02796** | 13.73358642 | 1.801877543 | 3.96E-13 | 1.13E-11 | Cluster 3: L. iners | system, mannose fructose sorbose family IID component |
| **K05311** | 6.514495413 | 1.798426436 | 5.16E-08 | 3.84E-07 | Cluster 3: L. iners | Putative sugar-binding domain |
| **K13678** | 5.667069629 | 1.774612373 | 3.80E-06 | 1.64E-05 | Cluster 3: L. iners | Glycosyltransferase, group 1 family protein |

| | | | | | | |
|---|---|---|---|---|---|---|
| **K01274** | 9.386210602 | 1.766313166 | 1.30E-09 | 1.94E-08 | Cluster 3: L. iners | dipeptidase PepV |
| **K01439** | 17.17381108 | 1.747362452 | 3.39E-14 | 1.13E-12 | Cluster 3: L. iners | dipeptidase PepV |
| **K08368** | 10.60540462 | 1.742520279 | 2.59E-07 | 1.53E-06 | Cluster 3: L. iners | Sugar (and other) transporter |
| **K01819** | 1.400178526 | 1.740890565 | 0.000867456 | 0.002372571 | Cluster 3: L. iners | Ribose/Galactose Isomerase |
| **K02058** | 17.86526294 | 1.736500346 | 2.02E-07 | 1.25E-06 | Cluster 3: L. iners | ABC transporter substrate-binding protein PnrA-like |
| **K13677** | 5.96361542 | 1.731552973 | 5.95E-07 | 3.17E-06 | Cluster 3: L. iners | Glycosyltransferase, group 1 family protein |
| **K00691** | 18.04617791 | 1.730501417 | 2.03E-07 | 1.25E-06 | Cluster 3: L. iners | hydrolase, family 65, central catalytic |
| **K07335** | 18.08225135 | 1.729386565 | 4.40E-08 | 3.39E-07 | Cluster 3: L. iners | ABC transporter substrate-binding protein PnrA-like |
| **K03339** | 5.147521548 | 1.723149731 | 1.04E-06 | 5.16E-06 | Cluster 3: L. iners | Fructose-1,6-bisphosphate aldolase, class II |
| **K04087** | 5.882774958 | 1.722305587 | 1.08E-06 | 5.34E-06 | Cluster 3: L. iners | prohibitin homologues |
| **K02240** | 10.0887669 | 1.709921731 | 1.31E-07 | 8.59E-07 | Cluster 3: L. iners | Helicase C-terminal domain protein |
| **K03152** | 5.043318996 | 1.700432563 | 6.77E-11 | 1.42E-09 | Cluster 3: L. iners | DJ-1/PfpI family |

| | | | | | | |
|---|---|---|---|---|---|---|
| **K06885** | 13.47203616 | 1.693520045 | 2.18E-07 | 1.33E-06 | Cluster 3: L. iners | Metal dependent phosphohydrolases with conserved 'HD' motif. |
| **K07033** | 16.89888363 | 1.688846231 | 9.91E-09 | 1.01E-07 | Cluster 3: L. iners | assembly protein SufB |
| **K02086** | 1.90011258 | 1.683574693 | 4.33E-06 | 1.83E-05 | Cluster 3: L. iners | DnaD domain protein |
| **K16169** | 11.2644362 | 1.671473717 | 1.21E-06 | 5.91E-06 | Cluster 3: L. iners | xanthine permease |
| **K00561** | 1.09491363 | 1.671044886 | 0.00189406 | 0.004852019 | Cluster 3: L. iners | Belongs to the class I-like SAM-binding methyltransferase superfamily. rRNA adenine N(6)-methyltransferase family |
| **K08177** | 1.357137958 | 1.667454707 | 0.004678386 | 0.011209936 | Cluster 3: L. iners | Major Facilitator Superfamily |
| **K01493** | 3.510773192 | 1.663394499 | 1.25E-11 | 2.82E-10 | Cluster 3: L. iners | MafB19-like deaminase |
| **K01598** | 8.653675717 | 1.661739014 | 4.03E-08 | 3.15E-07 | Cluster 3: L. iners | Catalyzes two steps in the biosynthesis of coenzyme A. In the first step cysteine is conjugated to 4'-phosphopantothenate to form 4-phosphopantothenoylcysteine, in the latter compound is decarboxylated to form 4'-phosphopantotheine |
| **K07461** | 6.960908775 | 1.65590826 | 7.00E-08 | 4.99E-07 | Cluster 3: L. iners | Methyltransferase small domain |
| **K01258** | 19.55270028 | 1.653070647 | 6.73E-11 | 1.42E-09 | Cluster 3: L. iners | Cleaves the N-terminal amino acid of tripeptides |
| **K19784** | 4.078484726 | 1.645288942 | 1.69E-08 | 1.56E-07 | Cluster 3: L. iners | NADPH-dependent FMN reductase |

| | | | | | | |
|---|---|---|---|---|---|---|
| **K22393** | 2.849473832 | 1.636712803 | 2.50E-05 | 8.89E-05 | Cluster 3: L. iners | NADPH-dependent FMN reductase |
| **K01101** | 6.089124437 | 1.636051998 | 7.90E-07 | 4.02E-06 | Cluster 3: L. iners | Catalyzes the dephosphorylation of 2-6 carbon acid sugars in vitro |
| **K02244** | 7.20999347 | 1.626078308 | 8.80E-06 | 3.46E-05 | Cluster 3: L. iners | type II secretion system |
| **K00796** | 9.262949729 | 1.623992332 | 3.81E-05 | 0.000131641 | Cluster 3: L. iners | dihydropteroate synthase |
| **K09951** | 3.155429765 | 1.623354645 | 1.55E-05 | 5.79E-05 | Cluster 3: L. iners | CRISPR-associated endoribonuclease Cas2 |
| **K03740** | 15.33275723 | 1.622094104 | 1.63E-06 | 7.78E-06 | Cluster 3: L. iners | Protein involved in D-alanine esterification of lipoteichoic acid and wall teichoic acid (D-alanine transfer protein) |
| **K06286** | 11.75224364 | 1.620549137 | 1.11E-06 | 5.45E-06 | Cluster 3: L. iners | modulates the frequency and position of FtsZ ring formation. Inhibits FtsZ ring formation at polar sites. Interacts either with FtsZ or with one of its binding partners to promote depolymerization |
| **K22103** | 5.246448659 | 1.61929485 | 2.06E-05 | 7.46E-05 | Cluster 3: L. iners | DeoR C terminal sensor domain |
| **K12555** | 14.24813477 | 1.615133298 | 3.21E-06 | 1.40E-05 | Cluster 3: L. iners | penicillin-binding protein |
| **K15460** | 7.118181908 | 1.608218879 | 1.37E-08 | 1.31E-07 | Cluster 3: L. iners | Methyltransferase small domain |
| **K22394** | 2.406160248 | 1.607377552 | 8.60E-05 | 0.000274267 | Cluster 3: L. iners | NADPH-dependent FMN reductase |

| | | | | | | |
|---|---|---|---|---|---|---|
| **K17240** | 7.593789865 | 1.60684154 | 1.68E-05 | 6.23E-05 | Cluster 3: L. iners | ATPases associated with a variety of cellular activities |
| **K20707** | 9.525463276 | 1.606801257 | 3.89E-06 | 1.67E-05 | Cluster 3: L. iners | Catalyzes the interconversion of L-alanine and D-alanine. May also act on other amino acids |
| **K02248** | 3.047648744 | 1.605909475 | 1.76E-05 | 6.47E-05 | Cluster 3: L. iners | Putative Competence protein ComGF |
| **K06153** | 4.662905197 | 1.603932595 | 3.57E-07 | 2.02E-06 | Cluster 3: L. iners | Catalyzes the dephosphorylation of undecaprenyl diphosphate (UPP). Confers resistance to bacitracin |
| **K00882** | 10.75567019 | 1.600348771 | 7.85E-12 | 1.80E-10 | Cluster 3: L. iners | Belongs to the carbohydrate kinase PfkB family. LacC subfamily |
| **K07742** | 0.582410838 | 1.596072638 | 6.35E-05 | 0.000208322 | Cluster 3: L. iners | Protein of unknown function (DUF448) |
| **K07652** | 15.99677571 | 1.594676898 | 1.91E-07 | 1.19E-06 | Cluster 3: L. iners | Histidine kinase |
| **K07220** | 1.463374057 | 1.588548244 | 2.89E-05 | 0.000101146 | Cluster 3: L. iners | Protein of unknown function DUF47 |
| **K16511** | 3.998944193 | 1.584445026 | 1.91E-05 | 6.95E-05 | Cluster 3: L. iners | Enables the recognition and targeting of unfolded and aggregated proteins to the ClpC protease or to other proteins involved in proteolysis |
| **K13052** | 1.649205368 | 1.577181502 | 2.89E-06 | 1.27E-05 | Cluster 3: L. iners | Septum formation initiator |
| **K12552** | 12.53248308 | 1.574449564 | 2.68E-06 | 1.20E-05 | Cluster 3: L. iners | Penicillin-binding Protein |
| **K12556** | 12.53248308 | 1.574449564 | 2.68E-06 | 1.20E-05 | Cluster 3: L. iners | Penicillin-binding Protein |

| | | | | | | |
|---|---|---|---|---|---|---|
| **K08724** | 12.53671265 | 1.572715063 | 2.09E-06 | 9.76E-06 | Cluster 3: L. iners | Penicillin-binding Protein |
| **K19350** | 12.42482898 | 1.568082164 | 4.44E-07 | 2.46E-06 | Cluster 3: L. iners | ABC transporter |
| **K19124** | 7.046609504 | 1.559655526 | 0.000758935 | 0.002093176 | Cluster 3: L. iners | CRISPR system CASCADE complex protein CasC |
| **K03698** | 6.750428472 | 1.556215639 | 4.33E-06 | 1.83E-05 | Cluster 3: L. iners | Metal dependent phosphohydrolases with conserved 'HD' motif. |
| **K01515** | 7.679833941 | 1.553382227 | 2.24E-09 | 3.00E-08 | Cluster 3: L. iners | NUDIX domain |
| **K07507** | 7.904926891 | 1.547479143 | 3.46E-10 | 6.03E-09 | Cluster 3: L. iners | MgtC family |
| **K01495** | 1.564051183 | 1.543405609 | 1.59E-05 | 5.93E-05 | Cluster 3: L. iners | GTP cyclohydrolase 1 |
| **K03305** | 13.66310535 | 1.535142328 | 1.72E-06 | 8.10E-06 | Cluster 3: L. iners | amino acid peptide transporter |
| **K17836** | 7.386162913 | 1.532200696 | 1.08E-07 | 7.37E-07 | Cluster 3: L. iners | Beta-lactamase enzyme family |
| **K20265** | 11.86348616 | 1.521050242 | 3.06E-05 | 0.000106731 | Cluster 3: L. iners | amino acid |
| **K18890** | 9.976236958 | 1.513543946 | 1.00E-06 | 5.00E-06 | Cluster 3: L. iners | ABC transporter |
| **K07444** | 6.990242042 | 1.508576118 | 1.98E-07 | 1.23E-06 | Cluster 3: L. iners | Belongs to the methyltransferase superfamily |

| | | | | | | |
|---|---|---|---|---|---|---|
| **K08591** | 4.752515484 | 1.501065271 | 2.62E-10 | 4.74E-09 | Cluster 3: L. iners | Catalyzes the transfer of an acyl group from acyl-phosphate (acyl-PO(4)) to glycerol-3-phosphate (G3P) to form lysophosphatidic acid (LPA). This enzyme utilizes acyl-phosphate as fatty acyl donor, but not acyl-CoA or acyl-ACP |
| **K07404** | 8.450852602 | 1.495079492 | 2.32E-06 | 1.07E-05 | Cluster 3: L. iners | Lactonase, 7-bladed beta-propeller |
| **K01271** | 21.25934296 | 1.491093306 | 1.21E-07 | 8.06E-07 | Cluster 3: L. iners | Creatinase/Prolidase N-terminal domain |
| **K18692** | 8.884234345 | 1.490516928 | 1.04E-05 | 4.05E-05 | Cluster 3: L. iners | DEAD-box RNA helicase. May work in conjunction with the cold shock proteins to ensure proper initiation of transcription at low and optimal temperatures |
| **K02243** | 6.669251197 | 1.487129601 | 1.67E-05 | 6.22E-05 | Cluster 3: L. iners | Type II IV secretion system protein |
| **K19005** | 25.66037065 | 1.477233589 | 3.95E-05 | 0.000135735 | Cluster 3: L. iners | Phosphoglycerol transferase and related proteins, alkaline phosphatase superfamily |
| **K00243** | 4.595943478 | 1.475765624 | 4.72E-07 | 2.60E-06 | Cluster 3: L. iners | S1 domain |
| **K07334** | 0.47115698 | 1.474574301 | 0.023453218 | 0.047049588 | Cluster 3: L. iners | RelE-like toxin of type II toxin-antitoxin system HigB |
| **K11071** | 4.634011068 | 1.474508196 | 3.10E-08 | 2.52E-07 | Cluster 3: L. iners | ABC transporter permease |
| **K02859** | 0.515566301 | 1.473928405 | 0.000710232 | 0.00196987 | Cluster 3: L. iners | acetyltransferase |
| **K06400** | 14.33115785 | 1.473741467 | 8.08E-08 | 5.67E-07 | Cluster 3: L. iners | ABC transporter |

| | | | | | | |
|---|---|---|---|---|---|---|
| **K02242** | 3.74454376 | 1.4708939 | 1.68E-06 | 7.98E-06 | Cluster 3: L. iners | Competence protein |
| **K14540** | 4.960445191 | 1.469164793 | 3.26E-09 | 4.11E-08 | Cluster 3: L. iners | Required for a late step of 50S ribosomal subunit assembly. Has GTPase activity |
| **K01232** | 10.37711977 | 1.448193586 | 0.000296095 | 0.000867608 | Cluster 3: L. iners | Family 4 glycosyl hydrolase C-terminal domain |
| **K01419** | 1.092890714 | 1.447340532 | 5.52E-05 | 0.000183759 | Cluster 3: L. iners | Protease subunit of a proteasome-like degradation complex believed to be a general protein degrading machinery |
| **K05362** | 12.36849385 | 1.44681268 | 1.81E-05 | 6.63E-05 | Cluster 3: L. iners | Catalyzes the addition of an amino acid to the nucleotide precursor UDP-N-acetylmuramoyl-L-alanyl-D-glutamate (UMAG) in the biosynthesis of bacterial cell-wall peptidoglycan |
| **K15780** | 10.64697878 | 1.43984172 | 1.35E-05 | 5.08E-05 | Cluster 3: L. iners | Ligates lysine onto the cytidine present at position 34 of the AUA codon-specific tRNA(Ile) that contains the anticodon CAU, in an ATP-dependent manner. Cytidine is converted to lysidine, thus changing the amino acid specificity of the tRNA from methionine to isoleucine |
| **K03758** | 3.155871084 | 1.436333289 | 0.001141783 | 0.003059233 | Cluster 3: L. iners | Arginine ornithine antiporter |
| **K01270** | 11.0010604 | 1.435051079 | 7.01E-11 | 1.44E-09 | Cluster 3: L. iners | dipeptidase PepV |
| **K01914** | 9.521528419 | 1.433427136 | 1.71E-09 | 2.42E-08 | Cluster 3: L. iners | aspartate--ammonia ligase |
| **K04771** | 13.60826185 | 1.430144352 | 3.23E-08 | 2.59E-07 | Cluster 3: L. iners | serine protease |

| | | | | | | |
|---|---|---|---|---|---|---|
| **K09787** | 0.674743053 | 1.429262654 | 0.000153188 | 0.00046908 | Cluster 3: L. iners | Might take part in the signal recognition particle (SRP) pathway. This is inferred from the conservation of its genetic proximity to ftsY ffh. May be a regulatory protein |
| **K03590** | 8.131005338 | 1.426898011 | 2.62E-06 | 1.18E-05 | Cluster 3: L. iners | Cell division protein that is involved in the assembly of the Z ring. May serve as a membrane anchor for the Z ring |
| **K21567** | 5.675260039 | 1.425396163 | 3.19E-05 | 0.000110908 | Cluster 3: L. iners | Ferredoxin--NADP reductase |
| **K03367** | 15.24068898 | 1.424876577 | 1.04E-05 | 4.05E-05 | Cluster 3: L. iners | Catalyzes the first step in the D-alanylation of lipoteichoic acid (LTA), the activation of D-alanine and its transfer onto the D-alanyl carrier protein (Dcp) DltC. In an ATP- dependent two-step reaction, forms a high energy D-alanyl-AMP intermediate, followed by transfer of the D-alanyl residue as a thiol ester to the phosphopantheinyl prosthetic group of the Dcp. D-alanylation of LTA plays an important role in modulating the properties of the cell wall in Gram-positive bacteria, influencing the net charge of the cell wall |
| **K02622** | 9.164146922 | 1.42006926 | 5.99E-07 | 3.18E-06 | Cluster 3: L. iners | Topoisomerase IV is essential for chromosome segregation. It relaxes supercoiled DNA. Performs the decatenation events required during the replication of a circular DNA molecule |
| **K03763** | 23.63473518 | 1.416538027 | 4.17E-09 | 4.96E-08 | Cluster 3: L. iners | Required for replicative DNA synthesis. This DNA polymerase also exhibits 3' to 5' exonuclease activity |
| **K03667** | 6.90195886 | 1.415123922 | 8.43E-06 | 3.32E-05 | Cluster 3: L. iners | this subunit has chaperone activity. The binding of ATP and its subsequent hydrolysis by HslU are essential for unfolding of protein substrates |

| | | | | | | |
|---|---|---|---|---|---|---|
| | | | | | | subsequently hydrolyzed by HslV. HslU recognizes the N-terminal part of its protein substrates and unfolds these before they are guided to HslV for hydrolysis |
| **K12573** | 17.77158641 | 1.408237578 | 3.85E-09 | 4.66E-08 | Cluster 3: L. iners | 3'-5' exoribonuclease that releases 5'-nucleoside monophosphates and is involved in maturation of structured RNAs |
| **K12132** | 94.87371343 | 1.408125877 | 2.99E-06 | 1.31E-05 | Cluster 3: L. iners | Protein tyrosine kinase |
| **K07027** | 5.222649675 | 1.407830137 | 2.32E-07 | 1.39E-06 | Cluster 3: L. iners | Catalyzes the transfer of a lysyl group from L-lysyl-tRNA(Lys) to membrane-bound phosphatidylglycerol (PG), which produces lysylphosphatidylglycerol (LPG), a major component of the bacterial membrane with a positive net charge. LPG synthesis contributes to bacterial virulence as it is involved in the resistance mechanism against cationic antimicrobial peptides (CAMP) produces by the host's immune system (defensins, cathelicidins) and by the competing microorganisms |
| **K03436** | 4.512387419 | 1.406818733 | 3.67E-06 | 1.58E-05 | Cluster 3: L. iners | DeoR C terminal sensor domain |
| **K08723** | 4.360018993 | 1.405267727 | 5.42E-06 | 2.25E-05 | Cluster 3: L. iners | Haloacid dehalogenase-like hydrolase |
| **K11145** | 1.032066761 | 1.404571412 | 3.96E-05 | 0.000135916 | Cluster 3: L. iners | Involved in correct processing of both the 5' and 3' ends of 23S rRNA precursor. Processes 30S rRNA precursor transcript even in absence of ribonuclease 3 (Rnc) |

| | | | | | | |
|---|---|---|---|---|---|---|
| **K00980** | 1.312149476 | 1.400380595 | 0.000452479 | 0.001293173 | Cluster 3: L. iners | Glycerol-3-phosphate cytidylyltransferase |
| **K20461** | 7.739937186 | 1.39913763 | 0.00258727 | 0.006450186 | Cluster 3: L. iners | ABC-2 family transporter protein |
| **K11072** | 8.593710685 | 1.389648377 | 6.58E-07 | 3.42E-06 | Cluster 3: L. iners | Part of the ABC transporter complex PotABCD involved in spermidine putrescine import. Responsible for energy coupling to the transport system |
| **K05592** | 22.30374025 | 1.389060242 | 3.00E-08 | 2.45E-07 | Cluster 3: L. iners | DEAD-box RNA helicase possibly involved in RNA degradation. Unwinds dsRNA in both 5'- and 3'-directions, has RNA- dependent ATPase activity |
| **K00950** | 3.098343549 | 1.386866783 | 8.85E-05 | 0.000282085 | Cluster 3: L. iners | 7,8-dihydro-6-hydroxymethylpterin-pyrophosphokinase (HPPK) |
| **K16899** | 17.39092069 | 1.383815545 | 2.33E-07 | 1.39E-06 | Cluster 3: L. iners | The heterodimer acts as both an ATP-dependent DNA helicase and an ATP-dependent, dual-direction single-stranded exonuclease. Recognizes the chi site generating a DNA molecule suitable for the initiation of homologous recombination. This subunit has 5' - 3' nuclease activity |
| **K03547** | 8.864910509 | 1.383655541 | 2.04E-10 | 3.87E-09 | Cluster 3: L. iners | Ser Thr phosphatase family protein |
| **K20460** | 3.699582248 | 1.383377067 | 0.002404455 | 0.006040237 | Cluster 3: L. iners | ABC-2 family transporter protein |
| **K11070** | 2.63400171 | 1.382125763 | 8.84E-07 | 4.42E-06 | Cluster 3: L. iners | ABC transporter permease |
| **K03650** | 12.14605872 | 1.370772537 | 1.12E-08 | 1.11E-07 | Cluster 3: L. iners | Exhibits a very high intrinsic GTPase hydrolysis rate. Involved in the addition of a carboxymethylaminomethyl (cmnm) group at the |

| | | | | | | |
|---|---|---|---|---|---|---|
| | | | | | | wobble position (U34) of certain tRNAs, forming tRNA- cmnm(5)s(2)U34 |
| **K03569** | 14.3618045 | 1.366273165 | 4.15E-09 | 4.96E-08 | Cluster 3: L. iners | cell shape determining protein MreB |
| **K02056** | 14.50608816 | 1.36117284 | 5.05E-07 | 2.76E-06 | Cluster 3: L. iners | ABC transporter |
| **K06023** | 6.133224497 | 1.358372957 | 1.45E-06 | 7.05E-06 | Cluster 3: L. iners | Catalyzes the ATP- as well as the pyrophosphate-dependent phosphorylation of a specific serine residue in HPr, a phosphocarrier protein of the phosphoenolpyruvate-dependent sugar phosphotransferase system (PTS). HprK P also catalyzes the pyrophosphate-producing, inorganic phosphate-dependent dephosphorylation (phosphorolysis) of seryl-phosphorylated HPr (P- Ser-HPr). The two antagonistic activities of HprK P are regulated by several intracellular metabolites, which change their concentration in response to the absence or presence of rapidly metabolisable carbon sources (glucose, fructose, etc.) in the growth medium. Therefore, by controlling the phosphorylation state of HPr, HPrK P is a sensor enzyme that plays a major role in the regulation of carbon metabolism and sugar transport it mediates carbon catabolite repression (CCR), and regulates PTS-catalyzed carbohydrate uptake and inducer exclusion |
| **K02621** | 11.99894835 | 1.357875099 | 6.19E-07 | 3.25E-06 | Cluster 3: L. iners | Topoisomerase IV is essential for chromosome segregation. It relaxes supercoiled DNA. Performs the decatenation events required during the replication of a circular DNA molecule |

| | | | | | | |
|---|---|---|---|---|---|---|
| **K03700** | 2.707226441 | 1.353965488 | 4.21E-05 | 0.000143538 | Cluster 3: L. iners | Endonuclease that resolves Holliday junction intermediates in genetic recombination. Cleaves mobile four-strand junctions by introducing symmetrical nicks in paired strands. Promotes annealing of linear ssDNA with homologous dsDNA. Required for DNA repair, homologous recombination and chromosome segregation |
| **K03346** | 9.332817546 | 1.353651875 | 6.73E-05 | 0.000218877 | Cluster 3: L. iners | Replication initiation and membrane attachment |
| **K07533** | 7.360791154 | 1.351077957 | 8.05E-06 | 3.19E-05 | Cluster 3: L. iners | Plays a major role in protein secretion by helping the post-translocational extracellular folding of several secreted proteins |
| **K00364** | 6.646808082 | 1.348962664 | 2.86E-06 | 1.26E-05 | Cluster 3: L. iners | Catalyzes the irreversible NADPH-dependent deamination of GMP to IMP. It functions in the conversion of nucleobase, nucleoside and nucleotide derivatives of G to A nucleotides, and in maintaining the intracellular balance of A and G nucleotides |
| **K07658** | 4.786333689 | 1.344161577 | 1.16E-07 | 7.83E-07 | Cluster 3: L. iners | Transcriptional regulatory protein, C-terminal domain protein |
| **K10040** | 5.733336888 | 1.342284555 | 6.99E-05 | 0.000225918 | Cluster 3: L. iners | ABC transporter permease |
| **K00687** | 11.22284617 | 1.339318413 | 0.000132837 | 0.000409305 | Cluster 3: L. iners | Penicillin-binding Protein |
| **K12553** | 11.22284617 | 1.339318413 | 0.000132837 | 0.000409305 | Cluster 3: L. iners | Penicillin-binding Protein |
| **K21465** | 11.22284617 | 1.339318413 | 0.000132837 | 0.000409305 | Cluster 3: L. iners | Penicillin-binding Protein |

| | | | | | | |
|---|---|---|---|---|---|---|
| **K21466** | 11.22284617 | 1.339318413 | 0.000132837 | 0.000409305 | Cluster 3: L. iners | Penicillin-binding Protein |
| **K16052** | 7.891620084 | 1.335907967 | 2.50E-05 | 8.89E-05 | Cluster 3: L. iners | mechanosensitive ion channel |
| **K03048** | 2.131286283 | 1.335054262 | 0.000274179 | 0.000810616 | Cluster 3: L. iners | Participates in both the initiation and recycling phases of transcription. In the presence of the delta subunit, RNAP displays an increased specificity of transcription, a decreased affinity for nucleic acids, and an increased efficiency of RNA synthesis because of enhanced recycling |
| **K01153** | 44.88715359 | 1.332484102 | 3.73E-07 | 2.11E-06 | Cluster 3: L. iners | Subunit R is required for both nuclease and ATPase activities, but not for modification |
| **K02057** | 15.06558459 | 1.329244712 | 1.19E-06 | 5.84E-06 | Cluster 3: L. iners | Belongs to the binding-protein-dependent transport system permease family |
| **K00557** | 9.579549794 | 1.328790092 | 8.05E-05 | 0.000257828 | Cluster 3: L. iners | Belongs to the class I-like SAM-binding methyltransferase superfamily. RNA M5U methyltransferase family |
| **K09685** | 5.812144578 | 1.32867304 | 5.61E-05 | 0.000186515 | Cluster 3: L. iners | pur operon repressor |
| **K03571** | 2.637030679 | 1.326451871 | 6.15E-07 | 3.24E-06 | Cluster 3: L. iners | rod shape-determining protein MreD |
| **K01809** | 9.385716896 | 1.325579678 | 3.98E-08 | 3.14E-07 | Cluster 3: L. iners | mannose-6-phosphate isomerase |
| **K00938** | 5.844510037 | 1.315006439 | 0.000118801 | 0.000373049 | Cluster 3: L. iners | phosphomevalonate kinase |

| | | | | | | |
|---|---|---|---|---|---|---|
| **K05878** | 8.314829778 | 1.312837034 | 5.41E-06 | 2.25E-05 | Cluster 3: L. iners | Dak1 domain |
| **K11102** | 1.487777704 | 1.309153844 | 0.005909484 | 0.013791127 | Cluster 3: L. iners | Belongs to the dicarboxylate amino acid cation symporter (DAACS) (TC 2.A.23) family |
| **K09698** | 12.93326556 | 1.308717779 | 7.02E-07 | 3.63E-06 | Cluster 3: L. iners | Catalyzes the attachment of glutamate to tRNA(Glu) in a two-step reaction glutamate is first activated by ATP to form Glu-AMP and then transferred to the acceptor end of tRNA(Glu) |
| **K08602** | 15.18658415 | 1.290897533 | 4.68E-05 | 0.000157171 | Cluster 3: L. iners | oligoendopeptidase F |
| **K03581** | 13.81424611 | 1.290645871 | 9.52E-08 | 6.59E-07 | Cluster 3: L. iners | DNA-dependent ATPase and ATP-dependent 5'-3' DNA helicase. Has no activity on blunt DNA or DNA with 3'-overhangs, requires at least 10 bases of 5'-ssDNA for helicase activity |
| **K00784** | 8.80320172 | 1.288631287 | 6.42E-06 | 2.58E-05 | Cluster 3: L. iners | Zinc phosphodiesterase, which displays some tRNA 3'- processing endonuclease activity. Probably involved in tRNA maturation, by removing a 3'-trailer from precursor tRNA |
| **K00857** | 2.50178102 | 1.284853874 | 2.05E-06 | 9.62E-06 | Cluster 3: L. iners | thymidine kinase |
| **K06948** | 7.941297451 | 1.284106963 | 0.00014609 | 0.000448719 | Cluster 3: L. iners | Ribosome biogenesis GTPase YqeH |
| **K14761** | 0.538260304 | 1.281464546 | 0.001674264 | 0.004344274 | Cluster 3: L. iners | S4 domain protein YaaA |
| **K16898** | 17.71435304 | 1.281013179 | 1.52E-07 | 9.78E-07 | Cluster 3: L. iners | ATP-dependent helicase nuclease subunit A |

| | | | | | | |
|---|---|---|---|---|---|---|
| **K01208** | 14.16343799 | 1.276020922 | 7.50E-07 | 3.83E-06 | Cluster 3: L. iners | Belongs to the glycosyl hydrolase 13 family |
| **K03315** | 10.61200347 | 1.272179088 | 9.62E-06 | 3.77E-05 | Cluster 3: L. iners | Na H antiporter NhaC |
| **K18940** | 9.580995972 | 1.270817046 | 0.001489972 | 0.003927169 | Cluster 3: L. iners | Histidine kinase |
| **K00558** | 16.52307058 | 1.26566062 | 0.000596682 | 0.001676149 | Cluster 3: L. iners | C-5 cytosine-specific DNA methylase |
| **K02746** | 4.441857167 | 1.264042223 | 2.02E-08 | 1.77E-07 | Cluster 3: L. iners | PTS system |
| **K01215** | 11.70236407 | 1.256527702 | 0.000343867 | 0.000994289 | Cluster 3: L. iners | Alpha amylase, catalytic domain protein |
| **K13038** | 10.29211921 | 1.254954264 | 2.17E-09 | 2.97E-08 | Cluster 3: L. iners | Catalyzes two steps in the biosynthesis of coenzyme A. In the first step cysteine is conjugated to 4'-phosphopantothenate to form 4-phosphopantothenoylcysteine, in the latter compound is decarboxylated to form 4'-phosphopantotheine |
| **K01926** | 3.407513316 | 1.251918651 | 1.86E-05 | 6.80E-05 | Cluster 3: L. iners | Modulates transcription in response to changes in cellular NADH NAD( ) redox state |
| **K01281** | 13.4619518 | 1.247190518 | 0.000542369 | 0.001532309 | Cluster 3: L. iners | Removes N-terminal dipeptides sequentially from polypeptides having unsubstituted N-termini provided that the penultimate residue is proline |
| **K06177** | 5.683321171 | 1.238069527 | 2.42E-07 | 1.44E-06 | Cluster 3: L. iners | Responsible for synthesis of pseudouridine from uracil |
| **K00783** | 4.70911208 | 1.232662456 | 2.43E-05 | 8.70E-05 | Cluster 3: L. iners | Specifically methylates the pseudouridine at position 1915 (m3Psi1915) in 23S rRNA |

| | | | | | | |
|---|---|---|---|---|---|---|
| **K09817** | 4.464966974 | 1.229939922 | 4.59E-08 | 3.48E-07 | Cluster 3: L. iners | ABC transporter |
| **K02794** | 10.03884962 | 1.225037166 | 1.70E-09 | 2.42E-08 | Cluster 3: L. iners | PTS system sorbose subfamily IIB component |
| **K11069** | 8.287427104 | 1.220687977 | 1.22E-05 | 4.66E-05 | Cluster 3: L. iners | ABC transporter |
| **K15051** | 14.58210311 | 1.21370636 | 0.000132735 | 0.000409305 | Cluster 3: L. iners | DNA RNA non-specific endonuclease |
| **K03572** | 13.57663936 | 1.211196763 | 2.85E-07 | 1.66E-06 | Cluster 3: L. iners | This protein is involved in the repair of mismatches in DNA. It is required for dam-dependent methyl-directed DNA mismatch repair. May act as a molecular matchmaker , a protein that promotes the formation of a stable complex between two or more DNA-binding proteins in an ATP-dependent manner without itself being part of a final effector complex |
| **K17074** | 14.31199026 | 1.21043014 | 0.000102828 | 0.000324962 | Cluster 3: L. iners | ABC transporter permease |
| **K18682** | 11.95843724 | 1.208418544 | 1.73E-07 | 1.09E-06 | Cluster 3: L. iners | Endoribonuclease that initiates mRNA decay |
| **K03739** | 9.482143303 | 1.205783555 | 0.000128061 | 0.000399583 | Cluster 3: L. iners | MBOAT, membrane-bound O-acyltransferase family |
| **K01791** | 9.006758456 | 1.20526512 | 2.85E-06 | 1.26E-05 | Cluster 3: L. iners | Belongs to the UDP-N-acetylglucosamine 2-epimerase family |
| **K00942** | 9.365483083 | 1.202712037 | 1.33E-09 | 1.98E-08 | Cluster 3: L. iners | Essential for recycling GMP and indirectly, cGMP |

| | | | | | | |
|---|---|---|---|---|---|---|
| **K00850** | 5.502189797 | 1.202454553 | 4.17E-08 | 3.22E-07 | Cluster 3: L. iners | Catalyzes the phosphorylation of D-fructose 6-phosphate to fructose 1,6-bisphosphate by ATP, the first committing step of glycolysis |
| **K07012** | 12.05554008 | 1.202150763 | 0.009329012 | 0.020953088 | Cluster 3: L. iners | CRISPR-associated helicase cas3 |
| **K07030** | 11.78290595 | 1.200562148 | 6.58E-07 | 3.42E-06 | Cluster 3: L. iners | DAK2 domain fusion protein YloV |
| **K05837** | 10.37474568 | 1.199221073 | 6.48E-08 | 4.66E-07 | Cluster 3: L. iners | Belongs to the SEDS family |
| **K11749** | 8.745765315 | 1.1979297 | 1.07E-08 | 1.08E-07 | Cluster 3: L. iners | zinc metalloprotease |
| **K00832** | 8.114941343 | 1.197714207 | 2.00E-05 | 7.26E-05 | Cluster 3: L. iners | Aminotransferase |
| **K11144** | 4.068023225 | 1.193821368 | 0.000114631 | 0.000360529 | Cluster 3: L. iners | Primosomal protein DnaI |
| **K07029** | 5.973729525 | 1.192361334 | 7.71E-05 | 0.000248515 | Cluster 3: L. iners | Lipid kinase |
| **K01071** | 3.396339053 | 1.192257696 | 0.000760195 | 0.002093721 | Cluster 3: L. iners | Acyl-ACP thioesterase |
| **K03495** | 17.29674884 | 1.192231664 | 2.79E-06 | 1.24E-05 | Cluster 3: L. iners | NAD-binding protein involved in the addition of a carboxymethylaminomethyl (cmnm) group at the wobble position (U34) of certain tRNAs, forming tRNA-cmnm(5)s(2)U34 |
| **K03458** | 24.19965812 | 1.186816684 | 8.59E-09 | 8.92E-08 | Cluster 3: L. iners | xanthine permease |

| | | | | | | |
|---|---|---|---|---|---|---|
| **K02793** | 9.25295486 | 1.181497988 | 1.74E-06 | 8.21E-06 | Cluster 3: L. iners | PTS system sorbose subfamily IIB component |
| **K22044** | 8.732196432 | 1.166003065 | 5.08E-05 | 0.000169942 | Cluster 3: L. iners | mechanosensitive ion channel |
| **K00566** | 9.519684678 | 1.157087193 | 5.98E-08 | 4.35E-07 | Cluster 3: L. iners | Catalyzes the 2-thiolation of uridine at the wobble position (U34) of tRNA, leading to the formation of s(2)U34 |
| **K02787** | 5.890225188 | 1.156273921 | 0.015038029 | 0.032233688 | Cluster 3: L. iners | Phosphotransferase system, EIIC |
| **K02788** | 5.890225188 | 1.156273921 | 0.015038029 | 0.032233688 | Cluster 3: L. iners | Phosphotransferase system, EIIC |
| **K03529** | 25.37835214 | 1.154742394 | 1.21E-07 | 8.06E-07 | Cluster 3: L. iners | Required for chromosome condensation and partitioning |
| **K03570** | 5.778257161 | 1.154698654 | 2.84E-08 | 2.35E-07 | Cluster 3: L. iners | Involved in formation and maintenance of cell shape |
| **K03555** | 16.83318496 | 1.151034525 | 3.97E-06 | 1.70E-05 | Cluster 3: L. iners | that it carries out the mismatch recognition step. This protein has a weak ATPase activity |
| **K06878** | 4.390396105 | 1.150874054 | 0.000816282 | 0.002245061 | Cluster 3: L. iners | Belongs to the phenylalanyl-tRNA synthetase beta subunit family. Type 1 subfamily |
| **K03481** | 7.01666334 | 1.150342962 | 0.003443016 | 0.008382257 | Cluster 3: L. iners | Helix-turn-helix domain, rpiR family |
| **K02687** | 6.092145287 | 1.144036711 | 2.53E-05 | 8.94E-05 | Cluster 3: L. iners | Ribosomal protein L11 methyltransferase |
| **K17076** | 4.955408514 | 1.143881056 | 1.10E-05 | 4.25E-05 | Cluster 3: L. iners | ABC transporter, ATP-binding protein |

| | | | | | | |
|---|---|---|---|---|---|---|
| **K13789** | 5.017554865 | 1.141901125 | 3.13E-06 | 1.36E-05 | Cluster 3: L. iners | Belongs to the FPP GGPP synthase family |
| **K10947** | 1.248775633 | 1.139163547 | 0.001697386 | 0.004398483 | Cluster 3: L. iners | Asp23 family, cell envelope-related function |
| **K02775** | 13.73217166 | 1.132416803 | 0.000339218 | 0.000983731 | Cluster 3: L. iners | PTS system sugar-specific permease component |
| **K01421** | 23.64799158 | 1.1260833 | 0.000843751 | 0.002314154 | Cluster 3: L. iners | YhgE Pip domain protein |
| **K21471** | 39.23252247 | 1.118693597 | 2.78E-06 | 1.24E-05 | Cluster 3: L. iners | Transposase |
| **K04487** | 16.47837827 | 1.117607076 | 4.10E-08 | 3.19E-07 | Cluster 3: L. iners | Aminotransferase class V |
| **K20375** | 6.186307253 | 1.114721618 | 0.013855721 | 0.029927144 | Cluster 3: L. iners | Transcriptional regulator |
| **K01838** | 2.657076438 | 1.113493399 | 0.000267759 | 0.000794016 | Cluster 3: L. iners | Haloacid dehalogenase-like hydrolase |
| **K07058** | 6.321333598 | 1.112691475 | 1.51E-05 | 5.68E-05 | Cluster 3: L. iners | Belongs to the UPF0761 family |
| **K09773** | 5.031493433 | 1.110977689 | 1.24E-06 | 6.04E-06 | Cluster 3: L. iners | Bifunctional serine threonine kinase and phosphorylase involved in the regulation of the pyruvate, phosphate dikinase (PPDK) by catalyzing its phosphorylation dephosphorylation |
| **K01916** | 4.732908018 | 1.105464226 | 4.11E-05 | 0.000140426 | Cluster 3: L. iners | Catalyzes the ATP-dependent amidation of deamido-NAD to form NAD. Uses ammonia as a nitrogen source |

| | | | | | | |
|---|---|---|---|---|---|---|
| **K16211** | 10.98401536 | 1.100931752 | 0.002566623 | 0.006423071 | Cluster 3: L. iners | Major Facilitator |
| **K02781** | 0.850032286 | 1.095638769 | 0.009934921 | 0.022162516 | Cluster 3: L. iners | PTS system glucitol/sorbitol-specific IIA component |
| **K02837** | 10.54619519 | 1.091683416 | 1.74E-05 | 6.41E-05 | Cluster 3: L. iners | Increases the formation of ribosomal termination complexes and stimulates activities of RF-1 and RF-2. It binds guanine nucleotides and has strong preference for UGA stop codons. It may interact directly with the ribosome. The stimulation of RF- 1 and RF-2 is significantly reduced by GTP and GDP, but not by GMP |
| **K06024** | 2.020440326 | 1.085029098 | 2.53E-05 | 8.94E-05 | Cluster 3: L. iners | Participates in chromosomal partition during cell division. May act via the formation of a condensin-like complex containing Smc and ScpA that pull DNA away from mid-cell into both cell halves |
| **K01104** | 25.32174901 | 1.082854011 | 1.19E-05 | 4.56E-05 | Cluster 3: L. iners | Tyrosine phosphatase family |
| **K05540** | 6.514097211 | 1.076674544 | 0.000171108 | 0.000519117 | Cluster 3: L. iners | Catalyzes the synthesis of 5,6-dihydrouridine (D), a modified base found in the D-loop of most tRNAs, via the reduction of the C5-C6 double bond in target uridines |
| **K07456** | 22.5743496 | 1.058009663 | 0.000223808 | 0.00066871 | Cluster 3: L. iners | Endonuclease that is involved in the suppression of homologous recombination and may therefore have a key role in the control of bacterial genetic diversity |
| **K04094** | 7.280089295 | 1.052159679 | 6.29E-06 | 2.55E-05 | Cluster 3: L. iners | Catalyzes the folate-dependent formation of 5-methyl-uridine at position 54 (M-5-U54) in all tRNAs |

| | | | | | | |
|---|---|---|---|---|---|---|
| **K00625** | 8.977918476 | 1.049939808 | 5.85E-06 | 2.41E-05 | Cluster 3: L. iners | phosphate acetyltransferase |
| **K07668** | 5.105630843 | 1.049760754 | 0.001005105 | 0.002704048 | Cluster 3: L. iners | response regulator |
| **K05946** | 3.203204888 | 1.047642176 | 0.001527683 | 0.004004872 | Cluster 3: L. iners | Catalyzes the conversion of GlcNAc-PP-undecaprenol into ManNAc-GlcNAc-PP-undecaprenol, the first committed lipid intermediate in the de novo synthesis of teichoic acid |
| **K01878** | 5.538742909 | 1.040994392 | 7.78E-06 | 3.09E-05 | Cluster 3: L. iners | glycyl-tRNA synthetase alpha subunit |
| **K12700** | 6.679462675 | 1.039738931 | 0.001217196 | 0.003248052 | Cluster 3: L. iners | Nucleoside |
| **K02760** | 0.593931771 | 1.037487613 | 0.024204332 | 0.048408664 | Cluster 3: L. iners | PTS system, Lactose/Cellobiose specific IIB subunit |
| **K07462** | 15.51832943 | 1.03181463 | 1.03E-06 | 5.13E-06 | Cluster 3: L. iners | Single-stranded-DNA-specific exonuclease RecJ |
| **K09747** | 0.908498495 | 1.028919867 | 0.004821883 | 0.011497888 | Cluster 3: L. iners | Binds to DNA and alters its conformation. May be involved in regulation of gene expression, nucleoid organization and DNA protection |
| **K01185** | 3.836786552 | 1.028831543 | 0.00035165 | 0.001015306 | Cluster 3: L. iners | mannosyl-glycoprotein |
| **K06967** | 2.297790359 | 1.028749674 | 0.000505198 | 0.001433453 | Cluster 3: L. iners | SAM-dependent methyltransferase |
| **K11068** | 3.156386751 | 1.018221321 | 6.24E-05 | 0.000205456 | Cluster 3: L. iners | protein, hemolysin III |

| | | | | | | |
|---|---|---|---|---|---|---|
| **K17073** | 15.52239176 | 1.008593024 | 6.50E-05 | 0.000212786 | Cluster 3: L. iners | ABC transporter permease |
| **K20107** | 2.565429697 | 1.001937354 | 0.009903471 | 0.022117378 | Cluster 3: L. iners | phosphoenolpyruvate-dependent sugar phosphotransferase system, EIIA 1 |
| **K20108** | 2.565429697 | 1.001937354 | 0.009903471 | 0.022117378 | Cluster 3: L. iners | phosphoenolpyruvate-dependent sugar phosphotransferase system, EIIA 1 |
| **K03500** | 9.58552401 | 1.00110345 | 2.80E-06 | 1.25E-05 | Cluster 3: L. iners | Specifically methylates the cytosine at position 967 (m5C967) of 16S rRNA |
| **K09816** | 6.736242353 | 1.000695871 | 2.22E-06 | 1.03E-05 | Cluster 3: L. iners | ABC 3 transport family |

**Supplemental Table 4. KEGG pathways significantly enriched across microbiome community states**

| Cluster | Pathway_ID | Pathway_Name | BaseMean | $Log_2$FoldChange | $Log_2$FC_SE | Wald_Statistic | p_value | BH_adjusted_p |
|---|---|---|---|---|---|---|---|---|
| Cluster 1: *L. crispatus* | 00906 | Carotenoid biosynthesis | 3.18466808 | 3.36865396 | 0.45176263 | 7.45669025 | 8.8723E-14 | 2.2269E-12 |
| Cluster 1: *L. crispatus* | 00120 | Primary bile acid biosynthesis | 2.4367897 | 2.58239467 | 0.39527107 | 6.53322454 | 6.4369E-11 | 8.9758E-10 |
| Cluster 1: *L. crispatus* | 00121 | Secondary bile acid biosynthesis | 2.4564403 | 2.56802884 | 0.37307252 | 6.88345758 | 5.8417E-12 | 9.7751E-11 |
| Cluster 1: *L. crispatus* | 05211 | Renal cell carcinoma | 5.49661114 | 1.71791848 | 0.28558726 | 6.01538915 | 1.7946E-09 | 1.3649E-08 |
| Cluster 1: *L. crispatus* | 04934 | Cushing syndrome | 5.5001708 | 1.71428144 | 0.28609397 | 5.99202222 | 2.0725E-09 | 1.53E-08 |
| Cluster 1: *L. crispatus* | 00740 | Riboflavin metabolism | 39.2367517 | 1.42741769 | 0.12214757 | 11.68601 | 1.5028E-31 | 3.7721E-29 |
| Cluster 1: *L. crispatus* | 00621 | Dioxin degradation | 0.4111713 | 1.39028262 | 0.53249802 | 2.61086909 | 0.00903125 | 0.01937472 |
| Cluster 1: *L. crispatus* | 00053 | Ascorbate and aldarate metabolism | 27.8223335 | 1.29675815 | 0.24955371 | 5.19630876 | 2.0328E-07 | 9.2772E-07 |
| Cluster 1: *L. crispatus* | 01501 | beta-Lactam resistance | 123.772147 | 1.19758536 | 0.10712976 | 11.17883 | 5.1768E-29 | 6.4969E-27 |
| Cluster 1: *L. crispatus* | 00333 | Prodigiosin biosynthesis | 5.63566039 | 1.07441485 | 0.25306339 | 4.24563524 | 2.1797E-05 | 7.5989E-05 |
| Cluster 1: *L. crispatus* | 04917 | Prolactin signaling pathway | 8.34007744 | 1.010474 | 0.25299902 | 3.99398388 | 6.4972E-05 | 0.00020173 |

| | | | | | | | | |
|---|---|---|---|---|---|---|---|---|
| Cluster 2: *Gardnerella* | 04973 | Carbohydrate digestion and absorption | 14.59372 | 2.92683121 | 0.31563919 | 9.27271172 | 1.8147E-20 | 1.6145E-19 |
| Cluster 2: *Gardnerella* | 00642 | Ethylbenzene degradation | 3.75326363 | 2.88161282 | 0.33162467 | 8.68938025 | 3.6442E-18 | 2.3505E-17 |
| Cluster 2: *Gardnerella* | 04974 | Protein digestion and absorption | 6.35348588 | 2.86976105 | 0.31153573 | 9.21165954 | 3.2112E-20 | 2.7616E-19 |
| Cluster 2: *Gardnerella* | 04640 | Hematopoietic cell lineage | 5.72087329 | 2.83359161 | 0.3308895 | 8.56355865 | 1.0944E-17 | 6.7225E-17 |
| Cluster 2: *Gardnerella* | 00592 | alpha-Linolenic acid metabolism | 4.09630148 | 2.76843932 | 0.31851652 | 8.69166628 | 3.5716E-18 | 2.3505E-17 |
| Cluster 2: *Gardnerella* | 04614 | Renin-angiotensin system | 6.20535234 | 2.7343776 | 0.30406491 | 8.99274304 | 2.4114E-19 | 1.8298E-18 |
| Cluster 2: *Gardnerella* | 00281 | Geraniol degradation | 4.45910547 | 2.72725855 | 0.31567664 | 8.63940575 | 5.6506E-18 | 3.5558E-17 |
| Cluster 2: *Gardnerella* | 05143 | African trypanosomiasis | 7.09886508 | 2.67995408 | 0.35023431 | 7.65188897 | 1.9805E-14 | 8.9643E-14 |
| Cluster 2: *Gardnerella* | 04216 | Ferroptosis | 15.8781035 | 2.66493952 | 0.27787147 | 9.59054737 | 8.7619E-22 | 9.419E-21 |
| Cluster 2: *Gardnerella* | 00440 | Phosphonate and phosphinate metabolism | 3.44360908 | 2.59729044 | 0.28498767 | 9.11369407 | 7.9624E-20 | 6.4197E-19 |
| Cluster 2: *Gardnerella* | 04910 | Insulin signaling pathway | 12.3150144 | 2.5725723 | 0.27947762 | 9.20493142 | 3.4189E-20 | 2.8454E-19 |
| Cluster 2: *Gardnerella* | 05142 | Chagas disease | 6.06626565 | 2.55200279 | 0.33747951 | 7.56194878 | 3.9707E-14 | 1.7364E-13 |

| | | | | | | | | |
|---|---|---|---|---|---|---|---|---|
| Cluster 2: *Gardnerella* | 04920 | Adipocytokine signaling pathway | 16.7371452 | 2.50418132 | 0.281609 | 8.8924052 | 5.98E-19 | 4.2857E-18 |
| Cluster 2: *Gardnerella* | 00332 | Carbapenem biosynthesis | 8.23724455 | 2.48934033 | 0.2830849 | 8.79361743 | 1.4482E-18 | 9.8325E-18 |
| Cluster 2: *Gardnerella* | 04714 | Thermogenesis | 17.7784231 | 2.34738879 | 0.28093918 | 8.35550538 | 6.5153E-17 | 3.7354E-16 |
| Cluster 2: *Gardnerella* | 04214 | Apoptosis - fly | 1.96658146 | 2.30146685 | 0.29004777 | 7.93478563 | 2.1086E-15 | 1.0462E-14 |
| Cluster 2: *Gardnerella* | 04213 | Longevity regulating pathway - multiple species | 12.4691392 | 2.08638117 | 0.27068861 | 7.70767999 | 1.2813E-14 | 5.9029E-14 |
| Cluster 2: *Gardnerella* | 04142 | Lysosome | 6.97005375 | 1.95682824 | 0.32871852 | 5.95289924 | 2.6343E-09 | 8.3908E-09 |
| Cluster 2: *Gardnerella* | 01055 | Biosynthesis of vancomycin group antibiotics | 4.69529311 | 1.91876048 | 0.27961002 | 6.86227361 | 6.7773E-12 | 2.548E-11 |
| Cluster 2: *Gardnerella* | 04622 | RIG-I-like receptor signaling pathway | 1.21640181 | 1.87262099 | 0.37645437 | 4.9743638 | 6.5462E-07 | 1.7593E-06 |
| Cluster 2: *Gardnerella* | 01051 | Biosynthesis of ansamycins | 8.27269809 | 1.8668228 | 0.25869298 | 7.21636445 | 5.3396E-13 | 2.2584E-12 |
| Cluster 2: *Gardnerella* | 04146 | Peroxisome | 24.9615393 | 1.82183508 | 0.16301435 | 11.1759182 | 5.3494E-29 | 9.8582E-28 |
| Cluster 2: *Gardnerella* | 00524 | Neomycin, kanamycin and gentamicin biosynthesis | 4.58229979 | 1.77699499 | 0.25678536 | 6.92015705 | 4.5114E-12 | 1.7372E-11 |
| Cluster 2: *Gardnerella* | 00400 | Phenylalanine, tyrosine and | 61.7435863 | 1.77418032 | 0.13993916 | 12.6782263 | 7.8079E-37 | 3.3574E-35 |

| | | | | | | | | |
|---|---|---|---|---|---|---|---|---|
| | | tryptophan biosynthesis | | | | | | |
| Cluster 2: *Gardnerella* | 04931 | Insulin resistance | 48.7133393 | 1.77347015 | 0.13666614 | 12.9766612 | 1.6596E-38 | 1.0704E-36 |
| Cluster 2: Gardnerella | 00061 | Fatty acid biosynthesis | 58.841614 | 1.77067791 | 0.17679524 | 10.015416 | 1.3041E-23 | 1.6823E-22 |
| Cluster 2: *Gardnerella* | 05418 | Fluid shear stress and atherosclerosis | 6.65455218 | 1.72799832 | 0.18529688 | 9.32556602 | 1.1039E-20 | 1.0172E-19 |
| Cluster 2: *Gardnerella* | 02026 | Biofilm formation - Escherichia coli | 28.7507006 | 1.72283941 | 0.14327283 | 12.0248859 | 2.6295E-33 | 6.1674E-32 |
| Cluster 2: *Gardnerella* | 04659 | Th17 cell differentiation | 1.25895679 | 1.67383829 | 0.37715657 | 4.43804628 | 9.0779E-06 | 2.1292E-05 |
| Cluster 2: *Gardnerella* | 04657 | IL-17 signaling pathway | 1.24299894 | 1.67009459 | 0.38262135 | 4.36487564 | 1.272E-05 | 2.93E-05 |
| Cluster 2: *Gardnerella* | 04914 | Progesterone-mediated oocyte maturation | 1.34800135 | 1.66657056 | 0.38202718 | 4.36243979 | 1.2862E-05 | 2.9366E-05 |
| Cluster 2: *Gardnerella* | 04915 | Estrogen signaling pathway | 1.32079493 | 1.66116162 | 0.38400036 | 4.3259377 | 1.5188E-05 | 3.4075E-05 |
| Cluster 2: *Gardnerella* | 00523 | Polyketide sugar unit biosynthesis | 10.2750187 | 1.65940215 | 0.26931229 | 6.16162791 | 7.2001E-10 | 2.4125E-09 |
| Cluster 2: *Gardnerella* | 04612 | Antigen processing and presentation | 1.25415585 | 1.63775552 | 0.3772025 | 4.34184689 | 1.4129E-05 | 3.1976E-05 |
| Cluster 2: *Gardnerella* | 05215 | Prostate cancer | 1.32599606 | 1.6085662 | 0.38968103 | 4.12790487 | 3.6608E-05 | 8.0726E-05 |

| | | | | | | | | |
|---|---|---|---|---|---|---|---|---|
| Cluster 2: *Gardnerella* | 00290 | Valine, leucine and isoleucine biosynthesis | 27.7769792 | 1.51179576 | 0.18589582 | 8.13248938 | 4.2056E-16 | 2.2144E-15 |
| Cluster 2: *Gardnerella* | 05120 | Epithelial cell signaling in Helicobacter pylori infection | 3.96279688 | 1.50931213 | 0.32509248 | 4.64271622 | 3.4386E-06 | 8.6132E-06 |
| Cluster 2: *Gardnerella* | 00340 | Histidine metabolism | 11.7929706 | 1.48171716 | 0.24462244 | 6.0571595 | 1.3855E-09 | 4.5247E-09 |
| Cluster 2: *Gardnerella* | 04217 | Necroptosis | 32.3207468 | 1.45481598 | 0.12092098 | 12.0311297 | 2.438E-33 | 6.1674E-32 |
| Cluster 2: *Gardnerella* | 00401 | Novobiocin biosynthesis | 29.3356654 | 1.37643721 | 0.14276328 | 9.64139498 | 5.3457E-22 | 5.9964E-21 |
| Cluster 2: *Gardnerella* | 00540 | Lipopolysaccharide biosynthesis | 5.71839504 | 1.35671655 | 0.45975857 | 2.95093263 | 0.00316816 | 0.00556044 |
| Cluster 2: *Gardnerella* | 01210 | 2-Oxocarboxylic acid metabolism | 59.3988669 | 1.35579052 | 0.15044084 | 9.01211758 | 2.0212E-19 | 1.5802E-18 |
| Cluster 2: *Gardnerella* | 04151 | PI3K-Akt signaling pathway | 2.56102472 | 1.29449227 | 0.35746151 | 3.62134725 | 0.00029307 | 0.00057282 |
| Cluster 2: *Gardnerella* | 01212 | Fatty acid metabolism | 76.0285092 | 1.28421318 | 0.11894909 | 10.796326 | 3.5825E-27 | 6.162E-26 |
| Cluster 2: *Gardnerella* | 01053 | Biosynthesis of siderophore group nonribosomal peptides | 2.80860351 | 1.24367561 | 0.48278792 | 2.57602885 | 0.00999423 | 0.01685302 |
| Cluster 2: *Gardnerella* | 05132 | Salmonella infection | 1.17541507 | 1.24312221 | 0.508943 | 2.44255685 | 0.01458363 | 0.02396546 |

| | | | | | | | | |
|---|---|---|---|---|---|---|---|---|
| Cluster 2: *Gardnerella* | 00860 | Porphyrin metabolism | 28.8863078 | 1.24146476 | 0.21573516 | 5.75457788 | 8.6859E-09 | 2.6678E-08 |
| Cluster 2: *Gardnerella* | 00410 | beta-Alanine metabolism | 6.17815796 | 1.21586267 | 0.32180186 | 3.77829597 | 0.00015791 | 0.00032299 |
| Cluster 2: *Gardnerella* | 00920 | Sulfur metabolism | 26.3276822 | 1.1728775 | 0.11845047 | 9.90183936 | 4.0869E-23 | 5.021E-22 |
| Cluster 2: *Gardnerella* | 00220 | Arginine biosynthesis | 56.0003663 | 1.13008259 | 0.09362187 | 12.070711 | 1.5083E-33 | 4.3237E-32 |
| Cluster 2: *Gardnerella* | 04621 | NOD-like receptor signaling pathway | 4.05409224 | 1.10479532 | 0.2309856 | 4.78296189 | 1.7273E-06 | 4.4123E-06 |
| Cluster 2: *Gardnerella* | 00525 | Acarbose and validamycin biosynthesis | 7.54809903 | 1.08820722 | 0.29149981 | 3.73313183 | 0.00018911 | 0.00038118 |
| Cluster 2: *Gardnerella* | 03008 | Ribosome biogenesis in eukaryotes | 7.01894001 | 1.08518912 | 0.15480409 | 7.01008025 | 2.3818E-12 | 9.3566E-12 |
| Cluster 2: *Gardnerella* | 00750 | Vitamin B6 metabolism | 17.0005946 | 1.03826934 | 0.10865004 | 9.55608773 | 1.2229E-21 | 1.2621E-20 |
| Cluster 2: *Gardnerella* | 00521 | Streptomycin biosynthesis | 36.5838406 | 1.03588053 | 0.09973195 | 10.3866467 | 2.852E-25 | 4.5989E-24 |
| Cluster 2: *Gardnerella* | 00903 | Limonene degradation | 2.24442594 | 1.02476804 | 0.41525272 | 2.46781776 | 0.01359395 | 0.0224823 |
| Cluster 2: *Gardnerella* | 00531 | Glycosaminoglycan degradation | 4.19429537 | 1.02361342 | 0.38428737 | 2.66366655 | 0.00772941 | 0.01329459 |
| Cluster 3: *L. iners* | 02040 | Flagellar assembly | 17.1920751 | 2.78002363 | 0.37968555 | 7.32191055 | 2.4447E-13 | 3.7281E-12 |

| | | | | | | | | |
|---|---|---|---|---|---|---|---|---|
| Cluster 3: *L. iners* | 02030 | Bacterial chemotaxis | 22.1807834 | 2.1107047 | 0.28526197 | 7.39918012 | 1.3703E-13 | 2.229E-12 |
| Cluster 3: *L. iners* | 00627 | Aminobenzoate degradation | 12.5548878 | 1.68132597 | 0.26392357 | 6.3705032 | 1.8841E-10 | 1.8389E-09 |
| Cluster 3: *L. iners* | 05206 | MicroRNAs in cancer | 12.5686187 | 1.40233069 | 0.28811628 | 4.86723871 | 1.1317E-06 | 4.6714E-06 |
| Cluster 3: *L. iners* | 00311 | Penicillin and cephalosporin biosynthesis | 7.14593753 | 1.29313161 | 0.29982636 | 4.31293503 | 1.611E-05 | 4.736E-05 |
| Cluster 3: *L. iners* | 00643 | Styrene degradation | 7.33291166 | 1.18843703 | 0.30777257 | 3.86141302 | 0.00011273 | 0.00028653 |
| Cluster 3: *L. iners* | 04122 | Sulfur relay system | 35.9259349 | 1.08024529 | 0.12751221 | 8.47170074 | 2.4184E-17 | 8.4297E-16 |
| Cluster 3: *L. iners* | 04152 | AMPK signaling pathway | 6.65181329 | 1.02862597 | 0.24024022 | 4.28165598 | 1.8551E-05 | 5.3886E-05 |
| Cluster 3: *L. iners* | 00511 | Other glycan degradation | 43.3408437 | 1.01618513 | 0.16131049 | 6.2995601 | 2.9849E-10 | 2.8012E-09 |
| Cluster 3: *L. iners* | 00940 | Phenylpropanoid biosynthesis | 7.94483784 | 1.00248065 | 0.36548778 | 2.7428568 | 0.00609072 | 0.01238447 |

Differentially abundant KEGG pathways were identified using DESeq2 in one-versus-rest comparisons for the *L. crispatus*-, *Gardnerella*-, and *L. iners*-predominated microbiome communities. Pathways with a Benjamini–Hochberg-adjusted $p<0.05$ and $\log_2$ fold change >1 were considered significantly enriched. The table includes the KEGG pathway identifier and name, base mean, $\log_2$ fold change, standard error, Wald statistic, raw $p$ value, and Benjamini–Hochberg-adjusted $p$ value.

**Supplemental Table 5. Taxa significantly correlated with core motility- and chemotaxis-associated KEGG Orthology (KO) identifiers**

| Taxon | n_core_motility_KOs | genes | mean_rho | max_rho | min_FDR |
|---|---|---|---|---|---|
| *Sphingomonas koreensis* | 16 | *cheA, cheB, flgE, flgK, flgL, flhA, fliA, fliD, fliF, fliG, fliI, fliK, fliM, fliP, fliS, motA* | 0.48134569 | 0.612975 | 4.56E-14 |
| *Alloscardovia omnicolens* | 16 | *cheA, cheB, flgE, flgK, flgL, flhA, fliA, fliD, fliF, fliG, fliI, fliK, fliM, fliP, fliS, motA* | 0.47812357 | 0.56945331 | 6.74E-12 |
| *Bifidobacterium breve* | 16 | *cheA, cheB, flgE, flgK, flgL, flhA, fliA, fliD, fliF, fliG, fliI, fliK, fliM, fliP, fliS, motA* | 0.46520428 | 0.5971344 | 3.06E-13 |
| *Corynebacterium glucuronolyticum* | 16 | *cheA, cheB, flgE, flgK, flgL, flhA, fliA, fliD, fliF, fliG, fliI, fliK, fliM, fliP, fliS, motA* | 0.38473185 | 0.47947378 | 2.74E-08 |
| *Streptococcus agalactiae* | 16 | *cheA, cheB, flgE, flgK, flgL, flhA, fliA, fliD, fliF, fliG, fliI, fliK, fliM, fliP, fliS, motA* | 0.31936912 | 0.37379977 | 3.23E-05 |
| *Amygdalobacter indicium* | 10 | *cheA, cheB, flgE, flgK, fliA, fliD, fliI, fliK, fliM, fliP* | 0.29053137 | 0.35006399 | 0.00011429 |
| *Lactobacillus acidophilus* | 8 | *cheB, flgL, fliA, fliG, fliK, fliM, fliS, motA* | 0.30992445 | 0.38833167 | 1.41E-05 |
| *Cutibacterium acnes* | 4 | *cheB, fliG, fliS, motA* | 0.26735748 | 0.30974329 | 0.00076602 |
| *Lactobacillus jensenii* | 2 | *fliG, fliS* | 0.28806236 | 0.30169285 | 0.00108352 |
| *Lactobacillus amylovorus* | 2 | *fliG, fliS* | 0.27495242 | 0.27796164 | 0.0028442 |
| *Fannyhessea vaginae* | 2 | *cheA, fliI* | 0.25814605 | 0.27017604 | 0.00382486 |
| *Gardnerella greenwoodii* | 1 | *cheA* | 0.29494838 | 0.29494838 | 0.00143833 |

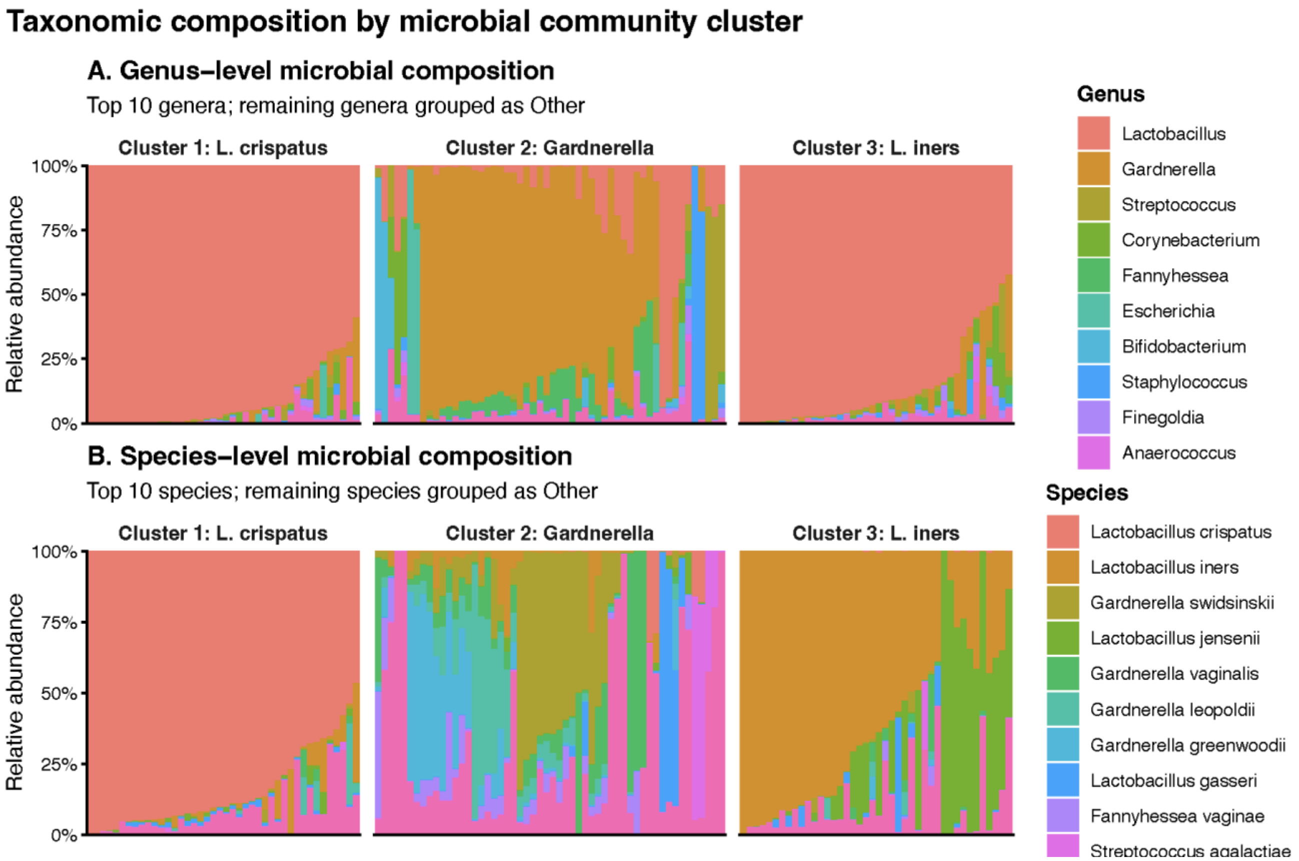


**Supplemental Figure 1. Taxonomic composition of the three urinary microbial community clusters at the genus and species levels.** Stacked bar plots show the relative abundance of microbial taxa across individual samples stratified by the three community clusters: Cluster 1 (*L. crispatus*-predominated), Cluster 2 (*Gardnerella*-predominated), and Cluster 3 (*L. iners*-predominated). (A) Genus-level composition showing the 10 most abundant genera. (B) Species-level composition showing the 10 most abundant species. Each vertical bar represents an individual's sample, and colors indicate the relative abundance of the corresponding taxon. Samples are displayed within each cluster according to their taxonomic composition. The genus-level profiles demonstrate the predominance of *Lactobacillus* in Clusters 1 and 3 and the greater contribution of *Gardnerella* and other taxa in Cluster 2, while species-level resolution distinguishes the *L. crispatus*- and *L. iners*-predominated community states and highlights the greater taxonomic heterogeneity of the *Gardnerella*-predominated cluster.
